# Combining small-molecule bioconjugation and hydrogen-deuterium exchange mass spectrometry (HDX-MS) to expose allostery: the case of human cytochrome P450 3A4

**DOI:** 10.1101/2020.06.09.142851

**Authors:** Julie Ducharme, Christopher J. Thibodeaux, Karine Auclair

## Abstract

We report herein a novel approach to study allostery which combines the use of carefully selected bioconjugates and hydrogen-deuterium exchange mass spectrometry (HDX-MS). The utility of our method is demonstrated using human cytochrome P450 3A4 (CYP3A4). CYP3A4 is arguably the most important drug-metabolizing enzyme, and as such, is involved in numerous drug interactions. Diverse allosteric ligand effects have been reported for this enzyme, yet the structural mechanism of these phenomena remain poorly understood. We have described different CYP3A4-effector bioconjugates, some of which mimic the allosteric effect of positive effectors on CYP3A4, while others show activity enhancement even though the label does not occupy the allosteric pocket (agonistic), or do not show activation while still blocking the allosteric site (antagonistic). These bioonjugates were studied here by HDX-MS, which enabled us to better define the position of the allosteric site, and to identify important regions involved in CYP3A4 activation.

## INTRODUCTION

Allostery is a ubiquitous biological phenomenon by which structural information is propagated spatially between distinct sites across the structure of macromolecular systems. Typically, an effector binds to an allosteric site, and this binding event modulates the functional activity at a remote site by triggering a reorganization of protein intramolecular interactions along a propagation path. Modern definitions of allostery consider proteins as dynamic conformational ensembles whose populations are altered by environmental changes, such as the binding of an effector.^1^ Despite its central role in biology, the molecular basis of allostery remains largely uncharacterized in most enzymes,^2,3^ hampering our ability to fully understand the biophysical mechanisms underlying both life and disease.

Human cytochrome P450 enzymes (CYPs) are heme monooxygenases that frequently exhibit atypical cooperative and allosteric behaviors.^4–6^ Among the 57 different human CYPs, CYP3A4 is the most abundant. This enzyme has attracted considerable interest for its contribution to the metabolism of approximately 50% of all clinical drugs^7^ and for its involvement in adverse drug interactions and drug resistance. The exceptionally broad substrate specificity of CYP3A4 is a result of its active site plasticity, which can even accommodate multiple ligands simultaneously.^6,8^ The kinetic behavior of CYP3A4 is further complicated by ligand binding at allosteric site(s).^9^ Whereas the allosteric modulation of CYP3A4 has been kinetically defined for several effectors,^10–12^ the structural mechanisms involved are challenging to characterize given the large number of possible enzyme-ligand(s) complexes and the possibility that ligands relocalize after initial binding.^13^ Furthermore, most known effectors of CYP3A4 also compete with other substrate molecules for oxidation.^13^ Importantly, the location of the allosteric site remains a matter of debate. It may overlap with the active site (Figure S10A) and even be dependent on the nature of the effector.^14,15^ All of these characteristics make deconvolution of the different ligand binding events, as well as the relationship of these events to allosteric activation and catalytic activity particularly challenging to investigate.

Hydrogen-deuterium exchange mass spectrometry (HDX-MS) is a powerful approach for characterizing protein conformational flexibility. This technique probes reorganization in the hydrogen-bond network of secondary structural elements by monitoring the time-dependent exchange of deuterium atoms between the solvent and the amide moities of the protein backbone.^16^ The ability of CYP3A4 to bind multiple ligands concurrently and the possible relocalisation of ligands in the enzyme may lead to heterogeneity in the system, thereby complicating HDX-MS data analysis and interpretation. In addition, the structural perturbations induced by transient ligand binding at an allosteric site may be difficult to capture on the time-scales typically employed in continuous exchange HDX-MS workflows. Furthermore, the current methods do not discriminate between dynamic changes implicated in ligand binding from those involved in rate enhancement. To overcome these intrinsic limitations, we report herein a novel approach that combines bioconjugation and HDX-MS. Covalently attaching the effector greatly reduces sample heterogeneity and noise, thereby enhancing our ability to capture the conformational changes that are specifically involved in enzyme activation.

Progesterone is a well-studied substrate and positive effector of CYP3A4. We recently demonstrated that, as expected for a selective binding process, the impact of bioconjugation of a maleimide-containing progesterone derivative (PGM) on enzyme kinetics varies greatly depending on the location of the attachment point, which affects binding orientation.^17,18^ The CYP3A4 F108C-PGM and F215C-PGM bioconjugates were found to effectively mimic progesterone allosteric activation towards 7-benzyloxy-4-trifluoromethylcoumarin (BFC) oxidation.^18^ We also constructed a PGM conjugate for which allosteric activation was antagonized (G481C-PGM), and a conjugate (L482C-PGM) which emulated activation without the PGM moiety occupying the allosteric site (agonist-like).^18^ Given the interesting differences in these functional effects, we reasoned that this collection of CYP3A4-PGM bioconjugates would enable a robust HDX-MS characterization of allosteric mechanisms in CYP3A4. The results presented herein reveal that, as expected, the conformational dynamics of CYP3A4 are significantly affected by the location of the covalently-tethered PGM ligand. Importantly, comparison of the HDX-MS data for all four CYP3A4 mutants before and after PGM bioconjugation, allowed us to not only confirm the location of the allosteric site, but also to identify critical CYP3A4 structural elements involved in allosteric activation, thereby demonstrating the utility of our approach.

## RESULTS

### Description of the HDX-MS workflow

Our study employs a typical continuous-labeling, bottom-up HDX-MS approach (Table S1). HDX was achieved by incubating enzymes in a buffered D_2_O solution (pD = 7.0, >98% deuterium) for different periods of time (0.5-240 min). During this continuous exchange, the amide moieties of the protein backbone acquire solvent deuteria at a rate that is highly sensitive to the local hydrogen bonding environment of that amide.^19^ Ligand binding (or PGM conjugation) can induce local and/or global fluctuations in the conformation of the protein, which may alter hydrogen bonding networks and, consequently, the rate of deuterium uptake at a given amide. Relative decreases in deuterium exchange typically reflect a structural organization or an intermolecular interaction with a ligand, whereas relative increases in deuterium exchange suggest enhanced flexibility of the protein backbone. Note that all of the CYP3A4 enzymes investigated in this work are truncated by 10 amino acids on the *N*-terminus. Truncation of this transmembrane α-helix is commonly employed to enhance CYP3A4 expression yields and solubility, and does not significantly impact enzymatic function under *in vitro* conditions (Figure S2 and S11B).^20,21^ In addition, many previous studies looking at CYP3A4 allostery in solution have employed similarly truncated variants,^22,23^ allowing direct comparison between our results and previous literature reports.

The extent of deuterium uptake for wild type CYP3A4 after 240 min of H/D exchange is shown in Figure 1A. Our workflow resulted in a total of 81 unique, reproducibly-detected peptic peptides spanning 88% of the CYP3A4 amino acid sequence. Overlapping peptides were identified across most of the protein sequence, providing further validation for the uptake data. A total of seven exchange time points were collected over a time regime spanning nearly four orders of magnitude (0.5-240 min).^16^ The deuterium uptake for each peptide over time can be visualized in Figure 1B, where the relative fractional deuterium uptake is plotted along the protein sequence. These data reveal that, for example, residues 150-270 undergo a more gradual deuterium uptake over time compared to other regions of the protein (*e*.*g*. residues 360-470), for which nearly complete exchange is achieved in 30 seconds. This data suggests that the *N*-terminus of CYP3A4 is generally more structured than the *C*-terminus. Interestingly, the most flexible regions (Figure 1C, colored in red) are located mostly at positions distal to the heme group, and include the B/C-loop, part of the D-helix, and the F-G region, as well as the J’-helix of the proximal side. The most rigid regions are found on the proximal side and in the direct vicinity of the heme, encompassing the *N*-terminal A-helices, the C, J-, K- and L-helices, and most of the active site residues.

**Figure 1.**
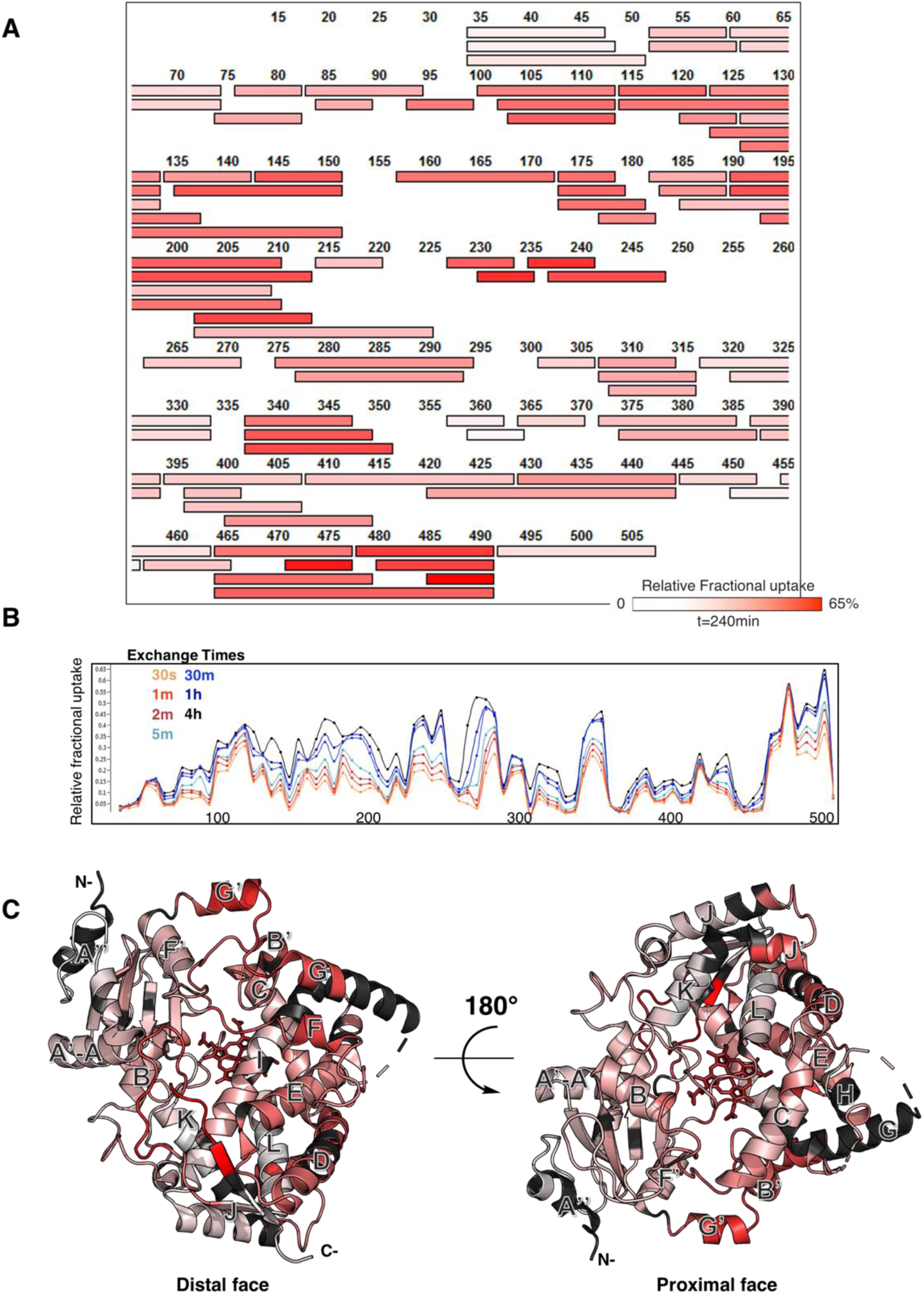
HDX profile of wild type CYP3A4. **A)** Coverage map after 240 min of exposure to D_2_O. The bars represent peptic peptides colored according to their relative deuterium uptake. The CYP3A4 amino acid sequence numbering is indicated. A total of 81 peptides was obtained providing a coverage of 88% and a redundancy of 2.26 peptides/amino acid. **B)** Deuterium relative fractional uptake (RFU) as a function of exchange time and amino acid sequence position for each detected peptide. The RFU represents the ratio of the measured deuterium uptake to the theoretical maximum uptake for a given peptide. This plot gives an overview of uptake rate across the CYP3A4 amino acid sequence **C)** Relative deuterium uptake (at 240 min) mapped onto the CYP3A4 crystal structure (PDB: 1W0F). A white-red color gradient is utilized, where white represents the regions undergoing little deteurium uptake over time (more structured) and red represents the more flexbile regions. The dark gray are regions not covered with HDX peptides

### Differential HDX profiles of the PGM-conjugates which best mimic allosteric activation highlight regions potentially involved in allosteric activation

To determine how PGM bioconjugation affects protein conformation, the HDX profiles were compared before and after PGM-bioconjugation (differential HDX) for the F108C and F215C mutants of CYP3A4. These bioconjugates have been previously shown to mimic progesterone allosteric activation.^14^ The relative fractional uptake (RFU) differences between the free F108C and F215C enzymes and their PGM-bioconjugates are shown in Figure 2. In this representation of the HDX data, regions of the protein shaded blue and red undergo less and more deuterium exchange, respectively, in the presence of the conjugated PGM moiety. The RFU differences measured in triplicate at each exchange time point were summed together, enabling us to capture more subtle changes in HDX with higher statistical confidence (all HDX difference data in this study are reported at the 98% confidence interval).^24^ Several regions of the F108C and F215C mutant enzymes exhibited similar changes in deuterium uptake upon PGM conjugation (Figure 3, S5, and S6). We reasoned that these regions would potentially be reporting on allosteric signaling networks shared by both the F108C and F215C enzymes. Namely, the first portion of the E-helix, the F’-helix, the G’-helix, the K/β1-loop, the β1-sheet, and part of the I-helix all undergo rigidification in both enzymes upon PGM labeling. In addition, the C and F helices become more flexible upon PGM conjugation to the enzyme. The rigidification of peptides in the F’-G’ helices, which comprise a portion of the previously proposed allosteric binding site,^25,22^ are consistent with binding of the PGM label at this location. Interestingly, both the β1-sheet and K/β1-loop (wich also rigidify upon PGM conjugation) are located in the active site, providing evidence that binding of PGM to the allosteric pocket may help to organize CYP3A4 elements that directly flank the redox-active heme. Although largely similar, the differential HDX profiles for these two mutants do show some minor differences, such as in the A’-A, G-helix, and D/E-loop which become more rigid in the F215C-PGM conjugate and more flexbile in the F108C-PGM conjugate. These minor disparities may be attributable to the different covalent PGM attachment sites, which may affect the PGM binding orientation in the allosteric pocket.

**Figure 2.**
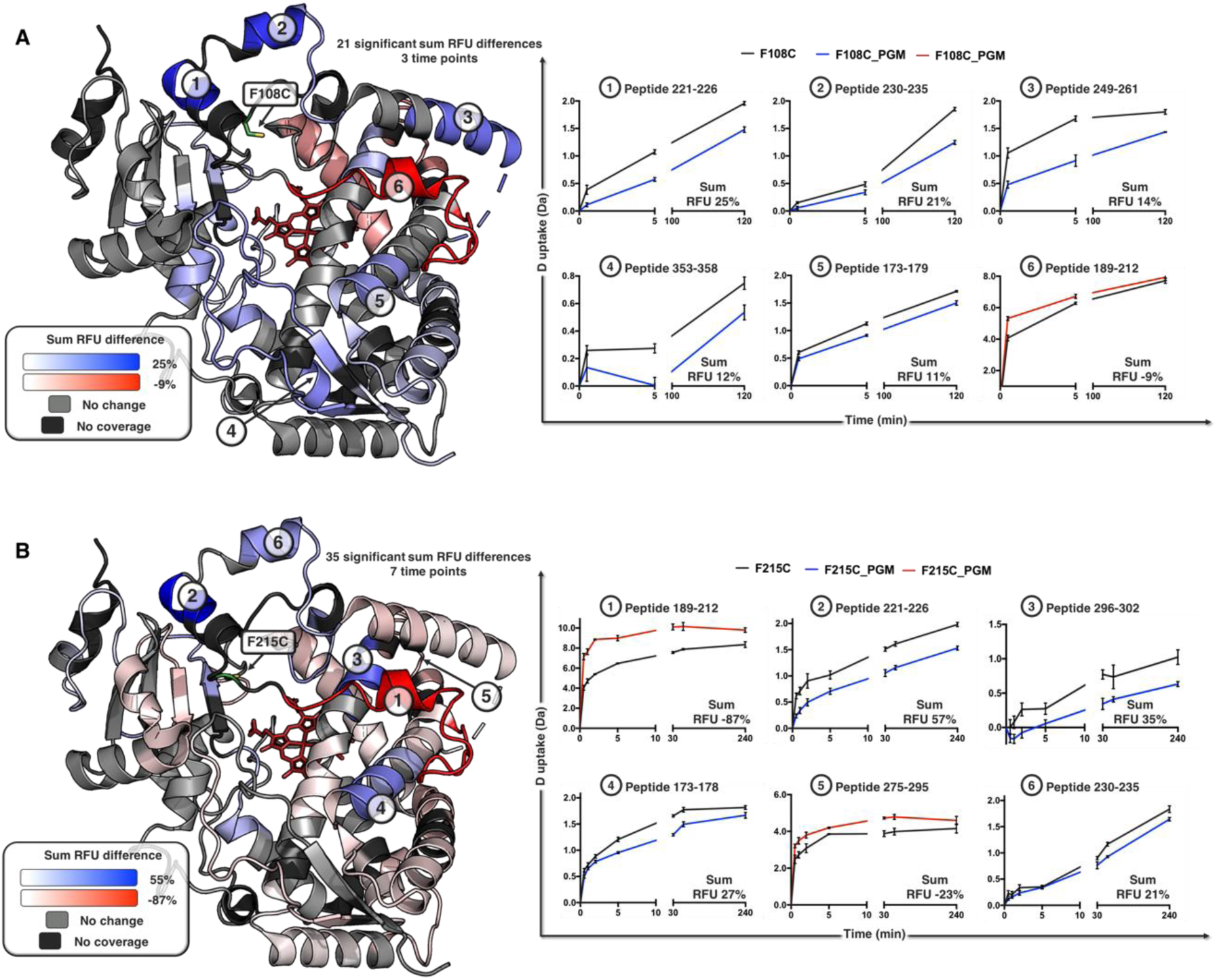
Differential HDX profiles of the allostery-mimicking PGM-bioconjugates. Changes in relative fractional uptake (RFU) of the CYP3A4 F108C (**A**) and CYP3A4 F215C (**B**) mutants upon PGM bioconjugation. The RFU of the bioconjugate was subtracted from the RFU of the free enzyme for each peptide at each exchange time point. The RFU difference for each peptide at each time point was then summed together to generate a final list of 21 and 35 peptides (for F108C and F215C, respectively) that showed a significant RFU change upon PGM conjugation (CI 98%). The significant peptides are mapped onto the CYP3A4 structure (PDB: 1W0F), where the blue and red colors indicate protein segments that exchange less and more deuterium, respectively, upon PGM conjugation. The light gray regions did not undergo a significant change in uptake upon PGM bioconjugation. The dark gray regions were not significantly covered by peptides derived from the proteolysis step. The cysteine bioconjugation handles can be found in the structure where the mutation number is shown. Deuterium uptake plots are shown for the six peptides that underwent the largest change in the summed RFU for each enzyme upon PGM conjugation. The summed RFU difference value is indicated at the bottom of each graph. A complete list of peptides that underwent significant H/D exchange upon PGM conjugation is provided in Figures S5 (F108C) and S6 (F215C).

**Figure 3.**
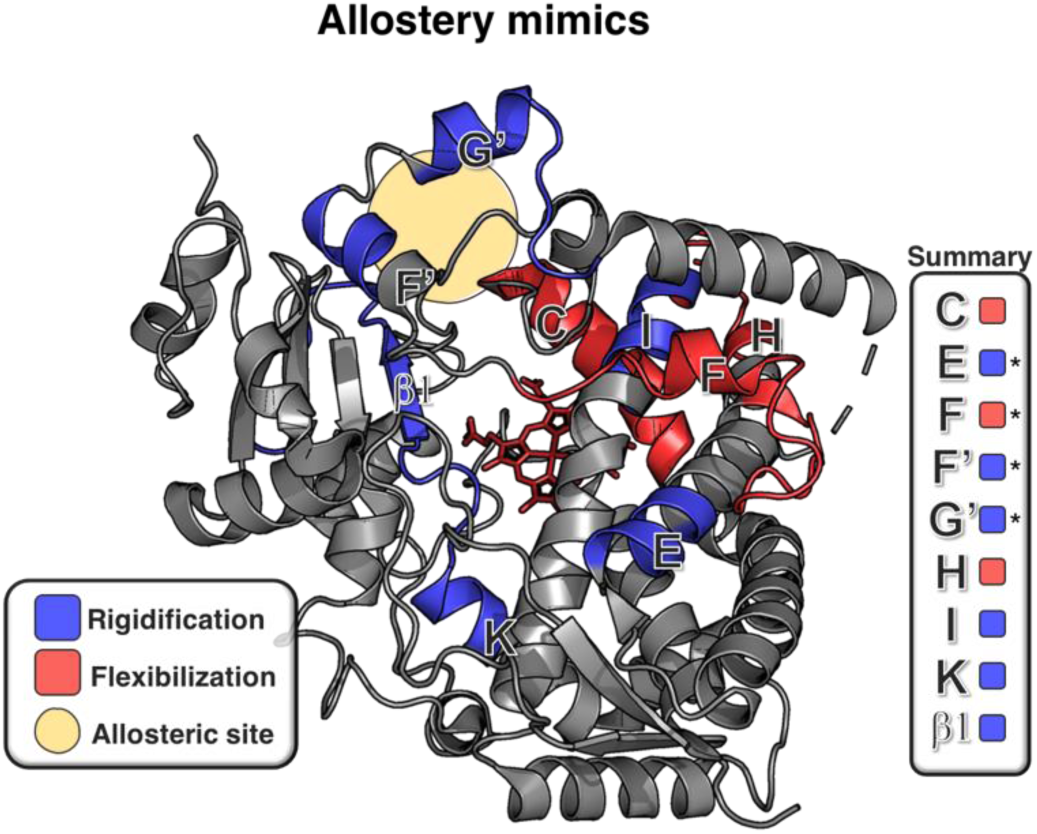
HDX profile differences that are common to the allostery-mimicking F108C-PGM and F215C-PGM systems. Regions highlighted in blue are, for both systems, becoming more rigid after PGM bioconjugation, whereas regions in red are becoming more flexible. A yellow circle designates the approximate position of the putative allosteric site of CYP3A4, which became more rigid upon PGM conjugation in both enzymes. A summary table of all the common regions affected by PGM is given on the right, with regions commonly showing the largest uptake difference identified by a star. These included the E-helix (peptide 173-178), F-helix (peptide 189-212), F’-helix (peptide 221-225) and G’-helix (peptide 230-235).

### Differential HDX of the antagonist- and agonist-mimicking PGM-bioconjugates help better define functionally relevant changes implicated in allosteric activation

To further confirm the conclusions reached using the allostery-mimicking PGM-conjugates, we next compared the HDX difference profiles of these enzymes with those of the antagonistic (G481C-PGM) and agonistic (L482C-PGM) bioconjugates. In the antagonistic system, the PGM moiety occupies the allosteric pocket and prevents allosteric activation by progesterone, yet does not activate the enzyme. As such, the G481C-PGM bioconjugate is expected to have a similar HDX profile as the allostery-mimicking systems for structural elements involved in binding the allosteric effector, but not for those elements implicated in activity enhancement. In contrast, the agonistic system is activated even though the PGM label does not occupy the progesterone allosteric pocket; this bioconjugate is further activated by progesterone. We therefore anticipate the HDX profile of L482C-PGM to be comparable to that of the allostery-mimicking systems for structural elements implicated in activity improvement, but not those associated with effector binding.

Comparison between the antagonistic (G481C) and the allostery-mimicking (F108C and F215C) enzymes may help us confirm the location of the allosteric site (from the HDX uptake patterns they share) and to identify conformational changes specific to allosteric activation (by identifying where their HDX uptake patterns differ). Our data show that the four most affected regions of CYP3A4 G481C (the F, F’, H-helices and the K/β1-loop) all become more rigid after PGM bioconjugation (Figure 4A, S3A, S7). Similar to the allostery-mimicking enzymes, rigidification of the F’-helix (peptide 221-226) is observed for the antagonistic system, consistent with PGM occupying the same putative allosteric site, but likely in an orientation that does not trigger activation. Interestingly, the F-helix (peptide 193-213), which experienced a drastic increase in flexibility in the allostery-mimicking bioconjugates, undergoes a significant rigidification in the antagonist bioconjugate. It is therefore tempting to speculate that flexibility in the F-helix is somehow linked to allosteric activation.

**Figure 4.**
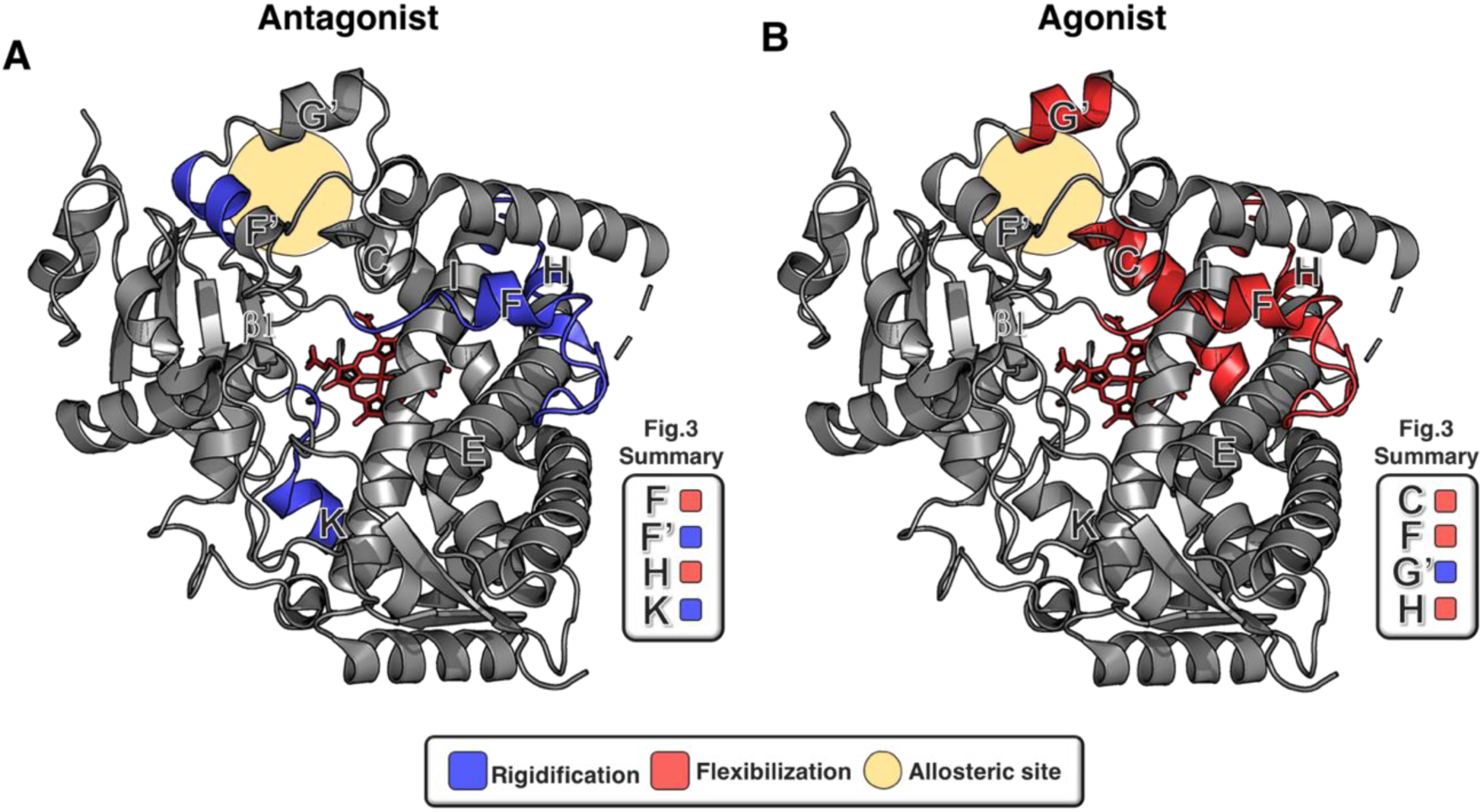
Common features of the differential HDX profiles for the antagonistic CYP3A4 G481C (**A**) and agonistic CYP3A4 L482C (**B**) PGM-bioconjugates, as compared to the allostery-mimicking F108C and F215C systems. Summary tables from Figure 3 are shown as a reminder of the deuterium uptake changes observed in critical structural elements of the allostery-mimicking systems. Regions highlighted in blue are becoming more rigid after PGM bioconjugation, while regions in red are becoming more flexible. A yellow circle designates the approximate position of the putative allosteric site of CYP3A4. The full differential HDX profiles for the antagonist and agonist systems are provided in Figure S3, S7, and S8.

In the agonistic (L482C) system, bioconjugation of PGM positively modulates enzyme activity but does not prevent progesterone from activating CYP3A4. Thus, the tethered PGM moiety in this variant likely occupies a binding site that is distinct from the progesterone allosteric site. Consistent with this claim, and in contrast to the allostery-mimicking and antagonistic systems, no rigidification is observed around the allosteric pocket (*e*.*g*. F’-helix) upon PGM conjugation to L482C. The regions of the L482C protein most affected by PGM bioconjugation all increase in flexibility (Figure 4B, S3B, S8), including the F-helix (peptide 189-212) that also became more flexible in the allostery-mimicking enzymes. Thus, the HDX-MS data for the agonistic system further suggests that flexibility in the F-helix may be important to allosteric activation of the enzyme.

### PGM bioconjugation is a powerful tool to capture progesterone-induced allosteric conformational changes

At concentrations below 25 μM, progesterone positively modulates the activity of CYP3A4 towards several substrates (*e*.*g*. BFC, carbamazepine, nevirapine).^18,26,27^ Above this concentration, progesterone was reported to compete with the substrate for oxidation.^18,28^ If free progesterone is binding to the allosteric site at concentrations less than 25 μm, we reasoned that the differential HDX profile of unlabeled CYP3A4 F215C (known to be activated by progesterone) in the presence and absence of progesterone (20 μM) should resemble the HDX profiles of the allostery-mimicking bioconjugates. Surprisingly, progesterone binding to F215C resulted in H/D uptake behavior that contrasted with the F215C-PGM conjugate in several critical regions of the protein (Figure 5, Figure S9). Namely, peptides derived from the E-helix, G’-helix and K/β1-loop show an increase in flexibility in the presence of progesterone, instead of becoming more rigid as in the allostery-mimicking bioconjugates. In addition, the F-helix region (peptide 189-212) became more rigid in the presence of progesterone, contrasting with the flexibility induced in this region by PGM conjugation. Interestingly, the F’ and G’-helices of the allosteric site did not undergo significant changes in deuterium uptake in the presence of 20 μM progesterone (as seen for the allostery-mimicking bioconjugates), suggesting that progesterone may not bind very tightly to this region under our conditions. In the absence of another substrate for oxidation (*e*.*g*. BFC), it appears that progesterone may not have a long residence time in the allosteric pocket, and may relocate to the active site (Figure 5B). Cumulatively, these interesting data suggest that the conformational changes triggered by PGM binding to the allosteric site differ from the conformational changes associated with binding and oxidation of free progesterone. This result highlights the utility of the bioconjugation approach, which proved necessary in this case to reveal the structural perturbations associated specifically with allosteric activation.

**Figure 5.**
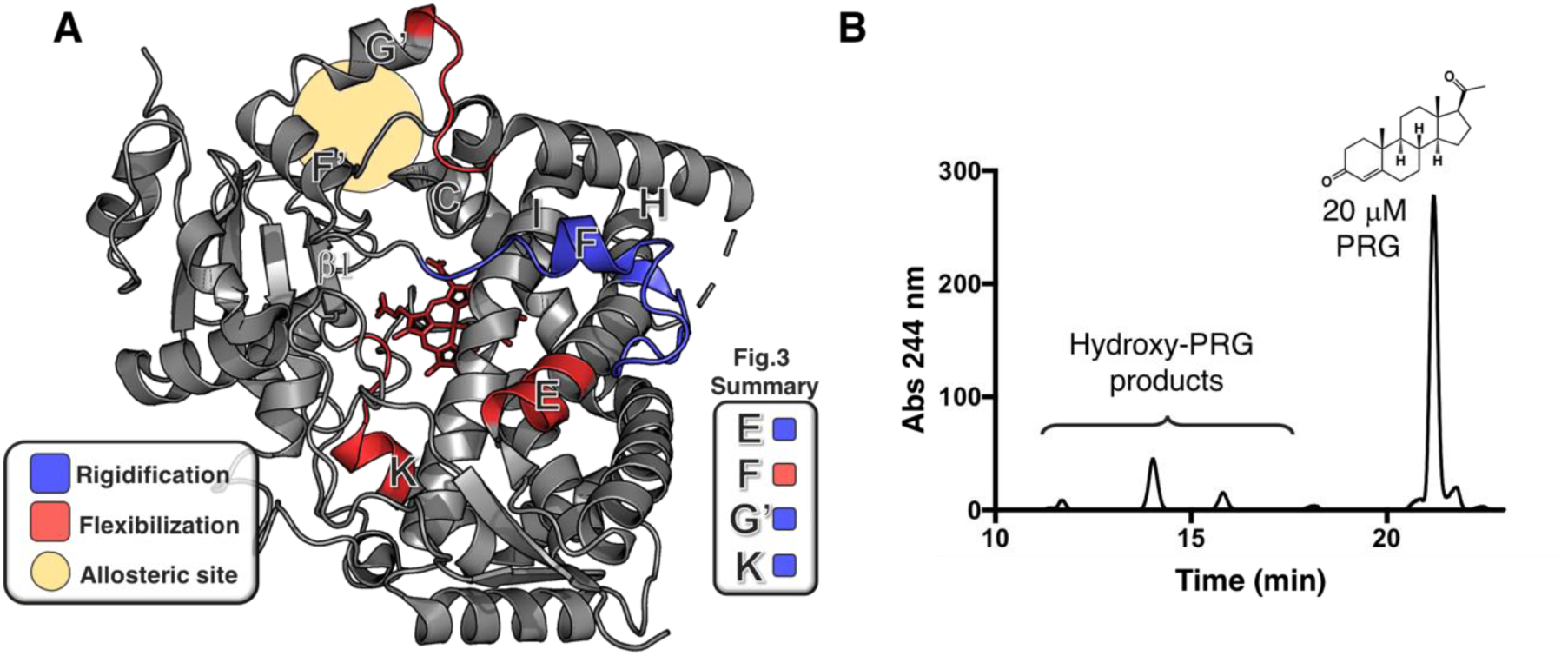
Common features of the differential HDX profiles of the CYP3A4 F215C mutant in the presence of progesterone (PRG), as compared to the allostery-mimicking systems. **A)** Regions highlighted in blue are becoming more rigid in presence of progesterone, while regions in red are becoming more flexible. A yellow circle designates the position of the putative allosteric site of CYP3A4. A summary table from Figure 3 is shown as a reminder of the deuterium uptake changes observed in critical structural elements of the F215C-PGM and F108C-PGM allostery-mimicking systems. **B)** HPLC-based activity assay of the CYP3A4 F215C mutant in the presence of 20 μM progesterone, a concentration at which progesterone behaves as a positive effector in the presence of other substrates. The elution of hydroxylated progesterone products between 12-16 min indicates that progesterone must also be binding to the active site and potentially serving as a substrate. This relocation of PRG to the active site could be responsible for why the differential HDX profile of [F215C + PRG] differs from the profile of the F215C-PGM bioconjugate (Figure 2). The full differential HDX profiles for [F215C + PRG] and F215C-PGM are shown in Figures S9 and S4, respectively.

## DISCUSSION

Allostery is a challenging phenomenon to study, especially when multiple ligands bind simultaneously and lead to atypical enzyme kinetics.^4^ The CYP3A4 system is further complicated by the probable overlap between the allosteric and active sites, the dual effector/substrate activity of several ligands, and the possible relocalization of bound ligands to the active site. Important insight into the location of the CYP3A4 allosteric site came from a crystal structure in which the substrate progesterone was found to bind at a site distal to the heme, near the F’-helix (Figure 6A).^25^ Although it has been speculated that this peripheral binding site may stem from a crystallographic artifact, many biochemical and biophysical studies support the hypothesis that the F’-helix is involved in effector binding and allosteric activation.^14,22,23,26,29–35^ How a binding event in this region may lead to CYP3A4 catalytic enhancement remains unknown. Structural changes leading to allosteric activation are often dynamic, and can be transient, making them difficult to fully capture with X-ray crystallography. Indeed, comparing the structures of the substrate-free CYP3A4 (PDB: 1W0E, 1TQN)^25,36^ to the progesterone-bound complex (PDB: 1W0F, 5A1P),^25,37^ reveals no drastic reorganization, even though the biochemical evidence for allosteric activation is strong.

**Figure 6.**
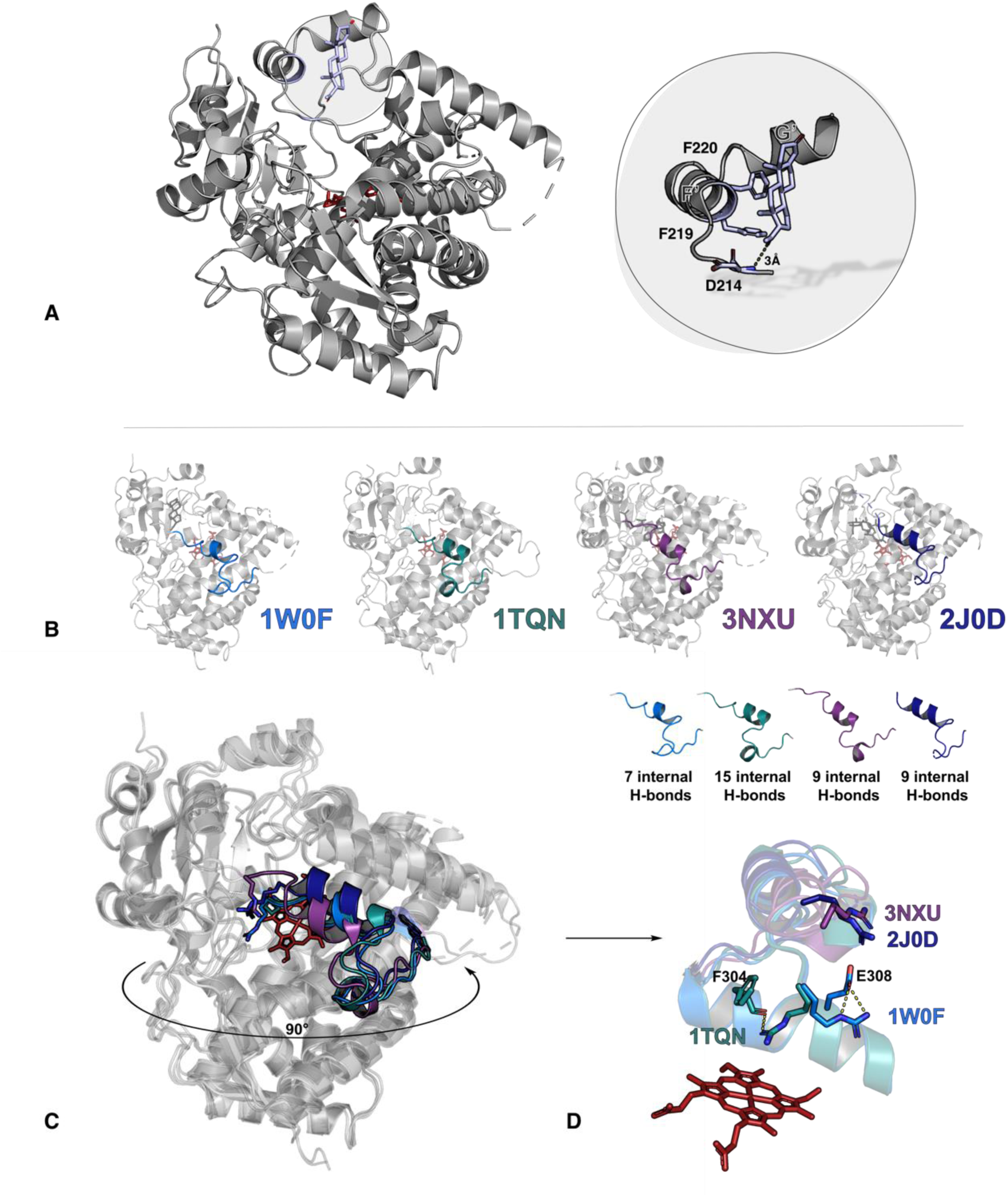
Important structural features of CYP3A4 highlighted in various crystal structures. **A**) Published crystal structure of CYP3A4 in complex with progesterone (PDB: 1W0F). Progesterone interacts with residues D214, F219 and F220 (light blue) of the F’ helix, approximately 18 Å away from heme iron. The peripheral site to which progesterone is bound is postulated to be an allosteric site. **B)** The F-helix adopts different conformations when ligands bind at the active site. CYP3A4 structures 1W0F (progesterone-bound), 1TQN (substrate-free), 3NXU (ritonavir-bound) and 2J0D (erythromycin-bound), all showing different conformations of the F-helical region. This reorganization affects the total number of internal hydrogen bonds holding the secondary structure in place. Changes in such intramolecular hydrogen-bonding are likely to be detected by HDX-MS measurements. **C)** All four structures from (A) superimposed in order to expose the observed range of F-helix conformations. **D)** Arg212, located on the F/F’-loop, reorients greatly upon restructuring of the F-helix. In the progesterone-bound (1W0F) and substrate-free structures (1TQN), the F/F’-loop closes in towards the I-helix where Arg212 forms salt bridges with Phe304 and Glu308. In the ritonavir (3NXU) and erythromycin-bound structures (2J0D), the F/F’-loop has moved away from the active site and the Arg212 side chain is no longer in contact with the I-helix.

In previous work,^18^ we developed a set of functionally distinct CYP3A4-PGM bioconjugates that exhibited a range of kinetic behaviors. Interestingly, with the CYP3A4 F108C and F215C mutants, PGM bioconjugation was found to mimic progesterone allostery and prevent further activation by progesterone, consistent with the colvalently tethered PGM moiety occupying the allosteric site and increasing the enzyme activity. In contrast, PGM bioconjugation to the CYP3A4 G481C mutant led to an antagonistic behavior, where PGM occupies the allosteric site (prevents progesterone from activating the enzyme) but does not itself activate the enzyme. Finally, conjugation of PGM to the CYP3A4 L482C mutant resulted in an enzyme with agonistic behavior, where the enzyme was activated but the allosteric site remained available for progesterone binding. Together, these bioconjugates provide a unique toolbox to study the underlying mechanisms of allostery in CYP3A4. Namely, these modified enzymes enabled us to distinguish structural effects relevant to allosteric enzyme activation from those that reflect ligand binding or bioconjugation. The covalent attachment of progesterone also proved essential to prevent relocalization of the effector ligand from the allosteric site to the active site.

To date, several HDX-MS studies have looked at the dynamic properties of CYPs. These include investigations of active site plasticity (P450cam, CYP2B4),^38,39^ the impact of ligand binding (CYP46A1, CYP2B4),^40,41^ and allosteric site mapping (CYP46A1).^42^ To our knowledge, only two HDX-MS studies have looked specifically at CYP3A4, one investigating membrane association and inhibitor binding,^38^ and the other examining the binding of the substrate and the positive effector, midazolam.^43,44^ Herein, using CYP3A4 as a model system, we demonstrate the utility of combining bioconjugation and HDX-MS to characterize allosteric mechanisms. The results presented also advance our mechanistic understanding of CYP3A4 in several ways: i) they confirm the existence of functionally relevant flexible regions in wild type CYP3A4; ii) confirm that the F’-helix of CYP3A4 is part of the allosteric site; iii) identify coupled structural elements involved in CYP3A4 allosteric activation; and iv) demonstrate the importance of F-helix flexibility in CYP3A4 allosteric activation.

The most dynamic regions identified herein for wild type CYP3A4 comprise the B/C-loop, the F-G region (excluding the F’-helix), and the C-terminal-loop (Figure 1C). These regions were previously identified by Ekroos *et al*. as showing the highest variation in Cα RSMD based on alignment of the available crystal structures of CYP3A4.^45^ In their recent HDX-MS study of CYP3A4, Treuheit *et al*. also identified the F-G region, B/C-loop and C-terminus as the most flexible areas of the enzyme.^43^ These regions constitute the lining of the active site, and their motion is likely responsible for the well-known active site plasticity and relaxed substrate specificity of CYPs. Interestingly, all of these regions possess high sequence variability in different CYPs, consistent with the distinct substrate specificity profiles of these enzymes.^46,47^ These dynamic regions include four of the six characterized CYP substrate recognition sites (SRSs): SRS-1 (B/C-loop), SRS-2 (F/F’-loop), SRS-3 (G’/G-loop) and SRS-6 (C-terminus). SRS-4 and SRS-5 are located deeper in the active site (Figure S10B), and were found to be more protected from deuterium exchange in our study. Interestingly, the dynamic B/C-loop (SRS-1) separates two main substrate access channels, previously termed 2a and 2e (Figure S12A),^47^ and the motion of the B/C loop may modulate substrate access and/or product egress via these routes.^47^ A phenylalanine cluster (comprised of Phe108, Phe219, Phe220, Phe241 and Phe304) is found in the vicinity of the B/C-loop and F-G region, and is thought to contribute to hydrophobic packing that closes the active site upon ligand binding (Figure S11A).^48,25^

The short F-helix is an unusual structural characteristic of CYP3A4 and contains residues implicated in catalysis.^49–51^ Dynamic motions within the F-helix are therefore likely to affect activity. A key residue of the F/F’ loop is Arg212,^36^ which in the absence of ligand points towards the active site and interacts with the I-helix. Arg212 easily reorients upon ligand binding, owing to the flexibility of this region. In 3D structures of both substrate-free (PDB: 1TQN)^36^ and progesterone-bound (PDB:1W0F) CYP3A4,^25^ the Arg212 side chain is pointing towards the active site, where it interacts with I-helix residues Phe304 and Glu308, respectively. Conversely, in structures of the inhibitor-bound (PDB: 3NXU)^52^ and a substrate-bound form (PDB: 2J0D) of CYP3A4,^45^ the Arg212 side chain points away from the active site and the I-helix (Figure 6C,D). As noted by Benkaidali *et al*,^53^ movement of the F/F’-loop is also involved in the transition of CYP3A4 from the closed to the open state. Interestingly, in our HDX results with wild type CYP3A4, the F’-helix, which is part of the putative allosteric site, is the least dynamic portion of the F-G region. Its higher level of organization is likely important for effector binding, as further supported by our results with CYP3A4 bioconjugates.

To determine the impact of PGM bioconjugation on the conformational dynamics of CYP3A4, HDX-MS profiles of both unconjugated and PGM-conjugated mutants were compared. Different regions of the protein partially gained or lost their ability to uptake deuterium upon bioconjugation, suggesting that PGM indeed triggers changes in CYP3A4 conformational dynamics. Some of these dynamic perturbations may reflect structural changes involved in allostery, while others may result from the covalent modification itself. Therefore, not all deuterium uptake changes observed upon bioconjugation are necessarily relevant to allostery or are strictly associated with ligand binding. Focusing on the similarities in the deuterium uptake differences of two different allostery-mimicking bioconjugates (F108C and F215C) enabled us to assign dynamic changes induced specifically by PGM binding to the allosteric site (Figure 3). These regions include the C-, F-, and H-helices (which become more flexible), and the E-, F’-G’-, I-, K-helices (which become more rigid). Some of these elements are within the most highly dynamic regions of the ligand-free wild type CYP3A4 (Figure 1), suggesting that their flexibility may be required to sample allosterically-activated enzyme conformations.

We also compared the deuterium uptake profiles of these allosteric mimics with CYP3A4-PGM conjugates that exhibit antagonistic and agonistic behavior. The antagonistic bioconjugate (G481C mutant) and the allostery-mimicking (L482C) systems exhibited similar RFU changes in the F’, H- and K-helices upon PGM conjugation. These HDX perturbations likely reflect binding interactions between PGM and the enzyme, rather than an allosteric process that results in enzyme activation. On the other hand, comparison of the agonistic bioconjugate (L482C mutant) with the allostery-mimicking sytems enabled us to identify linked elements (the C-, F-, G’ and H-helices) with shared H/D uptake patterns that may report directly on allosteric enzyme activation (Figure 4). In particular, the largest change in deuterium uptake in the allostery-mimicking bioconjugates occurs in the F-helix, a region which shows highly variable conformations between the different crystal structures of CYP3A4 (Figure 6B-D). The F-helix also becomes more flexible in the agonist mimic (Figure 4B) but not in the antagonist mimic (Figure 4A). These data suggest that flexibility in the F-helix, triggered by effector binding, may be correlated with allosteric activation. Reorganization of the F-helix may involve a lost interaction between Arg212 and Glu308 in the I-helix, and a subsequent reorientation of Arg212 away from Glu308. This movement could facilitate proton transfer by the adjacent Thr309 residue in the I-helix, which is essential to oxygen activation by the heme.^25,54–57^ In addition to the F-helix, the C-helix also becomes very flexible in all of the activated bioconjugate enzymes, but not in the antagonist system. Noticeably, the C-helix is located below the entrance of the putative product egress channel 2e and is also involved in binding of the redox partner. Its conformational motions may thus play a role in the trafficking of small molecules and/or in the interaction with the cytochrome P450 reductase.^47,58^

The HDX-MS studies also provided insight into the effector binding site. Differential HDX profiles for the allostery-mimicking and antagonist systems revealed a rigidification of the F’-helix, consistent with binding of the covalently-tethered PGM label to the allosteric site deduced originally from X-ray crystal strcutures.^25^ As expected, organization of the F’-helix in the allosteric site was not observed for the agonistic L482C system, consistent with the available progesterone binding site in this conjugate suggested by earlier kinetic studies.^18^ In addition, rigidification of the K/β1-loop in the active site was observed only in the allostery-mimicking and antagonist bioconjugates, but not in the agonist conjugate, implying that K/β1-loop rigidification may be the result of coupled conformational changes between the F’-helix and the active site. Being near the heme prosthetic group, the K/β1-loop (SRS-5) is expected to participate in substrate orientation during catalysis.^59^ Residue Arg375 on the β1-sheet directly interacts with one of the heme propionate groups and is also believed to be involved in water channel gating (Figure S12B).^60^ Reorganization of the K/β1-loop region (triggered by effector binding to the allosteric site) may organize the substrate binding pocket in the active site, and impact the circulation of water molecules into and out of the active site. Somewhat surprisingly, we were unable to observe structural organization of the F’-helix and the allosteric site of the unmodified F215C enzyme in the presence of the effector progesterone. We attribute this lack of an HDX perturbation to weak binding of progesterone to the allosteric site – a finding that validates the effectiveness of our covalent conjugation strategy to reveal allosteric mechanisms.

## CONCLUSION

In summary, we have demonstrated that combining bioconjugation and HDX-MS is a powerful strategy to study the underlying mechanisms of allostery. This unique approach allowed us to overcome issues related to ligand relocalization and to separate dynamic changes caused by binding (or bioconjugation) from those involved in activity enhancement. The approach proved useful not only to confirm the location of the previously suggested CYP3A4 allosteric site, but also to identify a series of coupled structural elements (the K/β1-loop and F-helix) specifically involved in activity enhancement. Allosteric activation or inhibition of CYP3A4 by small pharmaceuticals (*e*.*g*. midazolam and warfarin) is an important mechanism associated with drug interactions in humans.^61–63^ Such adverse interactions are currently established empirically, but a better mechanistic understanding of the phenomena may improve *in silico* predictions, thereby facilitating the rational design of pharmaceuticals with lower potential for adverse drug interactions. Considering that allostery is ubiquitous in biological systems, the general approach outlined in this work should find much broader application in unraveling the allosteric properties of other enzyme systems.

## Supporting information

Tablesof peptides

## Acknowledgements

This research was funded by the National Science and Engineering Research Council of Canada (NSERC), the Center in Green Chemistry and Catalysis (CGCC), and the Canadian Foundation for Innovation (CFI). Julie Ducharme was supported by scholarships from the CGCC and the Fonds de recherche du Quebec - nature et technologies (FRQNT). We would like to thank Dr. A. S. Wahba for his work on protein LC-MS-QToF, Dr. J. R. Halpert for providing the CYP3A4 plasmid and Dr. C. B. Kasper for the CPR plasmid.

## Methods

### 1. Generation of mutants with a single reactive

cysteine residue CYP3A4 mutants were generated using the plasmid of a previously reported *N*-terminally truncated cysteine depleted mutant.^18^ All mutants generated have 5 cysteines mutated to other residues (C58T/C64A/C98S/C239S/C468G) with a 10 amino acid long truncation on the *N*-terminus along with a newly introduced cysteine for the site specific bioconjugation, and a His_4_-tag. The oligonucelotide primers used to generate the variant enzymes are found in Table S13. The mutants used in this study have been characterized previously^18,64^ and have been shown to be active for 7-benzyloxy-4-trifluoromethylcoumarin (BFC) oxidation. The fact that these enzymes can still be stimulated by progesterone suggests that the allosteric site is preserved.

### 2. Protein expression and purification

All mutants and wild type CYP3A4 were expressed and purified as previously described in similar yields.^18^ Briefly, wildtype CYP3A4 and its mutants were expressed in DH5-α *E. coli* cells grown in TB broth supplemented with D-Ala, IPTG and trace metals. The purification was performed in two steps. The first was an affinity purification using a Ni-NTA agarose resin (Qiagen). The protein was eluted with 200 mM imidazole in potassium phosphate buffer (0.05 M, pH 7.4, 10% glycerol). The second purification step was an ion exchange purification where a Macro-Prep High S resin (BioRad) was used. The protein was eluted with 0.5 M KCl in potassium phosphate buffer. The protein was stored at -80 °C in 0.1 M potassium phosphate, pH 7.4 containing 10% glycerol. Cytochrome P450 reductase (CPR) was expressed and purified as previously reported^17^ except for one important modification: the liquid growth medium was supplemented with a sterile, fully-dissolved riboflavin solution (0.2 mg/mL, 50:50 ACN: H_2_O, pH 11), which was added to the media at a volume ratio of 5 mL/L of medium at the time of inoculation. The CYP3A4 and CPR expression plasmids were generously provided by Dr. J. R. Halpert (University of Connecticut) and Dr. C. B. Kasper (University of Wisconsin-Madison), respectively.

### 3. Protein bioconjugation

The bioconjugation of CYP3A4 mutants with the maleimide-containing progesterone derivative (PGM) and characterization of the bioconjugates were performed as reported elsewhere.^18^ PGM was synthesized as previously described.^17^ Briefly, the mutants (2 *μ*M) were incubated in TCEP-containing (45 μM) potassium phosphate buffer (0.1 M, pH 7.4, 10% glycerol) for 20 min at 25 °C. The bioconjugation reaction with PGM (100 μM) was allowed to proceed with gentle shaking on an orbital shaker (2 h, 4°C, 60 rpm). The reaction was next quenched using DTT (2 mM) and desalted with Zeba Spin Desalting Columns (Pierce). The bioconjugation yield was determined by LC-MS-QToF (Figure S14).

### 4. Hydrogen-deuterium exchange protocol

Bioconjugates, unmodified mutants, and wild type CYP3A4 were concentrated to a final concentration of minimum 150 μM using Spin-X concentrators (30 kDa MWCO, 0.5 mL, Corning). The protein concentration was determined using the CO difference spectral assay to ensure that the protein concentration corresponded to the properly folded CYP3A4 form.^65^ All exchange reactions contained 1 μM of thus prepared enzyme in buffer (0.1 M KPi-D_2_O buffer, pD 7.4). This ensured a final D_2_O concentration >98% (D_2_O/H_2_O) which is critical for maximum deuterium uptake. When required, progesterone was added to a final concentration of 20 μM from a 20 mM stock solution in DMSO. Hydrogen-deuterium exchange was initiated by addition of the enzyme. At the desired time point (0.5 – 240 min), aliquots of the reaction mixture (50 μL) were immediately quenched with pH 1.7 buffer (100 μL, 100 mM KPi) to a final pH of 2.5, thus ensuring minimal amide H-D back exchange. The quenched samples were flash frozen in liquid nitrogen and stored at -80°C. Each reaction was performed in triplicate on the same day using the same stock solutions.

### 5. MS data acquisition

All HDX-MS experiments were performed on a Waters Synapt G2-Si with HDX technology, and MS data acquisition was performed as previously reported.^66^ One at the time, the quenched reaction samples (150 μL, stored at -80°C in 1.5 mL Eppendorf tubes) were thawed for 70 s in a water bath at 32°C and then loaded into a 40 μL injection loop of the HDX manager. The samples were injected in the HDX manager exactly 2 min after they were removed from the freezer. In the HDX manager, the protein was cleaved by elution (100 *μ*L/min flow rate of 0.1% formic acid in MilliQ water) through an online immobilized pepsin column for 3 min at 15°C. The resulting peptic peptides were trapped on a C18 guard column held at 0.4°C before reverse-phase chromatographic separation using a Waters BEH C18 UPLC column (1 × 100 mm). Prior to loading peptides, the column was equilibrated in 97:3 (v/v) MilliQ water:acetonitrile containing 0.1% formic acid. The column was eluted at 0.4°C with a linear gradient of 3-100% acetonitrile containing 0.1% formic acid applied over 10 min. The samples were ionized by electrospray ionization (ESI) with a capillary voltage of 2.8 kV, sampling cone of 30 V, source offset of 30 V, and desolvation temperature of 175 °C. Data were collected in positive ion and resolution modes. The ionized peptides were then further separated in the gas phase by travelling wave ion mobility using a wave velocity of 650 m/s, wave height of 40 V, a bias voltage of 3 V, and a nitrogen pressure of 3.1 mbar. After exiting the ion mobility region, the peptides were subjected to two alternating collision energy regimes applied over 0.4 s intervals: a low collision energy of 6 V and a high collision energy ramp from 21 – 44 V. The high energy regime fragments ions by collision induced dissociation (CID), while the low energy regime conserves the peptide precursor masses. This process allows the matching of peptide fragments with their respective parent ions and enables peptide identification with high confidence. A [glu-1]-fibrinopeptide B (Glu-Fib) external lockmass standard was, at all times, collected in parallel with all samples to enable correction of the peptide m/z values. An average variance in the determination of deuterium uptake values for this specific workflow was previously determined to be 0.087 Da with a standard deviation of 0.095 Da.^66^ These values are similar to previously reported peptide-level continuous exchange, bottom-up HDX-MS deuterium uptake differences.^67,68^

### 6. MS data processing

For each mutant, a peak list of the peptides that were reproducibly detected by our workflow was generated. These lists were determined by analyzing reference samples for each mutant prepared in triplicate in protiated potassium phosphate buffer (100 mM, pH 2.5) and subjected to the HDX-MS workflow described above. The peptide reference lists were generated by uploading the raw MS data in the Protein Lynx Global Server (PLGS) software (Waters) as previously described.^66^ PLGS scans the raw data and associates the detected peptide peaks with expected masses. Spectral assignments are scored by a variety of user-defined criteria: ppm error, difference in chromatographic retention time, difference in ion mobility drift time and number of MS/MS ion matches. The PLGS output is uploaded into DynamX 3.0 (Waters) for additional thresholding and data analysis. The final peptide list was restricted to include only peptides with m/z values within 5 ppm of the theoretical mass, 0.2 minimum products per amino acid, and present in all replicates. Depending on the mutant, this resulted in a list of 200 – 410 peptides covering 95.4% to 97.2% of CYP3A4 sequence with an amino acid redundancy of 5.43 to 13.26.

### 7. MS data analysis

DynamX 3.0 was used to quantify the level of deuterium uptake for each peptide under all conditions tested. DynamX 3.0 calculates a centroid mass from the isotopic distribution of deuterated peptide m/z signals. The centroid mass is then compared to the similarly determined centroid mass from the undeuterated reference peptides. The deuterium uptake is quantified based on this comparison and is reported as either relative uptake (in units of Da) or in relative fractional uptake (RFU, in units of %), where the absolute deuterium uptake is normalized by the total number of amide protons in the peptide. Data was curated manually to ensure that all peptide assignments by the software were correct. Differential HDX was used to compare two states by subtracting the deuterium uptake data for each peptide for the two states of interest. We ensured that all the compared peptides where from the same charge state, and that the reference and deuterated peptide had matching chromatographic retention times, ion mobility drift times, and masses.

### 8. Assessing HDX-MS variance

The overall reproducibility of the workflow was established as previously described.^66^ The significant deuterium uptake difference between compared states was determined using the Deuteros software package developed by Politis and coworkers.^24^ This software calculates confidence intervals based on the standard deviation obtained for all the deuterium uptake measurements for each specific time points. All difference data reported in this work represent a sum of the HDX difference values for each peptide at each of the analyzed HDX time points. A 98% confidence limit was used as the threshold to define a significant uptake difference.

### 9. Progesterone hydroxylation assay

Progesterone hydroxylation was determined by LC-MS. The reaction mixture contained CYP3A4 (1 *μ*M), Progesterone (20 *μ*M from an initial stock of 20 mM in DMSO), CPR (1 *μ*M) and NADPH (1 mM from a 25 mM stock in MilliQ water) in potassium phosphate buffer (100 mM, pH 7.4, 10% glycerol). Except for NADPH, all reaction components were pre-incubated at 37°C and 250 rpm for 5 min. The reaction was initiated with addition of NADPH and allowed to proceed for 1 h. The final reaction volume was 150 *μ*l and all experiments were performed in triplicate. The reactions were quenched by addition of DCM (0.5 mL). The samples were extracted in DCM (2 x 0.5 mL) and the combined organic layers were dried under a flow of N_2_. The dried samples were re-dissolved in acetonitrile (ACN) (100 *μ*l) before LC-UV-MS analysis. The samples were run on a C-18 Omega Polar HPLC column (Phenomenex, 250 x 4.6mm, <u>5</u>*<u>μ</u>*<u>m particle</u> size) with the following method: 50%-50% ACN:H_2_O for 3 min followed by a 9 min linear increase to 80% ACN and 8 min linear increase to a final 95% concentration of ACN. Hydroxylated progesterone products eluted between 12-16 min and progesterone eluted at 21 min.

## Notes

### Competing Interest Statement

The authors have declared no competing interest.

