## Supplemental Figures and Tables for "Combining small-molecule bioconjugation and hydrogen-deuterium exchange mass spectrometry (HDX-MS) to expose allostery: the case of human cytochrome P450 3A4"

| Data Set | WILDTYPE |
|---|---|
| HDX reaction details | 0.1 M KPi, pD = 7.4, RT, 1 μM protein |
| Exchange time points (min) | 0.5, 1, 2, 5, 30, 60, 240 |
| # of Peptides | 81 |
| Sequence coverage | 87.9% |
| Redundancy | 2.26 |
| Replicates | 3 technical |
| Repeatability (maximum SD between replicates, Da) | 0.21 |

| Data Set | MUTANT F215C | BIOCONJUGATE F215C_PGM | MUTANT F215C with PRG |
|---|---|---|---|
| HDX reaction details | 0.1 M KPi, pD = 7.4, RT, 1 μM protein | 0.1 M KPi, pD = 7.4, RT, 1 μM protein | 0.1 M KPi, pD = 7.4, RT, 1 μM protein |
| Exchange time points (min) | 0.5, 1, 2, 5, 30, 60, 240 | 0.5, 1, 2, 5, 30, 60, 240 | 5, 60 |
| # of Peptides | 72 | 72 | 76 |
| Sequence coverage | 87.1% | 87.1% | 87.3% |
| Redundancy | 1.85 | 1.85 | 1,97 |
| Replicates | 3 technical | 3 technical | 3 technical |
| Repeatability (maximum SD between replicates, Da) | 0.40 | 0.68 | 0.19 |
| Significant differences threshold (Exposure time sum) | | 0.53 Da (98% CI) | 0.50 Da (98% CI) |

| Data Set | MUTANT F108C | BIOCONJUGATE F108C_PGM |
|---|---|---|
| HDX reaction details | 0.1 M KPi, pD = 7.4, RT, 1 μM protein | 0.1 M KPi, pD = 7.4, RT, 1 μM protein |
| Exchange time points (min) | 0.5, 30, 120 | 0.5, 30, 120 |
| # of Peptides | 72 | 72 |
| Sequence coverage | 87.1% | 87.1% |
| Redundancy | 1.85 | 1.85 |
| Replicates | 3 technical | 3 technical |
| Repeatability (maximum SD between replicates, Da) | 0.25 | 0.24 |
| Significant differences threshold (Exposure time sum) | 0.41 Da (98% CI) | |

| Data Set | MUTANT G481C | BIOCONJUGATE G481C_PGM |
|---|---|---|
| HDX reaction details | 0.1 M KPi, pD = 7.4, RT, 1 μM protein | 0.1 M KPi, pD = 7.4, RT, 1 μM protein |
| Exchange time points (min) | 5, 60 | 5, 60 |
| # of Peptides | 81 | 80 |
| Sequence coverage | 87.5% | 87% |
| Redundancy | 2.16 | 2.16 |
| Replicates | 3 technical | 3 technical |
| Repeatability (maximum SD between replicates, Da) | 0.33 | 0.24 |
| Significant differences threshold (Exposure time sum) | 0.39 Da (98% CI) | |

| Data Set | MUTANT L482C | BIOCONJUGATE L482C_PGM |
|---|---|---|
| HDX reaction details | 0.1 M KPi, pD = 7.4, RT, 1 μM protein | 0.1 M KPi, pD = 7.4, RT, 1 μM protein |
| Exchange time points (min) | 5, 60 | 5, 60 |
| # of Peptides | 77 | 77 |
| Sequence coverage | 85.9% | 85.9% |
| Redundancy | 2.09 | 2.09 |
| Replicates | 3 technical | 3 technical |
| Repeatability (maximum SD between replicates, Da) | 0.29 | 0.15 |
| Significant differences threshold (Exposure time sum) | 0.46 Da (98% CI) | |

**Table S1.** Summary tables of HDX-MS results for all experimental conditions tested. Experiments were all performed under the same conditions, but the number of $D_2O$ exposure time points varied. The number and nature of the detected peptides varied slightly, but the percent sequence coverage remained higher than 85% for all states. Redundancy is a measure of degree of overlap between the detected peptides. The significant difference threshold is used when determing a significant uptake difference between two biochemical states (e.g. the free enzyme and its PGM-bioconjugate).

Δ                A″ –Helix

1 MALIPDLAME MALLLAVFLVLLYLGT **HSHGLFKKLG** IPGPTPLPFLG

A'- & A- Helix                Helix B

49 **NILSYHKGFCMFDMECHKKYG** KVWGFYDGQQPVLAIT **DPDMIKTVLV**

Helix B'                Helix C

96 KECYSVFTNRRPFGP **VGFMKSA** ISIA **EDEEWKRLRSLLSPTFT** S

Helix D                Helix E

140 **GKLKEMVPIIAQYGDVLVRNLRREAET** GKPVT **LKDVFGAYSMDVITSTSFG**

Helix F                Helix F′                Helix G′

191 VNIDSLNNPQDP **FVENTK** KLLRFDFL **DPFFLSITVF** PF **LIPILEVL** NICVF

Helix G                Helix H

242 **PREVTNFLRKSVKRMKESR** LEDTQKHRV **DFLQLMID** SQNSKETESHKALSDL

Helix I                Helix J

294 **ELVAQSIIFIFAGYETTSSVLSFIMYEL** AT **HPDVQQKLQEEIDAVL** PNKAPP

Helix J – K

346 **TYDTVLQMEYLDMVVNETLRLF** PIAMRLERVCKKDVEINGMFIPKGVVVMI **PSYALHR**

Helix L

404 DPKYWTEPEKF **LPERFS** KKNKDNIDPYIYTPFGSGPRNCI **GMRFALMNMKLALIRVLQ**

His – tag

462 NFSFKPCKETQIPLKLSLGGLLQPEKPVVLKVESRDGTVSGA HHHH

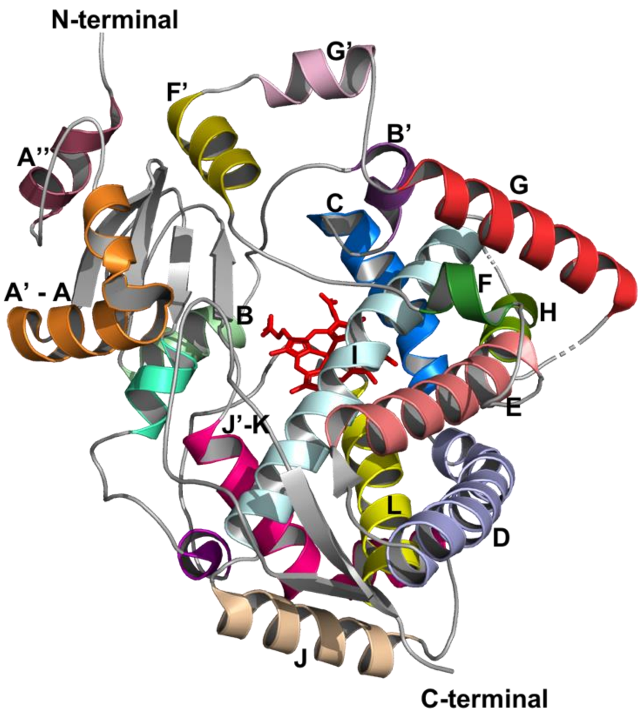


**Figure S2.** Sequence of the *N*-terminally truncated wild type CYP3A4. Residues that are part of α-helices are shown in bold in the sequence and are labeled according to the established P450 nomenclature. Each α-helix is color-coded and mapped onto the tertiary structure (PDB: 1W0F).[1] The enzyme used for crystallization of 1W0F is 13 amino acids shorter at the *N*-terminus than the CYP3A4 construct used in this study.

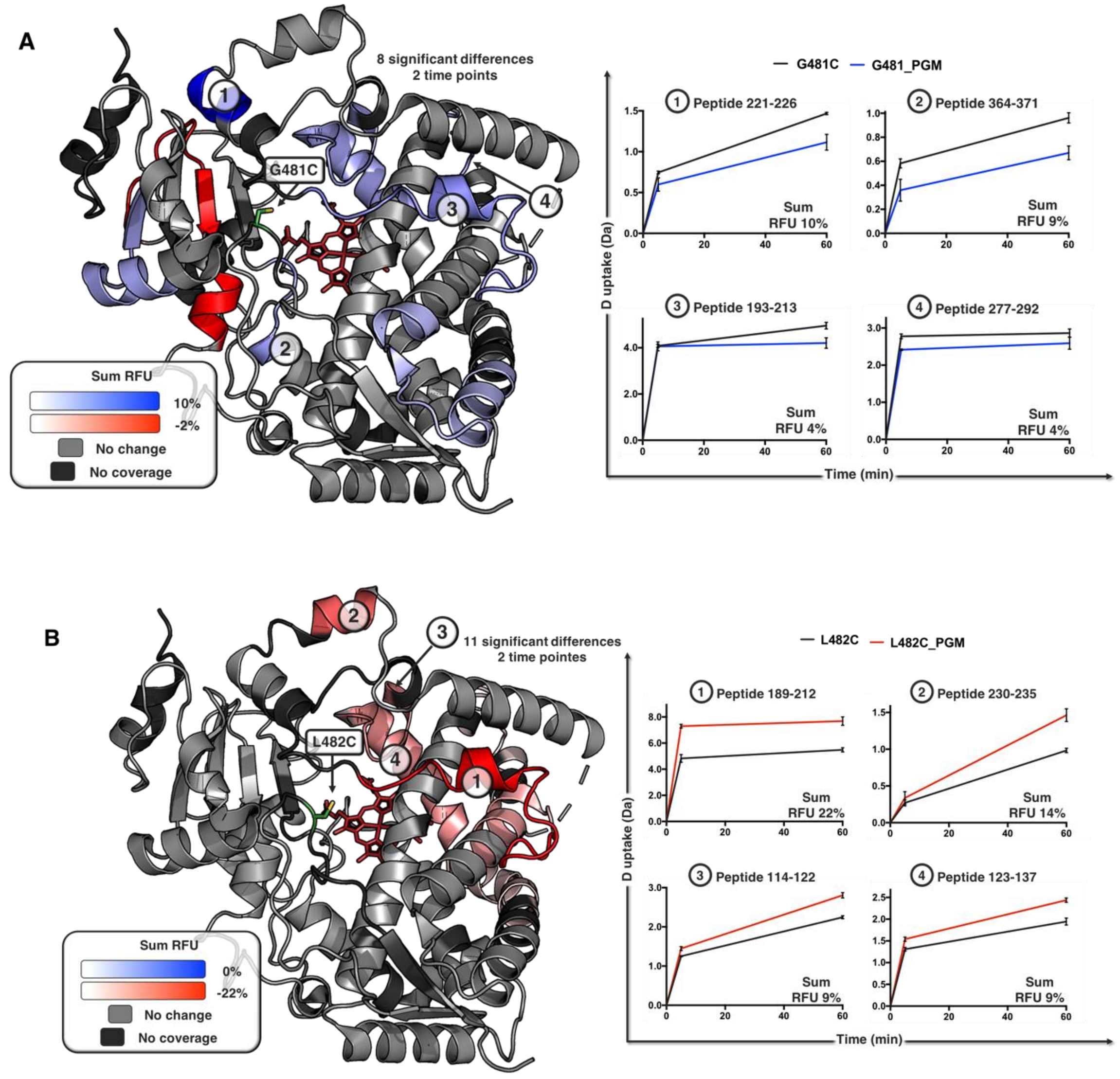


**Figure S3.** Differential HDX profiles of antagonistic and agonistic PGM-bioconjugates. **A)** Sum of the RFU difference for the CYP3A4 G481C mutant before and after PGM bioconjugation (antagonistic). A total of eight peptides exhibited a significant sum RFU difference across two exchange time points (5, 60 min). **B)** Sum of the RFU difference for the CYP3A4 L482C mutant before and after PGM bioconjugation (agonistic). A total of 11 peptides exhibited a significant sum RFU difference across two exchange times (5, 60 min). The summed RFU difference (CI 98%) is mapped onto the CYP3A4 structure (PDB: 1W0F). Protein regions shown in blue undergo a decrease in the extent of deuterium uptake upon bioconjugation, while elements colored in red undergo an increase in deuterium uptake. Regions displayed in light gray did not undergo any significant change in deuterium uptake after PGM-labeling, and those for which coverage is not significant are shown in dark gray. The cysteine bioconjugation handles can be found in the structure where the mutation number is shown. For both state comparisons, the four most affected regions are ranked from 1 to 4. Deuterium uptake plots for peptides derived from these regions are shown on the right. The summed RFU difference across both exchange time points is shown at the bottom of each graph.

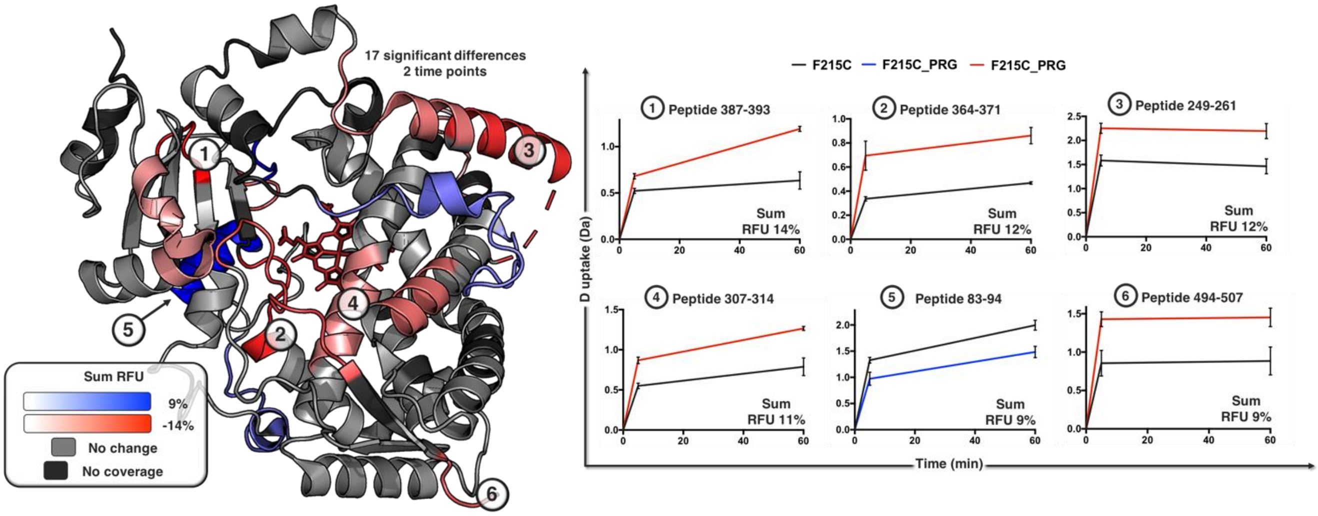


**Figure S4.** Differential HDX profiles for the CYP3A4 F215C mutant bound to progesterone (PRG). Sum of the RFU difference with and without progesterone. A total of 17 peptides exhibited significant RFU differences between the two states across two exchange times (5, 60 min). The summed relative fractional uptake (RFU) difference (CI 98%) is mapped onto the CYP3A4 structure (PDB: 1W0F). Protein regions shown in blue undergo a decrease in the extent of deuterium uptake upon bioconjugation, while elements colored in red undergo an increase in deuterium uptake. Regions displayed in light gray did not undergo any significant change in deuterium uptake after PGM-labeling, and those for which coverage is not significant are shown in dark gray. The six most affected regions are ranked from 1 to 6. Deuterium uptake plots for peptides derived from these regions are shown on the right. The summed RFU difference across both exchange time points is shown at the bottom of each graph.

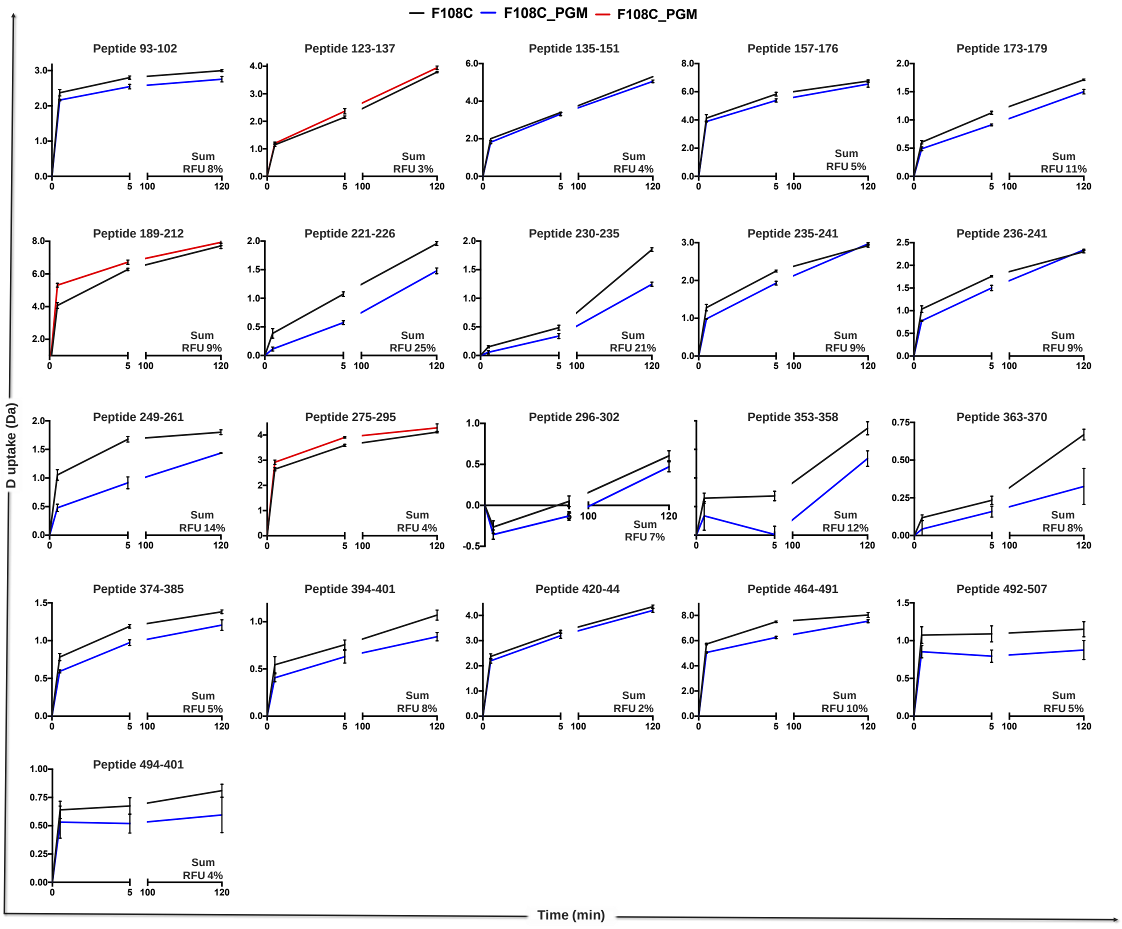


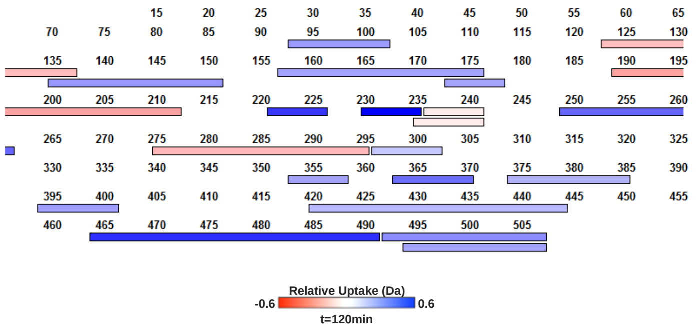


| Start | End | Significant (CI 98%) deuterium uptake difference (Da) | | | | |
|---|---|---|---|---|---|---|
| | | t= 0.5 min | t= 5 min | t=120 min | Sum | Sum %RFU |
| 93 | 102 | 0 | 0.60 | 0 | 0.70 | 7.83 |
| 135 | 151 | 0 | 0 | 0 | 0.52 | 3.71 |
| 157 | 176 | 0.24 | 0.45 | 0 | 0.91 | 5.00 |
| 173 | 178 | 0 | 0 | 0 | 0.54 | 10.82 |
| 221 | 226 | 0.27 | 0.50 | 0.48 | 1.26 | 24.95 |
| 230 | 235 | 0 | 0 | 0.61 | 0.85 | 21.25 |
| 235 | 241 | 0.30 | 0.32 | 0 | 0.57 | 9.54 |
| 236 | 241 | 0.26 | 0.26 | 0 | 0.48 | 9.53 |
| 249 | 261 | 0.57 | 0.76 | 0.36 | 1.70 | 14.18 |
| 296 | 302 | 0 | 0 | 0 | 0.40 | 6.75 |
| 353 | 358 | 0 | 0.27 | 0 | 0.60 | 12.11 |
| 363 | 370 | 0 | 0 | 0.34 | 0.49 | 8.22 |
| 374 | 385 | 0 | 0 | 0 | 0.58 | 5.29 |
| 394 | 401 | 0 | 0 | 0 | 0.49 | 8.22 |
| 420 | 444 | 0 | 0 | 0 | 0.49 | 2.33 |
| 464 | 491 | 0.66 | 1.24 | 0.49 | 2.40 | 10.43 |
| 492 | 507 | 0 | 0.29 | 0.27 | 0.79 | 5.27 |
| 494 | 507 | 0 | 0 | 0 | 0.48 | 3.69 |
| 123 | 137 | 0 | 0 | 0 | - 0.43 | - 3.30 |
| 189 | 212 | - 1.24 | - 0.4414 | 0 | - 1.90 | - 9.06 |
| 275 | 295 | - 0.28 | - 0.3132 | 0 | - 0.76 | - 3.80 |

**Coverage= 87.1% Total peptides= 73, Significant peptides= 21**
**Min= 0.41, Max= 3.25**

**Figure S5.** Significant deuterium uptake differences for the CYP3A4 F108C mutant before and after PGM bioconjugation presented in three ways (CI 98%): 1) the deuterium uptake plots for each significant peptide; 2) a coverage map strictly showing the peptides undergoing significant uptake difference; and 3) a table detailing the deuterium uptake (Da) per exposure time point, the total uptake sum (Da), and the sum of the relative fractional uptake (RFU) difference. The blue color indicates peptides showing a decrease in deuterium uptake after PGM-labeling (rigidification), and the red color represents peptides undergoing an increase in deuterium uptake over time (flexibilization).

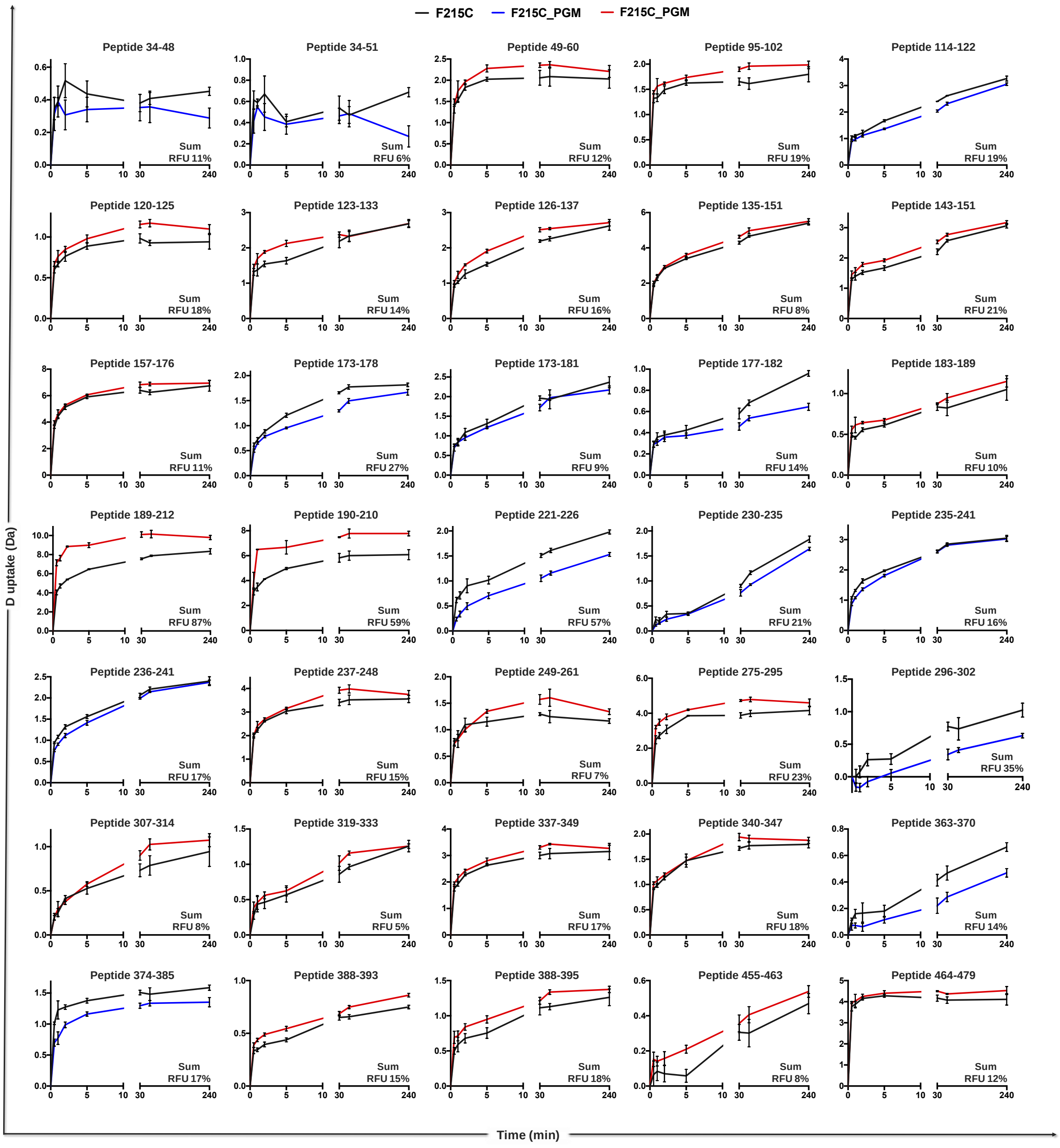

F215C
F215C_PGM
F215C_PGM
Peptide 34-48
Sum RFU 11%
Peptide 34-51
Sum RFU 6%
Peptide 49-60
Sum RFU 12%
Peptide 95-102
Sum RFU 19%
Peptide 114-122
Sum RFU 19%
Peptide 120-125
Sum RFU 18%
Peptide 123-133
Sum RFU 14%
Peptide 126-137
Sum RFU 16%
Peptide 135-151
Sum RFU 8%
Peptide 143-151
Sum RFU 21%
Peptide 157-176
Sum RFU 11%
Peptide 173-178
Sum RFU 27%
Peptide 173-181
Sum RFU 9%
Peptide 177-182
Sum RFU 14%
Peptide 183-189
Sum RFU 10%
Peptide 189-212
Sum RFU 87%
Peptide 190-210
Sum RFU 59%
Peptide 221-226
Sum RFU 57%
Peptide 230-235
Sum RFU 21%
Peptide 235-241
Sum RFU 16%
Peptide 236-241
Sum RFU 17%
Peptide 237-248
Sum RFU 15%
Peptide 249-261
Sum RFU 7%
Peptide 275-295
Sum RFU 23%
Peptide 296-302
Sum RFU 35%
Peptide 307-314
Sum RFU 8%
Peptide 319-333
Sum RFU 5%
Peptide 337-349
Sum RFU 17%
Peptide 340-347
Sum RFU 18%
Peptide 363-370
Sum RFU 14%
Peptide 374-385
Sum RFU 17%
Peptide 388-393
Sum RFU 15%
Peptide 388-395
Sum RFU 18%
Peptide 455-463
Sum RFU 8%
Peptide 464-479
Sum RFU 12%
D uptake (Da)
Time (min)

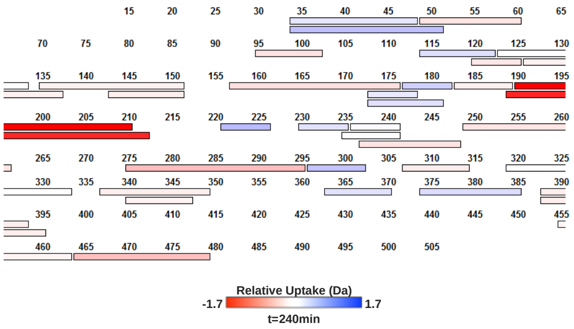


| Start | End | Deuterium uptake difference (Da) | | | | | | | | |
|---|---|---|---|---|---|---|---|---|---|---|
| | | t=0.5 min | t=1 min | t=2 min | t=5 min | t=30 min | t=60 min | t=240 min | Sum | Sum %RFU |
| 34 | 48 | 0 | 0 | 0 | 0 | 0 | 0 | 0 | 0.58 | 5.81 |
| 34 | 51 | 0 | 0 | 0 | 0 | 0 | 0 | 0.42 | 0.96 | 7.36 |
| 114 | 122 | 0 | 0 | 0 | 0.30 | 0.36 | 0 | 0 | 1.56 | 19.51 |
| 173 | 178 | 0 | 0 | 0 | 0 | 0.36 | 0 | 0 | 1.33 | 26.64 |
| 173 | 181 | 0 | 0 | 0 | 0 | 0 | 0 | 0 | 0.69 | 8.63 |
| 177 | 182 | 0 | 0 | 0 | 0 | 0 | 0 | 0 | 0.70 | 14.07 |
| 221 | 226 | 0.38 | 0.37 | 0.41 | 0.31 | 0.45 | 0.45 | 0.45 | 2.83 | 56.64 |
| 230 | 235 | 0 | 0 | 0 | 0 | 0 | 0 | 0 | 0.83 | 20.85 |
| 235 | 241 | 0 | 0 | 0.27 | 0 | 0 | 0 | 0 | 0.94 | 15.62 |
| 236 | 241 | 0 | 0 | 0 | 0 | 0 | 0 | 0 | 0.87 | 17.41 |
| 296 | 302 | 0 | 0 | 0.34 | 0 | 0.43 | 0 | 0.39 | 2.10 | 35.14 |
| 363 | 370 | 0 | 0 | 0 | 0 | 0 | 0 | 0 | 0.86 | 14.30 |
| 374 | 385 | 0.31 | 0.45 | 0.29 | 0 | 0 | 0 | 0 | 1.86 | 16.00 |
| 49 | 60 | 0 | 0 | 0 | - 0.25 | - 0.30 | 0 | 0 | - 1.36 | - 12.39 |
| 95 | 102 | 0 | 0 | 0 | 0 | 0 | - 0.34 | 0 | - 1.36 | - 19.46 |
| 120 | 125 | 0 | 0 | 0 | 0 | 0 | 0 | 0 | - 0.91 | - 18.14 |
| 123 | 133 | 0 | 0 | - 0.34 | - 0.49 | 0 | 0 | 0 | - 1.45 | - 14.46 |
| 126 | 137 | 0 | 0 | - 0.26 | - 0.37 | - 0.32 | 0 | 0 | - 1.62 | - 16.21 |
| 135 | 151 | 0 | 0 | 0 | 0 | - 0.33 | 0 | 0 | - 1.17 | - 8.38 |
| 143 | 151 | 0 | 0 | - 0.26 | 0 | - 0.32 | 0 | 0 | - 1.4 | - 20.76 |
| 157 | 176 | - 0.30 | 0 | 0 | 0 | - 0.44 | - 0.63 | 0 | - 2.05 | - 11.36 |
| 183 | 189 | 0 | 0 | 0 | 0 | 0 | 0 | 0 | - 0.63 | - 10.54 |
| 190 | 210 | - 0.86 | - 3.00 | 0 | - 1.70 | - 1.66 | - 1.77 | - 1.70 | - 10.69 | - 59.42 |
| 189 | 212 | -3.11 | - 2.92 | - 3.46 | - 2.52 | - 2.55 | - 2.28 | - 1.44 | - 18.27 | - 86.99 |
| 237 | 248 | 0 | 0 | 0 | 0 | - 0.52 | - 0.46 | 0 | - 1.53 | - 15.27 |
| 249 | 261 | 0 | 0 | 0 | 0 | - 0.28 | - 0.36 | 0 | - 0.85 | - 7.10 |
| 275 | 295 | - 0.73 | - 0.77 | - 0.71 | - 0.34 | - 0.85 | - 0.81 | - 0.43 | - 4.63 | - 23.16 |
| 307 | 314 | 0 | 0 | 0 | 0 | 0 | 0 | 0 | - 0.58 | - 8.28 |
| 319 | 333 | 0 | 0 | 0 | 0 | 0 | 0 | 0 | - 0.64 | - 4.92 |
| 337 | 349 | 0 | 0 | 0 | 0 | - 0.30 | - 0.36 | 0 | - 1.52 | - 16.89 |
| 340 | 347 | 0 | 0 | 0 | 0 | 0 | 0 | 0 | - 0.70 | - 17.58 |
| 388 | 393 | 0 | 0 | 0 | 0 | 0 | 0 | 0 | - 0.59 | - 14.65 |
| 388 | 395 | 0 | 0 | 0 | 0 | 0 | 0 | 0 | - 1.09 | - 18.19 |
| 455 | 463 | 0 | 0 | 0 | 0 | 0 | 0 | 0 | - 0.61 | - 7.60 |
| 464 | 479 | 0 | 0 | 0 | 0 | - 0.33 | 0 | - 0.42 | - 1.59 | - 12.22 |

**Coverage= 87.1% Total peptides= 70, Significant peptides= 35, Min= 0.58, Max= 18.27**

**Figure S6.** Significant deuterium uptake differences for the CYP3A4 F215C mutant before and after PGM bioconjugation presented in three ways (CI 98%): 1) the deuterium uptake plots for each significant peptide; 2) a coverage map strictly showing the peptides undergoing significant uptake difference; and 3) a table detailing the deuterium uptake (Da) per exposure time point, the total uptake sum (Da), and the sum of the relative fractional uptake (RFU) difference. The blue color indicates peptides showing a decrease in deuterium uptake after PGM-labeling (rigidification), and the red color represents peptides undergoing an increase in deuterium uptake over time (flexibilization).

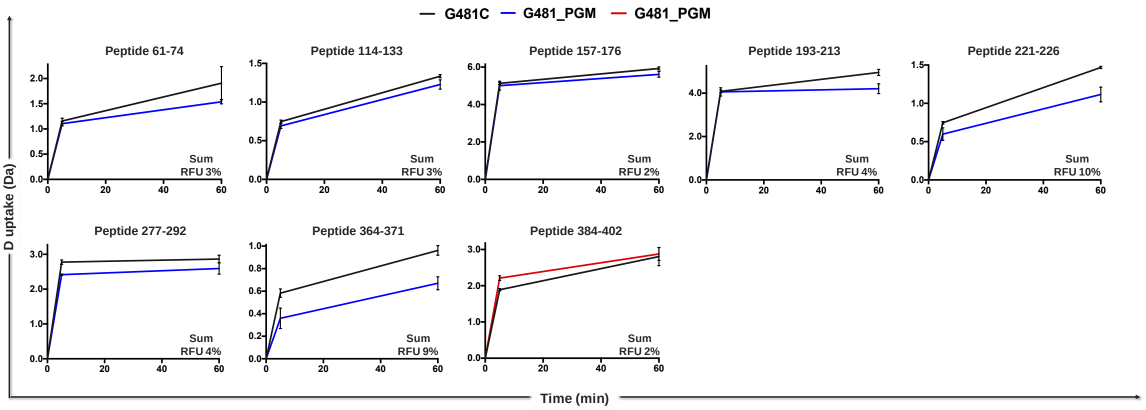


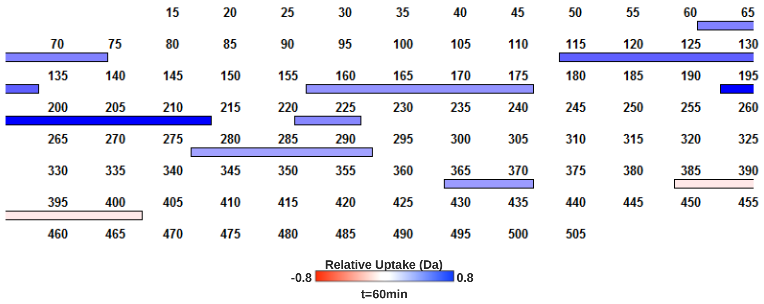


| Start | End | Deuterium uptake difference (Da) t=5min | t=60min | Sum | Sum % RFU |
|---|---|---|---|---|---|
| 61 | 74 | 0 | 0.37 | 0.42 | 3.24 |
| 114 | 133 | 0 | 0.47 | 0.62 | 3.24 |
| 157 | 176 | 0 | 0.32 | 0.44 | 2.46 |
| 193 | 213 | 0 | 0.75 | 0.77 | 4.30 |
| 221 | 226 | 0 | 0.35 | 0.50 | 10.00 |
| 277 | 292 | 0.36 | 0.27 | 0.63 | 4.20 |
| 364 | 371 | 0.22 | 0.29 | 0.51 | 8.59 |
| 384 | 402 | 0.32 | 0 | -0.39 | -2.45 |

Coverage= 87% Total peptides= 80, Significant peptides= 8, Min= 0.39, Max= 0.77

**Figure S7.** Significant deuterium uptake differences for the CYP3A4 G481C mutant before and after PGM bioconjugation presented in three ways (CI 98%): 1) the deuterium uptake plots for each significant peptide; 2) a coverage map strictly showing the peptides undergoing significant uptake difference; and 3) a table detailing the deuterium uptake (Da) per exposure time point, the total uptake sum (Da), and the sum of the relative fractional uptake (RFU) difference. The blue color indicates peptides showing a decrease in deuterium uptake after PGM-labeling (rigidification), and the red color represents peptides undergoing an increase in deuterium uptake over time (flexibilization).

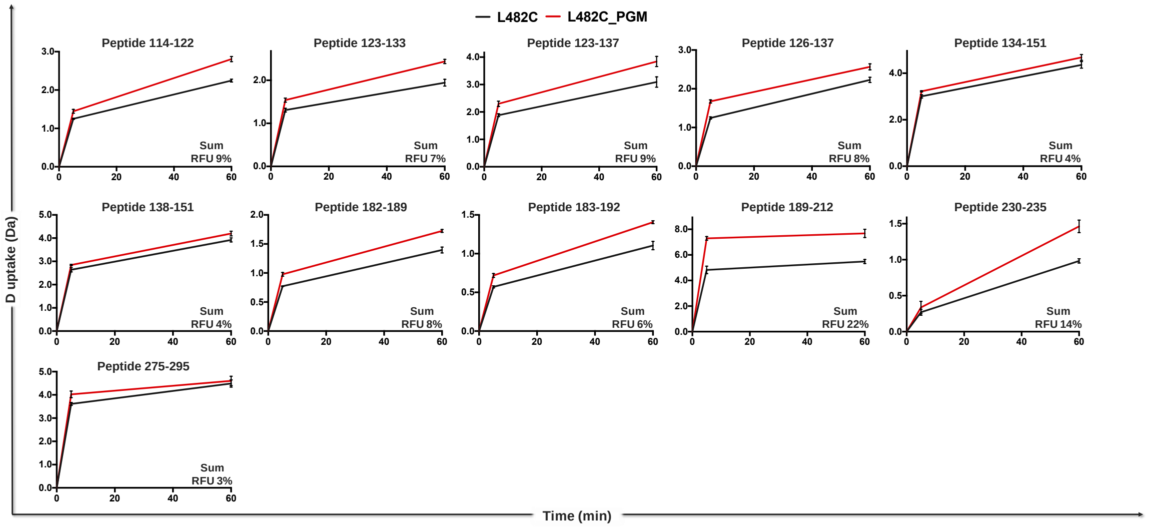


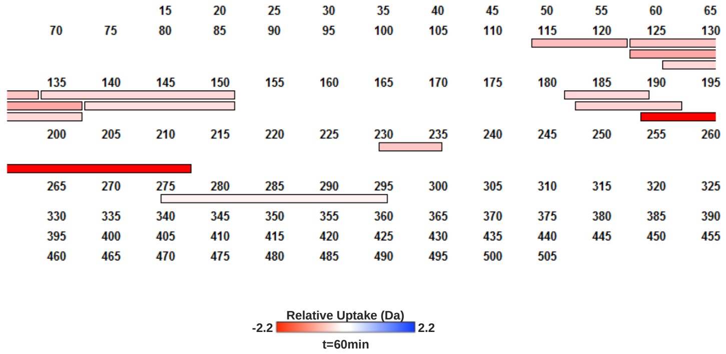


| Start | End | Deuterium uptake difference (Da) | | | |
|---|---|---|---|---|---|
| | | t=5 min | t=60 min | Sum | Sum %RFU |
| 114 | 122 | 0 | - 0.56 | - 0.75 | -9.44 |
| 123 | 133 | - 0.27 | - 0.49 | - 0.73 | -7.26 |
| 123 | 137 | - 0.42 | - 0.75 | - 1.16 | -8.96 |
| 126 | 137 | - 0.43 | - 0.34 | - 0.77 | -7.66 |
| 134 | 151 | - 0.22 | - 0.32 | - 0.54 | -3.63 |
| 138 | 151 | 0 | 0 | - 0.48 | -4.01 |
| 182 | 189 | 0 | - 0.33 | - 0.53 | -7.62 |
| 183 | 192 | 0 | - 0.37 | - 0.54 | -5.98 |
| 189 | 212 | - 2.47 | - 2.20 | - 4.67 | -22.24 |
| 230 | 235 | 0 | - 0.48 | - 0.55 | -13.68 |
| 275 | 295 | - 0.41 | 0 | - 0.52 | -2.62 |

Coverage= 85.9% Total peptides= 77, Significant peptides= 11, Min= 0.48, Max= 4.67

**Figure S8.** Significant deuterium uptake differences for the CYP3A4 L482C mutant before and after PGM bioconjugation presented in three ways (CI 98%): 1) the deuterium uptake plots for each significant peptide; 2) a coverage map strictly showing the peptides undergoing significant uptake difference; and 3) a table detailing the deuterium uptake (Da) per exposure time point, the total uptake sum (Da), and the sum of the relative fractional uptake (RFU) difference. The blue color indicates peptides showing a decrease in deuterium uptake after PGM-labeling (rigidification), and the red color represents peptides undergoing an increase in deuterium uptake over time (flexibilization).

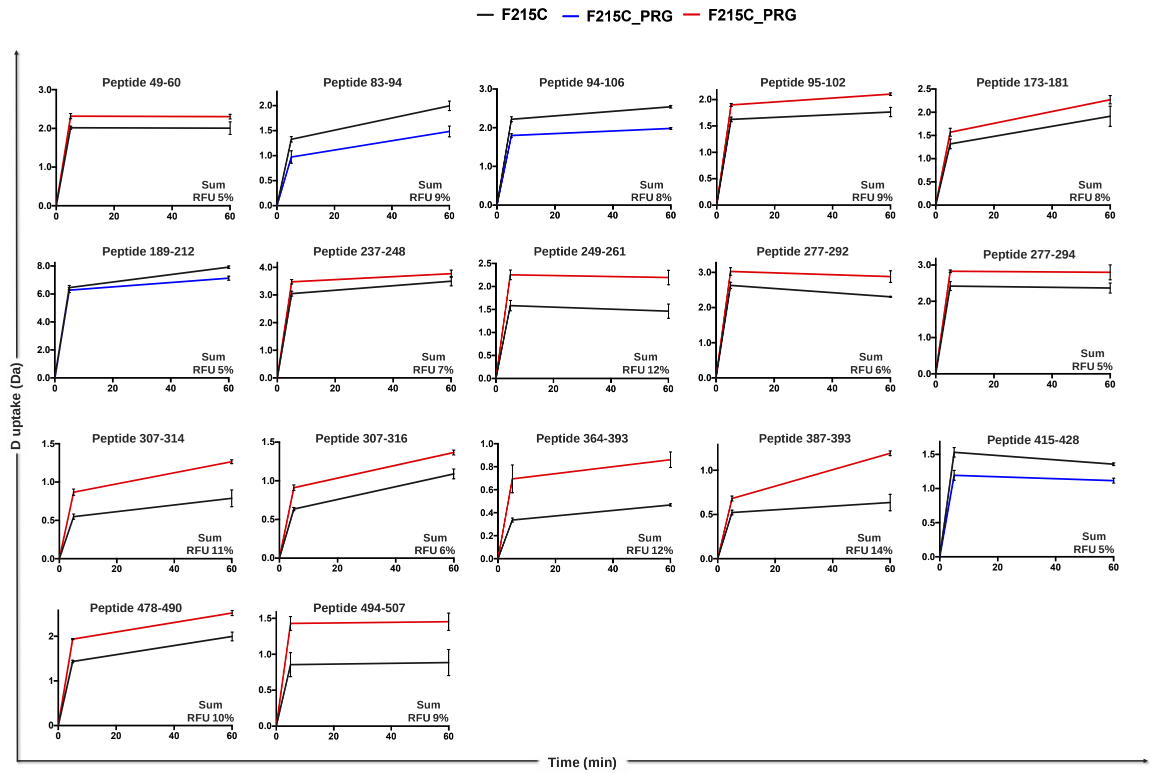


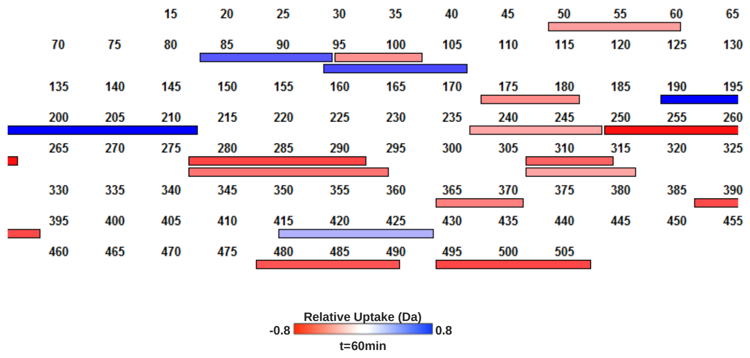


| Start | End | Deuterium uptake difference (Da) | | | |
|---|---|---|---|---|---|
| | | t=5 min | t=60 min | Sum | Sum %RFU |
| 83 | 94 | 0.35 | 0.52 | 0.87 | 8.67 |
| 94 | 106 | 0.42 | 0.56 | 0.98 | 8.18 |
| 189 | 212 | 0 | 0.78 | 0.96 | 4.57 |
| 415 | 428 | 0.34 | 0 | 0.58 | 4.81 |
| 49 | 60 | - 0.30 | 0 | - 0.60 | - 5.44 |
| 95 | 102 | - 0.27 | - 0.34 | - 0.61 | - 8.75 |
| 173 | 181 | - 0.25 | - 0.36 | - 0.61 | - 7.61 |
| 237 | 248 | - 0.43 | 0 | - 0.70 | - 7.00 |
| 249 | 261 | - 0.67 | - 0.73 | - 1.40 | - 11.66 |
| 277 | 292 | - 0.40 | - 0.57 | - 0.97 | - 6.48 |
| 277 | 294 | - 0.41 | - 0.43 | - 0.84 | - 4.94 |
| 307 | 314 | - 0.32 | - 0.48 | - 0.80 | - 11.39 |
| 307 | 316 | - 0.28 | 0 | - 0.55 | - 6.14 |
| 364 | 371 | - 0.36 | - 0.39 | - 0.75 | - 12.54 |
| 387 | 393 | 0 | - 0.56 | - 0.71 | - 14.31 |
| 478 | 490 | - 0.50 | - 0.52 | - 1.02 | - 10.23 |
| 494 | 507 | - 0.57 | - 0.57 | - 1.14 | - 8.77 |

Coverage= 87.3% Total peptides= 73, Significant peptides= 17, Min= 0.55, Max= 1.40

**Figure S9.** Significant deuterium uptake differences for the CYP3A4 F215C mutant with and without progesterone added presented in three ways (CI 98%): 1) the deuterium uptake plots for each significant peptide; 2) a coverage map strictly showing the peptides undergoing significant uptake difference; and 3) a table detailing the deuterium uptake (Da) per exposure time point, the total uptake sum (Da), and the sum of the relative fractional uptake (RFU) difference. The blue color indicates peptides showing a decrease in deuterium uptake in the presence of progesterone (rigidification), and the red color represents peptides undergoing an increase in deuterium uptake over time (flexibilization).

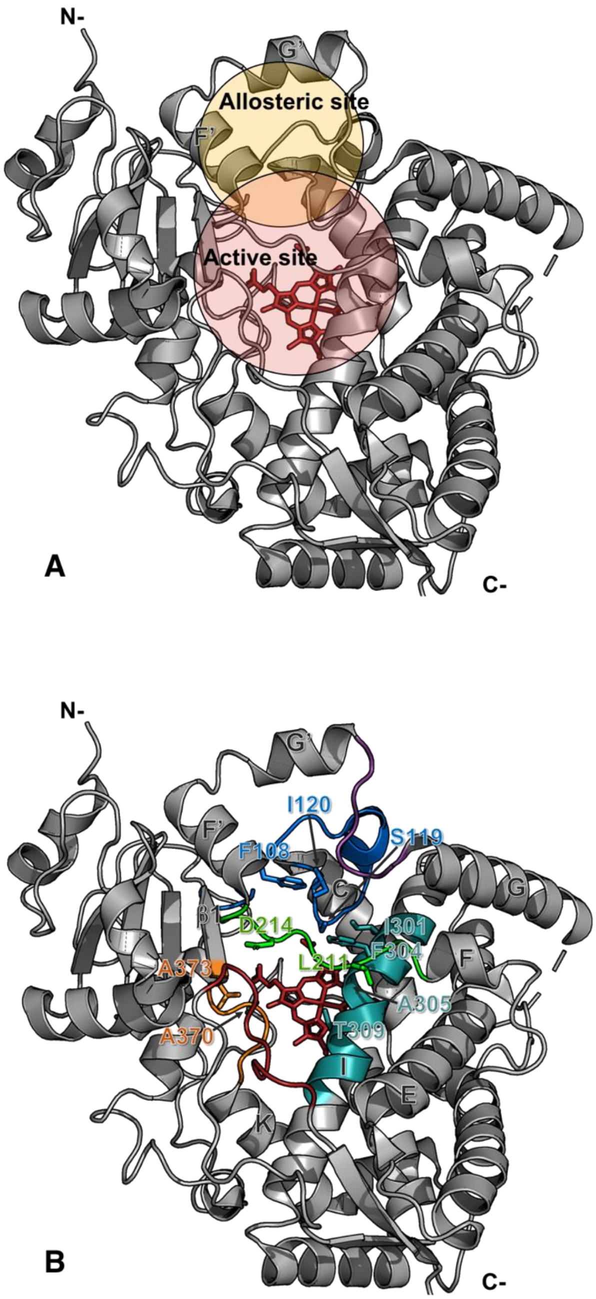


**Figure S10.** Important structural features of CYP3A4. **A)** Approximate position of the CYP3A4 active site showing the heme prosthetic group (red) and putative allosteric site (yellow). The two sites likely overlap. **B)** The six substrate recognition regions are highlighted: SRS-1; marine (B/C-loop), SRS-2; green (F/F'-loop), SRS-3; purple (G'/G-loop), SRS-4; teal (I-helix), SRS-5; orange (K/b1-loop), SRS-6; red (C-terminal loop).[2] SRS-4 and SRS-5 are deeper in the active site, running just over the heme group. The residues shown in the SRSs regions have previously been associated with cooperativity and are important in the regioselectivity of substrate oxidation.[3] Thr309 is essential for the proton transfer required to activate $O_2$.

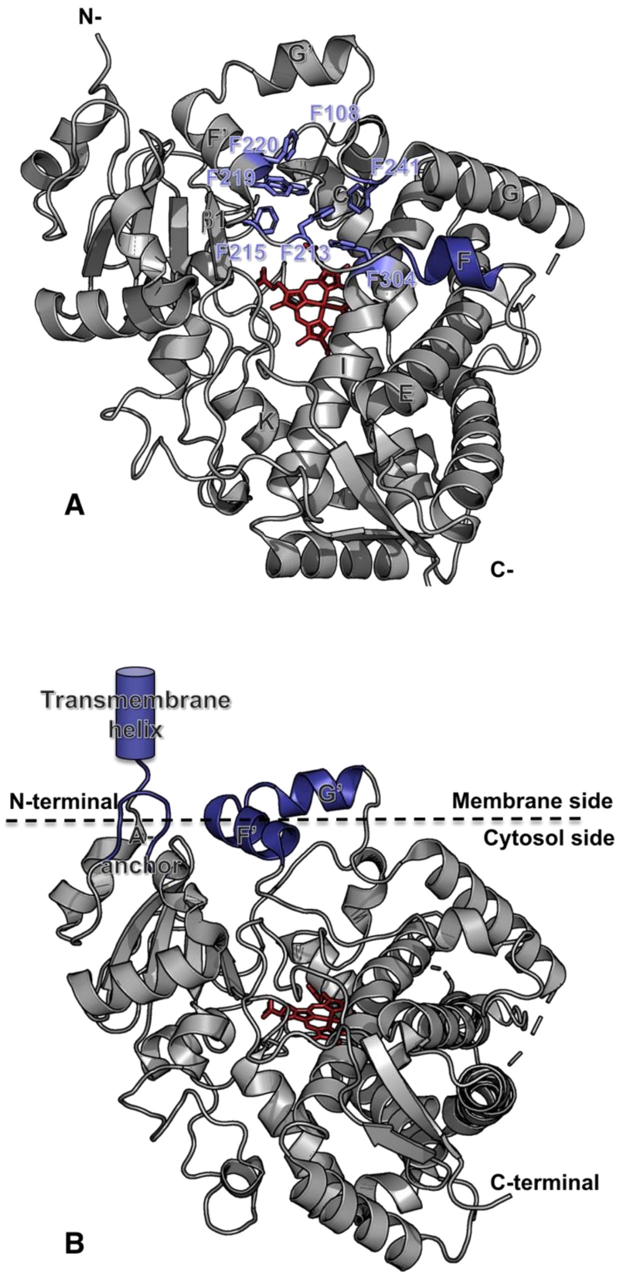


**Figure S11.** Important structural features of CYP3A4. **A)** The key unusual structural features of CYP3A4 as compared to other CYPs include an active site Phe-cluster and a shorter F-helix.[3] The cluster's phenylalanine residues are shown in light purple and the F-helix is shown in deep blue. **B)** In its native environment, CYP3A4 is bound to the ER membrane using a transmembrane helix anchor. Only the F'-helix, G'-helix and A''/A'-loop are thought to interact with the membrane.[4] P450 redox partners (such as cytochrome P450 reductase and cytochrome b5) interact with CYP3A4 on the cytosolic side.

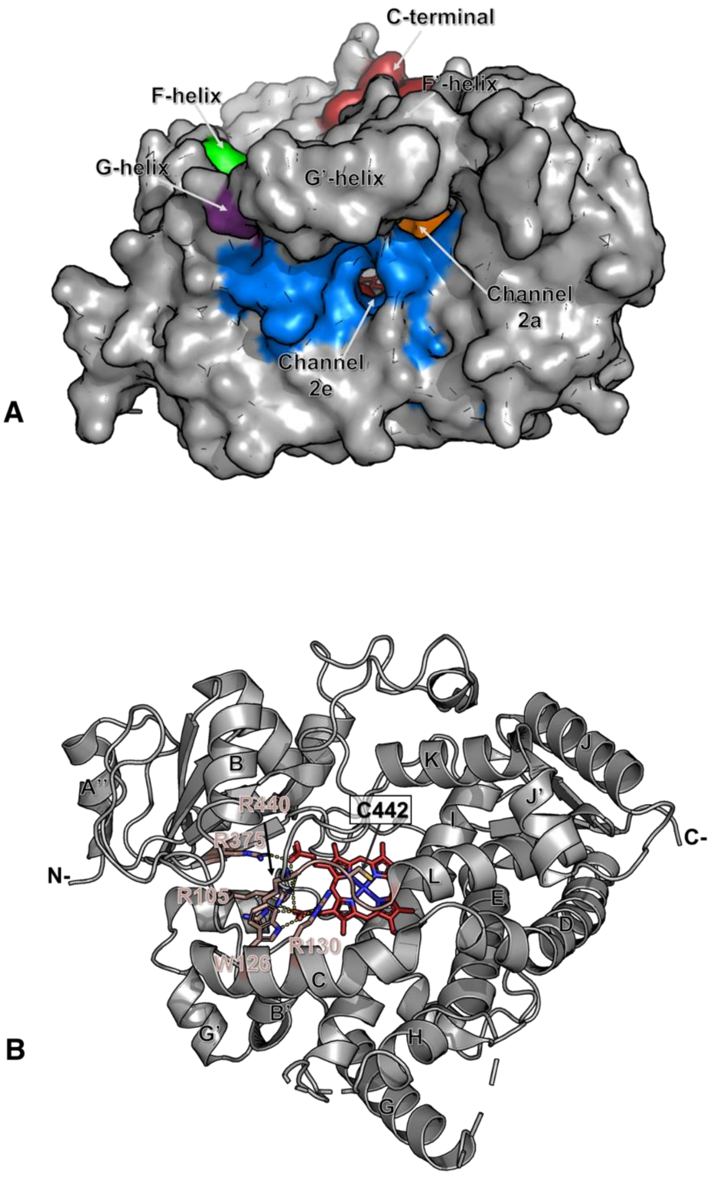


**Figure S12.** Important structural features of CYP3A4. **A)** The two main substrate-access channels (**2a** and **2e**) are separated by the B/C-loop (marine) and are open in most of the available CYP3A4 structures.[5] The **2a** and **2e** channels are located adjacent to the F'- and G'-helices, which form part of the putative allosteric site. **B)** Heme proximal side view showing the heme propionate salt bridges formed with four arginine and one tryptophan residues. Arg375 is involved in gating the active site water channel.[6] The cysteine at position 442 coordinates with the heme iron and is held in place by a conserved rigid B-bulge segment which controls the iron redox potential.[7]

| Mutation | Primer sequence |
|---|---|
| F108C | 5’-CCACTGGACCACAAGGCCTCCGGTTTGT-3’ |
| G481C | 5’-CCCCTGAAATTAAGCTTAGGATGCCTTCTTCAACCTGAAAAACCC-3’ |
| L482C | 5’-CCCTGAAATTAAGCTTAGGAGGATGTCTTCAACCTGAAAAACCCG-3’ |
| F215C | 5’-GAAAACACCAAGAAGCTTTTAAGATTTGATTGTTTGGATCCATT CTTTC-3’ |

**Table S13.** Oligonucleotide primers used for constructing the CYP3A4 mutants used in this study. Only the forward primers are shown.

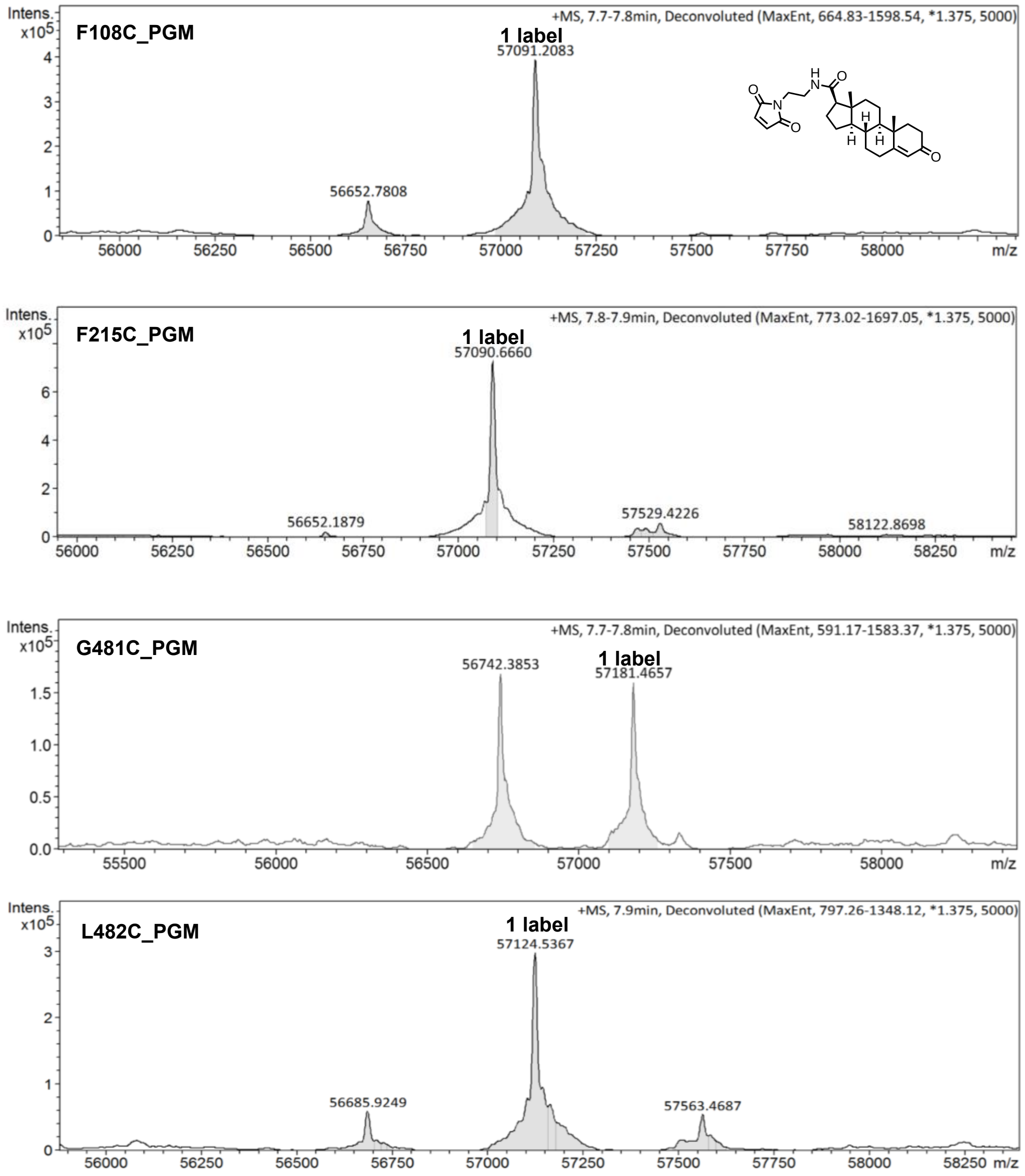


**Figure S14.** PGM-bioconjugation yields. Deconvoluted mass spectra for the different CYP3A4 mutants bioconjugated to PGM. Peaks correspond either to the unlabeled protein or the PGM-bioconjugate, singly or doubly labeled.
