## Supplementary material for "Combining small-molecule bioconjugation and hydrogen-deuterium exchange mass spectrometry (HDX-MS) to expose allostery: the case of human cytochrome P450 3A4": Tablesof peptides

| Deuterium uptake data for F108C and F108C_PGM CYP3A4 states | | | | | | |
|---|---|---|---|---|---|---|
| Start | End | Sequence | State | Exposure (min) | Uptake (Da) | Uptake SD (Da) |
| 33 | 48 | FKKLGIPGPTPLPFLG | F108C | 0 | 0 | 0 |
| 33 | 48 | FKKLGIPGPTPLPFLG | F108C | 0.5 | 0.3165 | 0.1165 |
| 33 | 48 | FKKLGIPGPTPLPFLG | F108C | 5 | 0.2732 | 0.0984 |
| 33 | 48 | FKKLGIPGPTPLPFLG | F108C | 120 | 0.3354 | 0.1043 |
| 33 | 48 | FKKLGIPGPTPLPFLG | F108C_PGM | 0 | 0 | 0 |
| 33 | 48 | FKKLGIPGPTPLPFLG | F108C_PGM | 0.5 | 0.3074 | 0.0744 |
| 33 | 48 | FKKLGIPGPTPLPFLG | F108C_PGM | 5 | 0.2623 | 0.1256 |
| 33 | 48 | FKKLGIPGPTPLPFLG | F108C_PGM | 120 | 0.1736 | 0.0875 |
| 33 | 51 | FKKLGIPGPTPLPFLGNIL | F108C | 0 | 0 | 0 |
| 33 | 51 | FKKLGIPGPTPLPFLGNIL | F108C | 0.5 | 0.5156 | 0.143 |
| 33 | 51 | FKKLGIPGPTPLPFLGNIL | F108C | 5 | 0.4249 | 0.1498 |
| 33 | 51 | FKKLGIPGPTPLPFLGNIL | F108C | 120 | 0.493 | 0.1464 |
| 33 | 51 | FKKLGIPGPTPLPFLGNIL | F108C_PGM | 0 | 0 | 0 |
| 33 | 51 | FKKLGIPGPTPLPFLGNIL | F108C_PGM | 0.5 | 0.4208 | 0.0562 |
| 33 | 51 | FKKLGIPGPTPLPFLGNIL | F108C_PGM | 5 | 0.3797 | 0.0387 |
| 33 | 51 | FKKLGIPGPTPLPFLGNIL | F108C_PGM | 120 | 0.3423 | 0.0261 |
| 34 | 48 | KKLGIPGPTPLPFLG | F108C | 0 | 0 | 0 |
| 34 | 48 | KKLGIPGPTPLPFLG | F108C | 0.5 | 0.3683 | 0.0641 |
| 34 | 48 | KKLGIPGPTPLPFLG | F108C | 5 | 0.3399 | 0.0463 |
| 34 | 48 | KKLGIPGPTPLPFLG | F108C | 120 | 0.4272 | 0.0591 |
| 34 | 48 | KKLGIPGPTPLPFLG | F108C_PGM | 0 | 0 | 0 |
| 34 | 48 | KKLGIPGPTPLPFLG | F108C_PGM | 0.5 | 0.2639 | 0.0269 |
| 34 | 48 | KKLGIPGPTPLPFLG | F108C_PGM | 5 | 0.3303 | 0.0697 |
| 34 | 48 | KKLGIPGPTPLPFLG | F108C_PGM | 120 | 0.2693 | 0.0295 |
| 34 | 51 | KKLGIPGPTPLPFLGNIL | F108C | 0 | 0 | 0 |
| 34 | 51 | KKLGIPGPTPLPFLGNIL | F108C | 0.5 | 0.5077 | 0.1035 |
| 34 | 51 | KKLGIPGPTPLPFLGNIL | F108C | 5 | 0.3664 | 0.0175 |
| 34 | 51 | KKLGIPGPTPLPFLGNIL | F108C | 120 | 0.5011 | 0.1853 |
| 34 | 51 | KKLGIPGPTPLPFLGNIL | F108C_PGM | 0 | 0 | 0 |
| 34 | 51 | KKLGIPGPTPLPFLGNIL | F108C_PGM | 0.5 | 0.3933 | 0.0336 |
| 34 | 51 | KKLGIPGPTPLPFLGNIL | F108C_PGM | 5 | 0.4885 | 0.1464 |
| 34 | 51 | KKLGIPGPTPLPFLGNIL | F108C_PGM | 120 | 0.3087 | 0.0725 |
| 49 | 60 | NILSYHKGFTMF | F108C | 0 | 0 | 0 |
| 49 | 60 | NILSYHKGFTMF | F108C | 0.5 | 1.4701 | 0.0716 |
| 49 | 60 | NILSYHKGFTMF | F108C | 5 | 2.0247 | 0.0503 |
| 49 | 60 | NILSYHKGFTMF | F108C | 120 | 2.1166 | 0.034 |
| 49 | 60 | NILSYHKGFTMF | F108C_PGM | 0 | 0 | 0 |
| 49 | 60 | NILSYHKGFTMF | F108C_PGM | 0.5 | 1.2552 | 0.0612 |
| 49 | 60 | NILSYHKGFTMF | F108C_PGM | 5 | 2.0475 | 0.101 |
| 49 | 60 | NILSYHKGFTMF | F108C_PGM | 120 | 2.2728 | 0.0733 |
| 54 | 59 | HKGFTM | F108C | 0 | 0 | 0 |
| 54 | 59 | HKGFTM | F108C | 0.5 | 1.0639 | 0.0671 |

| | | | | | | |
|---|---|---|---|---|---|---|
| 54 | 59 | HKGFTM | F108C | 5 | 1.1222 | 0.0709 |
| 54 | 59 | HKGFTM | F108C | 120 | 1.077 | 0.0816 |
| 54 | 59 | HKGFTM | F108C_PGM | 0 | 0 | 0 |
| 54 | 59 | HKGFTM | F108C_PGM | 0.5 | 0.9899 | 0.0802 |
| 54 | 59 | HKGFTM | F108C_PGM | 5 | 0.9365 | 0.0851 |
| 54 | 59 | HKGFTM | F108C_PGM | 120 | 0.9535 | 0.1426 |
| 61 | 73 | DMEAHKKYGKVWG | F108C | 0 | 0 | 0 |
| 61 | 73 | DMEAHKKYGKVWG | F108C | 0.5 | 0.6569 | 0.0532 |
| 61 | 73 | DMEAHKKYGKVWG | F108C | 5 | 1.0658 | 0.061 |
| 61 | 73 | DMEAHKKYGKVWG | F108C | 120 | 1.9191 | 0.0502 |
| 61 | 73 | DMEAHKKYGKVWG | F108C_PGM | 0 | 0 | 0 |
| 61 | 73 | DMEAHKKYGKVWG | F108C_PGM | 0.5 | 0.6331 | 0.0419 |
| 61 | 73 | DMEAHKKYGKVWG | F108C_PGM | 5 | 0.8335 | 0.0616 |
| 61 | 73 | DMEAHKKYGKVWG | F108C_PGM | 120 | 1.7874 | 0.1472 |
| 61 | 74 | DMEAHKKYGKVWGF | F108C | 0 | 0 | 0 |
| 61 | 74 | DMEAHKKYGKVWGF | F108C | 0.5 | 0.6015 | 0.0785 |
| 61 | 74 | DMEAHKKYGKVWGF | F108C | 5 | 0.947 | 0.0724 |
| 61 | 74 | DMEAHKKYGKVWGF | F108C | 120 | 1.86 | 0.0865 |
| 61 | 74 | DMEAHKKYGKVWGF | F108C_PGM | 0 | 0 | 0 |
| 61 | 74 | DMEAHKKYGKVWGF | F108C_PGM | 0.5 | 0.5722 | 0.0149 |
| 61 | 74 | DMEAHKKYGKVWGF | F108C_PGM | 5 | 0.8261 | 0.013 |
| 61 | 74 | DMEAHKKYGKVWGF | F108C_PGM | 120 | 1.7186 | 0.0733 |
| 61 | 82 | DMEAHKKYGKVWGFYDGQQPVL | F108C | 0 | 0 | 0 |
| 61 | 82 | DMEAHKKYGKVWGFYDGQQPVL | F108C | 0.5 | 1.077 | 0.0713 |
| 61 | 82 | DMEAHKKYGKVWGFYDGQQPVL | F108C | 5 | 1.6282 | 0.061 |
| 61 | 82 | DMEAHKKYGKVWGFYDGQQPVL | F108C | 120 | 3.027 | 0.1244 |
| 61 | 82 | DMEAHKKYGKVWGFYDGQQPVL | F108C_PGM | 0 | 0 | 0 |
| 61 | 82 | DMEAHKKYGKVWGFYDGQQPVL | F108C_PGM | 0.5 | 0.9052 | 0.0427 |
| 61 | 82 | DMEAHKKYGKVWGFYDGQQPVL | F108C_PGM | 5 | 1.436 | 0.0367 |
| 61 | 82 | DMEAHKKYGKVWGFYDGQQPVL | F108C_PGM | 120 | 2.7005 | 0.1191 |
| 74 | 82 | FYDGQQPVL | F108C | 0 | 0 | 0 |
| 74 | 82 | FYDGQQPVL | F108C | 0.5 | 0.6584 | 0.0302 |
| 74 | 82 | FYDGQQPVL | F108C | 5 | 1.0195 | 0.0238 |
| 74 | 82 | FYDGQQPVL | F108C | 120 | 1.4034 | 0.0477 |
| 74 | 82 | FYDGQQPVL | F108C_PGM | 0 | 0 | 0 |
| 74 | 82 | FYDGQQPVL | F108C_PGM | 0.5 | 0.6531 | 0.0353 |
| 74 | 82 | FYDGQQPVL | F108C_PGM | 5 | 0.9372 | 0.0363 |
| 74 | 82 | FYDGQQPVL | F108C_PGM | 120 | 1.3411 | 0.0635 |
| 75 | 82 | YDGQQPVL | F108C | 0 | 0 | 0 |
| 75 | 82 | YDGQQPVL | F108C | 0.5 | 0.5344 | 0.0496 |
| 75 | 82 | YDGQQPVL | F108C | 5 | 0.8091 | 0.0557 |
| 75 | 82 | YDGQQPVL | F108C | 120 | 1.1816 | 0.0475 |
| 75 | 82 | YDGQQPVL | F108C_PGM | 0 | 0 | 0 |
| 75 | 82 | YDGQQPVL | F108C_PGM | 0.5 | 0.5283 | 0.0492 |
| 75 | 82 | YDGQQPVL | F108C_PGM | 5 | 0.7553 | 0.0676 |

| | | | | | | |
|---|---|---|---|---|---|---|
| 75 | 82 | YDGQQPVL | F108C_PGM | 120 | 1.173 | 0.0592 |
| 83 | 89 | AITDPDM | F108C | 0 | 0 | 0 |
| 83 | 89 | AITDPDM | F108C | 0.5 | 0.1912 | 0.0169 |
| 83 | 89 | AITDPDM | F108C | 5 | 0.4467 | 0.0303 |
| 83 | 89 | AITDPDM | F108C | 120 | 0.6401 | 0.0212 |
| 83 | 89 | AITDPDM | F108C_PGM | 0 | 0 | 0 |
| 83 | 89 | AITDPDM | F108C_PGM | 0.5 | 0.2545 | 0.0361 |
| 83 | 89 | AITDPDM | F108C_PGM | 5 | 0.4904 | 0.03 |
| 83 | 89 | AITDPDM | F108C_PGM | 120 | 0.7002 | 0.0568 |
| 83 | 94 | AITDPDMIKTVL | F108C | 0 | 0 | 0 |
| 83 | 94 | AITDPDMIKTVL | F108C | 0.5 | 0.5864 | 0.0366 |
| 83 | 94 | AITDPDMIKTVL | F108C | 5 | 1.2831 | 0.0423 |
| 83 | 94 | AITDPDMIKTVL | F108C | 120 | 1.9512 | 0.0214 |
| 83 | 94 | AITDPDMIKTVL | F108C_PGM | 0 | 0 | 0 |
| 83 | 94 | AITDPDMIKTVL | F108C_PGM | 0.5 | 0.5209 | 0.1 |
| 83 | 94 | AITDPDMIKTVL | F108C_PGM | 5 | 1.1521 | 0.066 |
| 83 | 94 | AITDPDMIKTVL | F108C_PGM | 120 | 1.8541 | 0.0334 |
| 93 | 102 | VLVKESYSVF | F108C | 0 | 0 | 0 |
| 93 | 102 | VLVKESYSVF | F108C | 0.5 | 2.3722 | 0.0918 |
| 93 | 102 | VLVKESYSVF | F108C | 5 | 2.8003 | 0.0459 |
| 93 | 102 | VLVKESYSVF | F108C | 120 | 2.9982 | 0.0283 |
| 93 | 102 | VLVKESYSVF | F108C_PGM | 0 | 0 | 0 |
| 93 | 102 | VLVKESYSVF | F108C_PGM | 0.5 | 2.1653 | 0.0227 |
| 93 | 102 | VLVKESYSVF | F108C_PGM | 5 | 2.5446 | 0.0654 |
| 93 | 102 | VLVKESYSVF | F108C_PGM | 120 | 2.7561 | 0.0784 |
| 94 | 106 | LVKESYSVFTNRR | F108C | 0 | 0 | 0 |
| 94 | 106 | LVKESYSVFTNRR | F108C | 0.5 | 1.5406 | 0.1016 |
| 94 | 106 | LVKESYSVFTNRR | F108C | 5 | 2.0999 | 0.081 |
| 94 | 106 | LVKESYSVFTNRR | F108C | 120 | 2.5581 | 0.0707 |
| 94 | 106 | LVKESYSVFTNRR | F108C_PGM | 0 | 0 | 0 |
| 94 | 106 | LVKESYSVFTNRR | F108C_PGM | 0.5 | 1.5688 | 0.0355 |
| 94 | 106 | LVKESYSVFTNRR | F108C_PGM | 5 | 2.1179 | 0.1105 |
| 94 | 106 | LVKESYSVFTNRR | F108C_PGM | 120 | 2.4385 | 0.0079 |
| 95 | 102 | VKESYSVF | F108C | 0 | 0 | 0 |
| 95 | 102 | VKESYSVF | F108C | 0.5 | 1.4272 | 0.0222 |
| 95 | 102 | VKESYSVF | F108C | 5 | 1.5886 | 0.0292 |
| 95 | 102 | VKESYSVF | F108C | 120 | 1.7458 | 0.0294 |
| 95 | 102 | VKESYSVF | F108C_PGM | 0 | 0 | 0 |
| 95 | 102 | VKESYSVF | F108C_PGM | 0.5 | 1.4414 | 0.0524 |
| 95 | 102 | VKESYSVF | F108C_PGM | 5 | 1.6486 | 0.0472 |
| 95 | 102 | VKESYSVF | F108C_PGM | 120 | 1.8254 | 0.0348 |
| 114 | 122 | MKSAISIAE | F108C | 0 | 0 | 0 |
| 114 | 122 | MKSAISIAE | F108C | 0.5 | 0.8475 | 0.074 |
| 114 | 122 | MKSAISIAE | F108C | 5 | 1.4968 | 0.0364 |
| 114 | 122 | MKSAISIAE | F108C | 120 | 2.8935 | 0.0522 |

| | | | | | | |
|---|---|---|---|---|---|---|
| 114 | 122 | MKSAISIAE | F108C_PGM | 0 | 0 | 0 |
| 114 | 122 | MKSAISIAE | F108C_PGM | 0.5 | 0.7041 | 0.011 |
| 114 | 122 | MKSAISIAE | F108C_PGM | 5 | 1.3494 | 0.0199 |
| 114 | 122 | MKSAISIAE | F108C_PGM | 120 | 2.9796 | 0.0259 |
| 120 | 125 | IAEDEE | F108C | 0 | 0 | 0 |
| 120 | 125 | IAEDEE | F108C | 0.5 | 0.6051 | 0.0412 |
| 120 | 125 | IAEDEE | F108C | 5 | 0.8768 | 0.0279 |
| 120 | 125 | IAEDEE | F108C | 120 | 0.9797 | 0.0334 |
| 120 | 125 | IAEDEE | F108C_PGM | 0 | 0 | 0 |
| 120 | 125 | IAEDEE | F108C_PGM | 0.5 | 0.5791 | 0.0302 |
| 120 | 125 | IAEDEE | F108C_PGM | 5 | 0.8666 | 0.0345 |
| 120 | 125 | IAEDEE | F108C_PGM | 120 | 1.0631 | 0.0321 |
| 123 | 133 | DEEWKRLRSLL | F108C | 0 | 0 | 0 |
| 123 | 133 | DEEWKRLRSLL | F108C | 0.5 | 1.0517 | 0.0656 |
| 123 | 133 | DEEWKRLRSLL | F108C | 5 | 1.7224 | 0.0801 |
| 123 | 133 | DEEWKRLRSLL | F108C | 120 | 2.5265 | 0.0912 |
| 123 | 133 | DEEWKRLRSLL | F108C_PGM | 0 | 0 | 0 |
| 123 | 133 | DEEWKRLRSLL | F108C_PGM | 0.5 | 1.1457 | 0.0585 |
| 123 | 133 | DEEWKRLRSLL | F108C_PGM | 5 | 2.1512 | 0.0436 |
| 123 | 133 | DEEWKRLRSLL | F108C_PGM | 120 | 3.7915 | 0.0246 |
| 123 | 137 | DEEWKRLRSLLSPTF | F108C | 0 | 0 | 0 |
| 123 | 137 | DEEWKRLRSLLSPTF | F108C | 0.5 | 1.2019 | 0.0502 |
| 123 | 137 | DEEWKRLRSLLSPTF | F108C | 5 | 2.3677 | 0.0969 |
| 123 | 137 | DEEWKRLRSLLSPTF | F108C | 120 | 3.948 | 0.0622 |
| 123 | 137 | DEEWKRLRSLLSPTF | F108C_PGM | 0 | 0 | 0 |
| 123 | 137 | DEEWKRLRSLLSPTF | F108C_PGM | 0.5 | 1.2019 | 0.0502 |
| 123 | 137 | DEEWKRLRSLLSPTF | F108C_PGM | 5 | 2.3677 | 0.0969 |
| 123 | 137 | DEEWKRLRSLLSPTF | F108C_PGM | 120 | 3.948 | 0.0622 |
| 126 | 137 | WKRLRSLLSPTF | F108C | 0 | 0 | 0 |
| 126 | 137 | WKRLRSLLSPTF | F108C | 0.5 | 0.9211 | 0.0343 |
| 126 | 137 | WKRLRSLLSPTF | F108C | 5 | 1.6025 | 0.0497 |
| 126 | 137 | WKRLRSLLSPTF | F108C | 120 | 2.5633 | 0.044 |
| 126 | 137 | WKRLRSLLSPTF | F108C_PGM | 0 | 0 | 0 |
| 126 | 137 | WKRLRSLLSPTF | F108C_PGM | 0.5 | 0.9078 | 0.028 |
| 126 | 137 | WKRLRSLLSPTF | F108C_PGM | 5 | 1.7155 | 0.0505 |
| 126 | 137 | WKRLRSLLSPTF | F108C_PGM | 120 | 2.5439 | 0.0812 |
| 135 | 151 | PTFTSGKLKEMVPIIAQ | F108C | 0 | 0 | 0 |
| 135 | 151 | PTFTSGKLKEMVPIIAQ | F108C | 0.5 | 2.0037 | 0.0034 |
| 135 | 151 | PTFTSGKLKEMVPIIAQ | F108C | 5 | 3.3989 | 0.012 |
| 135 | 151 | PTFTSGKLKEMVPIIAQ | F108C | 120 | 5.3008 | 0.0012 |
| 135 | 151 | PTFTSGKLKEMVPIIAQ | F108C_PGM | 0 | 0 | 0 |
| 135 | 151 | PTFTSGKLKEMVPIIAQ | F108C_PGM | 0.5 | 1.8189 | 0.0798 |
| 135 | 151 | PTFTSGKLKEMVPIIAQ | F108C_PGM | 5 | 3.3103 | 0.0855 |
| 135 | 151 | PTFTSGKLKEMVPIIAQ | F108C_PGM | 120 | 5.0548 | 0.0702 |
| 138 | 151 | TSGKLKEMVPIIAQ | F108C | 0 | 0 | 0 |

| | | | | | | |
|---|---|---|---|---|---|---|
| 138 | 151 | TSGKLKEMVPIIAQ | F108C | 0.5 | 1.6571 | 0.1105 |
| 138 | 151 | TSGKLKEMVPIIAQ | F108C | 5 | 2.8443 | 0.0684 |
| 138 | 151 | TSGKLKEMVPIIAQ | F108C | 120 | 4.5038 | 0.0715 |
| 138 | 151 | TSGKLKEMVPIIAQ | F108C_PGM | 0 | 0 | 0 |
| 138 | 151 | TSGKLKEMVPIIAQ | F108C_PGM | 0.5 | 1.4667 | 0.0653 |
| 138 | 151 | TSGKLKEMVPIIAQ | F108C_PGM | 5 | 2.7219 | 0.0408 |
| 138 | 151 | TSGKLKEMVPIIAQ | F108C_PGM | 120 | 4.4584 | 0.0481 |
| 143 | 151 | KEMVPIIAQ | F108C | 0 | 0 | 0 |
| 143 | 151 | KEMVPIIAQ | F108C | 0.5 | 1.236 | 0.0617 |
| 143 | 151 | KEMVPIIAQ | F108C | 5 | 1.6052 | 0.0214 |
| 143 | 151 | KEMVPIIAQ | F108C | 120 | 2.8571 | 0.0348 |
| 143 | 151 | KEMVPIIAQ | F108C_PGM | 0 | 0 | 0 |
| 143 | 151 | KEMVPIIAQ | F108C_PGM | 0.5 | 1.2709 | 0.0262 |
| 143 | 151 | KEMVPIIAQ | F108C_PGM | 5 | 1.7257 | 0.0591 |
| 143 | 151 | KEMVPIIAQ | F108C_PGM | 120 | 2.8047 | 0.1544 |
| 157 | 176 | VRNLRREAETGKPVTLKDVF | F108C | 0 | 0 | 0 |
| 157 | 176 | VRNLRREAETGKPVTLKDVF | F108C | 0.5 | 4.1298 | 0.2486 |
| 157 | 176 | VRNLRREAETGKPVTLKDVF | F108C | 5 | 5.8297 | 0.1323 |
| 157 | 176 | VRNLRREAETGKPVTLKDVF | F108C | 120 | 6.7588 | 0.088 |
| 157 | 176 | VRNLRREAETGKPVTLKDVF | F108C_PGM | 0 | 0 | 0 |
| 157 | 176 | VRNLRREAETGKPVTLKDVF | F108C_PGM | 0.5 | 3.891 | 0.0416 |
| 157 | 176 | VRNLRREAETGKPVTLKDVF | F108C_PGM | 5 | 5.3834 | 0.1129 |
| 157 | 176 | VRNLRREAETGKPVTLKDVF | F108C_PGM | 120 | 6.5435 | 0.217 |
| 173 | 178 | KDVFGA | F108C | 0 | 0 | 0 |
| 173 | 178 | KDVFGA | F108C | 0.5 | 0.6045 | 0.0302 |
| 173 | 178 | KDVFGA | F108C | 5 | 1.1264 | 0.0285 |
| 173 | 178 | KDVFGA | F108C | 120 | 1.7127 | 0.0139 |
| 173 | 178 | KDVFGA | F108C_PGM | 0 | 0 | 0 |
| 173 | 178 | KDVFGA | F108C_PGM | 0.5 | 0.4893 | 0.0328 |
| 173 | 178 | KDVFGA | F108C_PGM | 5 | 0.9131 | 0.0173 |
| 173 | 178 | KDVFGA | F108C_PGM | 120 | 1.5004 | 0.0396 |
| 173 | 181 | KDVFGAYSM | F108C | 0 | 0 | 0 |
| 173 | 181 | KDVFGAYSM | F108C | 0.5 | 0.7982 | 0.0381 |
| 173 | 181 | KDVFGAYSM | F108C | 5 | 1.3216 | 0.0626 |
| 173 | 181 | KDVFGAYSM | F108C | 120 | 2.36 | 0.0295 |
| 173 | 181 | KDVFGAYSM | F108C_PGM | 0 | 0 | 0 |
| 173 | 181 | KDVFGAYSM | F108C_PGM | 0.5 | 0.8184 | 0.0403 |
| 173 | 181 | KDVFGAYSM | F108C_PGM | 5 | 1.297 | 0.0215 |
| 173 | 181 | KDVFGAYSM | F108C_PGM | 120 | 2.1804 | 0.0412 |
| 177 | 182 | GAYSMD | F108C | 0 | 0 | 0 |
| 177 | 182 | GAYSMD | F108C | 0.5 | 0.3264 | 0.0338 |
| 177 | 182 | GAYSMD | F108C | 5 | 0.4337 | 0.0248 |
| 177 | 182 | GAYSMD | F108C | 120 | 1.0134 | 0.04 |
| 177 | 182 | GAYSMD | F108C_PGM | 0 | 0 | 0 |
| 177 | 182 | GAYSMD | F108C_PGM | 0.5 | 0.3128 | 0.0354 |

| | | | | | | |
|---|---|---|---|---|---|---|
| 177 | 182 | GAYSMD | F108C_PGM | 5 | 0.3953 | 0.0281 |
| 177 | 182 | GAYSMD | F108C_PGM | 120 | 0.7422 | 0.0305 |
| 182 | 189 | DVITSTSF | F108C | 0 | 0 | 0 |
| 182 | 189 | DVITSTSF | F108C | 0.5 | 0.7445 | 0.0358 |
| 182 | 189 | DVITSTSF | F108C | 5 | 0.9409 | 0.0332 |
| 182 | 189 | DVITSTSF | F108C | 120 | 1.8279 | 0.0255 |
| 182 | 189 | DVITSTSF | F108C_PGM | 0 | 0 | 0 |
| 182 | 189 | DVITSTSF | F108C_PGM | 0.5 | 0.7631 | 0.0284 |
| 182 | 189 | DVITSTSF | F108C_PGM | 5 | 0.9183 | 0.0162 |
| 182 | 189 | DVITSTSF | F108C_PGM | 120 | 1.7717 | 0.0492 |
| 183 | 189 | VITSTSFGVN | F108C | 0 | 0 | 0 |
| 183 | 189 | VITSTSFGVN | F108C | 0.5 | 0.5415 | 0.0131 |
| 183 | 189 | VITSTSFGVN | F108C | 5 | 0.7205 | 0.0055 |
| 183 | 189 | VITSTSFGVN | F108C | 120 | 1.2895 | 0.0206 |
| 183 | 189 | VITSTSFGVN | F108C_PGM | 0 | 0 | 0 |
| 183 | 189 | VITSTSFGVN | F108C_PGM | 0.5 | 0.5614 | 0.0322 |
| 183 | 189 | VITSTSFGVN | F108C_PGM | 5 | 0.7183 | 0.0285 |
| 183 | 189 | VITSTSFGVN | F108C_PGM | 120 | 1.3372 | 0.0612 |
| 189 | 212 | FGVNIDSLNNPQDPFVENTKKLLR | F108C | 0 | 0 | 0 |
| 189 | 212 | FGVNIDSLNNPQDPFVENTKKLLR | F108C | 0.5 | 4.072 | 0.1776 |
| 189 | 212 | FGVNIDSLNNPQDPFVENTKKLLR | F108C | 5 | 6.2766 | 0.0696 |
| 189 | 212 | FGVNIDSLNNPQDPFVENTKKLLR | F108C | 120 | 7.7176 | 0.1631 |
| 189 | 212 | FGVNIDSLNNPQDPFVENTKKLLR | F108C_PGM | 0 | 0 | 0 |
| 189 | 212 | FGVNIDSLNNPQDPFVENTKKLLR | F108C_PGM | 0.5 | 5.315 | 0.1159 |
| 189 | 212 | FGVNIDSLNNPQDPFVENTKKLLR | F108C_PGM | 5 | 6.718 | 0.1277 |
| 189 | 212 | FGVNIDSLNNPQDPFVENTKKLLR | F108C_PGM | 120 | 7.9355 | 0.2075 |
| 190 | 210 | GVNIDSLNNPQDPFVENTKKL | F108C | 0 | 0 | 0 |
| 190 | 210 | GVNIDSLNNPQDPFVENTKKL | F108C | 0.5 | 3.7755 | 0.2328 |
| 190 | 210 | GVNIDSLNNPQDPFVENTKKL | F108C | 5 | 5.6398 | 0.1984 |
| 190 | 210 | GVNIDSLNNPQDPFVENTKKL | F108C | 120 | 6.4664 | 0.1587 |
| 190 | 210 | GVNIDSLNNPQDPFVENTKKL | F108C_PGM | 0 | 0 | 0 |
| 190 | 210 | GVNIDSLNNPQDPFVENTKKL | F108C_PGM | 0.5 | 3.7256 | 0.1828 |
| 190 | 210 | GVNIDSLNNPQDPFVENTKKL | F108C_PGM | 5 | 5.6404 | 0.2085 |
| 190 | 210 | GVNIDSLNNPQDPFVENTKKL | F108C_PGM | 120 | 6.5511 | 0.1939 |
| 193 | 210 | IDSLNNPQDPFVENTKKL | F108C | 0 | 0 | 0 |
| 193 | 210 | IDSLNNPQDPFVENTKKL | F108C | 0.5 | 2.2983 | 0.1122 |
| 193 | 210 | IDSLNNPQDPFVENTKKL | F108C | 5 | 3.7533 | 0.0318 |
| 193 | 210 | IDSLNNPQDPFVENTKKL | F108C | 120 | 4.7273 | 0.0521 |
| 193 | 210 | IDSLNNPQDPFVENTKKL | F108C_PGM | 0 | 0 | 0 |
| 193 | 210 | IDSLNNPQDPFVENTKKL | F108C_PGM | 0.5 | 2.4012 | 0.0571 |
| 193 | 210 | IDSLNNPQDPFVENTKKL | F108C_PGM | 5 | 3.8858 | 0.1756 |
| 193 | 210 | IDSLNNPQDPFVENTKKL | F108C_PGM | 120 | 4.8334 | 0.17 |
| 221 | 226 | LSITVF | F108C | 0 | 0 | 0 |
| 221 | 226 | LSITVF | F108C | 0.5 | 0.3861 | 0.0834 |
| 221 | 226 | LSITVF | F108C | 5 | 1.0718 | 0.0412 |

| | | | | | | |
|---|---|---|---|---|---|---|
| 221 | 226 | LSITVF | F108C | 120 | 1.9555 | 0.033 |
| 221 | 226 | LSITVF | F108C_PGM | 0 | 0 | 0 |
| 221 | 226 | LSITVF | F108C_PGM | 0.5 | 0.1125 | 0.0343 |
| 221 | 226 | LSITVF | F108C_PGM | 5 | 0.5743 | 0.0332 |
| 221 | 226 | LSITVF | F108C_PGM | 120 | 1.4792 | 0.0498 |
| 230 | 235 | IPILEV | F108C | 0 | 0 | 0 |
| 230 | 235 | IPILEV | F108C | 0.5 | 0.1509 | 0.0192 |
| 230 | 235 | IPILEV | F108C | 5 | 0.4861 | 0.0441 |
| 230 | 235 | IPILEV | F108C | 120 | 1.8512 | 0.0324 |
| 230 | 235 | IPILEV | F108C_PGM | 0 | 0 | 0 |
| 230 | 235 | IPILEV | F108C_PGM | 0.5 | 0.0561 | 0.0271 |
| 230 | 235 | IPILEV | F108C_PGM | 5 | 0.3384 | 0.0447 |
| 230 | 235 | IPILEV | F108C_PGM | 120 | 1.2451 | 0.0367 |
| 233 | 247 | LEVLNISVFPREVTN | F108C | 0 | 0 | 0 |
| 233 | 247 | LEVLNISVFPREVTN | F108C | 0.5 | 6.6763 | 0.095 |
| 233 | 247 | LEVLNISVFPREVTN | F108C | 5 | 6.8348 | 0.0393 |
| 233 | 247 | LEVLNISVFPREVTN | F108C | 120 | 6.9938 | 0.0524 |
| 233 | 247 | LEVLNISVFPREVTN | F108C_PGM | 0 | 0 | 0 |
| 233 | 247 | LEVLNISVFPREVTN | F108C_PGM | 0.5 | 6.7672 | 0.0332 |
| 233 | 247 | LEVLNISVFPREVTN | F108C_PGM | 5 | 6.8003 | 0.0915 |
| 233 | 247 | LEVLNISVFPREVTN | F108C_PGM | 120 | 6.8986 | 0.2185 |
| 235 | 241 | VLNISVF | F108C | 0 | 0 | 0 |
| 235 | 241 | VLNISVF | F108C | 0.5 | 1.2824 | 0.0861 |
| 235 | 241 | VLNISVF | F108C | 5 | 2.2499 | 0.0263 |
| 235 | 241 | VLNISVF | F108C | 120 | 2.9213 | 0.0406 |
| 235 | 241 | VLNISVF | F108C_PGM | 0 | 0 | 0 |
| 235 | 241 | VLNISVF | F108C_PGM | 0.5 | 0.9839 | 0.0081 |
| 235 | 241 | VLNISVF | F108C_PGM | 5 | 1.9289 | 0.0498 |
| 235 | 241 | VLNISVF | F108C_PGM | 120 | 2.9684 | 0.0316 |
| 236 | 241 | LNISVF | F108C | 0 | 0 | 0 |
| 236 | 241 | LNISVF | F108C | 0.5 | 1.0347 | 0.0742 |
| 236 | 241 | LNISVF | F108C | 5 | 1.7577 | 0.015 |
| 236 | 241 | LNISVF | F108C | 120 | 2.3012 | 0.0278 |
| 236 | 241 | LNISVF | F108C_PGM | 0 | 0 | 0 |
| 236 | 241 | LNISVF | F108C_PGM | 0.5 | 0.7753 | 0.0134 |
| 236 | 241 | LNISVF | F108C_PGM | 5 | 1.5011 | 0.0583 |
| 236 | 241 | LNISVF | F108C_PGM | 120 | 2.3407 | 0.0145 |
| 237 | 248 | NISVFPREVTNF | F108C | 0 | 0 | 0 |
| 237 | 248 | NISVFPREVTNF | F108C | 0.5 | 2.4107 | 0.1161 |
| 237 | 248 | NISVFPREVTNF | F108C | 5 | 3.3715 | 0.0446 |
| 237 | 248 | NISVFPREVTNF | F108C | 120 | 3.6199 | 0.0621 |
| 237 | 248 | NISVFPREVTNF | F108C_PGM | 0 | 0 | 0 |
| 237 | 248 | NISVFPREVTNF | F108C_PGM | 0.5 | 2.237 | 0.1036 |
| 237 | 248 | NISVFPREVTNF | F108C_PGM | 5 | 3.2374 | 0.0979 |
| 237 | 248 | NISVFPREVTNF | F108C_PGM | 120 | 3.8404 | 0.0829 |

| | | | | | | |
|---|---|---|---|---|---|---|
| 249 | 261 | LRKSVKRMKESRL | F108C | 0 | 0 | 0 |
| 249 | 261 | LRKSVKRMKESRL | F108C | 0.5 | 1.0545 | 0.0915 |
| 249 | 261 | LRKSVKRMKESRL | F108C | 5 | 1.6779 | 0.0483 |
| 249 | 261 | LRKSVKRMKESRL | F108C | 120 | 1.8006 | 0.0427 |
| 249 | 261 | LRKSVKRMKESRL | F108C_PGM | 0 | 0 | 0 |
| 249 | 261 | LRKSVKRMKESRL | F108C_PGM | 0.5 | 0.48 | 0.0645 |
| 249 | 261 | LRKSVKRMKESRL | F108C_PGM | 5 | 0.9153 | 0.1046 |
| 249 | 261 | LRKSVKRMKESRL | F108C_PGM | 120 | 1.436 | 0.0053 |
| 262 | 271 | EDTQKHRVDF | F108C | 0 | 0 | 0 |
| 262 | 271 | EDTQKHRVDF | F108C | 0.5 | 0.7534 | 0.031 |
| 262 | 271 | EDTQKHRVDF | F108C | 5 | 0.8793 | 0.025 |
| 262 | 271 | EDTQKHRVDF | F108C | 120 | 1.0361 | 0.0194 |
| 262 | 271 | EDTQKHRVDF | F108C_PGM | 0 | 0 | 0 |
| 262 | 271 | EDTQKHRVDF | F108C_PGM | 0.5 | 0.7274 | 0.0343 |
| 262 | 271 | EDTQKHRVDF | F108C_PGM | 5 | 0.7803 | 0.044 |
| 262 | 271 | EDTQKHRVDF | F108C_PGM | 120 | 0.9768 | 0.1053 |
| 275 | 295 | MIDSQNSKETESHKALSDLEL | F108C | 0 | 0 | 0 |
| 275 | 295 | MIDSQNSKETESHKALSDLEL | F108C | 0.5 | 2.6402 | 0.0672 |
| 275 | 295 | MIDSQNSKETESHKALSDLEL | F108C | 5 | 3.591 | 0.0418 |
| 275 | 295 | MIDSQNSKETESHKALSDLEL | F108C | 120 | 4.1162 | 0.0314 |
| 275 | 295 | MIDSQNSKETESHKALSDLEL | F108C_PGM | 0 | 0 | 0 |
| 275 | 295 | MIDSQNSKETESHKALSDLEL | F108C_PGM | 0.5 | 2.9187 | 0.0855 |
| 275 | 295 | MIDSQNSKETESHKALSDLEL | F108C_PGM | 5 | 3.9043 | 0.0257 |
| 275 | 295 | MIDSQNSKETESHKALSDLEL | F108C_PGM | 120 | 4.2838 | 0.1592 |
| 296 | 302 | VAQSIIF | F108C | 0 | 0 | 0 |
| 296 | 302 | VAQSIIF | F108C | 0.5 | -0.2649 | 0.0761 |
| 296 | 302 | VAQSIIF | F108C | 5 | 0.0497 | 0.0657 |
| 296 | 302 | VAQSIIF | F108C | 120 | 0.6046 | 0.0612 |
| 296 | 302 | VAQSIIF | F108C_PGM | 0 | 0 | 0 |
| 296 | 302 | VAQSIIF | F108C_PGM | 0.5 | -0.3571 | 0.0568 |
| 296 | 302 | VAQSIIF | F108C_PGM | 5 | -0.1298 | 0.0513 |
| 296 | 302 | VAQSIIF | F108C_PGM | 120 | 0.4709 | 0.0612 |
| 301 | 306 | IFIFAG | F108C | 0 | 0 | 0 |
| 301 | 306 | IFIFAG | F108C | 0.5 | 0.0964 | 0.0289 |
| 301 | 306 | IFIFAG | F108C | 5 | 0.2045 | 0.0338 |
| 301 | 306 | IFIFAG | F108C | 120 | 0.5371 | 0.0251 |
| 301 | 306 | IFIFAG | F108C_PGM | 0 | 0 | 0 |
| 301 | 306 | IFIFAG | F108C_PGM | 0.5 | 0.0632 | 0.014 |
| 301 | 306 | IFIFAG | F108C_PGM | 5 | 0.1307 | 0.0157 |
| 301 | 306 | IFIFAG | F108C_PGM | 120 | 0.3808 | 0.083 |
| 307 | 313 | YETTSSV | F108C | 0 | 0 | 0 |
| 307 | 313 | YETTSSV | F108C | 0.5 | 0.4525 | 0.0528 |
| 307 | 313 | YETTSSV | F108C | 5 | 0.9379 | 0.0305 |
| 307 | 313 | YETTSSV | F108C | 120 | 1.4727 | 0.0261 |
| 307 | 313 | YETTSSV | F108C_PGM | 0 | 0 | 0 |

| | | | | | | |
|---|---|---|---|---|---|---|
| 307 | 313 | YETTSSV | F108C_PGM | 0.5 | 0.2877 | 0.0156 |
| 307 | 313 | YETTSSV | F108C_PGM | 5 | 0.7863 | 0.0236 |
| 307 | 313 | YETTSSV | F108C_PGM | 120 | 1.6458 | 0.1507 |
| 307 | 314 | YETTSSVL | F108C | 0 | 0 | 0 |
| 307 | 314 | YETTSSVL | F108C | 0.5 | 0.3413 | 0.0389 |
| 307 | 314 | YETTSSVL | F108C | 5 | 0.6338 | 0.0084 |
| 307 | 314 | YETTSSVL | F108C | 120 | 1.1111 | 0.0217 |
| 307 | 314 | YETTSSVL | F108C_PGM | 0 | 0 | 0 |
| 307 | 314 | YETTSSVL | F108C_PGM | 0.5 | 0.2974 | 0.0223 |
| 307 | 314 | YETTSSVL | F108C_PGM | 5 | 0.6791 | 0.0386 |
| 307 | 314 | YETTSSVL | F108C_PGM | 120 | 1.3109 | 0.0218 |
| 307 | 316 | YETTSSVLSF | F108C | 0 | 0 | 0 |
| 307 | 316 | YETTSSVLSF | F108C | 0.5 | 0.3186 | 0.0262 |
| 307 | 316 | YETTSSVLSF | F108C | 5 | 0.7198 | 0.0236 |
| 307 | 316 | YETTSSVLSF | F108C | 120 | 1.437 | 0.0246 |
| 307 | 316 | YETTSSVLSF | F108C_PGM | 0 | 0 | 0 |
| 307 | 316 | YETTSSVLSF | F108C_PGM | 0.5 | 0.2527 | 0.021 |
| 307 | 316 | YETTSSVLSF | F108C_PGM | 5 | 0.6319 | 0.0493 |
| 307 | 316 | YETTSSVLSF | F108C_PGM | 120 | 1.4742 | 0.0153 |
| 319 | 333 | YELATHPDVQQKLQE | F108C | 0 | 0 | 0 |
| 319 | 333 | YELATHPDVQQKLQE | F108C | 0.5 | 0.0833 | 0.0386 |
| 319 | 333 | YELATHPDVQQKLQE | F108C | 5 | 0.3572 | 0.0527 |
| 319 | 333 | YELATHPDVQQKLQE | F108C | 120 | 1.0518 | 0.0553 |
| 319 | 333 | YELATHPDVQQKLQE | F108C_PGM | 0 | 0 | 0 |
| 319 | 333 | YELATHPDVQQKLQE | F108C_PGM | 0.5 | 0.1328 | 0.108 |
| 319 | 333 | YELATHPDVQQKLQE | F108C_PGM | 5 | 0.4667 | 0.065 |
| 319 | 333 | YELATHPDVQQKLQE | F108C_PGM | 120 | 1.2543 | 0.0514 |
| 322 | 333 | ATHPDVQQKLQE | F108C | 0 | 0 | 0 |
| 322 | 333 | ATHPDVQQKLQE | F108C | 0.5 | -0.1716 | 0.0387 |
| 322 | 333 | ATHPDVQQKLQE | F108C | 5 | 0.1422 | 0.0323 |
| 322 | 333 | ATHPDVQQKLQE | F108C | 120 | 0.622 | 0.0303 |
| 322 | 333 | ATHPDVQQKLQE | F108C_PGM | 0 | 0 | 0 |
| 322 | 333 | ATHPDVQQKLQE | F108C_PGM | 0.5 | -0.2177 | 0.0321 |
| 322 | 333 | ATHPDVQQKLQE | F108C_PGM | 5 | -0.015 | 0.0326 |
| 322 | 333 | ATHPDVQQKLQE | F108C_PGM | 120 | 0.6016 | 0.0314 |
| 332 | 337 | QEEIDA | F108C | 0 | 0 | 0 |
| 332 | 337 | QEEIDA | F108C | 0.5 | 0.159 | 0.0477 |
| 332 | 337 | QEEIDA | F108C | 5 | 0.4563 | 0.0197 |
| 332 | 337 | QEEIDA | F108C | 120 | 1.3506 | 0.0217 |
| 332 | 337 | QEEIDA | F108C_PGM | 0 | 0 | 0 |
| 332 | 337 | QEEIDA | F108C_PGM | 0.5 | 0.069 | 0.0299 |
| 332 | 337 | QEEIDA | F108C_PGM | 5 | 0.4359 | 0.0233 |
| 332 | 337 | QEEIDA | F108C_PGM | 120 | 1.374 | 0.0784 |
| 337 | 349 | AVLPNKAPPTYDT | F108C | 0 | 0 | 0 |
| 337 | 349 | AVLPNKAPPTYDT | F108C | 0.5 | 1.8503 | 0.05 |

| | | | | | | |
|---|---|---|---|---|---|---|
| 337 | 349 | AVLPNKAPPTYDT | F108C | 5 | 2.6236 | 0.0421 |
| 337 | 349 | AVLPNKAPPTYDT | F108C | 120 | 3.0867 | 0.0142 |
| 337 | 349 | AVLPNKAPPTYDT | F108C_PGM | 0 | 0 | 0 |
| 337 | 349 | AVLPNKAPPTYDT | F108C_PGM | 0.5 | 1.9643 | 0.042 |
| 337 | 349 | AVLPNKAPPTYDT | F108C_PGM | 5 | 2.641 | 0.1164 |
| 337 | 349 | AVLPNKAPPTYDT | F108C_PGM | 120 | 3.1508 | 0.1291 |
| 340 | 347 | PNKAPPTY | F108C | 0 | 0 | 0 |
| 340 | 347 | PNKAPPTY | F108C | 0.5 | 0.9683 | 0.0278 |
| 340 | 347 | PNKAPPTY | F108C | 5 | 1.3699 | 0.0125 |
| 340 | 347 | PNKAPPTY | F108C | 120 | 1.7427 | 0.023 |
| 340 | 347 | PNKAPPTY | F108C_PGM | 0 | 0 | 0 |
| 340 | 347 | PNKAPPTY | F108C_PGM | 0.5 | 0.9342 | 0.0619 |
| 340 | 347 | PNKAPPTY | F108C_PGM | 5 | 1.3039 | 0.071 |
| 340 | 347 | PNKAPPTY | F108C_PGM | 120 | 1.6612 | 0.0909 |
| 353 | 358 | MEYLDM | F108C | 0 | 0 | 0 |
| 353 | 358 | MEYLDM | F108C | 0.5 | 0.259 | 0.0352 |
| 353 | 358 | MEYLDM | F108C | 5 | 0.2745 | 0.0335 |
| 353 | 358 | MEYLDM | F108C | 120 | 0.7477 | 0.0456 |
| 353 | 358 | MEYLDM | F108C_PGM | 0 | 0 | 0 |
| 353 | 358 | MEYLDM | F108C_PGM | 0.5 | 0.135 | 0.0999 |
| 353 | 358 | MEYLDM | F108C_PGM | 5 | 0.0048 | 0.0584 |
| 353 | 358 | MEYLDM | F108C_PGM | 120 | 0.5357 | 0.0537 |
| 357 | 363 | DMVVNET | F108C | 0 | 0 | 0 |
| 357 | 363 | DMVVNET | F108C | 0.5 | -0.0236 | 0.0259 |
| 357 | 363 | DMVVNET | F108C | 5 | -0.0007 | 0.044 |
| 357 | 363 | DMVVNET | F108C | 120 | 0.0737 | 0.0417 |
| 357 | 363 | DMVVNET | F108C_PGM | 0 | 0 | 0 |
| 357 | 363 | DMVVNET | F108C_PGM | 0.5 | -0.0234 | 0.0483 |
| 357 | 363 | DMVVNET | F108C_PGM | 5 | -0.0143 | 0.0383 |
| 357 | 363 | DMVVNET | F108C_PGM | 120 | -0.0073 | 0.0241 |
| 359 | 366 | VVNETLRL | F108C | 0 | 0 | 0 |
| 359 | 366 | VVNETLRL | F108C | 0.5 | 0.0247 | 0.0247 |
| 359 | 366 | VVNETLRL | F108C | 5 | 0.1021 | 0.0397 |
| 359 | 366 | VVNETLRL | F108C | 120 | 0.2754 | 0.0691 |
| 359 | 366 | VVNETLRL | F108C_PGM | 0 | 0 | 0 |
| 359 | 366 | VVNETLRL | F108C_PGM | 0.5 | 0.0324 | 0.015 |
| 359 | 366 | VVNETLRL | F108C_PGM | 5 | 0.059 | 0.0086 |
| 359 | 366 | VVNETLRL | F108C_PGM | 120 | 0.1803 | 0.0271 |
| 363 | 370 | TLRLFPIA | F108C | 0 | 0 | 0 |
| 363 | 370 | TLRLFPIA | F108C | 0.5 | 0.1179 | 0.0191 |
| 363 | 370 | TLRLFPIA | F108C | 5 | 0.2334 | 0.0277 |
| 363 | 370 | TLRLFPIA | F108C | 120 | 0.669 | 0.0358 |
| 363 | 370 | TLRLFPIA | F108C_PGM | 0 | 0 | 0 |
| 363 | 370 | TLRLFPIA | F108C_PGM | 0.5 | 0.043 | 0.0571 |
| 363 | 370 | TLRLFPIA | F108C_PGM | 5 | 0.1584 | 0.0361 |

| | | | | | | |
|---|---|---|---|---|---|---|
| 363 | 370 | TLRLFPIA | F108C_PGM | 120 | 0.3255 | 0.1195 |
| 364 | 371 | LRLFPIAM | F108C | 0 | 0 | 0 |
| 364 | 371 | LRLFPIAM | F108C | 0.5 | 0.2103 | 0.0934 |
| 364 | 371 | LRLFPIAM | F108C | 5 | 0.4198 | 0.0439 |
| 364 | 371 | LRLFPIAM | F108C | 120 | 0.841 | 0.1041 |
| 364 | 371 | LRLFPIAM | F108C_PGM | 0 | 0 | 0 |
| 364 | 371 | LRLFPIAM | F108C_PGM | 0.5 | 0.0348 | 0.135 |
| 364 | 371 | LRLFPIAM | F108C_PGM | 5 | 0.0237 | 0.1182 |
| 364 | 371 | LRLFPIAM | F108C_PGM | 120 | 0.2976 | 0.021 |
| 374 | 385 | ERVCKKDVEING | F108C | 0 | 0 | 0 |
| 374 | 385 | ERVCKKDVEING | F108C | 0.5 | 0.7824 | 0.0481 |
| 374 | 385 | ERVCKKDVEING | F108C | 5 | 1.1898 | 0.0247 |
| 374 | 385 | ERVCKKDVEING | F108C | 120 | 1.3814 | 0.0248 |
| 374 | 385 | ERVCKKDVEING | F108C_PGM | 0 | 0 | 0 |
| 374 | 385 | ERVCKKDVEING | F108C_PGM | 0.5 | 0.5931 | 0.0199 |
| 374 | 385 | ERVCKKDVEING | F108C_PGM | 5 | 0.9733 | 0.0383 |
| 374 | 385 | ERVCKKDVEING | F108C_PGM | 120 | 1.2052 | 0.0706 |
| 387 | 393 | FIPKGVV | F108C | 0 | 0 | 0 |
| 387 | 393 | FIPKGVV | F108C | 0.5 | 0.475 | 0.0321 |
| 387 | 393 | FIPKGVV | F108C | 5 | 0.5907 | 0.0345 |
| 387 | 393 | FIPKGVV | F108C | 120 | 0.7903 | 0.0653 |
| 387 | 393 | FIPKGVV | F108C_PGM | 0 | 0 | 0 |
| 387 | 393 | FIPKGVV | F108C_PGM | 0.5 | 0.4441 | 0.0355 |
| 387 | 393 | FIPKGVV | F108C_PGM | 5 | 0.5664 | 0.0407 |
| 387 | 393 | FIPKGVV | F108C_PGM | 120 | 0.7718 | 0.0332 |
| 388 | 393 | IPKGVV | F108C | 0 | 0 | 0 |
| 388 | 393 | IPKGVV | F108C | 0.5 | 0.3881 | 0.0172 |
| 388 | 393 | IPKGVV | F108C | 5 | 0.5398 | 0.023 |
| 388 | 393 | IPKGVV | F108C | 120 | 0.8537 | 0.0273 |
| 388 | 393 | IPKGVV | F108C_PGM | 0 | 0 | 0 |
| 388 | 393 | IPKGVV | F108C_PGM | 0.5 | 0.3441 | 0.0163 |
| 388 | 393 | IPKGVV | F108C_PGM | 5 | 0.5106 | 0.0185 |
| 388 | 393 | IPKGVV | F108C_PGM | 120 | 0.7997 | 0.0209 |
| 388 | 395 | IPKGVVVM | F108C | 0 | 0 | 0 |
| 388 | 395 | IPKGVVVM | F108C | 0.5 | 0.5778 | 0.0441 |
| 388 | 395 | IPKGVVVM | F108C | 5 | 0.9682 | 0.0373 |
| 388 | 395 | IPKGVVVM | F108C | 120 | 1.3826 | 0.0593 |
| 388 | 395 | IPKGVVVM | F108C_PGM | 0 | 0 | 0 |
| 388 | 395 | IPKGVVVM | F108C_PGM | 0.5 | 0.5275 | 0.0522 |
| 388 | 395 | IPKGVVVM | F108C_PGM | 5 | 0.8282 | 0.0545 |
| 388 | 395 | IPKGVVVM | F108C_PGM | 120 | 1.2245 | 0.0314 |
| 394 | 399 | VMIPSY | F108C | 0 | 0 | 0 |
| 394 | 399 | VMIPSY | F108C | 0.5 | 0.2475 | 0.0206 |
| 394 | 399 | VMIPSY | F108C | 5 | 0.3598 | 0.0271 |
| 394 | 399 | VMIPSY | F108C | 120 | 0.5495 | 0.0394 |

| | | | | | | |
|---|---|---|---|---|---|---|
| 394 | 399 | VMIPSY | F108C_PGM | 0 | 0 | 0 |
| 394 | 399 | VMIPSY | F108C_PGM | 0.5 | 0.2342 | 0.0251 |
| 394 | 399 | VMIPSY | F108C_PGM | 5 | 0.3346 | 0.0386 |
| 394 | 399 | VMIPSY | F108C_PGM | 120 | 0.5385 | 0.0166 |
| 394 | 401 | VMIPSYAL | F108C | 0 | 0 | 0 |
| 394 | 401 | VMIPSYAL | F108C | 0.5 | 0.5447 | 0.0866 |
| 394 | 401 | VMIPSYAL | F108C | 5 | 0.7561 | 0.0499 |
| 394 | 401 | VMIPSYAL | F108C | 120 | 1.0702 | 0.0528 |
| 394 | 401 | VMIPSYAL | F108C_PGM | 0 | 0 | 0 |
| 394 | 401 | VMIPSYAL | F108C_PGM | 0.5 | 0.407 | 0.043 |
| 394 | 401 | VMIPSYAL | F108C_PGM | 5 | 0.6297 | 0.0675 |
| 394 | 401 | VMIPSYAL | F108C_PGM | 120 | 0.8411 | 0.0442 |
| 400 | 407 | ALHRDPKY | F108C | 0 | 0 | 0 |
| 400 | 407 | ALHRDPKY | F108C | 0.5 | 0.0015 | 0.0569 |
| 400 | 407 | ALHRDPKY | F108C | 5 | 0.2344 | 0.0349 |
| 400 | 407 | ALHRDPKY | F108C | 120 | 0.3912 | 0.0413 |
| 400 | 407 | ALHRDPKY | F108C_PGM | 0 | 0 | 0 |
| 400 | 407 | ALHRDPKY | F108C_PGM | 0.5 | -0.0615 | 0.0435 |
| 400 | 407 | ALHRDPKY | F108C_PGM | 5 | 0.1352 | 0.0331 |
| 400 | 407 | ALHRDPKY | F108C_PGM | 120 | 0.3425 | 0.0576 |
| 408 | 414 | WTEPEKF | F108C | 0 | 0 | 0 |
| 408 | 414 | WTEPEKF | F108C | 0.5 | 1.0772 | 0.013 |
| 408 | 414 | WTEPEKF | F108C | 5 | 1.142 | 0.0324 |
| 408 | 414 | WTEPEKF | F108C | 120 | 1.1194 | 0.0207 |
| 408 | 414 | WTEPEKF | F108C_PGM | 0 | 0 | 0 |
| 408 | 414 | WTEPEKF | F108C_PGM | 0.5 | 1.2271 | 0.0167 |
| 408 | 414 | WTEPEKF | F108C_PGM | 5 | 1.1755 | 0.0451 |
| 408 | 414 | WTEPEKF | F108C_PGM | 120 | 1.2451 | 0.0751 |
| 420 | 444 | SKKNKDNIDPYIYTPFGSGPRNCIG | F108C | 0 | 0 | 0 |
| 420 | 444 | SKKNKDNIDPYIYTPFGSGPRNCIG | F108C | 0.5 | 2.3705 | 0.1032 |
| 420 | 444 | SKKNKDNIDPYIYTPFGSGPRNCIG | F108C | 5 | 3.3493 | 0.0622 |
| 420 | 444 | SKKNKDNIDPYIYTPFGSGPRNCIG | F108C | 120 | 4.3471 | 0.0637 |
| 420 | 444 | SKKNKDNIDPYIYTPFGSGPRNCIG | F108C_PGM | 0 | 0 | 0 |
| 420 | 444 | SKKNKDNIDPYIYTPFGSGPRNCIG | F108C_PGM | 0.5 | 2.1988 | 0.0957 |
| 420 | 444 | SKKNKDNIDPYIYTPFGSGPRNCIG | F108C_PGM | 5 | 3.1955 | 0.105 |
| 420 | 444 | SKKNKDNIDPYIYTPFGSGPRNCIG | F108C_PGM | 120 | 4.1987 | 0.0751 |
| 445 | 452 | MRFALMNM | F108C | 0 | 0 | 0 |
| 445 | 452 | MRFALMNM | F108C | 0.5 | -0.0535 | 0.0363 |
| 445 | 452 | MRFALMNM | F108C | 5 | 0.0475 | 0.1196 |
| 445 | 452 | MRFALMNM | F108C | 120 | 0.2553 | 0.0258 |
| 445 | 452 | MRFALMNM | F108C_PGM | 0 | 0 | 0 |
| 445 | 452 | MRFALMNM | F108C_PGM | 0.5 | -0.0132 | 0.0248 |
| 445 | 452 | MRFALMNM | F108C_PGM | 5 | 0.0965 | 0.0147 |
| 445 | 452 | MRFALMNM | F108C_PGM | 120 | 0.2149 | 0.037 |
| 446 | 452 | RFALMNM | F108C | 0 | 0 | 0 |

| | | | | | | |
|---|---|---|---|---|---|---|
| 446 | 452 | RFALMNM | F108C | 0.5 | 0.0721 | 0.0172 |
| 446 | 452 | RFALMNM | F108C | 5 | 0.1645 | 0.0269 |
| 446 | 452 | RFALMNM | F108C | 120 | 0.3835 | 0.0152 |
| 446 | 452 | RFALMNM | F108C_PGM | 0 | 0 | 0 |
| 446 | 452 | RFALMNM | F108C_PGM | 0.5 | 0.0649 | 0.0245 |
| 446 | 452 | RFALMNM | F108C_PGM | 5 | 0.1199 | 0.0329 |
| 446 | 452 | RFALMNM | F108C_PGM | 120 | 0.3923 | 0.022 |
| 455 | 463 | ALIRVLQNF | F108C | 0 | 0 | 0 |
| 455 | 463 | ALIRVLQNF | F108C | 0.5 | -0.021 | 0.0268 |
| 455 | 463 | ALIRVLQNF | F108C | 5 | 0.1082 | 0.0233 |
| 455 | 463 | ALIRVLQNF | F108C | 120 | 0.4529 | 0.0234 |
| 455 | 463 | ALIRVLQNF | F108C_PGM | 0 | 0 | 0 |
| 455 | 463 | ALIRVLQNF | F108C_PGM | 0.5 | -0.021 | 0.0265 |
| 455 | 463 | ALIRVLQNF | F108C_PGM | 5 | 0.0806 | 0.0347 |
| 455 | 463 | ALIRVLQNF | F108C_PGM | 120 | 0.4305 | 0.0302 |
| 457 | 463 | IRVLQNF | F108C | 0 | 0 | 0 |
| 457 | 463 | IRVLQNF | F108C | 0.5 | 0.1337 | 0.0259 |
| 457 | 463 | IRVLQNF | F108C | 5 | 0.2237 | 0.0237 |
| 457 | 463 | IRVLQNF | F108C | 120 | 0.4696 | 0.0286 |
| 457 | 463 | IRVLQNF | F108C_PGM | 0 | 0 | 0 |
| 457 | 463 | IRVLQNF | F108C_PGM | 0.5 | 0.1408 | 0.0297 |
| 457 | 463 | IRVLQNF | F108C_PGM | 5 | 0.2517 | 0.0245 |
| 457 | 463 | IRVLQNF | F108C_PGM | 120 | 0.4879 | 0.031 |
| 464 | 479 | SFKPGKETQIPLKLSL | F108C | 0 | 0 | 0 |
| 464 | 479 | SFKPGKETQIPLKLSL | F108C | 0.5 | 3.724 | 0.1176 |
| 464 | 479 | SFKPGKETQIPLKLSL | F108C | 5 | 3.9836 | 0.0569 |
| 464 | 479 | SFKPGKETQIPLKLSL | F108C | 120 | 4.0777 | 0.036 |
| 464 | 479 | SFKPGKETQIPLKLSL | F108C_PGM | 0 | 0 | 0 |
| 464 | 479 | SFKPGKETQIPLKLSL | F108C_PGM | 0.5 | 3.8092 | 0.0515 |
| 464 | 479 | SFKPGKETQIPLKLSL | F108C_PGM | 5 | 3.8734 | 0.1364 |
| 464 | 479 | SFKPGKETQIPLKLSL | F108C_PGM | 120 | 4.0605 | 0.2426 |
| 464 | 491 | SFKPGKETQIPLKLSLGGLLQPEKP | F108C | 0 | 0 | 0 |
| 464 | 491 | SFKPGKETQIPLKLSLGGLLQPEKP | F108C | 0.5 | 5.7238 | 0.0619 |
| 464 | 491 | SFKPGKETQIPLKLSLGGLLQPEKP | F108C | 5 | 7.4941 | 0.0632 |
| 464 | 491 | SFKPGKETQIPLKLSLGGLLQPEKP | F108C | 120 | 8.036 | 0.1977 |
| 464 | 491 | SFKPGKETQIPLKLSLGGLLQPEKP | F108C_PGM | 0 | 0 | 0 |
| 464 | 491 | SFKPGKETQIPLKLSLGGLLQPEKP | F108C_PGM | 0.5 | 5.0623 | 0.0547 |
| 464 | 491 | SFKPGKETQIPLKLSLGGLLQPEKP | F108C_PGM | 5 | 6.2505 | 0.094 |
| 464 | 491 | SFKPGKETQIPLKLSLGGLLQPEKP | F108C_PGM | 120 | 7.5418 | 0.1183 |
| 492 | 507 | KVESRDGTVSGAHHHH | F108C | 0 | 0 | 0 |
| 492 | 507 | KVESRDGTVSGAHHHH | F108C | 0.5 | 1.0725 | 0.1106 |
| 492 | 507 | KVESRDGTVSGAHHHH | F108C | 5 | 1.0899 | 0.1073 |
| 492 | 507 | KVESRDGTVSGAHHHH | F108C | 120 | 1.1507 | 0.1002 |
| 492 | 507 | KVESRDGTVSGAHHHH | F108C_PGM | 0 | 0 | 0 |
| 492 | 507 | KVESRDGTVSGAHHHH | F108C_PGM | 0.5 | 0.8525 | 0.082 |

| | | | | | | |
|---|---|---|---|---|---|---|
| 492 | 507 | KVESRDGTVSGAHHHH | F108C_PGM | 5 | 0.7945 | 0.0815 |
| 492 | 507 | KVESRDGTVSGAHHHH | F108C_PGM | 120 | 0.8759 | 0.1261 |
| 494 | 507 | ESRDGTVSGAHHHH | F108C | 0 | 0 | 0 |
| 494 | 507 | ESRDGTVSGAHHHH | F108C | 0.5 | 0.6408 | 0.0767 |
| 494 | 507 | ESRDGTVSGAHHHH | F108C | 5 | 0.6744 | 0.0737 |
| 494 | 507 | ESRDGTVSGAHHHH | F108C | 120 | 0.8108 | 0.0572 |
| 494 | 507 | ESRDGTVSGAHHHH | F108C_PGM | 0 | 0 | 0 |
| 494 | 507 | ESRDGTVSGAHHHH | F108C_PGM | 0.5 | 0.532 | 0.1418 |
| 494 | 507 | ESRDGTVSGAHHHH | F108C_PGM | 5 | 0.5197 | 0.0844 |
| 494 | 507 | ESRDGTVSGAHHHH | F108C_PGM | 120 | 0.5951 | 0.1562 |

| Deuterium uptake data for G481C and G481C_PGM CYP3A4 states | | | | | | |
|---|---|---|---|---|---|---|
| **Start** | **End** | **Sequence** | **State** | **Exposure (min)** | **Uptake (Da)** | **Uptake SD (Da)** |
| 43 | 56 | PLPFLGNILSYHKG | G481C | 0 | 0.000000 | 0.000000 |
| 43 | 56 | PLPFLGNILSYHKG | G481C | 5 | 3.352043 | 0.061796 |
| 43 | 56 | PLPFLGNILSYHKG | G481C | 60 | 5.597700 | 0.090357 |
| 43 | 56 | PLPFLGNILSYHKG | G481C_PGM | 0 | 0.000000 | 0.000000 |
| 43 | 56 | PLPFLGNILSYHKG | G481C_PGM | 5 | 3.402668 | 0.113649 |
| 43 | 56 | PLPFLGNILSYHKG | G481C_PGM | 60 | 5.511617 | 0.147972 |
| 49 | 60 | NILSYHKGFTMF | G481C | 0 | 0.000000 | 0.000000 |
| 49 | 60 | NILSYHKGFTMF | G481C | 5 | 1.866622 | 0.036689 |
| 49 | 60 | NILSYHKGFTMF | G481C | 60 | 2.014554 | 0.041459 |
| 49 | 60 | NILSYHKGFTMF | G481C_PGM | 0 | 0.000000 | 0.000000 |
| 49 | 60 | NILSYHKGFTMF | G481C_PGM | 5 | 1.795125 | 0.041166 |
| 49 | 60 | NILSYHKGFTMF | G481C_PGM | 60 | 1.992433 | 0.077517 |
| 54 | 60 | HKGFTMF | G481C | 0 | 0.000000 | 0.000000 |
| 54 | 60 | HKGFTMF | G481C | 5 | 0.869212 | 0.026678 |
| 54 | 60 | HKGFTMF | G481C | 60 | 0.878372 | 0.030189 |
| 54 | 60 | HKGFTMF | G481C_PGM | 0 | 0.000000 | 0.000000 |
| 54 | 60 | HKGFTMF | G481C_PGM | 5 | 0.827047 | 0.042547 |
| 54 | 60 | HKGFTMF | G481C_PGM | 60 | 0.849038 | 0.034459 |
| 60 | 74 | FDMEAHKKYGKVWGF | G481C | 0 | 0.000000 | 0.000000 |
| 60 | 74 | FDMEAHKKYGKVWGF | G481C | 5 | 0.898474 | 0.028332 |
| 60 | 74 | FDMEAHKKYGKVWGF | G481C | 60 | 1.421835 | 0.038734 |
| 60 | 74 | FDMEAHKKYGKVWGF | G481C_PGM | 0 | 0.000000 | 0.000000 |
| 60 | 74 | FDMEAHKKYGKVWGF | G481C_PGM | 5 | 0.874014 | 0.028276 |
| 60 | 74 | FDMEAHKKYGKVWGF | G481C_PGM | 60 | 1.351630 | 0.022060 |
| 61 | 74 | DMEAHKKYGKVWGF | G481C | 0 | 0.000000 | 0.000000 |
| 61 | 74 | DMEAHKKYGKVWGF | G481C | 5 | 1.156438 | 0.056128 |
| 61 | 74 | DMEAHKKYGKVWGF | G481C | 60 | 1.908156 | 0.328765 |
| 61 | 74 | DMEAHKKYGKVWGF | G481C_PGM | 0 | 0.000000 | 0.000000 |
| 61 | 74 | DMEAHKKYGKVWGF | G481C_PGM | 5 | 1.104208 | 0.045396 |
| 61 | 74 | DMEAHKKYGKVWGF | G481C_PGM | 60 | 1.538426 | 0.041379 |

| | | | | | | |
|---|---|---|---|---|---|---|
| 75 | 82 | YDGQQPVL | G481C | 0 | 0.000000 | 0.000000 |
| 75 | 82 | YDGQQPVL | G481C | 5 | 0.827464 | 0.012197 |
| 75 | 82 | YDGQQPVL | G481C | 60 | 1.150843 | 0.026509 |
| 75 | 82 | YDGQQPVL | G481C_PGM | 0 | 0.000000 | 0.000000 |
| 75 | 82 | YDGQQPVL | G481C_PGM | 5 | 0.793058 | 0.009229 |
| 75 | 82 | YDGQQPVL | G481C_PGM | 60 | 1.056044 | 0.039683 |
| 95 | 102 | VKESYSVF | G481C | 0 | 0.000000 | 0.000000 |
| 95 | 102 | VKESYSVF | G481C | 5 | 1.655308 | 0.028534 |
| 95 | 102 | VKESYSVF | G481C | 60 | 1.729975 | 0.038585 |
| 95 | 102 | VKESYSVF | G481C_PGM | 0 | 0.000000 | 0.000000 |
| 95 | 102 | VKESYSVF | G481C_PGM | 5 | 1.639335 | 0.039450 |
| 95 | 102 | VKESYSVF | G481C_PGM | 60 | 1.773461 | 0.130178 |
| 103 | 113 | TNRRPFGPVGF | G481C | 0 | 0.000000 | 0.000000 |
| 103 | 113 | TNRRPFGPVGF | G481C | 5 | 2.405364 | 0.057615 |
| 103 | 113 | TNRRPFGPVGF | G481C | 60 | 2.547156 | 0.072338 |
| 103 | 113 | TNRRPFGPVGF | G481C_PGM | 0 | 0.000000 | 0.000000 |
| 103 | 113 | TNRRPFGPVGF | G481C_PGM | 5 | 2.296582 | 0.121058 |
| 103 | 113 | TNRRPFGPVGF | G481C_PGM | 60 | 2.330808 | 0.107310 |
| 114 | 119 | MKSAIS | G481C | 0 | 0.000000 | 0.000000 |
| 114 | 119 | MKSAIS | G481C | 5 | 0.746430 | 0.019837 |
| 114 | 119 | MKSAIS | G481C | 60 | 1.334964 | 0.020778 |
| 114 | 119 | MKSAIS | G481C_PGM | 0 | 0.000000 | 0.000000 |
| 114 | 119 | MKSAIS | G481C_PGM | 5 | 0.692041 | 0.032889 |
| 114 | 119 | MKSAIS | G481C_PGM | 60 | 1.227181 | 0.060807 |
| 114 | 125 | MKSAISIAEDEE | G481C | 0 | 0.000000 | 0.000000 |
| 114 | 125 | MKSAISIAEDEE | G481C | 5 | 2.144307 | 0.021218 |
| 114 | 125 | MKSAISIAEDEE | G481C | 60 | 3.303349 | 0.053744 |
| 114 | 125 | MKSAISIAEDEE | G481C_PGM | 0 | 0.000000 | 0.000000 |
| 114 | 125 | MKSAISIAEDEE | G481C_PGM | 5 | 2.011472 | 0.077904 |
| 114 | 125 | MKSAISIAEDEE | G481C_PGM | 60 | 3.212079 | 0.101897 |
| 114 | 133 | MKSAISIAEDEEWKRLRSLL | G481C | 0 | 0.000000 | 0.000000 |
| 114 | 133 | MKSAISIAEDEEWKRLRSLL | G481C | 5 | 3.492180 | 0.064307 |
| 114 | 133 | MKSAISIAEDEEWKRLRSLL | G481C | 60 | 5.085046 | 0.042279 |
| 114 | 133 | MKSAISIAEDEEWKRLRSLL | G481C_PGM | 0 | 0.000000 | 0.000000 |
| 114 | 133 | MKSAISIAEDEEWKRLRSLL | G481C_PGM | 5 | 3.346801 | 0.127563 |
| 114 | 133 | MKSAISIAEDEEWKRLRSLL | G481C_PGM | 60 | 4.614074 | 0.040324 |
| 120 | 125 | IAEDEE | G481C | 0 | 0.000000 | 0.000000 |
| 120 | 125 | IAEDEE | G481C | 5 | 0.962221 | 0.022219 |
| 120 | 125 | IAEDEE | G481C | 60 | 1.065026 | 0.029720 |
| 120 | 125 | IAEDEE | G481C_PGM | 0 | 0.000000 | 0.000000 |
| 120 | 125 | IAEDEE | G481C_PGM | 5 | 0.931070 | 0.017579 |
| 120 | 125 | IAEDEE | G481C_PGM | 60 | 1.049970 | 0.030879 |
| 120 | 133 | IAEDEEWKRLRSLL | G481C | 0 | 0.000000 | 0.000000 |
| 120 | 133 | IAEDEEWKRLRSLL | G481C | 5 | 1.864266 | 0.057877 |
| 120 | 133 | IAEDEEWKRLRSLL | G481C | 60 | 2.996626 | 0.033732 |

| | | | | | | |
|---|---|---|---|---|---|---|
| 120 | 133 | IAEDEEWKRLRSLL | G481C_PGM | 0 | 0.000000 | 0.000000 |
| 120 | 133 | IAEDEEWKRLRSLL | G481C_PGM | 5 | 1.820170 | 0.076509 |
| 120 | 133 | IAEDEEWKRLRSLL | G481C_PGM | 60 | 2.692140 | 0.092192 |
| 123 | 133 | DEEWKRLRSLL | G481C | 0 | 0.000000 | 0.000000 |
| 123 | 133 | DEEWKRLRSLL | G481C | 5 | 2.044780 | 0.020984 |
| 123 | 133 | DEEWKRLRSLL | G481C | 60 | 2.597469 | 0.036834 |
| 123 | 133 | DEEWKRLRSLL | G481C_PGM | 0 | 0.000000 | 0.000000 |
| 123 | 133 | DEEWKRLRSLL | G481C_PGM | 5 | 2.106246 | 0.014361 |
| 123 | 133 | DEEWKRLRSLL | G481C_PGM | 60 | 2.552554 | 0.047418 |
| 123 | 137 | DEEWKRLRSLLSPTF | G481C | 0 | 0.000000 | 0.000000 |
| 123 | 137 | DEEWKRLRSLLSPTF | G481C | 5 | 2.473775 | 0.024603 |
| 123 | 137 | DEEWKRLRSLLSPTF | G481C | 60 | 3.658667 | 0.055139 |
| 123 | 137 | DEEWKRLRSLLSPTF | G481C_PGM | 0 | 0.000000 | 0.000000 |
| 123 | 137 | DEEWKRLRSLLSPTF | G481C_PGM | 5 | 2.517361 | 0.140807 |
| 123 | 137 | DEEWKRLRSLLSPTF | G481C_PGM | 60 | 3.363925 | 0.065780 |
| 126 | 133 | WKRLRSLL | G481C | 0 | 0.000000 | 0.000000 |
| 126 | 133 | WKRLRSLL | G481C | 5 | 0.776196 | 0.044313 |
| 126 | 133 | WKRLRSLL | G481C | 60 | 1.136874 | 0.039796 |
| 126 | 133 | WKRLRSLL | G481C_PGM | 0 | 0.000000 | 0.000000 |
| 126 | 133 | WKRLRSLL | G481C_PGM | 5 | 0.780152 | 0.055506 |
| 126 | 133 | WKRLRSLL | G481C_PGM | 60 | 1.014550 | 0.032642 |
| 126 | 137 | WKRLRSLLSPTF | G481C | 0 | 0.000000 | 0.000000 |
| 126 | 137 | WKRLRSLLSPTF | G481C | 5 | 1.509230 | 0.081380 |
| 126 | 137 | WKRLRSLLSPTF | G481C | 60 | 2.185424 | 0.107671 |
| 126 | 137 | WKRLRSLLSPTF | G481C_PGM | 0 | 0.000000 | 0.000000 |
| 126 | 137 | WKRLRSLLSPTF | G481C_PGM | 5 | 1.575351 | 0.105099 |
| 126 | 137 | WKRLRSLLSPTF | G481C_PGM | 60 | 2.057464 | 0.075976 |
| 134 | 151 | SPTFTSGKLKEMVPIIAQ | G481C | 0 | 0.000000 | 0.000000 |
| 134 | 151 | SPTFTSGKLKEMVPIIAQ | G481C | 5 | 3.247643 | 0.058745 |
| 134 | 151 | SPTFTSGKLKEMVPIIAQ | G481C | 60 | 4.456580 | 0.073325 |
| 134 | 151 | SPTFTSGKLKEMVPIIAQ | G481C_PGM | 0 | 0.000000 | 0.000000 |
| 134 | 151 | SPTFTSGKLKEMVPIIAQ | G481C_PGM | 5 | 3.232679 | 0.205269 |
| 134 | 151 | SPTFTSGKLKEMVPIIAQ | G481C_PGM | 60 | 4.189956 | 0.058813 |
| 138 | 151 | TSGKLKEMVPIIAQ | G481C | 0 | 0.000000 | 0.000000 |
| 138 | 151 | TSGKLKEMVPIIAQ | G481C | 5 | 2.812514 | 0.053638 |
| 138 | 151 | TSGKLKEMVPIIAQ | G481C | 60 | 4.221444 | 0.023611 |
| 138 | 151 | TSGKLKEMVPIIAQ | G481C_PGM | 0 | 0.000000 | 0.000000 |
| 138 | 151 | TSGKLKEMVPIIAQ | G481C_PGM | 5 | 2.878992 | 0.112392 |
| 138 | 151 | TSGKLKEMVPIIAQ | G481C_PGM | 60 | 4.078555 | 0.108331 |
| 143 | 151 | KEMVPIIAQ | G481C | 0 | 0.000000 | 0.000000 |
| 143 | 151 | KEMVPIIAQ | G481C | 5 | 0.931599 | 0.044261 |
| 143 | 151 | KEMVPIIAQ | G481C | 60 | 1.728926 | 0.027554 |
| 143 | 151 | KEMVPIIAQ | G481C_PGM | 0 | 0.000000 | 0.000000 |
| 143 | 151 | KEMVPIIAQ | G481C_PGM | 5 | 0.936401 | 0.054529 |
| 143 | 151 | KEMVPIIAQ | G481C_PGM | 60 | 1.860659 | 0.097233 |

| | | | | | | |
|---|---|---|---|---|---|---|
| 157 | 172 | VRNLRREAETGKPVTL | G481C | 0 | 0.000000 | 0.000000 |
| 157 | 172 | VRNLRREAETGKPVTL | G481C | 5 | 3.699799 | 0.068304 |
| 157 | 172 | VRNLRREAETGKPVTL | G481C | 60 | 4.203402 | 0.121128 |
| 157 | 172 | VRNLRREAETGKPVTL | G481C_PGM | 0 | 0.000000 | 0.000000 |
| 157 | 172 | VRNLRREAETGKPVTL | G481C_PGM | 5 | 3.657941 | 0.173495 |
| 157 | 172 | VRNLRREAETGKPVTL | G481C_PGM | 60 | 4.000564 | 0.157458 |
| 157 | 176 | VRNLRREAETGKPVTLKDVF | G481C | 0 | 0.000000 | 0.000000 |
| 157 | 176 | VRNLRREAETGKPVTLKDVF | G481C | 5 | 5.134219 | 0.059381 |
| 157 | 176 | VRNLRREAETGKPVTLKDVF | G481C | 60 | 5.930498 | 0.086417 |
| 157 | 176 | VRNLRREAETGKPVTLKDVF | G481C_PGM | 0 | 0.000000 | 0.000000 |
| 157 | 176 | VRNLRREAETGKPVTLKDVF | G481C_PGM | 5 | 5.009143 | 0.236722 |
| 157 | 176 | VRNLRREAETGKPVTLKDVF | G481C_PGM | 60 | 5.612448 | 0.151984 |
| 173 | 181 | KDVFGAYSM | G481C | 0 | 0.000000 | 0.000000 |
| 173 | 181 | KDVFGAYSM | G481C | 5 | 1.393291 | 0.026259 |
| 173 | 181 | KDVFGAYSM | G481C | 60 | 2.163260 | 0.024190 |
| 173 | 181 | KDVFGAYSM | G481C_PGM | 0 | 0.000000 | 0.000000 |
| 173 | 181 | KDVFGAYSM | G481C_PGM | 5 | 1.300303 | 0.030606 |
| 173 | 181 | KDVFGAYSM | G481C_PGM | 60 | 2.000564 | 0.006558 |
| 173 | 182 | KDVFGAYSMD | G481C | 0 | 0.000000 | 0.000000 |
| 173 | 182 | KDVFGAYSMD | G481C | 5 | 1.254352 | 0.094140 |
| 173 | 182 | KDVFGAYSMD | G481C | 60 | 2.258102 | 0.057448 |
| 173 | 182 | KDVFGAYSMD | G481C_PGM | 0 | 0.000000 | 0.000000 |
| 173 | 182 | KDVFGAYSMD | G481C_PGM | 5 | 1.145231 | 0.062258 |
| 173 | 182 | KDVFGAYSMD | G481C_PGM | 60 | 2.054866 | 0.042800 |
| 177 | 182 | GAYSMD | G481C | 0 | 0.000000 | 0.000000 |
| 177 | 182 | GAYSMD | G481C | 5 | 0.410745 | 0.048150 |
| 177 | 182 | GAYSMD | G481C | 60 | 0.693261 | 0.015665 |
| 177 | 182 | GAYSMD | G481C_PGM | 0 | 0.000000 | 0.000000 |
| 177 | 182 | GAYSMD | G481C_PGM | 5 | 0.432011 | 0.017968 |
| 177 | 182 | GAYSMD | G481C_PGM | 60 | 0.549693 | 0.010133 |
| 182 | 189 | DVITSTSF | G481C | 0 | 0.000000 | 0.000000 |
| 182 | 189 | DVITSTSF | G481C | 5 | 0.777013 | 0.034789 |
| 182 | 189 | DVITSTSF | G481C | 60 | 1.429711 | 0.017166 |
| 182 | 189 | DVITSTSF | G481C_PGM | 0 | 0.000000 | 0.000000 |
| 182 | 189 | DVITSTSF | G481C_PGM | 5 | 0.709702 | 0.021009 |
| 182 | 189 | DVITSTSF | G481C_PGM | 60 | 1.409918 | 0.023872 |
| 190 | 213 | GVNIDSLNNPQDPFVENTKKLLRF | G481C | 0 | 0.000000 | 0.000000 |
| 190 | 213 | GVNIDSLNNPQDPFVENTKKLLRF | G481C | 5 | 5.946781 | 0.107811 |
| 190 | 213 | GVNIDSLNNPQDPFVENTKKLLRF | G481C | 60 | 7.228096 | 0.188520 |
| 190 | 213 | GVNIDSLNNPQDPFVENTKKLLRF | G481C_PGM | 0 | 0.000000 | 0.000000 |
| 190 | 213 | GVNIDSLNNPQDPFVENTKKLLRF | G481C_PGM | 5 | 6.156522 | 0.181247 |
| 190 | 213 | GVNIDSLNNPQDPFVENTKKLLRF | G481C_PGM | 60 | 6.787248 | 0.206940 |
| 193 | 213 | IDSLNNPQDPFVENTKKLLRF | G481C | 0 | 0.000000 | 0.000000 |
| 193 | 213 | IDSLNNPQDPFVENTKKLLRF | G481C | 5 | 4.082065 | 0.068863 |
| 193 | 213 | IDSLNNPQDPFVENTKKLLRF | G481C | 60 | 4.958690 | 0.144390 |

| | | | | | | |
|---|---|---|---|---|---|---|
| 193 | 213 | IDSLNNPQDPFVENTKKLLRF | G481C_PGM | 0 | 0.000000 | 0.000000 |
| 193 | 213 | IDSLNNPQDPFVENTKKLLRF | G481C_PGM | 5 | 4.061772 | 0.188832 |
| 193 | 213 | IDSLNNPQDPFVENTKKLLRF | G481C_PGM | 60 | 4.205672 | 0.225215 |
| 219 | 232 | FFLSITVFPFLIPI | G481C | 0 | 0.000000 | 0.000000 |
| 219 | 232 | FFLSITVFPFLIPI | G481C | 5 | 2.958291 | 0.055199 |
| 219 | 232 | FFLSITVFPFLIPI | G481C | 60 | 4.977782 | 0.065064 |
| 219 | 232 | FFLSITVFPFLIPI | G481C_PGM | 0 | 0.000000 | 0.000000 |
| 219 | 232 | FFLSITVFPFLIPI | G481C_PGM | 5 | 2.999804 | 0.083429 |
| 219 | 232 | FFLSITVFPFLIPI | G481C_PGM | 60 | 4.730165 | 0.088396 |
| 221 | 226 | LSITVF | G481C | 0 | 0.000000 | 0.000000 |
| 221 | 226 | LSITVF | G481C | 5 | 0.746032 | 0.016085 |
| 221 | 226 | LSITVF | G481C | 60 | 1.468019 | 0.014150 |
| 221 | 226 | LSITVF | G481C_PGM | 0 | 0.000000 | 0.000000 |
| 221 | 226 | LSITVF | G481C_PGM | 5 | 0.598509 | 0.081649 |
| 221 | 226 | LSITVF | G481C_PGM | 60 | 1.115300 | 0.096959 |
| 230 | 235 | IPILEV | G481C | 0 | 0.000000 | 0.000000 |
| 230 | 235 | IPILEV | G481C | 5 | 0.351304 | 0.036191 |
| 230 | 235 | IPILEV | G481C | 60 | 0.872942 | 0.027544 |
| 230 | 235 | IPILEV | G481C_PGM | 0 | 0.000000 | 0.000000 |
| 230 | 235 | IPILEV | G481C_PGM | 5 | 0.352845 | 0.069896 |
| 230 | 235 | IPILEV | G481C_PGM | 60 | 0.845366 | 0.032534 |
| 235 | 241 | VLNISVF | G481C | 0 | 0.000000 | 0.000000 |
| 235 | 241 | VLNISVF | G481C | 5 | 2.014566 | 0.039022 |
| 235 | 241 | VLNISVF | G481C | 60 | 2.788316 | 0.036180 |
| 235 | 241 | VLNISVF | G481C_PGM | 0 | 0.000000 | 0.000000 |
| 235 | 241 | VLNISVF | G481C_PGM | 5 | 1.976493 | 0.034514 |
| 235 | 241 | VLNISVF | G481C_PGM | 60 | 2.672524 | 0.093100 |
| 236 | 241 | LNISVF | G481C | 0 | 0.000000 | 0.000000 |
| 236 | 241 | LNISVF | G481C | 5 | 1.528458 | 0.021476 |
| 236 | 241 | LNISVF | G481C | 60 | 2.093382 | 0.025543 |
| 236 | 241 | LNISVF | G481C_PGM | 0 | 0.000000 | 0.000000 |
| 236 | 241 | LNISVF | G481C_PGM | 5 | 1.454574 | 0.109239 |
| 236 | 241 | LNISVF | G481C_PGM | 60 | 1.966211 | 0.049073 |
| 237 | 248 | NISVFPREVTNF | G481C | 0 | 0.000000 | 0.000000 |
| 237 | 248 | NISVFPREVTNF | G481C | 5 | 3.092889 | 0.041656 |
| 237 | 248 | NISVFPREVTNF | G481C | 60 | 3.480937 | 0.043431 |
| 237 | 248 | NISVFPREVTNF | G481C_PGM | 0 | 0.000000 | 0.000000 |
| 237 | 248 | NISVFPREVTNF | G481C_PGM | 5 | 3.102550 | 0.056765 |
| 237 | 248 | NISVFPREVTNF | G481C_PGM | 60 | 3.454024 | 0.124179 |
| 242 | 248 | PREVTNF | G481C | 0 | 0.000000 | 0.000000 |
| 242 | 248 | PREVTNF | G481C | 5 | 1.991315 | 0.018359 |
| 242 | 248 | PREVTNF | G481C | 60 | 2.112311 | 0.049287 |
| 242 | 248 | PREVTNF | G481C_PGM | 0 | 0.000000 | 0.000000 |
| 242 | 248 | PREVTNF | G481C_PGM | 5 | 2.090898 | 0.097161 |
| 242 | 248 | PREVTNF | G481C_PGM | 60 | 2.133479 | 0.094916 |

| | | | | | | |
|---|---|---|---|---|---|---|
| 249 | 271 | LRKSVKRMKESRLEDTQKHRVDF | G481C | 0 | 0.000000 | 0.000000 |
| 249 | 271 | LRKSVKRMKESRLEDTQKHRVDF | G481C | 5 | 2.677260 | 0.041407 |
| 249 | 271 | LRKSVKRMKESRLEDTQKHRVDF | G481C | 60 | 2.892094 | 0.163051 |
| 249 | 271 | LRKSVKRMKESRLEDTQKHRVDF | G481C_PGM | 0 | 0.000000 | 0.000000 |
| 249 | 271 | LRKSVKRMKESRLEDTQKHRVDF | G481C_PGM | 5 | 2.626035 | 0.141419 |
| 249 | 271 | LRKSVKRMKESRLEDTQKHRVDF | G481C_PGM | 60 | 2.944525 | 0.157888 |
| 262 | 271 | EDTQKHRVDF | G481C | 0 | 0.000000 | 0.000000 |
| 262 | 271 | EDTQKHRVDF | G481C | 5 | 0.972299 | 0.047959 |
| 262 | 271 | EDTQKHRVDF | G481C | 60 | 1.096926 | 0.071060 |
| 262 | 271 | EDTQKHRVDF | G481C_PGM | 0 | 0.000000 | 0.000000 |
| 262 | 271 | EDTQKHRVDF | G481C_PGM | 5 | 0.959149 | 0.063181 |
| 262 | 271 | EDTQKHRVDF | G481C_PGM | 60 | 1.045839 | 0.093779 |
| 275 | 292 | MIDSQNSKETESHKALSD | G481C | 0 | 0.000000 | 0.000000 |
| 275 | 292 | MIDSQNSKETESHKALSD | G481C | 5 | 2.756692 | 0.055458 |
| 275 | 292 | MIDSQNSKETESHKALSD | G481C | 60 | 2.802530 | 0.084889 |
| 275 | 292 | MIDSQNSKETESHKALSD | G481C_PGM | 0 | 0.000000 | 0.000000 |
| 275 | 292 | MIDSQNSKETESHKALSD | G481C_PGM | 5 | 2.688163 | 0.055025 |
| 275 | 292 | MIDSQNSKETESHKALSD | G481C_PGM | 60 | 2.654687 | 0.080038 |
| 275 | 293 | MIDSQNSKETESHKALSDL | G481C | 0 | 0.000000 | 0.000000 |
| 275 | 293 | MIDSQNSKETESHKALSDL | G481C | 5 | 3.132396 | 0.117243 |
| 275 | 293 | MIDSQNSKETESHKALSDL | G481C | 60 | 3.150007 | 0.140970 |
| 275 | 293 | MIDSQNSKETESHKALSDL | G481C_PGM | 0 | 0.000000 | 0.000000 |
| 275 | 293 | MIDSQNSKETESHKALSDL | G481C_PGM | 5 | 2.986280 | 0.213485 |
| 275 | 293 | MIDSQNSKETESHKALSDL | G481C_PGM | 60 | 2.918057 | 0.046344 |
| 277 | 292 | DSQNSKETESHKALSD | G481C | 0 | 0.000000 | 0.000000 |
| 277 | 292 | DSQNSKETESHKALSD | G481C | 5 | 2.775457 | 0.063637 |
| 277 | 292 | DSQNSKETESHKALSD | G481C | 60 | 2.863096 | 0.110480 |
| 277 | 292 | DSQNSKETESHKALSD | G481C_PGM | 0 | 0.000000 | 0.000000 |
| 277 | 292 | DSQNSKETESHKALSD | G481C_PGM | 5 | 2.417238 | 0.021637 |
| 277 | 292 | DSQNSKETESHKALSD | G481C_PGM | 60 | 2.591864 | 0.162579 |
| 290 | 295 | LSDLEL | G481C | 0 | 0.000000 | 0.000000 |
| 290 | 295 | LSDLEL | G481C | 5 | 0.197860 | 0.027381 |
| 290 | 295 | LSDLEL | G481C | 60 | 0.247044 | 0.023696 |
| 290 | 295 | LSDLEL | G481C_PGM | 0 | 0.000000 | 0.000000 |
| 290 | 295 | LSDLEL | G481C_PGM | 5 | 0.064167 | 0.013008 |
| 290 | 295 | LSDLEL | G481C_PGM | 60 | 0.081151 | 0.046145 |
| 296 | 302 | VAQSIIF | G481C | 0 | 0.000000 | 0.000000 |
| 296 | 302 | VAQSIIF | G481C | 5 | 0.281920 | 0.028068 |
| 296 | 302 | VAQSIIF | G481C | 60 | 0.629445 | 0.038616 |
| 296 | 302 | VAQSIIF | G481C_PGM | 0 | 0.000000 | 0.000000 |
| 296 | 302 | VAQSIIF | G481C_PGM | 5 | 0.253752 | 0.052736 |
| 296 | 302 | VAQSIIF | G481C_PGM | 60 | 0.605640 | 0.045982 |
| 301 | 306 | IFIFAG | G481C | 0 | 0.000000 | 0.000000 |
| 301 | 306 | IFIFAG | G481C | 5 | 0.176902 | 0.012874 |
| 301 | 306 | IFIFAG | G481C | 60 | 0.362552 | 0.015718 |

| | | | | | | |
|---|---|---|---|---|---|---|
| 301 | 306 | IFIFAG | G481C_PGM | 0 | 0.000000 | 0.000000 |
| 301 | 306 | IFIFAG | G481C_PGM | 5 | 0.204524 | 0.017618 |
| 301 | 306 | IFIFAG | G481C_PGM | 60 | 0.365433 | 0.026291 |
| 307 | 313 | YETTSSV | G481C | 0 | 0.000000 | 0.000000 |
| 307 | 313 | YETTSSV | G481C | 5 | 1.054014 | 0.035941 |
| 307 | 313 | YETTSSV | G481C | 60 | 1.463699 | 0.017159 |
| 307 | 313 | YETTSSV | G481C_PGM | 0 | 0.000000 | 0.000000 |
| 307 | 313 | YETTSSV | G481C_PGM | 5 | 0.881934 | 0.028421 |
| 307 | 313 | YETTSSV | G481C_PGM | 60 | 1.384303 | 0.036484 |
| 307 | 316 | YETTSSVLSF | G481C | 0 | 0.000000 | 0.000000 |
| 307 | 316 | YETTSSVLSF | G481C | 5 | 0.568823 | 0.150119 |
| 307 | 316 | YETTSSVLSF | G481C | 60 | 1.291254 | 0.057657 |
| 307 | 316 | YETTSSVLSF | G481C_PGM | 0 | 0.000000 | 0.000000 |
| 307 | 316 | YETTSSVLSF | G481C_PGM | 5 | 0.599505 | 0.017073 |
| 307 | 316 | YETTSSVLSF | G481C_PGM | 60 | 1.121021 | 0.086226 |
| 317 | 333 | IMYELATHPDVQQKLQE | G481C | 0 | 0.000000 | 0.000000 |
| 317 | 333 | IMYELATHPDVQQKLQE | G481C | 5 | 0.607850 | 0.057957 |
| 317 | 333 | IMYELATHPDVQQKLQE | G481C | 60 | 1.162455 | 0.063407 |
| 317 | 333 | IMYELATHPDVQQKLQE | G481C_PGM | 0 | 0.000000 | 0.000000 |
| 317 | 333 | IMYELATHPDVQQKLQE | G481C_PGM | 5 | 0.652114 | 0.125020 |
| 317 | 333 | IMYELATHPDVQQKLQE | G481C_PGM | 60 | 1.131603 | 0.066810 |
| 321 | 333 | LATHPDVQQKLQE | G481C | 0 | 0.000000 | 0.000000 |
| 321 | 333 | LATHPDVQQKLQE | G481C | 5 | 0.483859 | 0.041728 |
| 321 | 333 | LATHPDVQQKLQE | G481C | 60 | 0.946980 | 0.023665 |
| 321 | 333 | LATHPDVQQKLQE | G481C_PGM | 0 | 0.000000 | 0.000000 |
| 321 | 333 | LATHPDVQQKLQE | G481C_PGM | 5 | 0.367452 | 0.018247 |
| 321 | 333 | LATHPDVQQKLQE | G481C_PGM | 60 | 1.075645 | 0.033955 |
| 322 | 336 | ATHPDVQQKLQEEID | G481C | 0 | 0.000000 | 0.000000 |
| 322 | 336 | ATHPDVQQKLQEEID | G481C | 5 | 0.517456 | 0.091182 |
| 322 | 336 | ATHPDVQQKLQEEID | G481C | 60 | 1.016854 | 0.096749 |
| 322 | 336 | ATHPDVQQKLQEEID | G481C_PGM | 0 | 0.000000 | 0.000000 |
| 322 | 336 | ATHPDVQQKLQEEID | G481C_PGM | 5 | 0.509756 | 0.091039 |
| 322 | 336 | ATHPDVQQKLQEEID | G481C_PGM | 60 | 1.027032 | 0.072923 |
| 337 | 348 | AVLPNKAPPTYD | G481C | 0 | 0.000000 | 0.000000 |
| 337 | 348 | AVLPNKAPPTYD | G481C | 5 | 1.900044 | 0.041424 |
| 337 | 348 | AVLPNKAPPTYD | G481C | 60 | 2.318864 | 0.079137 |
| 337 | 348 | AVLPNKAPPTYD | G481C_PGM | 0 | 0.000000 | 0.000000 |
| 337 | 348 | AVLPNKAPPTYD | G481C_PGM | 5 | 1.832465 | 0.092449 |
| 337 | 348 | AVLPNKAPPTYD | G481C_PGM | 60 | 2.090776 | 0.084392 |
| 337 | 351 | AVLPNKAPPTYDTVL | G481C | 0 | 0.000000 | 0.000000 |
| 337 | 351 | AVLPNKAPPTYDTVL | G481C | 5 | 3.156257 | 0.102031 |
| 337 | 351 | AVLPNKAPPTYDTVL | G481C | 60 | 3.912423 | 0.115528 |
| 337 | 351 | AVLPNKAPPTYDTVL | G481C_PGM | 0 | 0.000000 | 0.000000 |
| 337 | 351 | AVLPNKAPPTYDTVL | G481C_PGM | 5 | 3.157263 | 0.118614 |
| 337 | 351 | AVLPNKAPPTYDTVL | G481C_PGM | 60 | 3.834660 | 0.081726 |

| | | | | | | |
|---|---|---|---|---|---|---|
| 357 | 363 | DMVVNET | G481C | 0 | 0.000000 | 0.000000 |
| 357 | 363 | DMVVNET | G481C | 5 | 0.076051 | 0.051616 |
| 357 | 363 | DMVVNET | G481C | 60 | 0.000874 | 0.052874 |
| 357 | 363 | DMVVNET | G481C_PGM | 0 | 0.000000 | 0.000000 |
| 357 | 363 | DMVVNET | G481C_PGM | 5 | 0.146807 | 0.061427 |
| 357 | 363 | DMVVNET | G481C_PGM | 60 | 0.057557 | 0.059964 |
| 359 | 366 | VVNETLRL | G481C | 0 | 0.000000 | 0.000000 |
| 359 | 366 | VVNETLRL | G481C | 5 | 0.173123 | 0.034960 |
| 359 | 366 | VVNETLRL | G481C | 60 | 0.335832 | 0.032176 |
| 359 | 366 | VVNETLRL | G481C_PGM | 0 | 0.000000 | 0.000000 |
| 359 | 366 | VVNETLRL | G481C_PGM | 5 | 0.202182 | 0.024043 |
| 359 | 366 | VVNETLRL | G481C_PGM | 60 | 0.345303 | 0.018945 |
| 363 | 370 | TLRLFPIA | G481C | 0 | 0.000000 | 0.000000 |
| 363 | 370 | TLRLFPIA | G481C | 5 | 0.262289 | 0.027473 |
| 363 | 370 | TLRLFPIA | G481C | 60 | 0.448703 | 0.037233 |
| 363 | 370 | TLRLFPIA | G481C_PGM | 0 | 0.000000 | 0.000000 |
| 363 | 370 | TLRLFPIA | G481C_PGM | 5 | 0.220021 | 0.062873 |
| 363 | 370 | TLRLFPIA | G481C_PGM | 60 | 0.409157 | 0.028029 |
| 364 | 370 | LRLFPIA | G481C | 0 | 0.000000 | 0.000000 |
| 364 | 370 | LRLFPIA | G481C | 5 | 0.257892 | 0.020571 |
| 364 | 370 | LRLFPIA | G481C | 60 | 0.482347 | 0.033740 |
| 364 | 370 | LRLFPIA | G481C_PGM | 0 | 0.000000 | 0.000000 |
| 364 | 370 | LRLFPIA | G481C_PGM | 5 | 0.258938 | 0.036135 |
| 364 | 370 | LRLFPIA | G481C_PGM | 60 | 0.437076 | 0.039047 |
| 364 | 371 | LRLFPIAM | G481C | 0 | 0.000000 | 0.000000 |
| 364 | 371 | LRLFPIAM | G481C | 5 | 0.582694 | 0.037014 |
| 364 | 371 | LRLFPIAM | G481C | 60 | 0.962122 | 0.042824 |
| 364 | 371 | LRLFPIAM | G481C_PGM | 0 | 0.000000 | 0.000000 |
| 364 | 371 | LRLFPIAM | G481C_PGM | 5 | 0.359407 | 0.091627 |
| 364 | 371 | LRLFPIAM | G481C_PGM | 60 | 0.670251 | 0.056560 |
| 367 | 373 | FPIAMRL | G481C | 0 | 0.000000 | 0.000000 |
| 367 | 373 | FPIAMRL | G481C | 5 | 0.761924 | 0.024322 |
| 367 | 373 | FPIAMRL | G481C | 60 | 1.162730 | 0.030321 |
| 367 | 373 | FPIAMRL | G481C_PGM | 0 | 0.000000 | 0.000000 |
| 367 | 373 | FPIAMRL | G481C_PGM | 5 | 0.631540 | 0.025866 |
| 367 | 373 | FPIAMRL | G481C_PGM | 60 | 0.969895 | 0.017585 |
| 374 | 387 | ERVCKKDVEINGMF | G481C | 0 | 0.000000 | 0.000000 |
| 374 | 387 | ERVCKKDVEINGMF | G481C | 5 | 1.800726 | 0.034634 |
| 374 | 387 | ERVCKKDVEINGMF | G481C | 60 | 2.158269 | 0.047048 |
| 374 | 387 | ERVCKKDVEINGMF | G481C_PGM | 0 | 0.000000 | 0.000000 |
| 374 | 387 | ERVCKKDVEINGMF | G481C_PGM | 5 | 1.894394 | 0.030925 |
| 374 | 387 | ERVCKKDVEINGMF | G481C_PGM | 60 | 2.143216 | 0.046700 |
| 384 | 402 | NGMFIPKGVVVMIPSYALH | G481C | 0 | 0.000000 | 0.000000 |
| 384 | 402 | NGMFIPKGVVVMIPSYALH | G481C | 5 | 1.888344 | 0.032437 |
| 384 | 402 | NGMFIPKGVVVMIPSYALH | G481C | 60 | 2.804224 | 0.251734 |

| | | | | | | |
|---|---|---|---|---|---|---|
| 384 | 402 | NGMFIPKGVVVMIPSYALH | G481C_PGM | 0 | 0.000000 | 0.000000 |
| 384 | 402 | NGMFIPKGVVVMIPSYALH | G481C_PGM | 5 | 2.209863 | 0.063472 |
| 384 | 402 | NGMFIPKGVVVMIPSYALH | G481C_PGM | 60 | 2.874810 | 0.176708 |
| 388 | 393 | IPKGVV | G481C | 0 | 0.000000 | 0.000000 |
| 388 | 393 | IPKGVV | G481C | 5 | 0.622337 | 0.035259 |
| 388 | 393 | IPKGVV | G481C | 60 | 0.855691 | 0.019760 |
| 388 | 393 | IPKGVV | G481C_PGM | 0 | 0.000000 | 0.000000 |
| 388 | 393 | IPKGVV | G481C_PGM | 5 | 0.611856 | 0.027549 |
| 388 | 393 | IPKGVV | G481C_PGM | 60 | 0.845411 | 0.024085 |
| 394 | 399 | VMIPSY | G481C | 0 | 0.000000 | 0.000000 |
| 394 | 399 | VMIPSY | G481C | 5 | 0.258959 | 0.017761 |
| 394 | 399 | VMIPSY | G481C | 60 | 0.367519 | 0.017117 |
| 394 | 399 | VMIPSY | G481C_PGM | 0 | 0.000000 | 0.000000 |
| 394 | 399 | VMIPSY | G481C_PGM | 5 | 0.303127 | 0.009121 |
| 394 | 399 | VMIPSY | G481C_PGM | 60 | 0.336410 | 0.013279 |
| 400 | 407 | ALHRDPKY | G481C | 0 | 0.000000 | 0.000000 |
| 400 | 407 | ALHRDPKY | G481C | 5 | 0.464891 | 0.064467 |
| 400 | 407 | ALHRDPKY | G481C | 60 | 0.596014 | 0.030779 |
| 400 | 407 | ALHRDPKY | G481C_PGM | 0 | 0.000000 | 0.000000 |
| 400 | 407 | ALHRDPKY | G481C_PGM | 5 | 0.452645 | 0.086993 |
| 400 | 407 | ALHRDPKY | G481C_PGM | 60 | 0.517866 | 0.048317 |
| 408 | 414 | WTEPEKF | G481C | 0 | 0.000000 | 0.000000 |
| 408 | 414 | WTEPEKF | G481C | 5 | 1.046637 | 0.025174 |
| 408 | 414 | WTEPEKF | G481C | 60 | 1.009645 | 0.053640 |
| 408 | 414 | WTEPEKF | G481C_PGM | 0 | 0.000000 | 0.000000 |
| 408 | 414 | WTEPEKF | G481C_PGM | 5 | 1.018468 | 0.063595 |
| 408 | 414 | WTEPEKF | G481C_PGM | 60 | 1.001390 | 0.028290 |
| 408 | 419 | WTEPEKFLPERF | G481C | 0 | 0.000000 | 0.000000 |
| 408 | 419 | WTEPEKFLPERF | G481C | 5 | 1.358406 | 0.044498 |
| 408 | 419 | WTEPEKFLPERF | G481C | 60 | 1.529521 | 0.077272 |
| 408 | 419 | WTEPEKFLPERF | G481C_PGM | 0 | 0.000000 | 0.000000 |
| 408 | 419 | WTEPEKFLPERF | G481C_PGM | 5 | 1.354899 | 0.071347 |
| 408 | 419 | WTEPEKFLPERF | G481C_PGM | 60 | 1.475740 | 0.046227 |
| 414 | 437 | FLPERFSKKNKDNIDPYIYTPFGS | G481C | 0 | 0.000000 | 0.000000 |
| 414 | 437 | FLPERFSKKNKDNIDPYIYTPFGS | G481C | 5 | 3.347115 | 0.094625 |
| 414 | 437 | FLPERFSKKNKDNIDPYIYTPFGS | G481C | 60 | 3.438690 | 0.083820 |
| 414 | 437 | FLPERFSKKNKDNIDPYIYTPFGS | G481C_PGM | 0 | 0.000000 | 0.000000 |
| 414 | 437 | FLPERFSKKNKDNIDPYIYTPFGS | G481C_PGM | 5 | 3.392913 | 0.094372 |
| 414 | 437 | FLPERFSKKNKDNIDPYIYTPFGS | G481C_PGM | 60 | 3.524199 | 0.090999 |
| 415 | 444 | LPERFSKKNKDNIDPYIYTPFGSGP | G481C | 0 | 0.000000 | 0.000000 |
| 415 | 444 | LPERFSKKNKDNIDPYIYTPFGSGP | G481C | 5 | 3.030663 | 0.057313 |
| 415 | 444 | LPERFSKKNKDNIDPYIYTPFGSGP | G481C | 60 | 3.715531 | 0.080059 |
| 415 | 444 | LPERFSKKNKDNIDPYIYTPFGSGP | G481C_PGM | 0 | 0.000000 | 0.000000 |
| 415 | 444 | LPERFSKKNKDNIDPYIYTPFGSGP | G481C_PGM | 5 | 2.944608 | 0.151170 |
| 415 | 444 | LPERFSKKNKDNIDPYIYTPFGSGP | G481C_PGM | 60 | 3.567663 | 0.099589 |

| | | | | | | |
|---|---|---|---|---|---|---|
| 420 | 444 | SKKNKDNIDPYIYTPFGSGPRNCIG | G481C | 0 | 0.000000 | 0.000000 |
| 420 | 444 | SKKNKDNIDPYIYTPFGSGPRNCIG | G481C | 5 | 3.089618 | 0.076233 |
| 420 | 444 | SKKNKDNIDPYIYTPFGSGPRNCIG | G481C | 60 | 3.824810 | 0.057169 |
| 420 | 444 | SKKNKDNIDPYIYTPFGSGPRNCIG | G481C_PGM | 0 | 0.000000 | 0.000000 |
| 420 | 444 | SKKNKDNIDPYIYTPFGSGPRNCIG | G481C_PGM | 5 | 3.124191 | 0.118848 |
| 420 | 444 | SKKNKDNIDPYIYTPFGSGPRNCIG | G481C_PGM | 60 | 3.650417 | 0.077842 |
| 445 | 452 | MRFALMNM | G481C | 0 | 0.000000 | 0.000000 |
| 445 | 452 | MRFALMNM | G481C | 5 | 0.345240 | 0.029521 |
| 445 | 452 | MRFALMNM | G481C | 60 | 0.525429 | 0.048565 |
| 445 | 452 | MRFALMNM | G481C_PGM | 0 | 0.000000 | 0.000000 |
| 445 | 452 | MRFALMNM | G481C_PGM | 5 | 0.371916 | 0.035871 |
| 445 | 452 | MRFALMNM | G481C_PGM | 60 | 0.603071 | 0.041758 |
| 446 | 452 | RFALMNM | G481C | 0 | 0.000000 | 0.000000 |
| 446 | 452 | RFALMNM | G481C | 5 | 0.181673 | 0.025021 |
| 446 | 452 | RFALMNM | G481C | 60 | 0.367962 | 0.038956 |
| 446 | 452 | RFALMNM | G481C_PGM | 0 | 0.000000 | 0.000000 |
| 446 | 452 | RFALMNM | G481C_PGM | 5 | 0.232272 | 0.031939 |
| 446 | 452 | RFALMNM | G481C_PGM | 60 | 0.476073 | 0.031937 |
| 448 | 456 | ALMNMKLAL | G481C | 0 | 0.000000 | 0.000000 |
| 448 | 456 | ALMNMKLAL | G481C | 5 | 0.289003 | 0.057003 |
| 448 | 456 | ALMNMKLAL | G481C | 60 | 0.379709 | 0.071797 |
| 448 | 456 | ALMNMKLAL | G481C_PGM | 0 | 0.000000 | 0.000000 |
| 448 | 456 | ALMNMKLAL | G481C_PGM | 5 | 0.346300 | 0.068286 |
| 448 | 456 | ALMNMKLAL | G481C_PGM | 60 | 0.460631 | 0.066392 |
| 451 | 456 | NMKLAL | G481C | 0 | 0.000000 | 0.000000 |
| 451 | 456 | NMKLAL | G481C | 5 | 0.095077 | 0.032324 |
| 451 | 456 | NMKLAL | G481C | 60 | 0.131362 | 0.029358 |
| 451 | 456 | NMKLAL | G481C_PGM | 0 | 0.000000 | 0.000000 |
| 451 | 456 | NMKLAL | G481C_PGM | 5 | 0.115868 | 0.026270 |
| 451 | 456 | NMKLAL | G481C_PGM | 60 | 0.165734 | 0.034306 |
| 455 | 463 | ALIRVLQNF | G481C | 0 | 0.000000 | 0.000000 |
| 455 | 463 | ALIRVLQNF | G481C | 5 | 0.302999 | 0.049756 |
| 455 | 463 | ALIRVLQNF | G481C | 60 | 0.512443 | 0.053231 |
| 455 | 463 | ALIRVLQNF | G481C_PGM | 0 | 0.000000 | 0.000000 |
| 455 | 463 | ALIRVLQNF | G481C_PGM | 5 | 0.297990 | 0.057534 |
| 455 | 463 | ALIRVLQNF | G481C_PGM | 60 | 0.510771 | 0.052300 |
| 457 | 462 | IRVLQN | G481C | 0 | 0.000000 | 0.000000 |
| 457 | 462 | IRVLQN | G481C | 5 | 1.245799 | 0.034460 |
| 457 | 462 | IRVLQN | G481C | 60 | 1.878771 | 0.054220 |
| 457 | 462 | IRVLQN | G481C_PGM | 0 | 0.000000 | 0.000000 |
| 457 | 462 | IRVLQN | G481C_PGM | 5 | 1.300916 | 0.026716 |
| 457 | 462 | IRVLQN | G481C_PGM | 60 | 1.927643 | 0.059950 |
| 457 | 463 | IRVLQNF | G481C | 0 | 0.000000 | 0.000000 |
| 457 | 463 | IRVLQNF | G481C | 5 | 0.298458 | 0.061088 |
| 457 | 463 | IRVLQNF | G481C | 60 | 0.509453 | 0.058128 |

| | | | | | | |
|---|---|---|---|---|---|---|
| 457 | 463 | IRVLQNF | G481C_PGM | 0 | 0.000000 | 0.000000 |
| 457 | 463 | IRVLQNF | G481C_PGM | 5 | 0.358897 | 0.068315 |
| 457 | 463 | IRVLQNF | G481C_PGM | 60 | 0.561448 | 0.057042 |
| 463 | 479 | FSFKPGKETQIPLKLSL | G481C | 0 | 0.000000 | 0.000000 |
| 463 | 479 | FSFKPGKETQIPLKLSL | G481C | 5 | 3.912363 | 0.031233 |
| 463 | 479 | FSFKPGKETQIPLKLSL | G481C | 60 | 4.005954 | 0.077309 |
| 463 | 479 | FSFKPGKETQIPLKLSL | G481C_PGM | 0 | 0.000000 | 0.000000 |
| 463 | 479 | FSFKPGKETQIPLKLSL | G481C_PGM | 5 | 3.764260 | 0.136339 |
| 463 | 479 | FSFKPGKETQIPLKLSL | G481C_PGM | 60 | 3.812103 | 0.146898 |
| 464 | 479 | SFKPGKETQIPLKLSL | G481C | 0 | 0.000000 | 0.000000 |
| 464 | 479 | SFKPGKETQIPLKLSL | G481C | 5 | 3.866064 | 0.081144 |
| 464 | 479 | SFKPGKETQIPLKLSL | G481C | 60 | 3.878819 | 0.090243 |
| 464 | 479 | SFKPGKETQIPLKLSL | G481C_PGM | 0 | 0.000000 | 0.000000 |
| 464 | 479 | SFKPGKETQIPLKLSL | G481C_PGM | 5 | 3.707491 | 0.124824 |
| 464 | 479 | SFKPGKETQIPLKLSL | G481C_PGM | 60 | 3.731103 | 0.110731 |
| 471 | 484 | TQIPLKLSLGCLLQ | G481C | 0 | 0.000000 | 0.000000 |
| 471 | 484 | TQIPLKLSLGCLLQ | G481C | 5 | 0.391402 | 0.061771 |
| 471 | 484 | TQIPLKLSLGCLLQ | G481C | 60 | 0.641177 | 0.057188 |
| 474 | 479 | PLKLSL | G481C_PGM | 0 | 0.000000 | 0.000000 |
| 474 | 479 | PLKLSL | G481C_PGM | 5 | 1.475923 | 0.032697 |
| 474 | 479 | PLKLSL | G481C_PGM | 60 | 1.479601 | 0.029944 |
| 474 | 479 | PLKLSL | G481C | 0 | 0.000000 | 0.000000 |
| 474 | 479 | PLKLSL | G481C | 5 | 1.413771 | 0.056089 |
| 474 | 479 | PLKLSL | G481C | 60 | 1.436946 | 0.066292 |
| 483 | 491 | LQPEKPVVL | G481C_PGM | 0 | 0.000000 | 0.000000 |
| 483 | 491 | LQPEKPVVL | G481C_PGM | 5 | 2.321591 | 0.057934 |
| 483 | 491 | LQPEKPVVL | G481C_PGM | 60 | 2.521139 | 0.124705 |
| 483 | 491 | LQPEKPVVL | G481C | 0 | 0.000000 | 0.000000 |
| 483 | 491 | LQPEKPVVL | G481C | 5 | 2.200329 | 0.155415 |
| 483 | 491 | LQPEKPVVL | G481C | 60 | 2.380622 | 0.058226 |
| 485 | 491 | PEKPVVL | G481C_PGM | 0 | 0.000000 | 0.000000 |
| 485 | 491 | PEKPVVL | G481C_PGM | 5 | 1.852877 | 0.014910 |
| 485 | 491 | PEKPVVL | G481C_PGM | 60 | 2.024336 | 0.053942 |
| 485 | 491 | PEKPVVL | G481C | 0 | 0.000000 | 0.000000 |
| 485 | 491 | PEKPVVL | G481C | 5 | 1.729848 | 0.086308 |
| 485 | 491 | PEKPVVL | G481C | 60 | 1.854052 | 0.059018 |
| 491 | 507 | LKVESRDGTVSGAHHHH | G481C_PGM | 0 | 0.000000 | 0.000000 |
| 491 | 507 | LKVESRDGTVSGAHHHH | G481C_PGM | 5 | 0.907703 | 0.082660 |
| 491 | 507 | LKVESRDGTVSGAHHHH | G481C_PGM | 60 | 1.082143 | 0.073700 |
| 491 | 507 | LKVESRDGTVSGAHHHH | G481C | 0 | 0.000000 | 0.000000 |
| 491 | 507 | LKVESRDGTVSGAHHHH | G481C | 5 | 0.942254 | 0.096152 |
| 491 | 507 | LKVESRDGTVSGAHHHH | G481C | 60 | 0.974854 | 0.101923 |
| 501 | 507 | SGAHHHH | G481C_PGM | 0 | 0.000000 | 0.000000 |
| 501 | 507 | SGAHHHH | G481C_PGM | 5 | 0.340926 | 0.051276 |
| 501 | 507 | SGAHHHH | G481C_PGM | 60 | 0.349283 | 0.046634 |

| | | | | | | |
|---|---|---|---|---|---|---|
| 501 | 507 | SGAHHHH | G481C | 0 | 0.000000 | 0.000000 |
| 501 | 507 | SGAHHHH | G481C | 5 | 0.362988 | 0.038544 |
| 501 | 507 | SGAHHHH | G481C | 60 | 0.333667 | 0.050942 |

| Deuterium uptake data for L482C and L482C_PGM CYP3A4 states | | | | | | |
|---|---|---|---|---|---|---|
| **Start** | **End** | **Sequence** | **State** | **Exposure (min)** | **Uptake (Da)** | **Uptake SD (Da)** |
| 33 | 51 | FKKLGIPGPTPLPFLGNIL | L482C | 0 | 0.000000 | 0.000000 |
| 33 | 51 | FKKLGIPGPTPLPFLGNIL | L482C | 5 | 0.520301 | 0.110302 |
| 33 | 51 | FKKLGIPGPTPLPFLGNIL | L482C | 60 | 0.417758 | 0.021215 |
| 33 | 51 | FKKLGIPGPTPLPFLGNIL | L482C_PGM | 0 | 0.000000 | 0.000000 |
| 33 | 51 | FKKLGIPGPTPLPFLGNIL | L482C_PGM | 5 | 0.506898 | 0.066323 |
| 33 | 51 | FKKLGIPGPTPLPFLGNIL | L482C_PGM | 60 | 0.498736 | 0.084517 |
| 34 | 51 | KKLGIPGPTPLPFLGNIL | L482C | 0 | 0.000000 | 0.000000 |
| 34 | 51 | KKLGIPGPTPLPFLGNIL | L482C | 5 | 0.533619 | 0.127626 |
| 34 | 51 | KKLGIPGPTPLPFLGNIL | L482C | 60 | 0.533205 | 0.133360 |
| 34 | 51 | KKLGIPGPTPLPFLGNIL | L482C_PGM | 0 | 0.000000 | 0.000000 |
| 34 | 51 | KKLGIPGPTPLPFLGNIL | L482C_PGM | 5 | 0.577328 | 0.019207 |
| 34 | 51 | KKLGIPGPTPLPFLGNIL | L482C_PGM | 60 | 0.512128 | 0.062985 |
| 49 | 60 | NILSYHKGFTMF | L482C | 0 | 0.000000 | 0.000000 |
| 49 | 60 | NILSYHKGFTMF | L482C | 5 | 2.473474 | 0.027306 |
| 49 | 60 | NILSYHKGFTMF | L482C | 60 | 2.627093 | 0.130329 |
| 49 | 60 | NILSYHKGFTMF | L482C_PGM | 0 | 0.000000 | 0.000000 |
| 49 | 60 | NILSYHKGFTMF | L482C_PGM | 5 | 2.550152 | 0.056288 |
| 49 | 60 | NILSYHKGFTMF | L482C_PGM | 60 | 2.658486 | 0.028995 |
| 54 | 59 | HKGFTM | L482C | 0 | 0.000000 | 0.000000 |
| 54 | 59 | HKGFTM | L482C | 5 | 1.158693 | 0.084296 |
| 54 | 59 | HKGFTM | L482C | 60 | 0.996625 | 0.132829 |
| 54 | 59 | HKGFTM | L482C_PGM | 0 | 0.000000 | 0.000000 |
| 54 | 59 | HKGFTM | L482C_PGM | 5 | 0.911631 | 0.079762 |
| 54 | 59 | HKGFTM | L482C_PGM | 60 | 0.927189 | 0.098317 |
| 60 | 73 | FDMEAHKKYGKVWG | L482C | 0 | 0.000000 | 0.000000 |
| 60 | 73 | FDMEAHKKYGKVWG | L482C | 5 | 1.370237 | 0.039490 |
| 60 | 73 | FDMEAHKKYGKVWG | L482C | 60 | 2.061272 | 0.081109 |
| 60 | 73 | FDMEAHKKYGKVWG | L482C_PGM | 0 | 0.000000 | 0.000000 |
| 60 | 73 | FDMEAHKKYGKVWG | L482C_PGM | 5 | 1.442041 | 0.039581 |
| 60 | 73 | FDMEAHKKYGKVWG | L482C_PGM | 60 | 2.278853 | 0.056973 |
| 60 | 74 | FDMEAHKKYGKVWGF | L482C | 0 | 0.000000 | 0.000000 |
| 60 | 74 | FDMEAHKKYGKVWGF | L482C | 5 | 1.024366 | 0.059906 |
| 60 | 74 | FDMEAHKKYGKVWGF | L482C | 60 | 1.738821 | 0.071506 |
| 60 | 74 | FDMEAHKKYGKVWGF | L482C_PGM | 0 | 0.000000 | 0.000000 |
| 60 | 74 | FDMEAHKKYGKVWGF | L482C_PGM | 5 | 1.085276 | 0.052715 |
| 60 | 74 | FDMEAHKKYGKVWGF | L482C_PGM | 60 | 1.909681 | 0.078251 |
| 61 | 74 | DMEAHKKYGKVWGF | L482C | 0 | 0.000000 | 0.000000 |
| 61 | 74 | DMEAHKKYGKVWGF | L482C | 5 | 1.093843 | 0.032157 |

| | | | | | | |
|---|---|---|---|---|---|---|
| 61 | 74 | DMEAHKKYGKVWGF | L482C | 60 | 1.821825 | 0.079455 |
| 61 | 74 | DMEAHKKYGKVWGF | L482C_PGM | 0 | 0.000000 | 0.000000 |
| 61 | 74 | DMEAHKKYGKVWGF | L482C_PGM | 5 | 1.135482 | 0.032541 |
| 61 | 74 | DMEAHKKYGKVWGF | L482C_PGM | 60 | 2.070337 | 0.063780 |
| 61 | 82 | DMEAHKKYGKVWGFYDGQQPVL | L482C | 0 | 0.000000 | 0.000000 |
| 61 | 82 | DMEAHKKYGKVWGFYDGQQPVL | L482C | 5 | 1.728500 | 0.046150 |
| 61 | 82 | DMEAHKKYGKVWGFYDGQQPVL | L482C | 60 | 2.985940 | 0.120556 |
| 61 | 82 | DMEAHKKYGKVWGFYDGQQPVL | L482C_PGM | 0 | 0.000000 | 0.000000 |
| 61 | 82 | DMEAHKKYGKVWGFYDGQQPVL | L482C_PGM | 5 | 1.841610 | 0.056421 |
| 61 | 82 | DMEAHKKYGKVWGFYDGQQPVL | L482C_PGM | 60 | 3.189279 | 0.060825 |
| 74 | 82 | FYDGQQPVL | L482C | 0 | 0.000000 | 0.000000 |
| 74 | 82 | FYDGQQPVL | L482C | 5 | 1.068510 | 0.034675 |
| 74 | 82 | FYDGQQPVL | L482C | 60 | 1.515919 | 0.053654 |
| 74 | 82 | FYDGQQPVL | L482C_PGM | 0 | 0.000000 | 0.000000 |
| 74 | 82 | FYDGQQPVL | L482C_PGM | 5 | 1.152522 | 0.028358 |
| 74 | 82 | FYDGQQPVL | L482C_PGM | 60 | 1.630172 | 0.077335 |
| 75 | 82 | YDGQQPVL | L482C | 0 | 0.000000 | 0.000000 |
| 75 | 82 | YDGQQPVL | L482C | 5 | 0.835072 | 0.025076 |
| 75 | 82 | YDGQQPVL | L482C | 60 | 1.154566 | 0.063999 |
| 75 | 82 | YDGQQPVL | L482C_PGM | 0 | 0.000000 | 0.000000 |
| 75 | 82 | YDGQQPVL | L482C_PGM | 5 | 0.860170 | 0.014736 |
| 75 | 82 | YDGQQPVL | L482C_PGM | 60 | 1.249834 | 0.031993 |
| 83 | 89 | AITDPDM | L482C | 0 | 0.000000 | 0.000000 |
| 83 | 89 | AITDPDM | L482C | 5 | 0.590618 | 0.033887 |
| 83 | 89 | AITDPDM | L482C | 60 | 0.765230 | 0.031716 |
| 83 | 89 | AITDPDM | L482C_PGM | 0 | 0.000000 | 0.000000 |
| 83 | 89 | AITDPDM | L482C_PGM | 5 | 0.567077 | 0.022208 |
| 83 | 89 | AITDPDM | L482C_PGM | 60 | 0.817279 | 0.031720 |
| 83 | 94 | AITDPDMIKTVL | L482C | 0 | 0.000000 | 0.000000 |
| 83 | 94 | AITDPDMIKTVL | L482C | 5 | 1.284180 | 0.068292 |
| 83 | 94 | AITDPDMIKTVL | L482C | 60 | 1.900810 | 0.055604 |
| 83 | 94 | AITDPDMIKTVL | L482C_PGM | 0 | 0.000000 | 0.000000 |
| 83 | 94 | AITDPDMIKTVL | L482C_PGM | 5 | 1.216973 | 0.035621 |
| 83 | 94 | AITDPDMIKTVL | L482C_PGM | 60 | 2.009260 | 0.020546 |
| 93 | 102 | VLVKESYSVF | L482C | 0 | 0.000000 | 0.000000 |
| 93 | 102 | VLVKESYSVF | L482C | 5 | 2.735453 | 0.046860 |
| 93 | 102 | VLVKESYSVF | L482C | 60 | 2.844409 | 0.079972 |
| 93 | 102 | VLVKESYSVF | L482C_PGM | 0 | 0.000000 | 0.000000 |
| 93 | 102 | VLVKESYSVF | L482C_PGM | 5 | 2.705599 | 0.056700 |
| 93 | 102 | VLVKESYSVF | L482C_PGM | 60 | 2.867561 | 0.057980 |
| 94 | 106 | LVKESYSVFTNRR | L482C | 0 | 0.000000 | 0.000000 |
| 94 | 106 | LVKESYSVFTNRR | L482C | 5 | 2.040041 | 0.067959 |
| 94 | 106 | LVKESYSVFTNRR | L482C | 60 | 2.469952 | 0.093044 |
| 94 | 106 | LVKESYSVFTNRR | L482C_PGM | 0 | 0.000000 | 0.000000 |
| 94 | 106 | LVKESYSVFTNRR | L482C_PGM | 5 | 1.992576 | 0.015871 |

| | | | | | | |
|---|---|---|---|---|---|---|
| 94 | 106 | LVKESYSVFTNRR | L482C_PGM | 60 | 2.332258 | 0.029790 |
| 95 | 102 | VKESYSVF | L482C | 0 | 0.000000 | 0.000000 |
| 95 | 102 | VKESYSVF | L482C | 5 | 2.105360 | 0.017027 |
| 95 | 102 | VKESYSVF | L482C | 60 | 2.404931 | 0.088901 |
| 95 | 102 | VKESYSVF | L482C_PGM | 0 | 0.000000 | 0.000000 |
| 95 | 102 | VKESYSVF | L482C_PGM | 5 | 2.105452 | 0.045530 |
| 95 | 102 | VKESYSVF | L482C_PGM | 60 | 2.539424 | 0.020163 |
| 114 | 122 | MKSAISIAE | L482C | 0 | 0.000000 | 0.000000 |
| 114 | 122 | MKSAISIAE | L482C | 5 | 1.252392 | 0.015089 |
| 114 | 122 | MKSAISIAE | L482C | 60 | 2.245711 | 0.037264 |
| 114 | 122 | MKSAISIAE | L482C_PGM | 0 | 0.000000 | 0.000000 |
| 114 | 122 | MKSAISIAE | L482C_PGM | 5 | 1.447032 | 0.049774 |
| 114 | 122 | MKSAISIAE | L482C_PGM | 60 | 2.806546 | 0.067160 |
| 120 | 125 | IAEDEE | L482C | 0 | 0.000000 | 0.000000 |
| 120 | 125 | IAEDEE | L482C | 5 | 0.825090 | 0.027650 |
| 120 | 125 | IAEDEE | L482C | 60 | 0.997410 | 0.037616 |
| 120 | 125 | IAEDEE | L482C_PGM | 0 | 0.000000 | 0.000000 |
| 120 | 125 | IAEDEE | L482C_PGM | 5 | 0.851441 | 0.034716 |
| 120 | 125 | IAEDEE | L482C_PGM | 60 | 1.078510 | 0.054894 |
| 123 | 133 | DEEWKRLRSLL | L482C | 0 | 0.000000 | 0.000000 |
| 123 | 133 | DEEWKRLRSLL | L482C | 5 | 1.307845 | 0.043646 |
| 123 | 133 | DEEWKRLRSLL | L482C | 60 | 1.945387 | 0.079302 |
| 123 | 133 | DEEWKRLRSLL | L482C_PGM | 0 | 0.000000 | 0.000000 |
| 123 | 133 | DEEWKRLRSLL | L482C_PGM | 5 | 1.540585 | 0.046858 |
| 123 | 133 | DEEWKRLRSLL | L482C_PGM | 60 | 2.438309 | 0.050197 |
| 123 | 137 | DEEWKRLRSLLSPTF | L482C | 0 | 0.000000 | 0.000000 |
| 123 | 137 | DEEWKRLRSLLSPTF | L482C | 5 | 1.880972 | 0.053164 |
| 123 | 137 | DEEWKRLRSLLSPTF | L482C | 60 | 3.093118 | 0.189232 |
| 123 | 137 | DEEWKRLRSLLSPTF | L482C_PGM | 0 | 0.000000 | 0.000000 |
| 123 | 137 | DEEWKRLRSLLSPTF | L482C_PGM | 5 | 2.298163 | 0.099262 |
| 123 | 137 | DEEWKRLRSLLSPTF | L482C_PGM | 60 | 3.840666 | 0.187779 |
| 126 | 137 | WKRLRSLLSPTF | L482C | 0 | 0.000000 | 0.000000 |
| 126 | 137 | WKRLRSLLSPTF | L482C | 5 | 1.247083 | 0.028516 |
| 126 | 137 | WKRLRSLLSPTF | L482C | 60 | 2.227737 | 0.068686 |
| 126 | 137 | WKRLRSLLSPTF | L482C_PGM | 0 | 0.000000 | 0.000000 |
| 126 | 137 | WKRLRSLLSPTF | L482C_PGM | 5 | 1.674510 | 0.040284 |
| 126 | 137 | WKRLRSLLSPTF | L482C_PGM | 60 | 2.566286 | 0.077888 |
| 132 | 151 | LLSPTFTSGKLKEMVPIIAQ | L482C | 0 | 0.000000 | 0.000000 |
| 132 | 151 | LLSPTFTSGKLKEMVPIIAQ | L482C | 5 | 2.137482 | 0.075011 |
| 132 | 151 | LLSPTFTSGKLKEMVPIIAQ | L482C | 60 | 4.398015 | 0.099686 |
| 132 | 151 | LLSPTFTSGKLKEMVPIIAQ | L482C_PGM | 0 | 0.000000 | 0.000000 |
| 132 | 151 | LLSPTFTSGKLKEMVPIIAQ | L482C_PGM | 5 | 2.113406 | 0.091039 |
| 132 | 151 | LLSPTFTSGKLKEMVPIIAQ | L482C_PGM | 60 | 4.433343 | 0.082720 |
| 134 | 151 | SPTFTSGKLKEMVPIIAQ | L482C | 0 | 0.000000 | 0.000000 |
| 134 | 151 | SPTFTSGKLKEMVPIIAQ | L482C | 5 | 3.004194 | 0.071873 |

| | | | | | | |
|---|---|---|---|---|---|---|
| 134 | 151 | SPTFTSGKLKEMVPIIAQ | L482C | 60 | 4.358943 | 0.142840 |
| 134 | 151 | SPTFTSGKLKEMVPIIAQ | L482C_PGM | 0 | 0.000000 | 0.000000 |
| 134 | 151 | SPTFTSGKLKEMVPIIAQ | L482C_PGM | 5 | 3.224872 | 0.027478 |
| 134 | 151 | SPTFTSGKLKEMVPIIAQ | L482C_PGM | 60 | 4.682248 | 0.124602 |
| 138 | 151 | TSGKLKEMVPIIAQ | L482C | 0 | 0.000000 | 0.000000 |
| 138 | 151 | TSGKLKEMVPIIAQ | L482C | 5 | 2.638142 | 0.091557 |
| 138 | 151 | TSGKLKEMVPIIAQ | L482C | 60 | 3.921367 | 0.089696 |
| 138 | 151 | TSGKLKEMVPIIAQ | L482C_PGM | 0 | 0.000000 | 0.000000 |
| 138 | 151 | TSGKLKEMVPIIAQ | L482C_PGM | 5 | 2.841990 | 0.029022 |
| 138 | 151 | TSGKLKEMVPIIAQ | L482C_PGM | 60 | 4.198457 | 0.098486 |
| 143 | 151 | KEMVPIIAQ | L482C | 0 | 0.000000 | 0.000000 |
| 143 | 151 | KEMVPIIAQ | L482C | 5 | 1.004182 | 0.040895 |
| 143 | 151 | KEMVPIIAQ | L482C | 60 | 2.095392 | 0.010810 |
| 143 | 151 | KEMVPIIAQ | L482C_PGM | 0 | 0.000000 | 0.000000 |
| 143 | 151 | KEMVPIIAQ | L482C_PGM | 5 | 1.167659 | 0.022884 |
| 143 | 151 | KEMVPIIAQ | L482C_PGM | 60 | 2.358968 | 0.054950 |
| 157 | 176 | VRNLRREAETGKPVTLKDVF | L482C | 0 | 0.000000 | 0.000000 |
| 157 | 176 | VRNLRREAETGKPVTLKDVF | L482C | 5 | 6.004327 | 0.091256 |
| 157 | 176 | VRNLRREAETGKPVTLKDVF | L482C | 60 | 6.814507 | 0.222444 |
| 157 | 176 | VRNLRREAETGKPVTLKDVF | L482C_PGM | 0 | 0.000000 | 0.000000 |
| 157 | 176 | VRNLRREAETGKPVTLKDVF | L482C_PGM | 5 | 5.895484 | 0.146956 |
| 157 | 176 | VRNLRREAETGKPVTLKDVF | L482C_PGM | 60 | 6.772672 | 0.166898 |
| 157 | 178 | VRNLRREAETGKPVTLKDVFGA | L482C | 0 | 0.000000 | 0.000000 |
| 157 | 178 | VRNLRREAETGKPVTLKDVFGA | L482C | 5 | 6.616306 | 0.071742 |
| 157 | 178 | VRNLRREAETGKPVTLKDVFGA | L482C | 60 | 7.590621 | 0.239415 |
| 157 | 178 | VRNLRREAETGKPVTLKDVFGA | L482C_PGM | 0 | 0.000000 | 0.000000 |
| 157 | 178 | VRNLRREAETGKPVTLKDVFGA | L482C_PGM | 5 | 6.546958 | 0.120586 |
| 157 | 178 | VRNLRREAETGKPVTLKDVFGA | L482C_PGM | 60 | 7.733127 | 0.213449 |
| 173 | 181 | KDVFGAYSM | L482C | 0 | 0.000000 | 0.000000 |
| 173 | 181 | KDVFGAYSM | L482C | 5 | 1.480253 | 0.047844 |
| 173 | 181 | KDVFGAYSM | L482C | 60 | 2.321264 | 0.063437 |
| 173 | 181 | KDVFGAYSM | L482C_PGM | 0 | 0.000000 | 0.000000 |
| 173 | 181 | KDVFGAYSM | L482C_PGM | 5 | 1.516353 | 0.047303 |
| 173 | 181 | KDVFGAYSM | L482C_PGM | 60 | 2.493543 | 0.046261 |
| 177 | 182 | GAYSMD | L482C | 0 | 0.000000 | 0.000000 |
| 177 | 182 | GAYSMD | L482C | 5 | 0.235257 | 0.083777 |
| 177 | 182 | GAYSMD | L482C | 60 | 0.525626 | 0.026147 |
| 177 | 182 | GAYSMD | L482C_PGM | 0 | 0.000000 | 0.000000 |
| 177 | 182 | GAYSMD | L482C_PGM | 5 | 0.344360 | 0.053303 |
| 177 | 182 | GAYSMD | L482C_PGM | 60 | 0.742259 | 0.032755 |
| 182 | 189 | DVITSTSF | L482C | 0 | 0.000000 | 0.000000 |
| 182 | 189 | DVITSTSF | L482C | 5 | 0.773183 | 0.005065 |
| 182 | 189 | DVITSTSF | L482C | 60 | 1.395332 | 0.050070 |
| 182 | 189 | DVITSTSF | L482C_PGM | 0 | 0.000000 | 0.000000 |
| 182 | 189 | DVITSTSF | L482C_PGM | 5 | 0.977410 | 0.030970 |

| | | | | | | |
|---|---|---|---|---|---|---|
| 182 | 189 | DVITSTSF | L482C_PGM | 60 | 1.724622 | 0.026218 |
| 183 | 189 | VITSTSF | L482C | 0 | 0.000000 | 0.000000 |
| 183 | 189 | VITSTSF | L482C | 5 | 0.570826 | 0.014135 |
| 183 | 189 | VITSTSF | L482C | 60 | 1.104976 | 0.053746 |
| 183 | 189 | VITSTSF | L482C_PGM | 0 | 0.000000 | 0.000000 |
| 183 | 189 | VITSTSF | L482C_PGM | 5 | 0.718194 | 0.025396 |
| 183 | 189 | VITSTSF | L482C_PGM | 60 | 1.405974 | 0.018779 |
| 183 | 192 | VITSTSFGVN | L482C | 0 | 0.000000 | 0.000000 |
| 183 | 192 | VITSTSFGVN | L482C | 5 | 1.533605 | 0.099871 |
| 183 | 192 | VITSTSFGVN | L482C | 60 | 2.522071 | 0.111497 |
| 183 | 192 | VITSTSFGVN | L482C_PGM | 0 | 0.000000 | 0.000000 |
| 183 | 192 | VITSTSFGVN | L482C_PGM | 5 | 1.702032 | 0.064890 |
| 183 | 192 | VITSTSFGVN | L482C_PGM | 60 | 2.891752 | 0.098616 |
| 189 | 212 | FGVNIDSLNNPQDPFVENTKKLLR | L482C | 0 | 0.000000 | 0.000000 |
| 189 | 212 | FGVNIDSLNNPQDPFVENTKKLLR | L482C | 5 | 4.824950 | 0.291555 |
| 189 | 212 | FGVNIDSLNNPQDPFVENTKKLLR | L482C | 60 | 5.483400 | 0.160009 |
| 189 | 212 | FGVNIDSLNNPQDPFVENTKKLLR | L482C_PGM | 0 | 0.000000 | 0.000000 |
| 189 | 212 | FGVNIDSLNNPQDPFVENTKKLLR | L482C_PGM | 5 | 7.297207 | 0.143334 |
| 189 | 212 | FGVNIDSLNNPQDPFVENTKKLLR | L482C_PGM | 60 | 7.682628 | 0.326124 |
| 221 | 226 | LSITVF | L482C | 0 | 0.000000 | 0.000000 |
| 221 | 226 | LSITVF | L482C | 5 | 0.386881 | 0.045091 |
| 221 | 226 | LSITVF | L482C | 60 | 1.328673 | 0.033790 |
| 221 | 226 | LSITVF | L482C_PGM | 0 | 0.000000 | 0.000000 |
| 221 | 226 | LSITVF | L482C_PGM | 5 | 0.499371 | 0.042624 |
| 221 | 226 | LSITVF | L482C_PGM | 60 | 1.398840 | 0.068099 |
| 230 | 235 | IPILEV | L482C | 0 | 0.000000 | 0.000000 |
| 230 | 235 | IPILEV | L482C | 5 | 0.269470 | 0.041875 |
| 230 | 235 | IPILEV | L482C | 60 | 0.983966 | 0.030589 |
| 230 | 235 | IPILEV | L482C_PGM | 0 | 0.000000 | 0.000000 |
| 230 | 235 | IPILEV | L482C_PGM | 5 | 0.336749 | 0.085801 |
| 230 | 235 | IPILEV | L482C_PGM | 60 | 1.463912 | 0.087465 |
| 233 | 247 | LEVLNISVFPREVTN | L482C | 0 | 0.000000 | 0.000000 |
| 233 | 247 | LEVLNISVFPREVTN | L482C | 5 | 2.638155 | 0.022891 |
| 233 | 247 | LEVLNISVFPREVTN | L482C | 60 | 3.723859 | 0.080593 |
| 233 | 247 | LEVLNISVFPREVTN | L482C_PGM | 0 | 0.000000 | 0.000000 |
| 233 | 247 | LEVLNISVFPREVTN | L482C_PGM | 5 | 2.628952 | 0.013090 |
| 233 | 247 | LEVLNISVFPREVTN | L482C_PGM | 60 | 3.878822 | 0.060312 |
| 235 | 241 | VLNISVF | L482C | 0 | 0.000000 | 0.000000 |
| 235 | 241 | VLNISVF | L482C | 5 | 1.932007 | 0.023312 |
| 235 | 241 | VLNISVF | L482C | 60 | 2.826905 | 0.079272 |
| 235 | 241 | VLNISVF | L482C_PGM | 0 | 0.000000 | 0.000000 |
| 235 | 241 | VLNISVF | L482C_PGM | 5 | 2.162929 | 0.020084 |
| 235 | 241 | VLNISVF | L482C_PGM | 60 | 2.982742 | 0.077758 |
| 236 | 241 | LNISVF | L482C | 0 | 0.000000 | 0.000000 |
| 236 | 241 | LNISVF | L482C | 5 | 1.544575 | 0.038091 |

| | | | | | | |
|---|---|---|---|---|---|---|
| 236 | 241 | LNISVF | L482C | 60 | 2.259740 | 0.044666 |
| 236 | 241 | LNISVF | L482C_PGM | 0 | 0.000000 | 0.000000 |
| 236 | 241 | LNISVF | L482C_PGM | 5 | 1.734723 | 0.020748 |
| 236 | 241 | LNISVF | L482C_PGM | 60 | 2.407191 | 0.085952 |
| 237 | 248 | NISVFPREVTNF | L482C | 0 | 0.000000 | 0.000000 |
| 237 | 248 | NISVFPREVTNF | L482C | 5 | 3.661798 | 0.040358 |
| 237 | 248 | NISVFPREVTNF | L482C | 60 | 4.154430 | 0.079639 |
| 237 | 248 | NISVFPREVTNF | L482C_PGM | 0 | 0.000000 | 0.000000 |
| 237 | 248 | NISVFPREVTNF | L482C_PGM | 5 | 3.705388 | 0.087399 |
| 237 | 248 | NISVFPREVTNF | L482C_PGM | 60 | 4.081992 | 0.114772 |
| 249 | 261 | LRKSVKRMKESRL | L482C | 0 | 0.000000 | 0.000000 |
| 249 | 261 | LRKSVKRMKESRL | L482C | 5 | 1.593092 | 0.051242 |
| 249 | 261 | LRKSVKRMKESRL | L482C | 60 | 1.862768 | 0.136498 |
| 249 | 261 | LRKSVKRMKESRL | L482C_PGM | 0 | 0.000000 | 0.000000 |
| 249 | 261 | LRKSVKRMKESRL | L482C_PGM | 5 | 1.561075 | 0.072746 |
| 249 | 261 | LRKSVKRMKESRL | L482C_PGM | 60 | 1.943917 | 0.165198 |
| 262 | 271 | EDTQKHRVDF | L482C | 0 | 0.000000 | 0.000000 |
| 262 | 271 | EDTQKHRVDF | L482C | 5 | 0.997876 | 0.022357 |
| 262 | 271 | EDTQKHRVDF | L482C | 60 | 1.131472 | 0.092307 |
| 262 | 271 | EDTQKHRVDF | L482C_PGM | 0 | 0.000000 | 0.000000 |
| 262 | 271 | EDTQKHRVDF | L482C_PGM | 5 | 1.067297 | 0.032993 |
| 262 | 271 | EDTQKHRVDF | L482C_PGM | 60 | 1.195983 | 0.069497 |
| 275 | 295 | MIDSQNSKETESHKALSDLEL | L482C | 0 | 0.000000 | 0.000000 |
| 275 | 295 | MIDSQNSKETESHKALSDLEL | L482C | 5 | 3.610808 | 0.064540 |
| 275 | 295 | MIDSQNSKETESHKALSDLEL | L482C | 60 | 4.489154 | 0.155572 |
| 275 | 295 | MIDSQNSKETESHKALSDLEL | L482C_PGM | 0 | 0.000000 | 0.000000 |
| 275 | 295 | MIDSQNSKETESHKALSDLEL | L482C_PGM | 5 | 4.023199 | 0.140253 |
| 275 | 295 | MIDSQNSKETESHKALSDLEL | L482C_PGM | 60 | 4.600851 | 0.201702 |
| 277 | 294 | DSQNSKETESHKALSDLE | L482C | 0 | 0.000000 | 0.000000 |
| 277 | 294 | DSQNSKETESHKALSDLE | L482C | 5 | 3.037710 | 0.029498 |
| 277 | 294 | DSQNSKETESHKALSDLE | L482C | 60 | 3.128544 | 0.218590 |
| 277 | 294 | DSQNSKETESHKALSDLE | L482C_PGM | 0 | 0.000000 | 0.000000 |
| 277 | 294 | DSQNSKETESHKALSDLE | L482C_PGM | 5 | 3.191096 | 0.053929 |
| 277 | 294 | DSQNSKETESHKALSDLE | L482C_PGM | 60 | 3.287211 | 0.134542 |
| 296 | 302 | VAQSIIF | L482C | 0 | 0.000000 | 0.000000 |
| 296 | 302 | VAQSIIF | L482C | 5 | -0.216330 | 0.055165 |
| 296 | 302 | VAQSIIF | L482C | 60 | 0.185035 | 0.034846 |
| 296 | 302 | VAQSIIF | L482C_PGM | 0 | 0.000000 | 0.000000 |
| 296 | 302 | VAQSIIF | L482C_PGM | 5 | -0.187627 | 0.056919 |
| 296 | 302 | VAQSIIF | L482C_PGM | 60 | 0.295947 | 0.064321 |
| 301 | 306 | IFIFAG | L482C | 0 | 0.000000 | 0.000000 |
| 301 | 306 | IFIFAG | L482C | 5 | 0.110080 | 0.029934 |
| 301 | 306 | IFIFAG | L482C | 60 | 0.261989 | 0.026914 |
| 301 | 306 | IFIFAG | L482C_PGM | 0 | 0.000000 | 0.000000 |
| 301 | 306 | IFIFAG | L482C_PGM | 5 | 0.128883 | 0.020840 |

| | | | | | | |
|---|---|---|---|---|---|---|
| 301 | 306 | IFIFAG | L482C_PGM | 60 | 0.363621 | 0.046597 |
| 307 | 313 | YETTSSV | L482C | 0 | 0.000000 | 0.000000 |
| 307 | 313 | YETTSSV | L482C | 5 | 1.121221 | 0.059132 |
| 307 | 313 | YETTSSV | L482C | 60 | 1.666031 | 0.058619 |
| 307 | 313 | YETTSSV | L482C_PGM | 0 | 0.000000 | 0.000000 |
| 307 | 313 | YETTSSV | L482C_PGM | 5 | 1.057025 | 0.040025 |
| 307 | 313 | YETTSSV | L482C_PGM | 60 | 1.630786 | 0.071983 |
| 307 | 314 | YETTSSVL | L482C | 0 | 0.000000 | 0.000000 |
| 307 | 314 | YETTSSVL | L482C | 5 | 1.112940 | 0.041980 |
| 307 | 314 | YETTSSVL | L482C | 60 | 1.596444 | 0.059713 |
| 307 | 314 | YETTSSVL | L482C_PGM | 0 | 0.000000 | 0.000000 |
| 307 | 314 | YETTSSVL | L482C_PGM | 5 | 1.121532 | 0.016097 |
| 307 | 314 | YETTSSVL | L482C_PGM | 60 | 1.764417 | 0.129633 |
| 307 | 316 | YETTSSVLSF | L482C | 0 | 0.000000 | 0.000000 |
| 307 | 316 | YETTSSVLSF | L482C | 5 | 0.864405 | 0.020020 |
| 307 | 316 | YETTSSVLSF | L482C | 60 | 1.445339 | 0.045973 |
| 307 | 316 | YETTSSVLSF | L482C_PGM | 0 | 0.000000 | 0.000000 |
| 307 | 316 | YETTSSVLSF | L482C_PGM | 5 | 0.864339 | 0.016103 |
| 307 | 316 | YETTSSVLSF | L482C_PGM | 60 | 1.561195 | 0.021047 |
| 317 | 333 | IMYELATHPDVQQKLQE | L482C | 0 | 0.000000 | 0.000000 |
| 317 | 333 | IMYELATHPDVQQKLQE | L482C | 5 | 0.459008 | 0.077486 |
| 317 | 333 | IMYELATHPDVQQKLQE | L482C | 60 | 1.080432 | 0.082395 |
| 317 | 333 | IMYELATHPDVQQKLQE | L482C_PGM | 0 | 0.000000 | 0.000000 |
| 317 | 333 | IMYELATHPDVQQKLQE | L482C_PGM | 5 | 0.426575 | 0.052074 |
| 317 | 333 | IMYELATHPDVQQKLQE | L482C_PGM | 60 | 1.098536 | 0.039581 |
| 319 | 333 | YELATHPDVQQKLQE | L482C | 0 | 0.000000 | 0.000000 |
| 319 | 333 | YELATHPDVQQKLQE | L482C | 5 | 0.436939 | 0.035411 |
| 319 | 333 | YELATHPDVQQKLQE | L482C | 60 | 0.913394 | 0.030888 |
| 319 | 333 | YELATHPDVQQKLQE | L482C_PGM | 0 | 0.000000 | 0.000000 |
| 319 | 333 | YELATHPDVQQKLQE | L482C_PGM | 5 | 0.489654 | 0.056518 |
| 319 | 333 | YELATHPDVQQKLQE | L482C_PGM | 60 | 1.034379 | 0.028110 |
| 322 | 333 | ATHPDVQQKLQE | L482C | 0 | 0.000000 | 0.000000 |
| 322 | 333 | ATHPDVQQKLQE | L482C | 5 | 0.104818 | 0.031585 |
| 322 | 333 | ATHPDVQQKLQE | L482C | 60 | 0.647350 | 0.045381 |
| 322 | 333 | ATHPDVQQKLQE | L482C_PGM | 0 | 0.000000 | 0.000000 |
| 322 | 333 | ATHPDVQQKLQE | L482C_PGM | 5 | 0.144876 | 0.034022 |
| 322 | 333 | ATHPDVQQKLQE | L482C_PGM | 60 | 0.664361 | 0.041209 |
| 332 | 337 | QEEIDA | L482C | 0 | 0.000000 | 0.000000 |
| 332 | 337 | QEEIDA | L482C | 5 | 0.378381 | 0.046604 |
| 332 | 337 | QEEIDA | L482C | 60 | 1.173704 | 0.068759 |
| 332 | 337 | QEEIDA | L482C_PGM | 0 | 0.000000 | 0.000000 |
| 332 | 337 | QEEIDA | L482C_PGM | 5 | 0.376064 | 0.054189 |
| 332 | 337 | QEEIDA | L482C_PGM | 60 | 1.082840 | 0.011785 |
| 337 | 349 | AVLPNKAPPTYDT | L482C | 0 | 0.000000 | 0.000000 |
| 337 | 349 | AVLPNKAPPTYDT | L482C | 5 | 3.013135 | 0.086046 |

| | | | | | | |
|---|---|---|---|---|---|---|
| 337 | 349 | AVLPNKAPPTYDT | L482C | 60 | 3.540894 | 0.156601 |
| 337 | 349 | AVLPNKAPPTYDT | L482C_PGM | 0 | 0.000000 | 0.000000 |
| 337 | 349 | AVLPNKAPPTYDT | L482C_PGM | 5 | 2.927637 | 0.078028 |
| 337 | 349 | AVLPNKAPPTYDT | L482C_PGM | 60 | 3.540158 | 0.165326 |
| 340 | 347 | PNKAPPTY | L482C | 0 | 0.000000 | 0.000000 |
| 340 | 347 | PNKAPPTY | L482C | 5 | 1.387862 | 0.039105 |
| 340 | 347 | PNKAPPTY | L482C | 60 | 1.798798 | 0.069893 |
| 340 | 347 | PNKAPPTY | L482C_PGM | 0 | 0.000000 | 0.000000 |
| 340 | 347 | PNKAPPTY | L482C_PGM | 5 | 1.345041 | 0.034864 |
| 340 | 347 | PNKAPPTY | L482C_PGM | 60 | 1.738152 | 0.060652 |
| 348 | 353 | DTVLQM | L482C | 0 | 0.000000 | 0.000000 |
| 348 | 353 | DTVLQM | L482C | 5 | 0.844818 | 0.023987 |
| 348 | 353 | DTVLQM | L482C | 60 | 0.801759 | 0.017421 |
| 348 | 353 | DTVLQM | L482C_PGM | 0 | 0.000000 | 0.000000 |
| 348 | 353 | DTVLQM | L482C_PGM | 5 | 0.846373 | 0.028648 |
| 348 | 353 | DTVLQM | L482C_PGM | 60 | 0.848782 | 0.039273 |
| 357 | 363 | DMVVNET | L482C | 0 | 0.000000 | 0.000000 |
| 357 | 363 | DMVVNET | L482C | 5 | 0.013218 | 0.026025 |
| 357 | 363 | DMVVNET | L482C | 60 | -0.001726 | 0.025824 |
| 357 | 363 | DMVVNET | L482C_PGM | 0 | 0.000000 | 0.000000 |
| 357 | 363 | DMVVNET | L482C_PGM | 5 | -0.028601 | 0.032684 |
| 357 | 363 | DMVVNET | L482C_PGM | 60 | 0.037300 | 0.025207 |
| 359 | 366 | VVNETLRL | L482C | 0 | 0.000000 | 0.000000 |
| 359 | 366 | VVNETLRL | L482C | 5 | 0.000330 | 0.013131 |
| 359 | 366 | VVNETLRL | L482C | 60 | 0.054103 | 0.009210 |
| 359 | 366 | VVNETLRL | L482C_PGM | 0 | 0.000000 | 0.000000 |
| 359 | 366 | VVNETLRL | L482C_PGM | 5 | 0.009377 | 0.008098 |
| 359 | 366 | VVNETLRL | L482C_PGM | 60 | 0.047237 | 0.027243 |
| 361 | 366 | NETLRL | L482C | 0 | 0.000000 | 0.000000 |
| 361 | 366 | NETLRL | L482C | 5 | -0.015641 | 0.006268 |
| 361 | 366 | NETLRL | L482C | 60 | 0.016347 | 0.006724 |
| 361 | 366 | NETLRL | L482C_PGM | 0 | 0.000000 | 0.000000 |
| 361 | 366 | NETLRL | L482C_PGM | 5 | -0.020020 | 0.007065 |
| 361 | 366 | NETLRL | L482C_PGM | 60 | 0.019574 | 0.008251 |
| 363 | 370 | TLRLFPIA | L482C | 0 | 0.000000 | 0.000000 |
| 363 | 370 | TLRLFPIA | L482C | 5 | 0.136572 | 0.032869 |
| 363 | 370 | TLRLFPIA | L482C | 60 | 0.355534 | 0.035712 |
| 363 | 370 | TLRLFPIA | L482C_PGM | 0 | 0.000000 | 0.000000 |
| 363 | 370 | TLRLFPIA | L482C_PGM | 5 | 0.109496 | 0.056588 |
| 363 | 370 | TLRLFPIA | L482C_PGM | 60 | 0.323916 | 0.027833 |
| 374 | 385 | ERVCKKDVEING | L482C | 0 | 0.000000 | 0.000000 |
| 374 | 385 | ERVCKKDVEING | L482C | 5 | 1.186843 | 0.052175 |
| 374 | 385 | ERVCKKDVEING | L482C | 60 | 1.367429 | 0.070835 |
| 374 | 385 | ERVCKKDVEING | L482C_PGM | 0 | 0.000000 | 0.000000 |
| 374 | 385 | ERVCKKDVEING | L482C_PGM | 5 | 1.111521 | 0.045729 |

| | | | | | | |
|---|---|---|---|---|---|---|
| 374 | 385 | ERVCKKDVEING | L482C_PGM | 60 | 1.358308 | 0.053606 |
| 388 | 393 | IPKGVV | L482C | 0 | 0.000000 | 0.000000 |
| 388 | 393 | IPKGVV | L482C | 5 | 0.310686 | 0.022815 |
| 388 | 393 | IPKGVV | L482C | 60 | 0.537246 | 0.031966 |
| 388 | 393 | IPKGVV | L482C_PGM | 0 | 0.000000 | 0.000000 |
| 388 | 393 | IPKGVV | L482C_PGM | 5 | 0.340907 | 0.017372 |
| 388 | 393 | IPKGVV | L482C_PGM | 60 | 0.537737 | 0.022341 |
| 388 | 395 | IPKGVVVM | L482C | 0 | 0.000000 | 0.000000 |
| 388 | 395 | IPKGVVVM | L482C | 5 | 0.483444 | 0.016893 |
| 388 | 395 | IPKGVVVM | L482C | 60 | 0.785765 | 0.006417 |
| 388 | 395 | IPKGVVVM | L482C_PGM | 0 | 0.000000 | 0.000000 |
| 388 | 395 | IPKGVVVM | L482C_PGM | 5 | 0.501583 | 0.019404 |
| 388 | 395 | IPKGVVVM | L482C_PGM | 60 | 0.841876 | 0.012668 |
| 388 | 401 | IPKGVVVMIPSYAL | L482C | 0 | 0.000000 | 0.000000 |
| 388 | 401 | IPKGVVVMIPSYAL | L482C | 5 | 0.499614 | 0.198259 |
| 388 | 401 | IPKGVVVMIPSYAL | L482C | 60 | 0.597402 | 0.022723 |
| 388 | 401 | IPKGVVVMIPSYAL | L482C_PGM | 0 | 0.000000 | 0.000000 |
| 388 | 401 | IPKGVVVMIPSYAL | L482C_PGM | 5 | 0.633872 | 0.075102 |
| 388 | 401 | IPKGVVVMIPSYAL | L482C_PGM | 60 | 0.786948 | 0.078714 |
| 394 | 401 | VMIPSYAL | L482C | 0 | 0.000000 | 0.000000 |
| 394 | 401 | VMIPSYAL | L482C | 5 | 0.287875 | 0.052687 |
| 394 | 401 | VMIPSYAL | L482C | 60 | 0.498855 | 0.040063 |
| 394 | 401 | VMIPSYAL | L482C_PGM | 0 | 0.000000 | 0.000000 |
| 394 | 401 | VMIPSYAL | L482C_PGM | 5 | 0.317196 | 0.051249 |
| 394 | 401 | VMIPSYAL | L482C_PGM | 60 | 0.539635 | 0.050128 |
| 396 | 401 | IPSYAL | L482C | 0 | 0.000000 | 0.000000 |
| 396 | 401 | IPSYAL | L482C | 5 | 0.386400 | 0.044461 |
| 396 | 401 | IPSYAL | L482C | 60 | 0.519725 | 0.034658 |
| 396 | 401 | IPSYAL | L482C_PGM | 0 | 0.000000 | 0.000000 |
| 396 | 401 | IPSYAL | L482C_PGM | 5 | 0.405527 | 0.050658 |
| 396 | 401 | IPSYAL | L482C_PGM | 60 | 0.553179 | 0.034137 |
| 400 | 407 | ALHRDPKY | L482C | 0 | 0.000000 | 0.000000 |
| 400 | 407 | ALHRDPKY | L482C | 5 | 0.454361 | 0.023334 |
| 400 | 407 | ALHRDPKY | L482C | 60 | 0.512452 | 0.050402 |
| 400 | 407 | ALHRDPKY | L482C_PGM | 0 | 0.000000 | 0.000000 |
| 400 | 407 | ALHRDPKY | L482C_PGM | 5 | 0.428560 | 0.013514 |
| 400 | 407 | ALHRDPKY | L482C_PGM | 60 | 0.578119 | 0.021271 |
| 402 | 414 | HRDPKYWTEPEKF | L482C | 0 | 0.000000 | 0.000000 |
| 402 | 414 | HRDPKYWTEPEKF | L482C | 5 | 2.790456 | 0.124969 |
| 402 | 414 | HRDPKYWTEPEKF | L482C | 60 | 2.962024 | 0.137121 |
| 402 | 414 | HRDPKYWTEPEKF | L482C_PGM | 0 | 0.000000 | 0.000000 |
| 402 | 414 | HRDPKYWTEPEKF | L482C_PGM | 5 | 2.732587 | 0.061638 |
| 402 | 414 | HRDPKYWTEPEKF | L482C_PGM | 60 | 2.946017 | 0.069041 |
| 408 | 414 | WTEPEKF | L482C | 0 | 0.000000 | 0.000000 |
| 408 | 414 | WTEPEKF | L482C | 5 | 1.662944 | 0.033702 |

| | | | | | | |
|---|---|---|---|---|---|---|
| 408 | 414 | WTEPEKF | L482C | 60 | 1.742071 | 0.065075 |
| 408 | 414 | WTEPEKF | L482C_PGM | 0 | 0.000000 | 0.000000 |
| 408 | 414 | WTEPEKF | L482C_PGM | 5 | 1.732372 | 0.054026 |
| 408 | 414 | WTEPEKF | L482C_PGM | 60 | 1.773688 | 0.072797 |
| 415 | 444 | LPERFSKKNKDNIDPYIYTPFGSGP | L482C | 0 | 0.000000 | 0.000000 |
| 415 | 444 | LPERFSKKNKDNIDPYIYTPFGSGP | L482C | 5 | 3.027038 | 0.290093 |
| 415 | 444 | LPERFSKKNKDNIDPYIYTPFGSGP | L482C | 60 | 3.780067 | 0.094681 |
| 415 | 444 | LPERFSKKNKDNIDPYIYTPFGSGP | L482C_PGM | 0 | 0.000000 | 0.000000 |
| 415 | 444 | LPERFSKKNKDNIDPYIYTPFGSGP | L482C_PGM | 5 | 3.080907 | 0.109088 |
| 415 | 444 | LPERFSKKNKDNIDPYIYTPFGSGP | L482C_PGM | 60 | 3.777966 | 0.197780 |
| 420 | 444 | SKKNKDNIDPYIYTPFGSGPRNCIG | L482C | 0 | 0.000000 | 0.000000 |
| 420 | 444 | SKKNKDNIDPYIYTPFGSGPRNCIG | L482C | 5 | 3.153848 | 0.081666 |
| 420 | 444 | SKKNKDNIDPYIYTPFGSGPRNCIG | L482C | 60 | 3.867041 | 0.094193 |
| 420 | 444 | SKKNKDNIDPYIYTPFGSGPRNCIG | L482C_PGM | 0 | 0.000000 | 0.000000 |
| 420 | 444 | SKKNKDNIDPYIYTPFGSGPRNCIG | L482C_PGM | 5 | 3.200329 | 0.091599 |
| 420 | 444 | SKKNKDNIDPYIYTPFGSGPRNCIG | L482C_PGM | 60 | 4.036088 | 0.171418 |
| 445 | 452 | MRFALMNM | L482C | 0 | 0.000000 | 0.000000 |
| 445 | 452 | MRFALMNM | L482C | 5 | 0.455932 | 0.042574 |
| 445 | 452 | MRFALMNM | L482C | 60 | 0.592883 | 0.014080 |
| 445 | 452 | MRFALMNM | L482C_PGM | 0 | 0.000000 | 0.000000 |
| 445 | 452 | MRFALMNM | L482C_PGM | 5 | 0.388625 | 0.047426 |
| 445 | 452 | MRFALMNM | L482C_PGM | 60 | 0.673714 | 0.029610 |
| 446 | 452 | RFALMNM | L482C | 0 | 0.000000 | 0.000000 |
| 446 | 452 | RFALMNM | L482C | 5 | 0.080620 | 0.027781 |
| 446 | 452 | RFALMNM | L482C | 60 | 0.154752 | 0.048633 |
| 446 | 452 | RFALMNM | L482C_PGM | 0 | 0.000000 | 0.000000 |
| 446 | 452 | RFALMNM | L482C_PGM | 5 | 0.095446 | 0.022369 |
| 446 | 452 | RFALMNM | L482C_PGM | 60 | 0.288897 | 0.018419 |
| 455 | 463 | ALIRVLQNF | L482C | 0 | 0.000000 | 0.000000 |
| 455 | 463 | ALIRVLQNF | L482C | 5 | 0.025531 | 0.032317 |
| 455 | 463 | ALIRVLQNF | L482C | 60 | 0.297651 | 0.028206 |
| 455 | 463 | ALIRVLQNF | L482C_PGM | 0 | 0.000000 | 0.000000 |
| 455 | 463 | ALIRVLQNF | L482C_PGM | 5 | 0.059151 | 0.028652 |
| 455 | 463 | ALIRVLQNF | L482C_PGM | 60 | 0.314722 | 0.024076 |
| 457 | 463 | IRVLQNF | L482C | 0 | 0.000000 | 0.000000 |
| 457 | 463 | IRVLQNF | L482C | 5 | -0.056035 | 0.037638 |
| 457 | 463 | IRVLQNF | L482C | 60 | 0.136350 | 0.028469 |
| 457 | 463 | IRVLQNF | L482C_PGM | 0 | 0.000000 | 0.000000 |
| 457 | 463 | IRVLQNF | L482C_PGM | 5 | -0.041176 | 0.026814 |
| 457 | 463 | IRVLQNF | L482C_PGM | 60 | 0.449387 | 0.036815 |
| 463 | 477 | FSFKPGKETQIPLKL | L482C | 0 | 0.000000 | 0.000000 |
| 463 | 477 | FSFKPGKETQIPLKL | L482C | 5 | 3.472773 | 0.089038 |
| 463 | 477 | FSFKPGKETQIPLKL | L482C | 60 | 3.349505 | 0.168138 |
| 463 | 477 | FSFKPGKETQIPLKL | L482C_PGM | 0 | 0.000000 | 0.000000 |
| 463 | 477 | FSFKPGKETQIPLKL | L482C_PGM | 5 | 3.426193 | 0.098763 |

| | | | | | | |
|---|---|---|---|---|---|---|
| 463 | 477 | FSFKPGKETQIPLKL | L482C_PGM | 60 | 3.489610 | 0.113776 |
| 464 | 479 | SFKPGKETQIPLKLSL | L482C | 0 | 0.000000 | 0.000000 |
| 464 | 479 | SFKPGKETQIPLKLSL | L482C | 5 | 4.371998 | 0.118044 |
| 464 | 479 | SFKPGKETQIPLKLSL | L482C | 60 | 4.286432 | 0.244262 |
| 464 | 479 | SFKPGKETQIPLKLSL | L482C_PGM | 0 | 0.000000 | 0.000000 |
| 464 | 479 | SFKPGKETQIPLKLSL | L482C_PGM | 5 | 4.291338 | 0.086362 |
| 464 | 479 | SFKPGKETQIPLKLSL | L482C_PGM | 60 | 4.709682 | 0.096131 |
| 492 | 507 | KVESRDGTVSGAHHHH | L482C | 0 | 0.000000 | 0.000000 |
| 492 | 507 | KVESRDGTVSGAHHHH | L482C | 5 | 1.150868 | 0.074241 |
| 492 | 507 | KVESRDGTVSGAHHHH | L482C | 60 | 1.151307 | 0.141238 |
| 492 | 507 | KVESRDGTVSGAHHHH | L482C_PGM | 0 | 0.000000 | 0.000000 |
| 492 | 507 | KVESRDGTVSGAHHHH | L482C_PGM | 5 | 1.205424 | 0.093238 |
| 492 | 507 | KVESRDGTVSGAHHHH | L482C_PGM | 60 | 1.207081 | 0.065279 |
| 494 | 507 | ESRDGTVSGAHHHH | L482C | 0 | 0.000000 | 0.000000 |
| 494 | 507 | ESRDGTVSGAHHHH | L482C | 5 | 1.165280 | 0.114562 |
| 494 | 507 | ESRDGTVSGAHHHH | L482C | 60 | 0.942264 | 0.072176 |
| 494 | 507 | ESRDGTVSGAHHHH | L482C_PGM | 0 | 0.000000 | 0.000000 |
| 494 | 507 | ESRDGTVSGAHHHH | L482C_PGM | 5 | 1.066168 | 0.101482 |
| 494 | 507 | ESRDGTVSGAHHHH | L482C_PGM | 60 | 1.077652 | 0.173309 |

| Deuterium uptake data for F215C and F215C_PGM CYP3A4 states | | | | | | |
|---|---|---|---|---|---|---|
| **Start** | **End** | **Sequence** | **State** | **Exposure (min)** | **Uptake (Da)** | **Uptake SD (Da)** |
| 33 | 48 | FKKLGIPGPTPLPFLG | F215C | 0 | 0 | 0 |
| 33 | 48 | FKKLGIPGPTPLPFLG | F215C | 0.5 | 0.5193 | 0.0572 |
| 33 | 48 | FKKLGIPGPTPLPFLG | F215C | 1 | 0.5007 | 0.0957 |
| 33 | 48 | FKKLGIPGPTPLPFLG | F215C | 2 | 0.6429 | 0.1597 |
| 33 | 48 | FKKLGIPGPTPLPFLG | F215C | 5 | 0.5025 | 0.1053 |
| 33 | 48 | FKKLGIPGPTPLPFLG | F215C | 30 | 0.3888 | 0.0813 |
| 33 | 48 | FKKLGIPGPTPLPFLG | F215C | 60 | 0.4382 | 0.0852 |
| 33 | 48 | FKKLGIPGPTPLPFLG | F215C | 240 | 0.3019 | 0.0681 |
| 33 | 48 | FKKLGIPGPTPLPFLG | F215C_PGM | 0 | 0 | 0 |
| 33 | 48 | FKKLGIPGPTPLPFLG | F215C_PGM | 0.5 | 0.2703 | 0.1348 |
| 33 | 48 | FKKLGIPGPTPLPFLG | F215C_PGM | 1 | 0.3564 | 0.0999 |
| 33 | 48 | FKKLGIPGPTPLPFLG | F215C_PGM | 2 | 0.2619 | 0.1241 |
| 33 | 48 | FKKLGIPGPTPLPFLG | F215C_PGM | 5 | 0.2551 | 0.1812 |
| 33 | 48 | FKKLGIPGPTPLPFLG | F215C_PGM | 30 | 0.3006 | 0.103 |
| 33 | 48 | FKKLGIPGPTPLPFLG | F215C_PGM | 60 | 0.3437 | 0.1494 |
| 33 | 48 | FKKLGIPGPTPLPFLG | F215C_PGM | 240 | 0.2631 | 0.1111 |
| 33 | 51 | FKKLGIPGPTPLPFLGNIL | F215C | 0 | 0 | 0 |
| 33 | 51 | FKKLGIPGPTPLPFLGNIL | F215C | 0.5 | 0.6263 | 0.0671 |
| 33 | 51 | FKKLGIPGPTPLPFLGNIL | F215C | 1 | 0.6872 | 0.1341 |
| 33 | 51 | FKKLGIPGPTPLPFLGNIL | F215C | 2 | 0.7247 | 0.1431 |
| 33 | 51 | FKKLGIPGPTPLPFLGNIL | F215C | 5 | 0.4744 | 0.101 |
| 33 | 51 | FKKLGIPGPTPLPFLGNIL | F215C | 30 | 0.48 | 0.102 |

| | | | | | | |
|---|---|---|---|---|---|---|
| 33 | 51 | FKKLGIPGPTPLPFLGNIL | F215C | 60 | 0.3609 | 0.1689 |
| 33 | 51 | FKKLGIPGPTPLPFLGNIL | F215C | 240 | 0.7799 | 0.0196 |
| 33 | 51 | FKKLGIPGPTPLPFLGNIL | F215C_PGM | 0 | 0 | 0 |
| 33 | 51 | FKKLGIPGPTPLPFLGNIL | F215C_PGM | 0.5 | 0.6922 | 0.1003 |
| 33 | 51 | FKKLGIPGPTPLPFLGNIL | F215C_PGM | 1 | 0.6783 | 0.0196 |
| 33 | 51 | FKKLGIPGPTPLPFLGNIL | F215C_PGM | 2 | 0.5276 | 0.1322 |
| 33 | 51 | FKKLGIPGPTPLPFLGNIL | F215C_PGM | 5 | 0.48 | 0.2296 |
| 33 | 51 | FKKLGIPGPTPLPFLGNIL | F215C_PGM | 30 | 0.5525 | 0.0634 |
| 33 | 51 | FKKLGIPGPTPLPFLGNIL | F215C_PGM | 60 | 0.5924 | 0.1366 |
| 33 | 51 | FKKLGIPGPTPLPFLGNIL | F215C_PGM | 240 | 0.3462 | 0.02 |
| 34 | 48 | KKLGIPGPTPLPFLG | F215C | 0 | 0 | 0 |
| 34 | 48 | KKLGIPGPTPLPFLG | F215C | 0.5 | 0.347 | 0.069 |
| 34 | 48 | KKLGIPGPTPLPFLG | F215C | 1 | 0.3879 | 0.0252 |
| 34 | 48 | KKLGIPGPTPLPFLG | F215C | 2 | 0.5176 | 0.1042 |
| 34 | 48 | KKLGIPGPTPLPFLG | F215C | 5 | 0.4364 | 0.0797 |
| 34 | 48 | KKLGIPGPTPLPFLG | F215C | 30 | 0.3802 | 0.0535 |
| 34 | 48 | KKLGIPGPTPLPFLG | F215C | 60 | 0.4083 | 0.0365 |
| 34 | 48 | KKLGIPGPTPLPFLG | F215C | 240 | 0.4521 | 0.0258 |
| 34 | 48 | KKLGIPGPTPLPFLG | F215C_PGM | 0 | 0 | 0 |
| 34 | 48 | KKLGIPGPTPLPFLG | F215C_PGM | 0.5 | 0.3129 | 0.1012 |
| 34 | 48 | KKLGIPGPTPLPFLG | F215C_PGM | 1 | 0.3897 | 0.0948 |
| 34 | 48 | KKLGIPGPTPLPFLG | F215C_PGM | 2 | 0.3073 | 0.091 |
| 34 | 48 | KKLGIPGPTPLPFLG | F215C_PGM | 5 | 0.3403 | 0.0755 |
| 34 | 48 | KKLGIPGPTPLPFLG | F215C_PGM | 30 | 0.3528 | 0.0813 |
| 34 | 48 | KKLGIPGPTPLPFLG | F215C_PGM | 60 | 0.3569 | 0.0964 |
| 34 | 48 | KKLGIPGPTPLPFLG | F215C_PGM | 240 | 0.2884 | 0.0613 |
| 34 | 51 | KKLGIPGPTPLPFLGNIL | F215C | 0 | 0 | 0 |
| 34 | 51 | KKLGIPGPTPLPFLGNIL | F215C | 0.5 | 0.613 | 0.0868 |
| 34 | 51 | KKLGIPGPTPLPFLGNIL | F215C | 1 | 0.5904 | 0.0329 |
| 34 | 51 | KKLGIPGPTPLPFLGNIL | F215C | 2 | 0.6697 | 0.1709 |
| 34 | 51 | KKLGIPGPTPLPFLGNIL | F215C | 5 | 0.4096 | 0.0489 |
| 34 | 51 | KKLGIPGPTPLPFLGNIL | F215C | 30 | 0.5371 | 0.1139 |
| 34 | 51 | KKLGIPGPTPLPFLGNIL | F215C | 60 | 0.4718 | 0.0798 |
| 34 | 51 | KKLGIPGPTPLPFLGNIL | F215C | 240 | 0.6872 | 0.0433 |
| 34 | 51 | KKLGIPGPTPLPFLGNIL | F215C_PGM | 0 | 0 | 0 |
| 34 | 51 | KKLGIPGPTPLPFLGNIL | F215C_PGM | 0.5 | 0.4151 | 0.1147 |
| 34 | 51 | KKLGIPGPTPLPFLGNIL | F215C_PGM | 1 | 0.5482 | 0.0095 |
| 34 | 51 | KKLGIPGPTPLPFLGNIL | F215C_PGM | 2 | 0.4535 | 0.1272 |
| 34 | 51 | KKLGIPGPTPLPFLGNIL | F215C_PGM | 5 | 0.385 | 0.095 |
| 34 | 51 | KKLGIPGPTPLPFLGNIL | F215C_PGM | 30 | 0.4636 | 0.0729 |
| 34 | 51 | KKLGIPGPTPLPFLGNIL | F215C_PGM | 60 | 0.4842 | 0.1242 |
| 34 | 51 | KKLGIPGPTPLPFLGNIL | F215C_PGM | 240 | 0.2703 | 0.1002 |
| 49 | 60 | NILSYHKGFTMF | F215C | 0 | 0 | 0 |
| 49 | 60 | NILSYHKGFTMF | F215C | 0.5 | 1.3922 | 0.1663 |
| 49 | 60 | NILSYHKGFTMF | F215C | 1 | 1.5356 | 0.0597 |

| | | | | | | |
|---|---|---|---|---|---|---|
| 49 | 60 | NILSYHKGFTMF | F215C | 2 | 1.8304 | 0.0643 |
| 49 | 60 | NILSYHKGFTMF | F215C | 5 | 2.0238 | 0.0478 |
| 49 | 60 | NILSYHKGFTMF | F215C | 30 | 2.0573 | 0.1738 |
| 49 | 60 | NILSYHKGFTMF | F215C | 60 | 2.0871 | 0.2257 |
| 49 | 60 | NILSYHKGFTMF | F215C | 240 | 2.03 | 0.2172 |
| 49 | 60 | NILSYHKGFTMF | F215C_PGM | 0 | 0 | 0 |
| 49 | 60 | NILSYHKGFTMF | F215C_PGM | 0.5 | 1.4077 | 0.0235 |
| 49 | 60 | NILSYHKGFTMF | F215C_PGM | 1 | 1.7449 | 0.2412 |
| 49 | 60 | NILSYHKGFTMF | F215C_PGM | 2 | 1.9609 | 0.0445 |
| 49 | 60 | NILSYHKGFTMF | F215C_PGM | 5 | 2.2788 | 0.084 |
| 49 | 60 | NILSYHKGFTMF | F215C_PGM | 30 | 2.3547 | 0.0527 |
| 49 | 60 | NILSYHKGFTMF | F215C_PGM | 60 | 2.3655 | 0.0778 |
| 49 | 60 | NILSYHKGFTMF | F215C_PGM | 240 | 2.2068 | 0.1412 |
| 54 | 59 | HKGFTM | F215C | 0 | 0 | 0 |
| 54 | 59 | HKGFTM | F215C | 0.5 | 1.094 | 0.0512 |
| 54 | 59 | HKGFTM | F215C | 1 | 1.1492 | 0.0547 |
| 54 | 59 | HKGFTM | F215C | 2 | 1.186 | 0.0792 |
| 54 | 59 | HKGFTM | F215C | 5 | 1.1756 | 0.0601 |
| 54 | 59 | HKGFTM | F215C | 30 | 1.1772 | 0.0387 |
| 54 | 59 | HKGFTM | F215C | 60 | 1.1198 | 0.0486 |
| 54 | 59 | HKGFTM | F215C | 240 | 1.1368 | 0.0803 |
| 54 | 59 | HKGFTM | F215C_PGM | 0 | 0 | 0 |
| 54 | 59 | HKGFTM | F215C_PGM | 0.5 | 1.0103 | 0.0823 |
| 54 | 59 | HKGFTM | F215C_PGM | 1 | 1.0861 | 0.059 |
| 54 | 59 | HKGFTM | F215C_PGM | 2 | 1.1066 | 0.1052 |
| 54 | 59 | HKGFTM | F215C_PGM | 5 | 1.1288 | 0.0793 |
| 54 | 59 | HKGFTM | F215C_PGM | 30 | 1.1449 | 0.0735 |
| 54 | 59 | HKGFTM | F215C_PGM | 60 | 1.1489 | 0.065 |
| 54 | 59 | HKGFTM | F215C_PGM | 240 | 1.1054 | 0.0773 |
| 61 | 73 | DMEAHKKYGKVWG | F215C | 0 | 0 | 0 |
| 61 | 73 | DMEAHKKYGKVWG | F215C | 0.5 | 0.6896 | 0.0537 |
| 61 | 73 | DMEAHKKYGKVWG | F215C | 1 | 0.6996 | 0.0593 |
| 61 | 73 | DMEAHKKYGKVWG | F215C | 2 | 0.82 | 0.0713 |
| 61 | 73 | DMEAHKKYGKVWG | F215C | 5 | 0.916 | 0.0904 |
| 61 | 73 | DMEAHKKYGKVWG | F215C | 30 | 1.4519 | 0.0575 |
| 61 | 73 | DMEAHKKYGKVWG | F215C | 60 | 1.6329 | 0.0554 |
| 61 | 73 | DMEAHKKYGKVWG | F215C | 240 | 1.9257 | 0.1002 |
| 61 | 73 | DMEAHKKYGKVWG | F215C_PGM | 0 | 0 | 0 |
| 61 | 73 | DMEAHKKYGKVWG | F215C_PGM | 0.5 | 0.6742 | 0.0471 |
| 61 | 73 | DMEAHKKYGKVWG | F215C_PGM | 1 | 0.7622 | 0.1178 |
| 61 | 73 | DMEAHKKYGKVWG | F215C_PGM | 2 | 0.8857 | 0.038 |
| 61 | 73 | DMEAHKKYGKVWG | F215C_PGM | 5 | 1.0338 | 0.0525 |
| 61 | 73 | DMEAHKKYGKVWG | F215C_PGM | 30 | 1.4766 | 0.0846 |
| 61 | 73 | DMEAHKKYGKVWG | F215C_PGM | 60 | 1.7026 | 0.1115 |
| 61 | 73 | DMEAHKKYGKVWG | F215C_PGM | 240 | 1.959 | 0.1284 |

| | | | | | | |
|---|---|---|---|---|---|---|
| 61 | 74 | DMEAHKKYGKVWGF | F215C | 0 | 0 | 0 |
| 61 | 74 | DMEAHKKYGKVWGF | F215C | 0.5 | 0.7396 | 0.0321 |
| 61 | 74 | DMEAHKKYGKVWGF | F215C | 1 | 0.7521 | 0.0432 |
| 61 | 74 | DMEAHKKYGKVWGF | F215C | 2 | 0.9378 | 0.0277 |
| 61 | 74 | DMEAHKKYGKVWGF | F215C | 5 | 1.0875 | 0.0489 |
| 61 | 74 | DMEAHKKYGKVWGF | F215C | 30 | 1.4621 | 0.0352 |
| 61 | 74 | DMEAHKKYGKVWGF | F215C | 60 | 1.5922 | 0.1205 |
| 61 | 74 | DMEAHKKYGKVWGF | F215C | 240 | 1.9573 | 0.0863 |
| 61 | 74 | DMEAHKKYGKVWGF | F215C_PGM | 0 | 0 | 0 |
| 61 | 74 | DMEAHKKYGKVWGF | F215C_PGM | 0.5 | 0.6896 | 0.0526 |
| 61 | 74 | DMEAHKKYGKVWGF | F215C_PGM | 1 | 0.7648 | 0.1143 |
| 61 | 74 | DMEAHKKYGKVWGF | F215C_PGM | 2 | 0.8569 | 0.0206 |
| 61 | 74 | DMEAHKKYGKVWGF | F215C_PGM | 5 | 0.9976 | 0.0415 |
| 61 | 74 | DMEAHKKYGKVWGF | F215C_PGM | 30 | 1.4167 | 0.0688 |
| 61 | 74 | DMEAHKKYGKVWGF | F215C_PGM | 60 | 1.6362 | 0.0518 |
| 61 | 74 | DMEAHKKYGKVWGF | F215C_PGM | 240 | 1.907 | 0.0379 |
| 61 | 82 | DMEAHKKYGKVWGFYDGQQPVL | F215C | 0 | 0 | 0 |
| 61 | 82 | DMEAHKKYGKVWGFYDGQQPVL | F215C | 0.5 | 0.8939 | 0.0323 |
| 61 | 82 | DMEAHKKYGKVWGFYDGQQPVL | F215C | 1 | 1.0622 | 0.0581 |
| 61 | 82 | DMEAHKKYGKVWGFYDGQQPVL | F215C | 2 | 1.2434 | 0.2104 |
| 61 | 82 | DMEAHKKYGKVWGFYDGQQPVL | F215C | 5 | 1.5569 | 0.2601 |
| 61 | 82 | DMEAHKKYGKVWGFYDGQQPVL | F215C | 30 | 1.966 | 0.1116 |
| 61 | 82 | DMEAHKKYGKVWGFYDGQQPVL | F215C | 60 | 1.8611 | 0.1085 |
| 61 | 82 | DMEAHKKYGKVWGFYDGQQPVL | F215C | 240 | 1.8735 | 0.0647 |
| 61 | 82 | DMEAHKKYGKVWGFYDGQQPVL | F215C_PGM | 0 | 0 | 0 |
| 61 | 82 | DMEAHKKYGKVWGFYDGQQPVL | F215C_PGM | 0.5 | 1.0219 | 0.1251 |
| 61 | 82 | DMEAHKKYGKVWGFYDGQQPVL | F215C_PGM | 1 | 1.1483 | 0.187 |
| 61 | 82 | DMEAHKKYGKVWGFYDGQQPVL | F215C_PGM | 2 | 1.2733 | 0.0618 |
| 61 | 82 | DMEAHKKYGKVWGFYDGQQPVL | F215C_PGM | 5 | 1.5487 | 0.0742 |
| 61 | 82 | DMEAHKKYGKVWGFYDGQQPVL | F215C_PGM | 30 | 2.168 | 0.0809 |
| 61 | 82 | DMEAHKKYGKVWGFYDGQQPVL | F215C_PGM | 60 | 2.4131 | 0.1356 |
| 61 | 82 | DMEAHKKYGKVWGFYDGQQPVL | F215C_PGM | 240 | 2.9889 | 0.1535 |
| 74 | 82 | FYDGQQPVL | F215C | 0 | 0 | 0 |
| 74 | 82 | FYDGQQPVL | F215C | 0.5 | 0.6613 | 0.0384 |
| 74 | 82 | FYDGQQPVL | F215C | 1 | 0.7131 | 0.055 |
| 74 | 82 | FYDGQQPVL | F215C | 2 | 0.8876 | 0.0293 |
| 74 | 82 | FYDGQQPVL | F215C | 5 | 1.007 | 0.0377 |
| 74 | 82 | FYDGQQPVL | F215C | 30 | 1.321 | 0.0786 |
| 74 | 82 | FYDGQQPVL | F215C | 60 | 1.412 | 0.1174 |
| 74 | 82 | FYDGQQPVL | F215C | 240 | 1.5752 | 0.1347 |
| 74 | 82 | FYDGQQPVL | F215C_PGM | 0 | 0 | 0 |
| 74 | 82 | FYDGQQPVL | F215C_PGM | 0.5 | 0.6423 | 0.0245 |
| 74 | 82 | FYDGQQPVL | F215C_PGM | 1 | 0.7499 | 0.0496 |
| 74 | 82 | FYDGQQPVL | F215C_PGM | 2 | 0.8464 | 0.0244 |
| 74 | 82 | FYDGQQPVL | F215C_PGM | 5 | 0.9807 | 0.0386 |

| | | | | | | |
|---|---|---|---|---|---|---|
| 74 | 82 | FYDGQQPVL | F215C_PGM | 30 | 1.3167 | 0.0392 |
| 74 | 82 | FYDGQQPVL | F215C_PGM | 60 | 1.2299 | 0.2104 |
| 74 | 82 | FYDGQQPVL | F215C_PGM | 240 | 1.4479 | 0.0769 |
| 83 | 89 | AITDPDM | F215C | 0 | 0 | 0 |
| 83 | 89 | AITDPDM | F215C | 0.5 | 0.2783 | 0.0299 |
| 83 | 89 | AITDPDM | F215C | 1 | 0.2676 | 0.0318 |
| 83 | 89 | AITDPDM | F215C | 2 | 0.3925 | 0.0429 |
| 83 | 89 | AITDPDM | F215C | 5 | 0.4871 | 0.0308 |
| 83 | 89 | AITDPDM | F215C | 30 | 0.6845 | 0.0355 |
| 83 | 89 | AITDPDM | F215C | 60 | 0.6037 | 0.0758 |
| 83 | 89 | AITDPDM | F215C | 240 | 0.8368 | 0.0922 |
| 83 | 89 | AITDPDM | F215C_PGM | 0 | 0 | 0 |
| 83 | 89 | AITDPDM | F215C_PGM | 0.5 | 0.3064 | 0.0362 |
| 83 | 89 | AITDPDM | F215C_PGM | 1 | 0.3623 | 0.0753 |
| 83 | 89 | AITDPDM | F215C_PGM | 2 | 0.46 | 0.017 |
| 83 | 89 | AITDPDM | F215C_PGM | 5 | 0.57 | 0.0525 |
| 83 | 89 | AITDPDM | F215C_PGM | 30 | 0.7282 | 0.0101 |
| 83 | 89 | AITDPDM | F215C_PGM | 60 | 0.693 | 0.0402 |
| 83 | 89 | AITDPDM | F215C_PGM | 240 | 0.7068 | 0.0436 |
| 83 | 94 | AITDPDMIKTVL | F215C | 0 | 0 | 0 |
| 83 | 94 | AITDPDMIKTVL | F215C | 0.5 | 0.6603 | 0.1229 |
| 83 | 94 | AITDPDMIKTVL | F215C | 1 | 0.7465 | 0.0595 |
| 83 | 94 | AITDPDMIKTVL | F215C | 2 | 0.9699 | 0.1402 |
| 83 | 94 | AITDPDMIKTVL | F215C | 5 | 1.2161 | 0.1767 |
| 83 | 94 | AITDPDMIKTVL | F215C | 30 | 1.6842 | 0.0633 |
| 83 | 94 | AITDPDMIKTVL | F215C | 60 | 1.8902 | 0.1731 |
| 83 | 94 | AITDPDMIKTVL | F215C | 240 | 2.1109 | 0.0509 |
| 83 | 94 | AITDPDMIKTVL | F215C_PGM | 0 | 0 | 0 |
| 83 | 94 | AITDPDMIKTVL | F215C_PGM | 0.5 | 0.6601 | 0.0834 |
| 83 | 94 | AITDPDMIKTVL | F215C_PGM | 1 | 0.7354 | 0.0793 |
| 83 | 94 | AITDPDMIKTVL | F215C_PGM | 2 | 0.8747 | 0.064 |
| 83 | 94 | AITDPDMIKTVL | F215C_PGM | 5 | 1.2036 | 0.1189 |
| 83 | 94 | AITDPDMIKTVL | F215C_PGM | 30 | 1.7011 | 0.099 |
| 83 | 94 | AITDPDMIKTVL | F215C_PGM | 60 | 1.8477 | 0.1185 |
| 83 | 94 | AITDPDMIKTVL | F215C_PGM | 240 | 1.997 | 0.1346 |
| 93 | 102 | VLVKESYSVF | F215C | 0 | 0 | 0 |
| 93 | 102 | VLVKESYSVF | F215C | 0.5 | 2.4192 | 0.0856 |
| 93 | 102 | VLVKESYSVF | F215C | 1 | 2.5998 | 0.0604 |
| 93 | 102 | VLVKESYSVF | F215C | 2 | 2.7889 | 0.0811 |
| 93 | 102 | VLVKESYSVF | F215C | 5 | 3.0148 | 0.0377 |
| 93 | 102 | VLVKESYSVF | F215C | 30 | 3.0881 | 0.0739 |
| 93 | 102 | VLVKESYSVF | F215C | 60 | 3.0501 | 0.0249 |
| 93 | 102 | VLVKESYSVF | F215C | 240 | 3.0874 | 0.1253 |
| 93 | 102 | VLVKESYSVF | F215C_PGM | 0 | 0 | 0 |
| 93 | 102 | VLVKESYSVF | F215C_PGM | 0.5 | 2.4005 | 0.0498 |

| | | | | | | |
|---|---|---|---|---|---|---|
| 93 | 102 | VLVKESYSVF | F215C_PGM | 1 | 2.6037 | 0.1452 |
| 93 | 102 | VLVKESYSVF | F215C_PGM | 2 | 2.696 | 0.0576 |
| 93 | 102 | VLVKESYSVF | F215C_PGM | 5 | 2.9152 | 0.0549 |
| 93 | 102 | VLVKESYSVF | F215C_PGM | 30 | 3.0151 | 0.1072 |
| 93 | 102 | VLVKESYSVF | F215C_PGM | 60 | 3.0193 | 0.0294 |
| 93 | 102 | VLVKESYSVF | F215C_PGM | 240 | 2.9771 | 0.1135 |
| 94 | 106 | LVKESYSVFTNRR | F215C | 0 | 0 | 0 |
| 94 | 106 | LVKESYSVFTNRR | F215C | 0.5 | 1.5325 | 0.0584 |
| 94 | 106 | LVKESYSVFTNRR | F215C | 1 | 1.7646 | 0.085 |
| 94 | 106 | LVKESYSVFTNRR | F215C | 2 | 2.0089 | 0.0122 |
| 94 | 106 | LVKESYSVFTNRR | F215C | 5 | 2.192 | 0.0314 |
| 94 | 106 | LVKESYSVFTNRR | F215C | 30 | 2.5804 | 0.0793 |
| 94 | 106 | LVKESYSVFTNRR | F215C | 60 | 2.5333 | 0.0479 |
| 94 | 106 | LVKESYSVFTNRR | F215C | 240 | 2.8247 | 0.0371 |
| 94 | 106 | LVKESYSVFTNRR | F215C_PGM | 0 | 0 | 0 |
| 94 | 106 | LVKESYSVFTNRR | F215C_PGM | 0.5 | 1.5847 | 0.0557 |
| 94 | 106 | LVKESYSVFTNRR | F215C_PGM | 1 | 1.811 | 0.0598 |
| 94 | 106 | LVKESYSVFTNRR | F215C_PGM | 2 | 1.96 | 0.0613 |
| 94 | 106 | LVKESYSVFTNRR | F215C_PGM | 5 | 2.2081 | 0.0721 |
| 94 | 106 | LVKESYSVFTNRR | F215C_PGM | 30 | 2.5649 | 0.035 |
| 94 | 106 | LVKESYSVFTNRR | F215C_PGM | 60 | 2.6245 | 0.0567 |
| 94 | 106 | LVKESYSVFTNRR | F215C_PGM | 240 | 2.7626 | 0.1331 |
| 95 | 102 | VKESYSVF | F215C | 0 | 0 | 0 |
| 95 | 102 | VKESYSVF | F215C | 0.5 | 1.3238 | 0.0897 |
| 95 | 102 | VKESYSVF | F215C | 1 | 1.3319 | 0.0758 |
| 95 | 102 | VKESYSVF | F215C | 2 | 1.4916 | 0.0748 |
| 95 | 102 | VKESYSVF | F215C | 5 | 1.6252 | 0.0439 |
| 95 | 102 | VKESYSVF | F215C | 30 | 1.6529 | 0.0715 |
| 95 | 102 | VKESYSVF | F215C | 60 | 1.6135 | 0.1155 |
| 95 | 102 | VKESYSVF | F215C | 240 | 1.798 | 0.1487 |
| 95 | 102 | VKESYSVF | F215C_PGM | 0 | 0 | 0 |
| 95 | 102 | VKESYSVF | F215C_PGM | 0.5 | 1.4412 | 0.033 |
| 95 | 102 | VKESYSVF | F215C_PGM | 1 | 1.5627 | 0.1524 |
| 95 | 102 | VKESYSVF | F215C_PGM | 2 | 1.6189 | 0.0252 |
| 95 | 102 | VKESYSVF | F215C_PGM | 5 | 1.7356 | 0.0465 |
| 95 | 102 | VKESYSVF | F215C_PGM | 30 | 1.898 | 0.0432 |
| 95 | 102 | VKESYSVF | F215C_PGM | 60 | 1.9584 | 0.0633 |
| 95 | 102 | VKESYSVF | F215C_PGM | 240 | 1.9844 | 0.0757 |
| 114 | 122 | MKSAISIAE | F215C | 0 | 0 | 0 |
| 114 | 122 | MKSAISIAE | F215C | 0.5 | 1.0368 | 0.064 |
| 114 | 122 | MKSAISIAE | F215C | 1 | 1.1095 | 0.0364 |
| 114 | 122 | MKSAISIAE | F215C | 2 | 1.2135 | 0.1003 |
| 114 | 122 | MKSAISIAE | F215C | 5 | 1.6712 | 0.0452 |
| 114 | 122 | MKSAISIAE | F215C | 30 | 2.4022 | 0.0041 |
| 114 | 122 | MKSAISIAE | F215C | 60 | 2.6156 | 0.0106 |

| | | | | | | |
|---|---|---|---|---|---|---|
| 114 | 122 | MKSAISIAE | F215C | 240 | 3.2663 | 0.0937 |
| 114 | 122 | MKSAISIAE | F215C_PGM | 0 | 0 | 0 |
| 114 | 122 | MKSAISIAE | F215C_PGM | 0.5 | 0.8969 | 0.0118 |
| 114 | 122 | MKSAISIAE | F215C_PGM | 1 | 0.9703 | 0.061 |
| 114 | 122 | MKSAISIAE | F215C_PGM | 2 | 1.1172 | 0.064 |
| 114 | 122 | MKSAISIAE | F215C_PGM | 5 | 1.3659 | 0.0275 |
| 114 | 122 | MKSAISIAE | F215C_PGM | 30 | 2.0392 | 0.0447 |
| 114 | 122 | MKSAISIAE | F215C_PGM | 60 | 2.3142 | 0.0561 |
| 114 | 122 | MKSAISIAE | F215C_PGM | 240 | 3.0504 | 0.0465 |
| 120 | 125 | IAEDEE | F215C | 0 | 0 | 0 |
| 120 | 125 | IAEDEE | F215C | 0.5 | 0.6009 | 0.0416 |
| 120 | 125 | IAEDEE | F215C | 1 | 0.6679 | 0.035 |
| 120 | 125 | IAEDEE | F215C | 2 | 0.7635 | 0.062 |
| 120 | 125 | IAEDEE | F215C | 5 | 0.8883 | 0.0384 |
| 120 | 125 | IAEDEE | F215C | 30 | 0.9836 | 0.0563 |
| 120 | 125 | IAEDEE | F215C | 60 | 0.9282 | 0.0331 |
| 120 | 125 | IAEDEE | F215C | 240 | 0.9405 | 0.0887 |
| 120 | 125 | IAEDEE | F215C_PGM | 0 | 0 | 0 |
| 120 | 125 | IAEDEE | F215C_PGM | 0.5 | 0.6685 | 0.0302 |
| 120 | 125 | IAEDEE | F215C_PGM | 1 | 0.7612 | 0.073 |
| 120 | 125 | IAEDEE | F215C_PGM | 2 | 0.8481 | 0.037 |
| 120 | 125 | IAEDEE | F215C_PGM | 5 | 0.9792 | 0.0475 |
| 120 | 125 | IAEDEE | F215C_PGM | 30 | 1.1546 | 0.0416 |
| 120 | 125 | IAEDEE | F215C_PGM | 60 | 1.1704 | 0.046 |
| 120 | 125 | IAEDEE | F215C_PGM | 240 | 1.098 | 0.054 |
| 123 | 133 | DEEWKRLRSLL | F215C | 0 | 0 | 0 |
| 123 | 133 | DEEWKRLRSLL | F215C | 0.5 | 1.3203 | 0.0995 |
| 123 | 133 | DEEWKRLRSLL | F215C | 1 | 1.3738 | 0.171 |
| 123 | 133 | DEEWKRLRSLL | F215C | 2 | 1.5434 | 0.0791 |
| 123 | 133 | DEEWKRLRSLL | F215C | 5 | 1.6374 | 0.0927 |
| 123 | 133 | DEEWKRLRSLL | F215C | 30 | 2.1893 | 0.1822 |
| 123 | 133 | DEEWKRLRSLL | F215C | 60 | 2.3379 | 0.1621 |
| 123 | 133 | DEEWKRLRSLL | F215C | 240 | 2.6821 | 0.0884 |
| 123 | 133 | DEEWKRLRSLL | F215C_PGM | 0 | 0 | 0 |
| 123 | 133 | DEEWKRLRSLL | F215C_PGM | 0.5 | 1.4363 | 0.1006 |
| 123 | 133 | DEEWKRLRSLL | F215C_PGM | 1 | 1.6922 | 0.1484 |
| 123 | 133 | DEEWKRLRSLL | F215C_PGM | 2 | 1.8865 | 0.038 |
| 123 | 133 | DEEWKRLRSLL | F215C_PGM | 5 | 2.1267 | 0.0883 |
| 123 | 133 | DEEWKRLRSLL | F215C_PGM | 30 | 2.3763 | 0.0747 |
| 123 | 133 | DEEWKRLRSLL | F215C_PGM | 60 | 2.3227 | 0.1378 |
| 123 | 133 | DEEWKRLRSLL | F215C_PGM | 240 | 2.6897 | 0.1169 |
| 123 | 137 | DEEWKRLRSLLSPTF | F215C | 0 | 0 | 0 |
| 123 | 137 | DEEWKRLRSLLSPTF | F215C | 0.5 | 1.4572 | 0.1421 |
| 123 | 137 | DEEWKRLRSLLSPTF | F215C | 1 | 1.6596 | 0.1331 |
| 123 | 137 | DEEWKRLRSLLSPTF | F215C | 2 | 2.0422 | 0.3269 |

| | | | | | | |
|---|---|---|---|---|---|---|
| 123 | 137 | DEEWKRLRSLLSPTF | F215C | 5 | 2.1924 | 0.125 |
| 123 | 137 | DEEWKRLRSLLSPTF | F215C | 30 | 3.25 | 0.0392 |
| 123 | 137 | DEEWKRLRSLLSPTF | F215C | 60 | 3.4773 | 0.1555 |
| 123 | 137 | DEEWKRLRSLLSPTF | F215C | 240 | 4.1543 | 0.0549 |
| 123 | 137 | DEEWKRLRSLLSPTF | F215C_PGM | 0 | 0 | 0 |
| 123 | 137 | DEEWKRLRSLLSPTF | F215C_PGM | 0.5 | 1.2972 | 0.1361 |
| 123 | 137 | DEEWKRLRSLLSPTF | F215C_PGM | 1 | 1.5324 | 0.1442 |
| 123 | 137 | DEEWKRLRSLLSPTF | F215C_PGM | 2 | 1.9127 | 0.0643 |
| 123 | 137 | DEEWKRLRSLLSPTF | F215C_PGM | 5 | 2.4595 | 0.0564 |
| 123 | 137 | DEEWKRLRSLLSPTF | F215C_PGM | 30 | 3.2237 | 0.0556 |
| 123 | 137 | DEEWKRLRSLLSPTF | F215C_PGM | 60 | 3.4684 | 0.0411 |
| 123 | 137 | DEEWKRLRSLLSPTF | F215C_PGM | 240 | 3.8844 | 0.1278 |
| 126 | 137 | WKRLRSLLSPTF | F215C | 0 | 0 | 0 |
| 126 | 137 | WKRLRSLLSPTF | F215C | 0.5 | 0.9358 | 0.0451 |
| 126 | 137 | WKRLRSLLSPTF | F215C | 1 | 1.0309 | 0.0395 |
| 126 | 137 | WKRLRSLLSPTF | F215C | 2 | 1.2578 | 0.1197 |
| 126 | 137 | WKRLRSLLSPTF | F215C | 5 | 1.5382 | 0.0562 |
| 126 | 137 | WKRLRSLLSPTF | F215C | 30 | 2.1945 | 0.0388 |
| 126 | 137 | WKRLRSLLSPTF | F215C | 60 | 2.2585 | 0.0639 |
| 126 | 137 | WKRLRSLLSPTF | F215C | 240 | 2.622 | 0.122 |
| 126 | 137 | WKRLRSLLSPTF | F215C_PGM | 0 | 0 | 0 |
| 126 | 137 | WKRLRSLLSPTF | F215C_PGM | 0.5 | 1.0206 | 0.0543 |
| 126 | 137 | WKRLRSLLSPTF | F215C_PGM | 1 | 1.2346 | 0.1033 |
| 126 | 137 | WKRLRSLLSPTF | F215C_PGM | 2 | 1.5195 | 0.0287 |
| 126 | 137 | WKRLRSLLSPTF | F215C_PGM | 5 | 1.9092 | 0.0535 |
| 126 | 137 | WKRLRSLLSPTF | F215C_PGM | 30 | 2.5146 | 0.0615 |
| 126 | 137 | WKRLRSLLSPTF | F215C_PGM | 60 | 2.5458 | 0.0296 |
| 126 | 137 | WKRLRSLLSPTF | F215C_PGM | 240 | 2.7144 | 0.0939 |
| 135 | 151 | PTFTSGKLKEMVPIIAQ | F215C | 0 | 0 | 0 |
| 135 | 151 | SPTFTSGKLKEMVPIIAQ | F215C | 0.5 | 1.8875 | 0.0751 |
| 135 | 151 | SPTFTSGKLKEMVPIIAQ | F215C | 1 | 2.3246 | 0.1687 |
| 135 | 151 | SPTFTSGKLKEMVPIIAQ | F215C | 2 | 2.8403 | 0.0291 |
| 135 | 151 | SPTFTSGKLKEMVPIIAQ | F215C | 5 | 3.3919 | 0.0572 |
| 135 | 151 | SPTFTSGKLKEMVPIIAQ | F215C | 30 | 4.2997 | 0.0858 |
| 135 | 151 | SPTFTSGKLKEMVPIIAQ | F215C | 60 | 4.664 | 0.0628 |
| 135 | 151 | SPTFTSGKLKEMVPIIAQ | F215C | 240 | 5.4049 | 0.1094 |
| 135 | 151 | SPTFTSGKLKEMVPIIAQ | F215C_PGM | 0 | 0 | 0 |
| 135 | 151 | SPTFTSGKLKEMVPIIAQ | F215C_PGM | 0.5 | 1.9933 | 0.1193 |
| 135 | 151 | SPTFTSGKLKEMVPIIAQ | F215C_PGM | 1 | 2.368 | 0.0644 |
| 135 | 151 | SPTFTSGKLKEMVPIIAQ | F215C_PGM | 2 | 2.9254 | 0.0774 |
| 135 | 151 | SPTFTSGKLKEMVPIIAQ | F215C_PGM | 5 | 3.5987 | 0.0914 |
| 135 | 151 | SPTFTSGKLKEMVPIIAQ | F215C_PGM | 30 | 4.6276 | 0.0842 |
| 135 | 151 | SPTFTSGKLKEMVPIIAQ | F215C_PGM | 60 | 4.9726 | 0.1619 |
| 135 | 151 | SPTFTSGKLKEMVPIIAQ | F215C_PGM | 240 | 5.5012 | 0.1519 |
| 138 | 151 | TSGKLKEMVPIIAQ | F215C | 0 | 0 | 0 |

| | | | | | | |
|---|---|---|---|---|---|---|
| 138 | 151 | TSGKLKEMVPIIAQ | F215C | 0.5 | 1.6542 | 0.0686 |
| 138 | 151 | TSGKLKEMVPIIAQ | F215C | 1 | 1.8532 | 0.1724 |
| 138 | 151 | TSGKLKEMVPIIAQ | F215C | 2 | 2.4115 | 0.0777 |
| 138 | 151 | TSGKLKEMVPIIAQ | F215C | 5 | 2.9723 | 0.1473 |
| 138 | 151 | TSGKLKEMVPIIAQ | F215C | 30 | 3.8878 | 0.0481 |
| 138 | 151 | TSGKLKEMVPIIAQ | F215C | 60 | 4.1626 | 0.0243 |
| 138 | 151 | TSGKLKEMVPIIAQ | F215C | 240 | 4.8744 | 0.1026 |
| 138 | 151 | TSGKLKEMVPIIAQ | F215C_PGM | 0 | 0 | 0 |
| 138 | 151 | TSGKLKEMVPIIAQ | F215C_PGM | 0.5 | 1.6776 | 0.0672 |
| 138 | 151 | TSGKLKEMVPIIAQ | F215C_PGM | 1 | 2.0908 | 0.1246 |
| 138 | 151 | TSGKLKEMVPIIAQ | F215C_PGM | 2 | 2.4812 | 0.0699 |
| 138 | 151 | TSGKLKEMVPIIAQ | F215C_PGM | 5 | 3.0289 | 0.0894 |
| 138 | 151 | TSGKLKEMVPIIAQ | F215C_PGM | 30 | 3.9348 | 0.0606 |
| 138 | 151 | TSGKLKEMVPIIAQ | F215C_PGM | 60 | 4.2526 | 0.0824 |
| 138 | 151 | TSGKLKEMVPIIAQ | F215C_PGM | 240 | 4.8125 | 0.1151 |
| 143 | 151 | KEMVPIIAQ | F215C | 0 | 0 | 0 |
| 143 | 151 | KEMVPIIAQ | F215C | 0.5 | 1.2975 | 0.0364 |
| 143 | 151 | KEMVPIIAQ | F215C | 1 | 1.3958 | 0.1245 |
| 143 | 151 | KEMVPIIAQ | F215C | 2 | 1.5245 | 0.0675 |
| 143 | 151 | KEMVPIIAQ | F215C | 5 | 1.6688 | 0.0757 |
| 143 | 151 | KEMVPIIAQ | F215C | 30 | 2.2071 | 0.0937 |
| 143 | 151 | KEMVPIIAQ | F215C | 60 | 2.5725 | 0.0447 |
| 143 | 151 | KEMVPIIAQ | F215C | 240 | 3.0641 | 0.0766 |
| 143 | 151 | KEMVPIIAQ | F215C_PGM | 0 | 0 | 0 |
| 143 | 151 | KEMVPIIAQ | F215C_PGM | 0.5 | 1.4574 | 0.1141 |
| 143 | 151 | KEMVPIIAQ | F215C_PGM | 1 | 1.5517 | 0.1279 |
| 143 | 151 | KEMVPIIAQ | F215C_PGM | 2 | 1.787 | 0.0662 |
| 143 | 151 | KEMVPIIAQ | F215C_PGM | 5 | 1.9206 | 0.0491 |
| 143 | 151 | KEMVPIIAQ | F215C_PGM | 30 | 2.5296 | 0.0632 |
| 143 | 151 | KEMVPIIAQ | F215C_PGM | 60 | 2.7683 | 0.0508 |
| 143 | 151 | KEMVPIIAQ | F215C_PGM | 240 | 3.1687 | 0.0647 |
| 157 | 176 | VRNLRREAETGKPVTLKDVF | F215C | 0 | 0 | 0 |
| 157 | 176 | VRNLRREAETGKPVTLKDVF | F215C | 0.5 | 3.6464 | 0.1238 |
| 157 | 176 | VRNLRREAETGKPVTLKDVF | F215C | 1 | 4.4412 | 0.1167 |
| 157 | 176 | VRNLRREAETGKPVTLKDVF | F215C | 2 | 5.1345 | 0.1865 |
| 157 | 176 | VRNLRREAETGKPVTLKDVF | F215C | 5 | 5.9017 | 0.1267 |
| 157 | 176 | VRNLRREAETGKPVTLKDVF | F215C | 30 | 6.3936 | 0.2103 |
| 157 | 176 | VRNLRREAETGKPVTLKDVF | F215C | 60 | 6.2467 | 0.1809 |
| 157 | 176 | VRNLRREAETGKPVTLKDVF | F215C | 240 | 6.7343 | 0.4027 |
| 157 | 176 | VRNLRREAETGKPVTLKDVF | F215C_PGM | 0 | 0 | 0 |
| 157 | 176 | VRNLRREAETGKPVTLKDVF | F215C_PGM | 0.5 | 3.9488 | 0.1559 |
| 157 | 176 | VRNLRREAETGKPVTLKDVF | F215C_PGM | 1 | 4.598 | 0.3368 |
| 157 | 176 | VRNLRREAETGKPVTLKDVF | F215C_PGM | 2 | 5.281 | 0.0863 |
| 157 | 176 | VRNLRREAETGKPVTLKDVF | F215C_PGM | 5 | 6.0645 | 0.0496 |
| 157 | 176 | VRNLRREAETGKPVTLKDVF | F215C_PGM | 30 | 6.8333 | 0.1956 |

| | | | | | | |
|---|---|---|---|---|---|---|
| 157 | 176 | VRNLRREAETGKPVTLKDVF | F215C_PGM | 60 | 6.8773 | 0.131 |
| 157 | 176 | VRNLRREAETGKPVTLKDVF | F215C_PGM | 240 | 6.9412 | 0.2162 |
| 173 | 178 | KDVFGA | F215C | 0 | 0 | 0 |
| 173 | 178 | KDVFGA | F215C | 0.5 | 0.6197 | 0.0324 |
| 173 | 178 | KDVFGA | F215C | 1 | 0.7226 | 0.0399 |
| 173 | 178 | KDVFGA | F215C | 2 | 0.8848 | 0.0332 |
| 173 | 178 | KDVFGA | F215C | 5 | 1.2063 | 0.0431 |
| 173 | 178 | KDVFGA | F215C | 30 | 1.657 | 0.0209 |
| 173 | 178 | KDVFGA | F215C | 60 | 1.7736 | 0.0442 |
| 173 | 178 | KDVFGA | F215C | 240 | 1.8157 | 0.0351 |
| 173 | 178 | KDVFGA | F215C_PGM | 0 | 0 | 0 |
| 173 | 178 | KDVFGA | F215C_PGM | 0.5 | 0.4998 | 0.0312 |
| 173 | 178 | KDVFGA | F215C_PGM | 1 | 0.6509 | 0.0211 |
| 173 | 178 | KDVFGA | F215C_PGM | 2 | 0.7832 | 0.0263 |
| 173 | 178 | KDVFGA | F215C_PGM | 5 | 0.9547 | 0.0175 |
| 173 | 178 | KDVFGA | F215C_PGM | 30 | 1.2958 | 0.0231 |
| 173 | 178 | KDVFGA | F215C_PGM | 60 | 1.4959 | 0.0475 |
| 173 | 178 | KDVFGA | F215C_PGM | 240 | 1.6675 | 0.0568 |
| 173 | 181 | KDVFGAYSM | F215C | 0 | 0 | 0 |
| 173 | 181 | KDVFGAYSM | F215C | 0.5 | 0.7624 | 0.0324 |
| 173 | 181 | KDVFGAYSM | F215C | 1 | 0.8338 | 0.0671 |
| 173 | 181 | KDVFGAYSM | F215C | 2 | 1.0885 | 0.1022 |
| 173 | 181 | KDVFGAYSM | F215C | 5 | 1.3112 | 0.1122 |
| 173 | 181 | KDVFGAYSM | F215C | 30 | 1.9611 | 0.0572 |
| 173 | 181 | KDVFGAYSM | F215C | 60 | 1.9319 | 0.2413 |
| 173 | 181 | KDVFGAYSM | F215C | 240 | 2.3663 | 0.1397 |
| 173 | 181 | KDVFGAYSM | F215C_PGM | 0 | 0 | 0 |
| 173 | 181 | KDVFGAYSM | F215C_PGM | 0.5 | 0.6984 | 0.0872 |
| 173 | 181 | KDVFGAYSM | F215C_PGM | 1 | 0.8386 | 0.0976 |
| 173 | 181 | KDVFGAYSM | F215C_PGM | 2 | 0.9522 | 0.0667 |
| 173 | 181 | KDVFGAYSM | F215C_PGM | 5 | 1.213 | 0.0331 |
| 173 | 181 | KDVFGAYSM | F215C_PGM | 30 | 1.7225 | 0.0835 |
| 173 | 181 | KDVFGAYSM | F215C_PGM | 60 | 1.9711 | 0.0698 |
| 173 | 181 | KDVFGAYSM | F215C_PGM | 240 | 2.1691 | 0.1015 |
| 177 | 182 | GAYSMD | F215C | 0 | 0 | 0 |
| 177 | 182 | GAYSMD | F215C | 0.5 | 0.2897 | 0.0246 |
| 177 | 182 | GAYSMD | F215C | 1 | 0.3576 | 0.0422 |
| 177 | 182 | GAYSMD | F215C | 2 | 0.3782 | 0.038 |
| 177 | 182 | GAYSMD | F215C | 5 | 0.4238 | 0.0448 |
| 177 | 182 | GAYSMD | F215C | 30 | 0.5816 | 0.0516 |
| 177 | 182 | GAYSMD | F215C | 60 | 0.6831 | 0.0272 |
| 177 | 182 | GAYSMD | F215C | 240 | 0.9595 | 0.0269 |
| 177 | 182 | GAYSMD | F215C_PGM | 0 | 0 | 0 |
| 177 | 182 | GAYSMD | F215C_PGM | 0.5 | 0.2911 | 0.0289 |
| 177 | 182 | GAYSMD | F215C_PGM | 1 | 0.3079 | 0.027 |

| | | | | | | |
|---|---|---|---|---|---|---|
| 177 | 182 | GAYSMD | F215C_PGM | 2 | 0.3578 | 0.0406 |
| 177 | 182 | GAYSMD | F215C_PGM | 5 | 0.3727 | 0.0242 |
| 177 | 182 | GAYSMD | F215C_PGM | 30 | 0.4589 | 0.0353 |
| 177 | 182 | GAYSMD | F215C_PGM | 60 | 0.5368 | 0.0245 |
| 177 | 182 | GAYSMD | F215C_PGM | 240 | 0.6448 | 0.0334 |
| 182 | 189 | DVITSTSF | F215C | 0 | 0 | 0 |
| 182 | 189 | DVITSTSF | F215C | 0.5 | 0.6854 | 0.0415 |
| 182 | 189 | DVITSTSF | F215C | 1 | 0.687 | 0.0379 |
| 182 | 189 | DVITSTSF | F215C | 2 | 0.8031 | 0.0276 |
| 182 | 189 | DVITSTSF | F215C | 5 | 0.8733 | 0.0295 |
| 182 | 189 | DVITSTSF | F215C | 30 | 1.1801 | 0.0903 |
| 182 | 189 | DVITSTSF | F215C | 60 | 1.2472 | 0.0762 |
| 182 | 189 | DVITSTSF | F215C | 240 | 1.6416 | 0.0699 |
| 182 | 189 | DVITSTSF | F215C_PGM | 0 | 0 | 0 |
| 182 | 189 | DVITSTSF | F215C_PGM | 0.5 | 0.6981 | 0.061 |
| 182 | 189 | DVITSTSF | F215C_PGM | 1 | 0.7314 | 0.0976 |
| 182 | 189 | DVITSTSF | F215C_PGM | 2 | 0.7856 | 0.0146 |
| 182 | 189 | DVITSTSF | F215C_PGM | 5 | 0.8558 | 0.0315 |
| 182 | 189 | DVITSTSF | F215C_PGM | 30 | 1.0856 | 0.0253 |
| 182 | 189 | DVITSTSF | F215C_PGM | 60 | 1.2454 | 0.0554 |
| 182 | 189 | DVITSTSF | F215C_PGM | 240 | 1.5416 | 0.0749 |
| 182 | 192 | DVITSTSFGVN | F215C | 0 | 0 | 0 |
| 182 | 192 | DVITSTSFGVN | F215C | 0.5 | 0.3758 | 0.0508 |
| 182 | 192 | DVITSTSFGVN | F215C | 1 | 0.4154 | 0.0198 |
| 182 | 192 | DVITSTSFGVN | F215C | 2 | 0.5151 | 0.0373 |
| 182 | 192 | DVITSTSFGVN | F215C | 5 | 0.5004 | 0.0323 |
| 182 | 192 | DVITSTSFGVN | F215C | 30 | 0.5227 | 0.0978 |
| 182 | 192 | DVITSTSFGVN | F215C | 60 | 0.4784 | 0.103 |
| 182 | 192 | DVITSTSFGVN | F215C | 240 | 0.4759 | 0.1321 |
| 182 | 192 | DVITSTSFGVN | F215C_PGM | 0 | 0 | 0 |
| 182 | 192 | DVITSTSFGVN | F215C_PGM | 0.5 | 0.3539 | 0.0405 |
| 182 | 192 | DVITSTSFGVN | F215C_PGM | 1 | 0.4243 | 0.102 |
| 182 | 192 | DVITSTSFGVN | F215C_PGM | 2 | 0.4232 | 0.0962 |
| 182 | 192 | DVITSTSFGVN | F215C_PGM | 5 | 0.5075 | 0.0296 |
| 182 | 192 | DVITSTSFGVN | F215C_PGM | 30 | 0.5172 | 0.0478 |
| 182 | 192 | DVITSTSFGVN | F215C_PGM | 60 | 0.5632 | 0.0987 |
| 182 | 192 | DVITSTSFGVN | F215C_PGM | 240 | 0.4767 | 0.0918 |
| 183 | 189 | VITSTSF | F215C | 0 | 0 | 0 |
| 183 | 189 | VITSTSF | F215C | 0.5 | 0.491 | 0.0267 |
| 183 | 189 | VITSTSF | F215C | 1 | 0.4589 | 0.017 |
| 183 | 189 | VITSTSF | F215C | 2 | 0.5558 | 0.0245 |
| 183 | 189 | VITSTSF | F215C | 5 | 0.6125 | 0.023 |
| 183 | 189 | VITSTSF | F215C | 30 | 0.8358 | 0.0418 |
| 183 | 189 | VITSTSF | F215C | 60 | 0.8242 | 0.0951 |
| 183 | 189 | VITSTSF | F215C | 240 | 1.0504 | 0.1322 |

| | | | | | | |
|---|---|---|---|---|---|---|
| 183 | 189 | VITSTSF | F215C_PGM | 0 | 0 | 0 |
| 183 | 189 | VITSTSF | F215C_PGM | 0.5 | 0.5568 | 0.0452 |
| 183 | 189 | VITSTSF | F215C_PGM | 1 | 0.6148 | 0.095 |
| 183 | 189 | VITSTSF | F215C_PGM | 2 | 0.6434 | 0.0157 |
| 183 | 189 | VITSTSF | F215C_PGM | 5 | 0.6742 | 0.0212 |
| 183 | 189 | VITSTSF | F215C_PGM | 30 | 0.8753 | 0.0122 |
| 183 | 189 | VITSTSF | F215C_PGM | 60 | 0.9474 | 0.0526 |
| 183 | 189 | VITSTSF | F215C_PGM | 240 | 1.1496 | 0.0696 |
| 189 | 212 | FGVNIDSLNNPQDPFVENTKKLLR | F215C | 0 | 0 | 0 |
| 189 | 212 | FGVNIDSLNNPQDPFVENTKKLLR | F215C | 0.5 | 4.0392 | 0.2501 |
| 189 | 212 | FGVNIDSLNNPQDPFVENTKKLLR | F215C | 1 | 4.7129 | 0.2115 |
| 189 | 212 | FGVNIDSLNNPQDPFVENTKKLLR | F215C | 2 | 5.3938 | 0.0441 |
| 189 | 212 | FGVNIDSLNNPQDPFVENTKKLLR | F215C | 5 | 6.4666 | 0.0439 |
| 189 | 212 | FGVNIDSLNNPQDPFVENTKKLLR | F215C | 30 | 7.5604 | 0.0951 |
| 189 | 212 | FGVNIDSLNNPQDPFVENTKKLLR | F215C | 60 | 7.8786 | 0.0744 |
| 189 | 212 | FGVNIDSLNNPQDPFVENTKKLLR | F215C | 240 | 8.3508 | 0.289 |
| 189 | 212 | FGVNIDSLNNPQDPFVENTKKLLR | F215C_PGM | 0 | 0 | 0 |
| 189 | 212 | FGVNIDSLNNPQDPFVENTKKLLR | F215C_PGM | 0.5 | 7.1489 | 0.3115 |
| 189 | 212 | FGVNIDSLNNPQDPFVENTKKLLR | F215C_PGM | 1 | 7.6352 | 0.286 |
| 189 | 212 | FGVNIDSLNNPQDPFVENTKKLLR | F215C_PGM | 2 | 8.8493 | 0.0481 |
| 189 | 212 | FGVNIDSLNNPQDPFVENTKKLLR | F215C_PGM | 5 | 8.9849 | 0.2585 |
| 189 | 212 | FGVNIDSLNNPQDPFVENTKKLLR | F215C_PGM | 30 | 10.1062 | 0.2803 |
| 189 | 212 | FGVNIDSLNNPQDPFVENTKKLLR | F215C_PGM | 60 | 10.1549 | 0.3944 |
| 189 | 212 | FGVNIDSLNNPQDPFVENTKKLLR | F215C_PGM | 240 | 9.7909 | 0.2278 |
| 190 | 210 | GVNIDSLNNPQDPFVENTKKL | F215C | 0 | 0 | 0 |
| 190 | 210 | GVNIDSLNNPQDPFVENTKKL | F215C | 0.5 | 3.1261 | 0.2805 |
| 190 | 210 | GVNIDSLNNPQDPFVENTKKL | F215C | 1 | 3.4902 | 0.312 |
| 190 | 210 | GVNIDSLNNPQDPFVENTKKL | F215C | 2 | 4.1201 | 0.0306 |
| 190 | 210 | GVNIDSLNNPQDPFVENTKKL | F215C | 5 | 4.9711 | 0.0869 |
| 190 | 210 | GVNIDSLNNPQDPFVENTKKL | F215C | 30 | 5.8226 | 0.3483 |
| 190 | 210 | GVNIDSLNNPQDPFVENTKKL | F215C | 60 | 6.0036 | 0.3661 |
| 190 | 210 | GVNIDSLNNPQDPFVENTKKL | F215C | 240 | 6.0798 | 0.4018 |
| 190 | 210 | GVNIDSLNNPQDPFVENTKKL | F215C_PGM | 0 | 0 | 0 |
| 190 | 210 | GVNIDSLNNPQDPFVENTKKL | F215C_PGM | 0.5 | 3.9858 | 0.6841 |
| 190 | 210 | GVNIDSLNNPQDPFVENTKKL | F215C_PGM | 1 | 6.4973 | 0.0156 |
| 190 | 210 | GVNIDSLNNPQDPFVENTKKL | F215C_PGM | 2 | | |
| 190 | 210 | GVNIDSLNNPQDPFVENTKKL | F215C_PGM | 5 | 6.6711 | 0.542 |
| 190 | 210 | GVNIDSLNNPQDPFVENTKKL | F215C_PGM | 30 | 7.4826 | 0.0374 |
| 190 | 210 | GVNIDSLNNPQDPFVENTKKL | F215C_PGM | 60 | 7.7765 | 0.3662 |
| 190 | 210 | GVNIDSLNNPQDPFVENTKKL | F215C_PGM | 240 | 7.7758 | 0.1874 |
| 221 | 226 | LSITVF | F215C | 0 | 0 | 0 |
| 221 | 226 | LSITVF | F215C | 0.5 | 0.6193 | 0.0534 |
| 221 | 226 | LSITVF | F215C | 1 | 0.7105 | 0.0727 |
| 221 | 226 | LSITVF | F215C | 2 | 0.904 | 0.1388 |
| 221 | 226 | LSITVF | F215C | 5 | 1.0174 | 0.0774 |

| | | | | | | |
|---|---|---|---|---|---|---|
| 221 | 226 | LSITVF | F215C | 30 | 1.5091 | 0.0394 |
| 221 | 226 | LSITVF | F215C | 60 | 1.6117 | 0.0453 |
| 221 | 226 | LSITVF | F215C | 240 | 1.9806 | 0.0412 |
| 221 | 226 | LSITVF | F215C_PGM | 0 | 0 | 0 |
| 221 | 226 | LSITVF | F215C_PGM | 0.5 | 0.2379 | 0.038 |
| 221 | 226 | LSITVF | F215C_PGM | 1 | 0.3388 | 0.0574 |
| 221 | 226 | LSITVF | F215C_PGM | 2 | 0.4957 | 0.069 |
| 221 | 226 | LSITVF | F215C_PGM | 5 | 0.7054 | 0.0592 |
| 221 | 226 | LSITVF | F215C_PGM | 30 | 1.0552 | 0.0655 |
| 221 | 226 | LSITVF | F215C_PGM | 60 | 1.157 | 0.0415 |
| 221 | 226 | LSITVF | F215C_PGM | 240 | 1.5305 | 0.0391 |
| 230 | 235 | IPILEV | F215C | 0 | 0 | 0 |
| 230 | 235 | IPILEV | F215C | 0.5 | 0.228 | 0.052 |
| 230 | 235 | IPILEV | F215C | 1 | 0.2014 | 0.0636 |
| 230 | 235 | IPILEV | F215C | 2 | 0.3369 | 0.0566 |
| 230 | 235 | IPILEV | F215C | 5 | 0.351 | 0.0304 |
| 230 | 235 | IPILEV | F215C | 30 | 0.9022 | 0.0303 |
| 230 | 235 | IPILEV | F215C | 60 | 1.1661 | 0.037 |
| 230 | 235 | IPILEV | F215C | 240 | 1.8343 | 0.0631 |
| 230 | 235 | IPILEV | F215C_PGM | 0 | 0 | 0 |
| 230 | 235 | IPILEV | F215C_PGM | 0.5 | 0.1175 | 0.0239 |
| 230 | 235 | IPILEV | F215C_PGM | 1 | 0.159 | 0.0225 |
| 230 | 235 | IPILEV | F215C_PGM | 2 | 0.2362 | 0.0403 |
| 230 | 235 | IPILEV | F215C_PGM | 5 | 0.3399 | 0.0293 |
| 230 | 235 | IPILEV | F215C_PGM | 30 | 0.7642 | 0.0669 |
| 230 | 235 | IPILEV | F215C_PGM | 60 | 0.9262 | 0.0178 |
| 230 | 235 | IPILEV | F215C_PGM | 240 | 1.643 | 0.0345 |
| 235 | 241 | VLNISVF | F215C | 0 | 0 | 0 |
| 235 | 241 | VLNISVF | F215C | 0.5 | 1.0763 | 0.0387 |
| 235 | 241 | VLNISVF | F215C | 1 | 1.3203 | 0.0321 |
| 235 | 241 | VLNISVF | F215C | 2 | 1.6352 | 0.0705 |
| 235 | 241 | VLNISVF | F215C | 5 | 1.9709 | 0.031 |
| 235 | 241 | VLNISVF | F215C | 30 | 2.6131 | 0.0486 |
| 235 | 241 | VLNISVF | F215C | 60 | 2.8524 | 0.0416 |
| 235 | 241 | VLNISVF | F215C | 240 | 3.0456 | 0.0938 |
| 235 | 241 | VLNISVF | F215C_PGM | 0 | 0 | 0 |
| 235 | 241 | VLNISVF | F215C_PGM | 0.5 | 0.8779 | 0.0606 |
| 235 | 241 | VLNISVF | F215C_PGM | 1 | 1.0807 | 0.0327 |
| 235 | 241 | VLNISVF | F215C_PGM | 2 | 1.3618 | 0.0475 |
| 235 | 241 | VLNISVF | F215C_PGM | 5 | 1.8122 | 0.0417 |
| 235 | 241 | VLNISVF | F215C_PGM | 30 | 2.6007 | 0.0394 |
| 235 | 241 | VLNISVF | F215C_PGM | 60 | 2.8198 | 0.042 |
| 235 | 241 | VLNISVF | F215C_PGM | 240 | 3.0238 | 0.0811 |
| 236 | 241 | LNISVF | F215C | 0 | 0 | 0 |
| 236 | 241 | LNISVF | F215C | 0.5 | 0.9427 | 0.0192 |

| | | | | | | |
|---|---|---|---|---|---|---|
| 236 | 241 | LNISVF | F215C | 1 | 1.0819 | 0.0371 |
| 236 | 241 | LNISVF | F215C | 2 | 1.3172 | 0.0499 |
| 236 | 241 | LNISVF | F215C | 5 | 1.5604 | 0.0478 |
| 236 | 241 | LNISVF | F215C | 30 | 2.0645 | 0.0498 |
| 236 | 241 | LNISVF | F215C | 60 | 2.2075 | 0.0502 |
| 236 | 241 | LNISVF | F215C | 240 | 2.3968 | 0.1035 |
| 236 | 241 | LNISVF | F215C_PGM | 0 | 0 | 0 |
| 236 | 241 | LNISVF | F215C_PGM | 0.5 | 0.7612 | 0.0293 |
| 236 | 241 | LNISVF | F215C_PGM | 1 | 0.9085 | 0.0287 |
| 236 | 241 | LNISVF | F215C_PGM | 2 | 1.1145 | 0.0501 |
| 236 | 241 | LNISVF | F215C_PGM | 5 | 1.4137 | 0.057 |
| 236 | 241 | LNISVF | F215C_PGM | 30 | 1.9895 | 0.0171 |
| 236 | 241 | LNISVF | F215C_PGM | 60 | 2.1444 | 0.0167 |
| 236 | 241 | LNISVF | F215C_PGM | 240 | 2.3685 | 0.0553 |
| 237 | 248 | NISVFPREVTNF | F215C | 0 | 0 | 0 |
| 237 | 248 | NISVFPREVTNF | F215C | 0.5 | 2.0007 | 0.1037 |
| 237 | 248 | NISVFPREVTNF | F215C | 1 | 2.2448 | 0.1003 |
| 237 | 248 | NISVFPREVTNF | F215C | 2 | 2.6174 | 0.0338 |
| 237 | 248 | NISVFPREVTNF | F215C | 5 | 3.0342 | 0.1019 |
| 237 | 248 | NISVFPREVTNF | F215C | 30 | 3.4079 | 0.1365 |
| 237 | 248 | NISVFPREVTNF | F215C | 60 | 3.5206 | 0.2051 |
| 237 | 248 | NISVFPREVTNF | F215C | 240 | 3.5578 | 0.1565 |
| 237 | 248 | NISVFPREVTNF | F215C_PGM | 0 | 0 | 0 |
| 237 | 248 | NISVFPREVTNF | F215C_PGM | 0.5 | 1.9954 | 0.0477 |
| 237 | 248 | NISVFPREVTNF | F215C_PGM | 1 | 2.4127 | 0.1878 |
| 237 | 248 | NISVFPREVTNF | F215C_PGM | 2 | 2.6946 | 0.0595 |
| 237 | 248 | NISVFPREVTNF | F215C_PGM | 5 | 3.1506 | 0.054 |
| 237 | 248 | NISVFPREVTNF | F215C_PGM | 30 | 3.9277 | 0.1272 |
| 237 | 248 | NISVFPREVTNF | F215C_PGM | 60 | 3.9827 | 0.1753 |
| 237 | 248 | NISVFPREVTNF | F215C_PGM | 240 | 3.7466 | 0.172 |
| 249 | 261 | LRKSVKRMKESRL | F215C | 0 | 0 | 0 |
| 249 | 261 | LRKSVKRMKESRL | F215C | 0.5 | 0.7714 | 0.0614 |
| 249 | 261 | LRKSVKRMKESRL | F215C | 1 | 0.8527 | 0.0603 |
| 249 | 261 | LRKSVKRMKESRL | F215C | 2 | 1.0939 | 0.1255 |
| 249 | 261 | LRKSVKRMKESRL | F215C | 5 | 1.1524 | 0.0843 |
| 249 | 261 | LRKSVKRMKESRL | F215C | 30 | 1.2954 | 0.0289 |
| 249 | 261 | LRKSVKRMKESRL | F215C | 60 | 1.2483 | 0.115 |
| 249 | 261 | LRKSVKRMKESRL | F215C | 240 | 1.1641 | 0.0428 |
| 249 | 261 | LRKSVKRMKESRL | F215C_PGM | 0 | 0 | 0 |
| 249 | 261 | LRKSVKRMKESRL | F215C_PGM | 0.5 | 0.7375 | 0.0498 |
| 249 | 261 | LRKSVKRMKESRL | F215C_PGM | 1 | 0.8239 | 0.1592 |
| 249 | 261 | LRKSVKRMKESRL | F215C_PGM | 2 | 1.0046 | 0.0487 |
| 249 | 261 | LRKSVKRMKESRL | F215C_PGM | 5 | 1.3469 | 0.0371 |
| 249 | 261 | LRKSVKRMKESRL | F215C_PGM | 30 | 1.5733 | 0.093 |
| 249 | 261 | LRKSVKRMKESRL | F215C_PGM | 60 | 1.6043 | 0.1614 |

| | | | | | | |
|---|---|---|---|---|---|---|
| 249 | 261 | LRKSVKRMKESRL | F215C_PGM | 240 | 1.3404 | 0.0542 |
| 262 | 271 | EDTQKHRVDF | F215C | 0 | 0 | 0 |
| 262 | 271 | EDTQKHRVDF | F215C | 0.5 | 0.8678 | 0.0381 |
| 262 | 271 | EDTQKHRVDF | F215C | 1 | 0.8124 | 0.0288 |
| 262 | 271 | EDTQKHRVDF | F215C | 2 | 0.9083 | 0.0217 |
| 262 | 271 | EDTQKHRVDF | F215C | 5 | 0.9478 | 0.0216 |
| 262 | 271 | EDTQKHRVDF | F215C | 30 | 1.0351 | 0.0483 |
| 262 | 271 | EDTQKHRVDF | F215C | 60 | 0.9961 | 0.0972 |
| 262 | 271 | EDTQKHRVDF | F215C | 240 | 1.0978 | 0.0779 |
| 262 | 271 | EDTQKHRVDF | F215C_PGM | 0 | 0 | 0 |
| 262 | 271 | EDTQKHRVDF | F215C_PGM | 0.5 | 0.8261 | 0.0333 |
| 262 | 271 | EDTQKHRVDF | F215C_PGM | 1 | 0.8221 | 0.102 |
| 262 | 271 | EDTQKHRVDF | F215C_PGM | 2 | 0.8853 | 0.06 |
| 262 | 271 | EDTQKHRVDF | F215C_PGM | 5 | 0.9582 | 0.0232 |
| 262 | 271 | EDTQKHRVDF | F215C_PGM | 30 | 1.1155 | 0.0587 |
| 262 | 271 | EDTQKHRVDF | F215C_PGM | 60 | 1.1609 | 0.0262 |
| 262 | 271 | EDTQKHRVDF | F215C_PGM | 240 | 1.0814 | 0.0698 |
| 275 | 295 | MIDSQNSKETESHKALSDLEL | F215C | 0 | 0 | 0 |
| 275 | 295 | MIDSQNSKETESHKALSDLEL | F215C | 0.5 | 2.4823 | 0.2289 |
| 275 | 295 | MIDSQNSKETESHKALSDLEL | F215C | 1 | 2.7209 | 0.17 |
| 275 | 295 | MIDSQNSKETESHKALSDLEL | F215C | 2 | 3.0844 | 0.2412 |
| 275 | 295 | MIDSQNSKETESHKALSDLEL | F215C | 5 | 3.8581 | 0.0256 |
| 275 | 295 | MIDSQNSKETESHKALSDLEL | F215C | 30 | 3.8776 | 0.1545 |
| 275 | 295 | MIDSQNSKETESHKALSDLEL | F215C | 60 | 3.9872 | 0.1806 |
| 275 | 295 | MIDSQNSKETESHKALSDLEL | F215C | 240 | 4.1565 | 0.2452 |
| 275 | 295 | MIDSQNSKETESHKALSDLEL | F215C_PGM | 0 | 0 | 0 |
| 275 | 295 | MIDSQNSKETESHKALSDLEL | F215C_PGM | 0.5 | 3.2092 | 0.0958 |
| 275 | 295 | MIDSQNSKETESHKALSDLEL | F215C_PGM | 1 | 3.4881 | 0.1753 |
| 275 | 295 | MIDSQNSKETESHKALSDLEL | F215C_PGM | 2 | 3.7976 | 0.1602 |
| 275 | 295 | MIDSQNSKETESHKALSDLEL | F215C_PGM | 5 | 4.199 | 0.0402 |
| 275 | 295 | MIDSQNSKETESHKALSDLEL | F215C_PGM | 30 | 4.7257 | 0.0589 |
| 275 | 295 | MIDSQNSKETESHKALSDLEL | F215C_PGM | 60 | 4.788 | 0.1309 |
| 275 | 295 | MIDSQNSKETESHKALSDLEL | F215C_PGM | 240 | 4.5922 | 0.2187 |
| 296 | 302 | VAQSIIF | F215C | 0 | 0 | 0 |
| 296 | 302 | VAQSIIF | F215C | 0.5 | 0.0056 | 0.1085 |
| 296 | 302 | VAQSIIF | F215C | 1 | 0.0761 | 0.0901 |
| 296 | 302 | VAQSIIF | F215C | 2 | 0.2619 | 0.094 |
| 296 | 302 | VAQSIIF | F215C | 5 | 0.2726 | 0.0818 |
| 296 | 302 | VAQSIIF | F215C | 30 | 0.7714 | 0.0669 |
| 296 | 302 | VAQSIIF | F215C | 60 | 0.7367 | 0.1715 |
| 296 | 302 | VAQSIIF | F215C | 240 | 1.0245 | 0.1068 |
| 296 | 302 | VAQSIIF | F215C_PGM | 0 | 0 | 0 |
| 296 | 302 | VAQSIIF | F215C_PGM | 0.5 | -0.1573 | 0.0641 |
| 296 | 302 | VAQSIIF | F215C_PGM | 1 | -0.1655 | 0.0659 |
| 296 | 302 | VAQSIIF | F215C_PGM | 2 | -0.0759 | 0.0788 |

| | | | | | | |
|---|---|---|---|---|---|---|
| 296 | 302 | VAQSIIF | F215C_PGM | 5 | 0.0549 | 0.0555 |
| 296 | 302 | VAQSIIF | F215C_PGM | 30 | 0.3431 | 0.0826 |
| 296 | 302 | VAQSIIF | F215C_PGM | 60 | 0.4098 | 0.0382 |
| 296 | 302 | VAQSIIF | F215C_PGM | 240 | 0.6312 | 0.0356 |
| 301 | 306 | IFIFAG | F215C | 0 | 0 | 0 |
| 301 | 306 | IFIFAG | F215C | 0.5 | 0.1028 | 0.0075 |
| 301 | 306 | IFIFAG | F215C | 1 | 0.1472 | 0.0106 |
| 301 | 306 | IFIFAG | F215C | 2 | 0.131 | 0.0374 |
| 301 | 306 | IFIFAG | F215C | 5 | 0.1681 | 0.0059 |
| 301 | 306 | IFIFAG | F215C | 30 | 0.3094 | 0.0342 |
| 301 | 306 | IFIFAG | F215C | 60 | 0.3868 | 0.0174 |
| 301 | 306 | IFIFAG | F215C | 240 | 0.5625 | 0.0439 |
| 301 | 306 | IFIFAG | F215C_PGM | 0 | 0 | 0 |
| 301 | 306 | IFIFAG | F215C_PGM | 0.5 | 0.0648 | 0.016 |
| 301 | 306 | IFIFAG | F215C_PGM | 1 | 0.0898 | 0.0085 |
| 301 | 306 | IFIFAG | F215C_PGM | 2 | 0.1088 | 0.017 |
| 301 | 306 | IFIFAG | F215C_PGM | 5 | 0.1754 | 0.0234 |
| 301 | 306 | IFIFAG | F215C_PGM | 30 | 0.2838 | 0.0346 |
| 301 | 306 | IFIFAG | F215C_PGM | 60 | 0.3185 | 0.044 |
| 301 | 306 | IFIFAG | F215C_PGM | 240 | 0.4392 | 0.0279 |
| 307 | 313 | YETTSSV | F215C | 0 | 0 | 0 |
| 307 | 313 | YETTSSV | F215C | 0.5 | 0.4667 | 0.0169 |
| 307 | 313 | YETTSSV | F215C | 1 | 0.5668 | 0.0536 |
| 307 | 313 | YETTSSV | F215C | 2 | 0.782 | 0.0433 |
| 307 | 313 | YETTSSV | F215C | 5 | 0.9846 | 0.0199 |
| 307 | 313 | YETTSSV | F215C | 30 | 1.3144 | 0.0361 |
| 307 | 313 | YETTSSV | F215C | 60 | 1.3589 | 0.0304 |
| 307 | 313 | YETTSSV | F215C | 240 | 1.5498 | 0.0777 |
| 307 | 313 | YETTSSV | F215C_PGM | 0 | 0 | 0 |
| 307 | 313 | YETTSSV | F215C_PGM | 0.5 | 0.3637 | 0.0569 |
| 307 | 313 | YETTSSV | F215C_PGM | 1 | 0.458 | 0.0263 |
| 307 | 313 | YETTSSV | F215C_PGM | 2 | 0.619 | 0.0174 |
| 307 | 313 | YETTSSV | F215C_PGM | 5 | 0.9218 | 0.0344 |
| 307 | 313 | YETTSSV | F215C_PGM | 30 | 1.3945 | 0.0517 |
| 307 | 313 | YETTSSV | F215C_PGM | 60 | 1.5747 | 0.0321 |
| 307 | 313 | YETTSSV | F215C_PGM | 240 | 1.6654 | 0.073 |
| 307 | 314 | YETTSSVL | F215C | 0 | 0 | 0 |
| 307 | 314 | YETTSSVL | F215C | 0.5 | 0.2055 | 0.0389 |
| 307 | 314 | YETTSSVL | F215C | 1 | 0.2537 | 0.0476 |
| 307 | 314 | YETTSSVL | F215C | 2 | 0.4123 | 0.0342 |
| 307 | 314 | YETTSSVL | F215C | 5 | 0.5266 | 0.0642 |
| 307 | 314 | YETTSSVL | F215C | 30 | 0.7324 | 0.072 |
| 307 | 314 | YETTSSVL | F215C | 60 | 0.7875 | 0.1103 |
| 307 | 314 | YETTSSVL | F215C | 240 | 0.9436 | 0.1668 |
| 307 | 314 | YETTSSVL | F215C_PGM | 0 | 0 | 0 |

| | | | | | | |
|---|---|---|---|---|---|---|
| 307 | 314 | YETTSSVL | F215C_PGM | 0.5 | 0.1951 | 0.0241 |
| 307 | 314 | YETTSSVL | F215C_PGM | 1 | 0.2948 | 0.0841 |
| 307 | 314 | YETTSSVL | F215C_PGM | 2 | 0.3766 | 0.0314 |
| 307 | 314 | YETTSSVL | F215C_PGM | 5 | 0.5763 | 0.0285 |
| 307 | 314 | YETTSSVL | F215C_PGM | 30 | 0.9007 | 0.0545 |
| 307 | 314 | YETTSSVL | F215C_PGM | 60 | 1.025 | 0.0636 |
| 307 | 314 | YETTSSVL | F215C_PGM | 240 | 1.073 | 0.0745 |
| 307 | 316 | YETTSSVLSF | F215C | 0 | 0 | 0 |
| 307 | 316 | YETTSSVLSF | F215C | 0.5 | 0.2669 | 0.0241 |
| 307 | 316 | YETTSSVLSF | F215C | 1 | 0.3217 | 0.0491 |
| 307 | 316 | YETTSSVLSF | F215C | 2 | 0.4828 | 0.0446 |
| 307 | 316 | YETTSSVLSF | F215C | 5 | 0.6338 | 0.0215 |
| 307 | 316 | YETTSSVLSF | F215C | 30 | 0.9845 | 0.05 |
| 307 | 316 | YETTSSVLSF | F215C | 60 | 1.0859 | 0.0691 |
| 307 | 316 | YETTSSVLSF | F215C | 240 | 1.406 | 0.0738 |
| 307 | 316 | YETTSSVLSF | F215C_PGM | 0 | 0 | 0 |
| 307 | 316 | YETTSSVLSF | F215C_PGM | 0.5 | 0.1875 | 0.0314 |
| 307 | 316 | YETTSSVLSF | F215C_PGM | 1 | 0.2975 | 0.0606 |
| 307 | 316 | YETTSSVLSF | F215C_PGM | 2 | 0.4012 | 0.0291 |
| 307 | 316 | YETTSSVLSF | F215C_PGM | 5 | 0.6003 | 0.0125 |
| 307 | 316 | YETTSSVLSF | F215C_PGM | 30 | 0.984 | 0.0443 |
| 307 | 316 | YETTSSVLSF | F215C_PGM | 60 | 1.1503 | 0.0355 |
| 307 | 316 | YETTSSVLSF | F215C_PGM | 240 | 1.335 | 0.0857 |
| 319 | 333 | YELATHPDVQQKLQE | F215C | 0 | 0 | 0 |
| 319 | 333 | YELATHPDVQQKLQE | F215C | 0.5 | 0.2757 | 0.0532 |
| 319 | 333 | YELATHPDVQQKLQE | F215C | 1 | 0.4362 | 0.1091 |
| 319 | 333 | YELATHPDVQQKLQE | F215C | 2 | 0.463 | 0.0898 |
| 319 | 333 | YELATHPDVQQKLQE | F215C | 5 | 0.5692 | 0.1029 |
| 319 | 333 | YELATHPDVQQKLQE | F215C | 30 | 0.8595 | 0.113 |
| 319 | 333 | YELATHPDVQQKLQE | F215C | 60 | 0.9647 | 0.0357 |
| 319 | 333 | YELATHPDVQQKLQE | F215C | 240 | 1.258 | 0.0318 |
| 319 | 333 | YELATHPDVQQKLQE | F215C_PGM | 0 | 0 | 0 |
| 319 | 333 | YELATHPDVQQKLQE | F215C_PGM | 0.5 | 0.3857 | 0.0812 |
| 319 | 333 | YELATHPDVQQKLQE | F215C_PGM | 1 | 0.4599 | 0.0982 |
| 319 | 333 | YELATHPDVQQKLQE | F215C_PGM | 2 | 0.5602 | 0.0489 |
| 319 | 333 | YELATHPDVQQKLQE | F215C_PGM | 5 | 0.6243 | 0.071 |
| 319 | 333 | YELATHPDVQQKLQE | F215C_PGM | 30 | 1.0182 | 0.1027 |
| 319 | 333 | YELATHPDVQQKLQE | F215C_PGM | 60 | 1.1595 | 0.0288 |
| 319 | 333 | YELATHPDVQQKLQE | F215C_PGM | 240 | 1.2579 | 0.0819 |
| 322 | 333 | ATHPDVQQKLQE | F215C | 0 | 0 | 0 |
| 322 | 333 | ATHPDVQQKLQE | F215C | 0.5 | 0.0212 | 0.0855 |
| 322 | 333 | ATHPDVQQKLQE | F215C | 1 | -0.0123 | 0.075 |
| 322 | 333 | ATHPDVQQKLQE | F215C | 2 | 0.0089 | 0.0521 |
| 322 | 333 | ATHPDVQQKLQE | F215C | 5 | 0.0975 | 0.0547 |
| 322 | 333 | ATHPDVQQKLQE | F215C | 30 | 0.5016 | 0.053 |

| | | | | | | |
|---|---|---|---|---|---|---|
| 322 | 333 | ATHPDVQQKLQE | F215C | 60 | 0.5201 | 0.0573 |
| 322 | 333 | ATHPDVQQKLQE | F215C | 240 | 0.6815 | 0.0612 |
| 322 | 333 | ATHPDVQQKLQE | F215C_PGM | 0 | 0 | 0 |
| 322 | 333 | ATHPDVQQKLQE | F215C_PGM | 0.5 | -0.1421 | 0.0376 |
| 322 | 333 | ATHPDVQQKLQE | F215C_PGM | 1 | -0.0942 | 0.0313 |
| 322 | 333 | ATHPDVQQKLQE | F215C_PGM | 2 | -0.0254 | 0.0344 |
| 322 | 333 | ATHPDVQQKLQE | F215C_PGM | 5 | 0.1022 | 0.0307 |
| 322 | 333 | ATHPDVQQKLQE | F215C_PGM | 30 | 0.4687 | 0.0406 |
| 322 | 333 | ATHPDVQQKLQE | F215C_PGM | 60 | 0.527 | 0.0305 |
| 322 | 333 | ATHPDVQQKLQE | F215C_PGM | 240 | 0.7422 | 0.0502 |
| 332 | 337 | QEEIDA | F215C | 0 | 0 | 0 |
| 332 | 337 | QEEIDA | F215C | 0.5 | 0.1546 | 0.0379 |
| 332 | 337 | QEEIDA | F215C | 1 | 0.1901 | 0.0308 |
| 332 | 337 | QEEIDA | F215C | 2 | 0.2866 | 0.023 |
| 332 | 337 | QEEIDA | F215C | 5 | 0.464 | 0.0177 |
| 332 | 337 | QEEIDA | F215C | 30 | 0.9557 | 0.0171 |
| 332 | 337 | QEEIDA | F215C | 60 | 1.033 | 0.0379 |
| 332 | 337 | QEEIDA | F215C | 240 | 1.3987 | 0.0129 |
| 332 | 337 | QEEIDA | F215C_PGM | 0 | 0 | 0 |
| 332 | 337 | QEEIDA | F215C_PGM | 0.5 | 0.1742 | 0.0171 |
| 332 | 337 | QEEIDA | F215C_PGM | 1 | 0.2296 | 0.0234 |
| 332 | 337 | QEEIDA | F215C_PGM | 2 | 0.3194 | 0.0131 |
| 332 | 337 | QEEIDA | F215C_PGM | 5 | 0.5076 | 0.0286 |
| 332 | 337 | QEEIDA | F215C_PGM | 30 | 1.0129 | 0.0238 |
| 332 | 337 | QEEIDA | F215C_PGM | 60 | 1.1817 | 0.0375 |
| 332 | 337 | QEEIDA | F215C_PGM | 240 | 1.4388 | 0.0307 |
| 337 | 349 | AVLPNKAPPTYDT | F215C | 0 | 0 | 0 |
| 337 | 349 | AVLPNKAPPTYDT | F215C | 0.5 | 1.7491 | 0.0901 |
| 337 | 349 | AVLPNKAPPTYDT | F215C | 1 | 1.9262 | 0.0838 |
| 337 | 349 | AVLPNKAPPTYDT | F215C | 2 | 2.2684 | 0.0327 |
| 337 | 349 | AVLPNKAPPTYDT | F215C | 5 | 2.631 | 0.0253 |
| 337 | 349 | AVLPNKAPPTYDT | F215C | 30 | 3.0005 | 0.1261 |
| 337 | 349 | AVLPNKAPPTYDT | F215C | 60 | 3.0722 | 0.1939 |
| 337 | 349 | AVLPNKAPPTYDT | F215C | 240 | 3.1515 | 0.3068 |
| 337 | 349 | AVLPNKAPPTYDT | F215C_PGM | 0 | 0 | 0 |
| 337 | 349 | AVLPNKAPPTYDT | F215C_PGM | 0.5 | 1.9531 | 0.0525 |
| 337 | 349 | AVLPNKAPPTYDT | F215C_PGM | 1 | 2.1415 | 0.1428 |
| 337 | 349 | AVLPNKAPPTYDT | F215C_PGM | 2 | 2.4207 | 0.0622 |
| 337 | 349 | AVLPNKAPPTYDT | F215C_PGM | 5 | 2.8007 | 0.1057 |
| 337 | 349 | AVLPNKAPPTYDT | F215C_PGM | 30 | 3.3042 | 0.0686 |
| 337 | 349 | AVLPNKAPPTYDT | F215C_PGM | 60 | 3.4317 | 0.0303 |
| 337 | 349 | AVLPNKAPPTYDT | F215C_PGM | 240 | 3.2677 | 0.1198 |
| 340 | 347 | PNKAPPTY | F215C | 0 | 0 | 0 |
| 340 | 347 | PNKAPPTY | F215C | 0.5 | 0.9159 | 0.0159 |
| 340 | 347 | PNKAPPTY | F215C | 1 | 0.9841 | 0.0384 |

| | | | | | | |
|---|---|---|---|---|---|---|
| 340 | 347 | PNKAPPTY | F215C | 2 | 1.1312 | 0.038 |
| 340 | 347 | PNKAPPTY | F215C | 5 | 1.4747 | 0.1292 |
| 340 | 347 | PNKAPPTY | F215C | 30 | 1.7177 | 0.0389 |
| 340 | 347 | PNKAPPTY | F215C | 60 | 1.7695 | 0.067 |
| 340 | 347 | PNKAPPTY | F215C | 240 | 1.7971 | 0.0699 |
| 340 | 347 | PNKAPPTY | F215C_PGM | 0 | 0 | 0 |
| 340 | 347 | PNKAPPTY | F215C_PGM | 0.5 | 1.0262 | 0.0259 |
| 340 | 347 | PNKAPPTY | F215C_PGM | 1 | 1.0757 | 0.075 |
| 340 | 347 | PNKAPPTY | F215C_PGM | 2 | 1.1915 | 0.0401 |
| 340 | 347 | PNKAPPTY | F215C_PGM | 5 | 1.4689 | 0.073 |
| 340 | 347 | PNKAPPTY | F215C_PGM | 30 | 1.9414 | 0.0723 |
| 340 | 347 | PNKAPPTY | F215C_PGM | 60 | 1.9135 | 0.0634 |
| 340 | 347 | PNKAPPTY | F215C_PGM | 240 | 1.8764 | 0.0593 |
| 353 | 358 | MEYLDM | F215C | 0 | 0 | 0 |
| 353 | 358 | MEYLDM | F215C | 0.5 | 0.2987 | 0.0366 |
| 353 | 358 | MEYLDM | F215C | 1 | 0.3752 | 0.0975 |
| 353 | 358 | MEYLDM | F215C | 2 | 0.3773 | 0.0339 |
| 353 | 358 | MEYLDM | F215C | 5 | 0.3856 | 0.0649 |
| 353 | 358 | MEYLDM | F215C | 30 | 0.4669 | 0.0396 |
| 353 | 358 | MEYLDM | F215C | 60 | 0.5637 | 0.0979 |
| 353 | 358 | MEYLDM | F215C | 240 | 0.8085 | 0.109 |
| 353 | 358 | MEYLDM | F215C_PGM | 0 | 0 | 0 |
| 353 | 358 | MEYLDM | F215C_PGM | 0.5 | 0.2608 | 0.0414 |
| 353 | 358 | MEYLDM | F215C_PGM | 1 | 0.2983 | 0.0834 |
| 353 | 358 | MEYLDM | F215C_PGM | 2 | 0.3622 | 0.0998 |
| 353 | 358 | MEYLDM | F215C_PGM | 5 | 0.2961 | 0.0351 |
| 353 | 358 | MEYLDM | F215C_PGM | 30 | 0.3382 | 0.0389 |
| 353 | 358 | MEYLDM | F215C_PGM | 60 | 0.4424 | 0.0342 |
| 353 | 358 | MEYLDM | F215C_PGM | 240 | 0.792 | 0.0968 |
| 357 | 363 | DMVVNET | F215C | 0 | 0 | 0 |
| 357 | 363 | DMVVNET | F215C | 0.5 | 0.0245 | 0.0373 |
| 357 | 363 | DMVVNET | F215C | 1 | 0.0153 | 0.0358 |
| 357 | 363 | DMVVNET | F215C | 2 | 0.0277 | 0.0655 |
| 357 | 363 | DMVVNET | F215C | 5 | -0.0013 | 0.0538 |
| 357 | 363 | DMVVNET | F215C | 30 | 0.083 | 0.0241 |
| 357 | 363 | DMVVNET | F215C | 60 | 0.0695 | 0.0284 |
| 357 | 363 | DMVVNET | F215C | 240 | 0.0447 | 0.0291 |
| 357 | 363 | DMVVNET | F215C_PGM | 0 | 0 | 0 |
| 357 | 363 | DMVVNET | F215C_PGM | 0.5 | -0.0059 | 0.0246 |
| 357 | 363 | DMVVNET | F215C_PGM | 1 | -0.0104 | 0.028 |
| 357 | 363 | DMVVNET | F215C_PGM | 2 | -0.0205 | 0.0305 |
| 357 | 363 | DMVVNET | F215C_PGM | 5 | 0.0056 | 0.0287 |
| 357 | 363 | DMVVNET | F215C_PGM | 30 | -0.0074 | 0.0287 |
| 357 | 363 | DMVVNET | F215C_PGM | 60 | 0.0219 | 0.0277 |
| 357 | 363 | DMVVNET | F215C_PGM | 240 | 0.0403 | 0.0276 |

| | | | | | | |
|---|---|---|---|---|---|---|
| 359 | 366 | VVNETLRL | F215C | 0 | 0 | 0 |
| 359 | 366 | VVNETLRL | F215C | 0.5 | 0.0307 | 0.0306 |
| 359 | 366 | VVNETLRL | F215C | 1 | 0.0582 | 0.0652 |
| 359 | 366 | VVNETLRL | F215C | 2 | -0.0027 | 0.0274 |
| 359 | 366 | VVNETLRL | F215C | 5 | 0.0234 | 0.0517 |
| 359 | 366 | VVNETLRL | F215C | 30 | 0.1481 | 0.0923 |
| 359 | 366 | VVNETLRL | F215C | 60 | 0.1047 | 0.0334 |
| 359 | 366 | VVNETLRL | F215C | 240 | 0.2108 | 0.0739 |
| 359 | 366 | VVNETLRL | F215C_PGM | 0 | 0 | 0 |
| 359 | 366 | VVNETLRL | F215C_PGM | 0.5 | 0.1033 | 0.0381 |
| 359 | 366 | VVNETLRL | F215C_PGM | 1 | 0.0694 | 0.0419 |
| 359 | 366 | VVNETLRL | F215C_PGM | 2 | 0.0913 | 0.0263 |
| 359 | 366 | VVNETLRL | F215C_PGM | 5 | 0.0984 | 0.0405 |
| 359 | 366 | VVNETLRL | F215C_PGM | 30 | 0.2043 | 0.0289 |
| 359 | 366 | VVNETLRL | F215C_PGM | 60 | 0.2172 | 0.0387 |
| 359 | 366 | VVNETLRL | F215C_PGM | 240 | 0.31 | 0.0303 |
| 363 | 370 | TLRLFPIA | F215C | 0 | 0 | 0 |
| 363 | 370 | TLRLFPIA | F215C | 0.5 | 0.1079 | 0.0201 |
| 363 | 370 | TLRLFPIA | F215C | 1 | 0.1602 | 0.0325 |
| 363 | 370 | TLRLFPIA | F215C | 2 | 0.1649 | 0.0774 |
| 363 | 370 | TLRLFPIA | F215C | 5 | 0.1797 | 0.0447 |
| 363 | 370 | TLRLFPIA | F215C | 30 | 0.4137 | 0.0439 |
| 363 | 370 | TLRLFPIA | F215C | 60 | 0.4675 | 0.0538 |
| 363 | 370 | TLRLFPIA | F215C | 240 | 0.6639 | 0.0327 |
| 363 | 370 | TLRLFPIA | F215C_PGM | 0 | 0 | 0 |
| 363 | 370 | TLRLFPIA | F215C_PGM | 0.5 | 0.0739 | 0.0377 |
| 363 | 370 | TLRLFPIA | F215C_PGM | 1 | 0.0716 | 0.0185 |
| 363 | 370 | TLRLFPIA | F215C_PGM | 2 | 0.0621 | 0.0349 |
| 363 | 370 | TLRLFPIA | F215C_PGM | 5 | 0.1152 | 0.0256 |
| 363 | 370 | TLRLFPIA | F215C_PGM | 30 | 0.2208 | 0.0582 |
| 363 | 370 | TLRLFPIA | F215C_PGM | 60 | 0.2872 | 0.0353 |
| 363 | 370 | TLRLFPIA | F215C_PGM | 240 | 0.4688 | 0.032 |
| 374 | 385 | ERVCKKDVEING | F215C | 0 | 0 | 0 |
| 374 | 385 | ERVCKKDVEING | F215C | 0.5 | 1.0125 | 0.02 |
| 374 | 385 | ERVCKKDVEING | F215C | 1 | 1.2274 | 0.146 |
| 374 | 385 | ERVCKKDVEING | F215C | 2 | 1.2733 | 0.0377 |
| 374 | 385 | ERVCKKDVEING | F215C | 5 | 1.3761 | 0.0387 |
| 374 | 385 | ERVCKKDVEING | F215C | 30 | 1.5075 | 0.0374 |
| 374 | 385 | ERVCKKDVEING | F215C | 60 | 1.4816 | 0.1043 |
| 374 | 385 | ERVCKKDVEING | F215C | 240 | 1.5858 | 0.042 |
| 374 | 385 | ERVCKKDVEING | F215C_PGM | 0 | 0 | 0 |
| 374 | 385 | ERVCKKDVEING | F215C_PGM | 0.5 | 0.7008 | 0.0566 |
| 374 | 385 | ERVCKKDVEING | F215C_PGM | 1 | 0.7734 | 0.1006 |
| 374 | 385 | ERVCKKDVEING | F215C_PGM | 2 | 0.9863 | 0.0458 |
| 374 | 385 | ERVCKKDVEING | F215C_PGM | 5 | 1.161 | 0.0336 |

| | | | | | | |
|---|---|---|---|---|---|---|
| 374 | 385 | ERVCKKDVEING | F215C_PGM | 30 | 1.296 | 0.0412 |
| 374 | 385 | ERVCKKDVEING | F215C_PGM | 60 | 1.3355 | 0.0447 |
| 374 | 385 | ERVCKKDVEING | F215C_PGM | 240 | 1.3522 | 0.0754 |
| 387 | 393 | FIPKGVV | F215C | 0 | 0 | 0 |
| 387 | 393 | FIPKGVV | F215C | 0.5 | 0.3156 | 0.056 |
| 387 | 393 | FIPKGVV | F215C | 1 | 0.4212 | 0.0356 |
| 387 | 393 | FIPKGVV | F215C | 2 | 0.4271 | 0.0297 |
| 387 | 393 | FIPKGVV | F215C | 5 | 0.4645 | 0.0609 |
| 387 | 393 | FIPKGVV | F215C | 30 | 0.5997 | 0.0507 |
| 387 | 393 | FIPKGVV | F215C | 60 | 0.6355 | 0.0932 |
| 387 | 393 | FIPKGVV | F215C | 240 | 0.7151 | 0.0609 |
| 387 | 393 | FIPKGVV | F215C_PGM | 0 | 0 | 0 |
| 387 | 393 | FIPKGVV | F215C_PGM | 0.5 | 0.4258 | 0.0822 |
| 387 | 393 | FIPKGVV | F215C_PGM | 1 | 0.4876 | 0.0341 |
| 387 | 393 | FIPKGVV | F215C_PGM | 2 | 0.4946 | 0.0361 |
| 387 | 393 | FIPKGVV | F215C_PGM | 5 | 0.5306 | 0.0255 |
| 387 | 393 | FIPKGVV | F215C_PGM | 30 | 0.6369 | 0.0269 |
| 387 | 393 | FIPKGVV | F215C_PGM | 60 | 0.732 | 0.0369 |
| 387 | 393 | FIPKGVV | F215C_PGM | 240 | 0.7118 | 0.0536 |
| 388 | 393 | IPKGVV | F215C | 0 | 0 | 0 |
| 388 | 393 | IPKGVV | F215C | 0.5 | 0.3371 | 0.0293 |
| 388 | 393 | IPKGVV | F215C | 1 | 0.3438 | 0.017 |
| 388 | 393 | IPKGVV | F215C | 2 | 0.3947 | 0.0227 |
| 388 | 393 | IPKGVV | F215C | 5 | 0.4377 | 0.0188 |
| 388 | 393 | IPKGVV | F215C | 30 | 0.6503 | 0.0254 |
| 388 | 393 | IPKGVV | F215C | 60 | 0.6567 | 0.0213 |
| 388 | 393 | IPKGVV | F215C | 240 | 0.7493 | 0.0171 |
| 388 | 393 | IPKGVV | F215C_PGM | 0 | 0 | 0 |
| 388 | 393 | IPKGVV | F215C_PGM | 0.5 | 0.3889 | 0.0215 |
| 388 | 393 | IPKGVV | F215C_PGM | 1 | 0.4369 | 0.0163 |
| 388 | 393 | IPKGVV | F215C_PGM | 2 | 0.4888 | 0.0153 |
| 388 | 393 | IPKGVV | F215C_PGM | 5 | 0.5449 | 0.0223 |
| 388 | 393 | IPKGVV | F215C_PGM | 30 | 0.686 | 0.0219 |
| 388 | 393 | IPKGVV | F215C_PGM | 60 | 0.7486 | 0.0151 |
| 388 | 393 | IPKGVV | F215C_PGM | 240 | 0.8617 | 0.0178 |
| 388 | 395 | IPKGVVVM | F215C | 0 | 0 | 0 |
| 388 | 395 | IPKGVVVM | F215C | 0.5 | 0.5003 | 0.0474 |
| 388 | 395 | IPKGVVVM | F215C | 1 | 0.5904 | 0.1041 |
| 388 | 395 | IPKGVVVM | F215C | 2 | 0.679 | 0.0666 |
| 388 | 395 | IPKGVVVM | F215C | 5 | 0.7547 | 0.075 |
| 388 | 395 | IPKGVVVM | F215C | 30 | 1.1137 | 0.1018 |
| 388 | 395 | IPKGVVVM | F215C | 60 | 1.1295 | 0.0448 |
| 388 | 395 | IPKGVVVM | F215C | 240 | 1.2613 | 0.1193 |
| 388 | 395 | IPKGVVVM | F215C_PGM | 0 | 0 | 0 |
| 388 | 395 | IPKGVVVM | F215C_PGM | 0.5 | 0.6937 | 0.0883 |

| | | | | | | |
|---|---|---|---|---|---|---|
| 388 | 395 | IPKGVVVM | F215C_PGM | 1 | 0.7133 | 0.0708 |
| 388 | 395 | IPKGVVVM | F215C_PGM | 2 | 0.8412 | 0.0465 |
| 388 | 395 | IPKGVVVM | F215C_PGM | 5 | 0.9478 | 0.0515 |
| 388 | 395 | IPKGVVVM | F215C_PGM | 30 | 1.2136 | 0.0498 |
| 388 | 395 | IPKGVVVM | F215C_PGM | 60 | 1.3366 | 0.0328 |
| 388 | 395 | IPKGVVVM | F215C_PGM | 240 | 1.3745 | 0.0448 |
| 394 | 399 | VMIPSY | F215C | 0 | 0 | 0 |
| 394 | 399 | VMIPSY | F215C | 0.5 | 0.2249 | 0.0101 |
| 394 | 399 | VMIPSY | F215C | 1 | 0.2533 | 0.0175 |
| 394 | 399 | VMIPSY | F215C | 2 | 0.29 | 0.0312 |
| 394 | 399 | VMIPSY | F215C | 5 | 0.3433 | 0.0283 |
| 394 | 399 | VMIPSY | F215C | 30 | 0.4384 | 0.0229 |
| 394 | 399 | VMIPSY | F215C | 60 | 0.4945 | 0.0477 |
| 394 | 399 | VMIPSY | F215C | 240 | 0.5733 | 0.0531 |
| 394 | 399 | VMIPSY | F215C_PGM | 0 | 0 | 0 |
| 394 | 399 | VMIPSY | F215C_PGM | 0.5 | 0.2388 | 0.0289 |
| 394 | 399 | VMIPSY | F215C_PGM | 1 | 0.2456 | 0.023 |
| 394 | 399 | VMIPSY | F215C_PGM | 2 | 0.2917 | 0.0234 |
| 394 | 399 | VMIPSY | F215C_PGM | 5 | 0.3253 | 0.019 |
| 394 | 399 | VMIPSY | F215C_PGM | 30 | 0.4456 | 0.0355 |
| 394 | 399 | VMIPSY | F215C_PGM | 60 | 0.5157 | 0.0228 |
| 394 | 399 | VMIPSY | F215C_PGM | 240 | 0.472 | 0.0192 |
| 394 | 401 | VMIPSYAL | F215C | 0 | 0 | 0 |
| 394 | 401 | VMIPSYAL | F215C | 0.5 | 0.5415 | 0.0399 |
| 394 | 401 | VMIPSYAL | F215C | 1 | 0.5405 | 0.069 |
| 394 | 401 | VMIPSYAL | F215C | 2 | 0.6592 | 0.0599 |
| 394 | 401 | VMIPSYAL | F215C | 5 | 0.7107 | 0.0975 |
| 394 | 401 | VMIPSYAL | F215C | 30 | 0.9099 | 0.0412 |
| 394 | 401 | VMIPSYAL | F215C | 60 | 0.9707 | 0.1152 |
| 394 | 401 | VMIPSYAL | F215C | 240 | 1.06 | 0.086 |
| 394 | 401 | VMIPSYAL | F215C_PGM | 0 | 0 | 0 |
| 394 | 401 | VMIPSYAL | F215C_PGM | 0.5 | 0.5522 | 0.0406 |
| 394 | 401 | VMIPSYAL | F215C_PGM | 1 | 0.618 | 0.0433 |
| 394 | 401 | VMIPSYAL | F215C_PGM | 2 | 0.6938 | 0.0771 |
| 394 | 401 | VMIPSYAL | F215C_PGM | 5 | 0.7603 | 0.0465 |
| 394 | 401 | VMIPSYAL | F215C_PGM | 30 | 0.891 | 0.1066 |
| 394 | 401 | VMIPSYAL | F215C_PGM | 60 | 0.8643 | 0.0443 |
| 394 | 401 | VMIPSYAL | F215C_PGM | 240 | 0.9752 | 0.057 |
| 400 | 407 | ALHRDPKY | F215C | 0 | 0 | 0 |
| 400 | 407 | ALHRDPKY | F215C | 0.5 | 0.1717 | 0.0516 |
| 400 | 407 | ALHRDPKY | F215C | 1 | 0.2216 | 0.0643 |
| 400 | 407 | ALHRDPKY | F215C | 2 | 0.2946 | 0.0483 |
| 400 | 407 | ALHRDPKY | F215C | 5 | 0.2331 | 0.1298 |
| 400 | 407 | ALHRDPKY | F215C | 30 | 0.4064 | 0.063 |
| 400 | 407 | ALHRDPKY | F215C | 60 | 0.4409 | 0.0713 |

| | | | | | | |
|---|---|---|---|---|---|---|
| 400 | 407 | ALHRDPKY | F215C | 240 | 0.5833 | 0.0805 |
| 400 | 407 | ALHRDPKY | F215C_PGM | 0 | 0 | 0 |
| 400 | 407 | ALHRDPKY | F215C_PGM | 0.5 | 0.0679 | 0.0353 |
| 400 | 407 | ALHRDPKY | F215C_PGM | 1 | 0.1234 | 0.0436 |
| 400 | 407 | ALHRDPKY | F215C_PGM | 2 | 0.227 | 0.0427 |
| 400 | 407 | ALHRDPKY | F215C_PGM | 5 | 0.2838 | 0.0347 |
| 400 | 407 | ALHRDPKY | F215C_PGM | 30 | 0.3547 | 0.0398 |
| 400 | 407 | ALHRDPKY | F215C_PGM | 60 | 0.3985 | 0.0452 |
| 400 | 407 | ALHRDPKY | F215C_PGM | 240 | 0.507 | 0.046 |
| 408 | 414 | WTEPEKF | F215C | 0 | 0 | 0 |
| 408 | 414 | WTEPEKF | F215C | 0.5 | 1.1084 | 0.0659 |
| 408 | 414 | WTEPEKF | F215C | 1 | 1.1249 | 0.031 |
| 408 | 414 | WTEPEKF | F215C | 2 | 1.22 | 0.0276 |
| 408 | 414 | WTEPEKF | F215C | 5 | 1.2117 | 0.0088 |
| 408 | 414 | WTEPEKF | F215C | 30 | 1.24 | 0.1505 |
| 408 | 414 | WTEPEKF | F215C | 60 | 1.2377 | 0.1645 |
| 408 | 414 | WTEPEKF | F215C | 240 | 1.1425 | 0.0758 |
| 408 | 414 | WTEPEKF | F215C_PGM | 0 | 0 | 0 |
| 408 | 414 | WTEPEKF | F215C_PGM | 0.5 | 1.1802 | 0.0155 |
| 408 | 414 | WTEPEKF | F215C_PGM | 1 | 1.1854 | 0.1206 |
| 408 | 414 | WTEPEKF | F215C_PGM | 2 | 1.2082 | 0.0292 |
| 408 | 414 | WTEPEKF | F215C_PGM | 5 | 1.2407 | 0.0396 |
| 408 | 414 | WTEPEKF | F215C_PGM | 30 | 1.2633 | 0.0225 |
| 408 | 414 | WTEPEKF | F215C_PGM | 60 | 1.2904 | 0.042 |
| 408 | 414 | WTEPEKF | F215C_PGM | 240 | 1.1671 | 0.066 |
| 420 | 444 | SKKNKDNIDPYIYTPFGSGPRNCIG | F215C | 0 | 0 | 0 |
| 420 | 444 | SKKNKDNIDPYIYTPFGSGPRNCIG | F215C | 0.5 | 2.2682 | 0.101 |
| 420 | 444 | SKKNKDNIDPYIYTPFGSGPRNCIG | F215C | 1 | 2.4645 | 0.0666 |
| 420 | 444 | SKKNKDNIDPYIYTPFGSGPRNCIG | F215C | 2 | 2.8617 | 0.0336 |
| 420 | 444 | SKKNKDNIDPYIYTPFGSGPRNCIG | F215C | 5 | 3.4064 | 0.0229 |
| 420 | 444 | SKKNKDNIDPYIYTPFGSGPRNCIG | F215C | 30 | 3.9211 | 0.0411 |
| 420 | 444 | SKKNKDNIDPYIYTPFGSGPRNCIG | F215C | 60 | 4.0874 | 0.0513 |
| 420 | 444 | SKKNKDNIDPYIYTPFGSGPRNCIG | F215C | 240 | 4.5632 | 0.1027 |
| 420 | 444 | SKKNKDNIDPYIYTPFGSGPRNCIG | F215C_PGM | 0 | 0 | 0 |
| 420 | 444 | SKKNKDNIDPYIYTPFGSGPRNCIG | F215C_PGM | 0.5 | 2.2626 | 0.0245 |
| 420 | 444 | SKKNKDNIDPYIYTPFGSGPRNCIG | F215C_PGM | 1 | 2.5984 | 0.111 |
| 420 | 444 | SKKNKDNIDPYIYTPFGSGPRNCIG | F215C_PGM | 2 | 2.9108 | 0.0539 |
| 420 | 444 | SKKNKDNIDPYIYTPFGSGPRNCIG | F215C_PGM | 5 | 3.4013 | 0.0538 |
| 420 | 444 | SKKNKDNIDPYIYTPFGSGPRNCIG | F215C_PGM | 30 | 3.9352 | 0.0344 |
| 420 | 444 | SKKNKDNIDPYIYTPFGSGPRNCIG | F215C_PGM | 60 | 4.1157 | 0.077 |
| 420 | 444 | SKKNKDNIDPYIYTPFGSGPRNCIG | F215C_PGM | 240 | 4.4021 | 0.1893 |
| 445 | 452 | MRFALMNM | F215C | 0 | 0 | 0 |
| 445 | 452 | MRFALMNM | F215C | 0.5 | -0.0908 | 0.0226 |
| 445 | 452 | MRFALMNM | F215C | 1 | -0.0082 | 0.0574 |
| 445 | 452 | MRFALMNM | F215C | 2 | 0.0164 | 0.0522 |

| | | | | | | |
|---|---|---|---|---|---|---|
| 445 | 452 | MRFALMNM | F215C | 5 | 0.0874 | 0.0284 |
| 445 | 452 | MRFALMNM | F215C | 30 | 0.1593 | 0.0333 |
| 445 | 452 | MRFALMNM | F215C | 60 | 0.2086 | 0.0504 |
| 445 | 452 | MRFALMNM | F215C | 240 | 0.4877 | 0.0553 |
| 445 | 452 | MRFALMNM | F215C_PGM | 0 | 0 | 0 |
| 445 | 452 | MRFALMNM | F215C_PGM | 0.5 | -0.0763 | 0.0375 |
| 445 | 452 | MRFALMNM | F215C_PGM | 1 | 0.0504 | 0.029 |
| 445 | 452 | MRFALMNM | F215C_PGM | 2 | 0.1223 | 0.0788 |
| 445 | 452 | MRFALMNM | F215C_PGM | 5 | 0.1642 | 0.0833 |
| 445 | 452 | MRFALMNM | F215C_PGM | 30 | 0.2275 | 0.1017 |
| 445 | 452 | MRFALMNM | F215C_PGM | 60 | 0.3037 | 0.0961 |
| 445 | 452 | MRFALMNM | F215C_PGM | 240 | 0.2633 | 0.1855 |
| 446 | 452 | RFALMNM | F215C | 0 | 0 | 0 |
| 446 | 452 | RFALMNM | F215C | 0.5 | 0.1599 | 0.0636 |
| 446 | 452 | RFALMNM | F215C | 1 | 0.1618 | 0.0402 |
| 446 | 452 | RFALMNM | F215C | 2 | 0.1562 | 0.0382 |
| 446 | 452 | RFALMNM | F215C | 5 | 0.1685 | 0.0263 |
| 446 | 452 | RFALMNM | F215C | 30 | 0.2761 | 0.0324 |
| 446 | 452 | RFALMNM | F215C | 60 | 0.3038 | 0.0404 |
| 446 | 452 | RFALMNM | F215C | 240 | 0.4513 | 0.0505 |
| 446 | 452 | RFALMNM | F215C_PGM | 0 | 0 | 0 |
| 446 | 452 | RFALMNM | F215C_PGM | 0.5 | 0.1172 | 0.0222 |
| 446 | 452 | RFALMNM | F215C_PGM | 1 | 0.118 | 0.0292 |
| 446 | 452 | RFALMNM | F215C_PGM | 2 | 0.1127 | 0.0412 |
| 446 | 452 | RFALMNM | F215C_PGM | 5 | 0.1477 | 0.0262 |
| 446 | 452 | RFALMNM | F215C_PGM | 30 | 0.2682 | 0.0526 |
| 446 | 452 | RFALMNM | F215C_PGM | 60 | 0.3358 | 0.0348 |
| 446 | 452 | RFALMNM | F215C_PGM | 240 | 0.484 | 0.0355 |
| 455 | 463 | ALIRVLQNF | F215C | 0 | 0 | 0 |
| 455 | 463 | ALIRVLQNF | F215C | 0.5 | 0.0652 | 0.0777 |
| 455 | 463 | ALIRVLQNF | F215C | 1 | 0.0836 | 0.0527 |
| 455 | 463 | ALIRVLQNF | F215C | 2 | 0.0704 | 0.0468 |
| 455 | 463 | ALIRVLQNF | F215C | 5 | 0.0584 | 0.0366 |
| 455 | 463 | ALIRVLQNF | F215C | 30 | 0.3055 | 0.0451 |
| 455 | 463 | ALIRVLQNF | F215C | 60 | 0.301 | 0.0784 |
| 455 | 463 | ALIRVLQNF | F215C | 240 | 0.4691 | 0.0576 |
| 455 | 463 | ALIRVLQNF | F215C_PGM | 0 | 0 | 0 |
| 455 | 463 | ALIRVLQNF | F215C_PGM | 0.5 | 0.1517 | 0.0409 |
| 455 | 463 | ALIRVLQNF | F215C_PGM | 1 | 0.1419 | 0.0274 |
| 455 | 463 | ALIRVLQNF | F215C_PGM | 2 | 0.1572 | 0.0384 |
| 455 | 463 | ALIRVLQNF | F215C_PGM | 5 | 0.2098 | 0.0226 |
| 455 | 463 | ALIRVLQNF | F215C_PGM | 30 | 0.3562 | 0.0481 |
| 455 | 463 | ALIRVLQNF | F215C_PGM | 60 | 0.4059 | 0.0463 |
| 455 | 463 | ALIRVLQNF | F215C_PGM | 240 | 0.5383 | 0.0331 |
| 457 | 463 | IRVLQNF | F215C | 0 | 0 | 0 |

| | | | | | | |
|---|---|---|---|---|---|---|
| 457 | 463 | IRVLQNF | F215C | 0.5 | 0.1407 | 0.025 |
| 457 | 463 | IRVLQNF | F215C | 1 | 0.1238 | 0.0244 |
| 457 | 463 | IRVLQNF | F215C | 2 | 0.1499 | 0.0243 |
| 457 | 463 | IRVLQNF | F215C | 5 | 0.1839 | 0.0226 |
| 457 | 463 | IRVLQNF | F215C | 30 | 0.3368 | 0.0266 |
| 457 | 463 | IRVLQNF | F215C | 60 | 0.3129 | 0.0293 |
| 457 | 463 | IRVLQNF | F215C | 240 | 0.4746 | 0.0513 |
| 457 | 463 | IRVLQNF | F215C_PGM | 0 | 0 | 0 |
| 457 | 463 | IRVLQNF | F215C_PGM | 0.5 | 0.1752 | 0.0348 |
| 457 | 463 | IRVLQNF | F215C_PGM | 1 | 0.1898 | 0.0305 |
| 457 | 463 | IRVLQNF | F215C_PGM | 2 | 0.2309 | 0.0334 |
| 457 | 463 | IRVLQNF | F215C_PGM | 5 | 0.2701 | 0.0448 |
| 457 | 463 | IRVLQNF | F215C_PGM | 30 | 0.3776 | 0.0395 |
| 457 | 463 | IRVLQNF | F215C_PGM | 60 | 0.4093 | 0.0427 |
| 457 | 463 | IRVLQNF | F215C_PGM | 240 | 0.5422 | 0.0347 |
| 464 | 479 | SFKPGKETQIPLKLSL | F215C | 0 | 0 | 0 |
| 464 | 479 | SFKPGKETQIPLKLSL | F215C | 0.5 | 3.7729 | 0.2122 |
| 464 | 479 | SFKPGKETQIPLKLSL | F215C | 1 | 3.8784 | 0.1346 |
| 464 | 479 | SFKPGKETQIPLKLSL | F215C | 2 | 4.1538 | 0.0706 |
| 464 | 479 | SFKPGKETQIPLKLSL | F215C | 5 | 4.2833 | 0.0706 |
| 464 | 479 | SFKPGKETQIPLKLSL | F215C | 30 | 4.1702 | 0.1459 |
| 464 | 479 | SFKPGKETQIPLKLSL | F215C | 60 | 4.0776 | 0.153 |
| 464 | 479 | SFKPGKETQIPLKLSL | F215C | 240 | 4.1101 | 0.2696 |
| 464 | 479 | SFKPGKETQIPLKLSL | F215C_PGM | 0 | 0 | 0 |
| 464 | 479 | SFKPGKETQIPLKLSL | F215C_PGM | 0.5 | 3.9599 | 0.0494 |
| 464 | 479 | SFKPGKETQIPLKLSL | F215C_PGM | 1 | 4.0299 | 0.3206 |
| 464 | 479 | SFKPGKETQIPLKLSL | F215C_PGM | 2 | 4.2479 | 0.1073 |
| 464 | 479 | SFKPGKETQIPLKLSL | F215C_PGM | 5 | 4.4012 | 0.123 |
| 464 | 479 | SFKPGKETQIPLKLSL | F215C_PGM | 30 | 4.5021 | 0.0266 |
| 464 | 479 | SFKPGKETQIPLKLSL | F215C_PGM | 60 | 4.3685 | 0.0216 |
| 464 | 479 | SFKPGKETQIPLKLSL | F215C_PGM | 240 | 4.526 | 0.191 |
| 464 | 491 | SFKPGKETQIPLKLSLGGLLQPEKP | F215C | 0 | 0 | 0 |
| 464 | 491 | SFKPGKETQIPLKLSLGGLLQPEKP | F215C | 0.5 | 5.4398 | 0.2149 |
| 464 | 491 | SFKPGKETQIPLKLSLGGLLQPEKP | F215C | 1 | 6.0703 | 0.2095 |
| 464 | 491 | SFKPGKETQIPLKLSLGGLLQPEKP | F215C | 2 | 7.0382 | 0.1727 |
| 464 | 491 | SFKPGKETQIPLKLSLGGLLQPEKP | F215C | 5 | | |
| 464 | 491 | SFKPGKETQIPLKLSLGGLLQPEKP | F215C | 30 | 7.6117 | 0.0198 |
| 464 | 491 | SFKPGKETQIPLKLSLGGLLQPEKP | F215C | 60 | 7.7313 | 0.029 |
| 464 | 491 | SFKPGKETQIPLKLSLGGLLQPEKP | F215C | 240 | 7.8655 | 0.0198 |
| 464 | 491 | SFKPGKETQIPLKLSLGGLLQPEKP | F215C_PGM | 0 | 0 | 0 |
| 464 | 491 | SFKPGKETQIPLKLSLGGLLQPEKP | F215C_PGM | 0.5 | 5.6213 | 0.0198 |
| 464 | 491 | SFKPGKETQIPLKLSLGGLLQPEKP | F215C_PGM | 1 | 5.9187 | 0.2537 |
| 464 | 491 | SFKPGKETQIPLKLSLGGLLQPEKP | F215C_PGM | 2 | 6.4873 | 0.0717 |
| 464 | 491 | SFKPGKETQIPLKLSLGGLLQPEKP | F215C_PGM | 5 | 7.0947 | 0.1334 |
| 464 | 491 | SFKPGKETQIPLKLSLGGLLQPEKP | F215C_PGM | 30 | 7.895 | 0.0198 |

| | | | | | | |
|---|---|---|---|---|---|---|
| 464 | 491 | SFKPGKETQIPLKLSLGGLLQPEKP | F215C_PGM | 60 | 7.7113 | 0.0958 |
| 464 | 491 | SFKPGKETQIPLKLSLGGLLQPEKP | F215C_PGM | 240 | 7.6249 | 0.3228 |
| 485 | 491 | PEKPVVL | F215C | 0 | 0 | 0 |
| 485 | 491 | PEKPVVL | F215C | 0.5 | 1.4409 | 0.0576 |
| 485 | 491 | PEKPVVL | F215C | 1 | 1.6528 | 0.0327 |
| 485 | 491 | PEKPVVL | F215C | 2 | 1.9565 | 0.0318 |
| 485 | 491 | PEKPVVL | F215C | 5 | 2.1989 | 0.0238 |
| 485 | 491 | PEKPVVL | F215C | 30 | 2.3308 | 0.0631 |
| 485 | 491 | PEKPVVL | F215C | 60 | 2.3259 | 0.0039 |
| 485 | 491 | PEKPVVL | F215C | 240 | 2.3615 | 0.0086 |
| 485 | 491 | PEKPVVL | F215C_PGM | 0 | 0 | 0 |
| 485 | 491 | PEKPVVL | F215C_PGM | 0.5 | 1.4849 | 0.0495 |
| 485 | 491 | PEKPVVL | F215C_PGM | 1 | 1.5935 | 0.0653 |
| 485 | 491 | PEKPVVL | F215C_PGM | 2 | 1.8928 | 0.0962 |
| 485 | 491 | PEKPVVL | F215C_PGM | 5 | 2.1706 | 0.018 |
| 485 | 491 | PEKPVVL | F215C_PGM | 30 | 2.4608 | 0.0735 |
| 485 | 491 | PEKPVVL | F215C_PGM | 60 | 2.4626 | 0.082 |
| 485 | 491 | PEKPVVL | F215C_PGM | 240 | 2.471 | 0.0841 |
| 492 | 507 | KVESRDGTVSGAHHHH | F215C | 0 | 0 | 0 |
| 492 | 507 | KVESRDGTVSGAHHHH | F215C | 0.5 | 0.8183 | 0.1392 |
| 492 | 507 | KVESRDGTVSGAHHHH | F215C | 1 | 0.8317 | 0.0766 |
| 492 | 507 | KVESRDGTVSGAHHHH | F215C | 2 | 0.9503 | 0.0911 |
| 492 | 507 | KVESRDGTVSGAHHHH | F215C | 5 | 0.9513 | 0.0937 |
| 492 | 507 | KVESRDGTVSGAHHHH | F215C | 30 | 0.9007 | 0.077 |
| 492 | 507 | KVESRDGTVSGAHHHH | F215C | 60 | 0.9044 | 0.121 |
| 492 | 507 | KVESRDGTVSGAHHHH | F215C | 240 | 0.9079 | 0.0994 |
| 492 | 507 | KVESRDGTVSGAHHHH | F215C_PGM | 0 | 0 | 0 |
| 492 | 507 | KVESRDGTVSGAHHHH | F215C_PGM | 0.5 | 0.9067 | 0.06 |
| 492 | 507 | KVESRDGTVSGAHHHH | F215C_PGM | 1 | 0.8993 | 0.1723 |
| 492 | 507 | KVESRDGTVSGAHHHH | F215C_PGM | 2 | 0.9664 | 0.1103 |
| 492 | 507 | KVESRDGTVSGAHHHH | F215C_PGM | 5 | 0.9681 | 0.0776 |
| 492 | 507 | KVESRDGTVSGAHHHH | F215C_PGM | 30 | 1.0291 | 0.064 |
| 492 | 507 | KVESRDGTVSGAHHHH | F215C_PGM | 60 | 1.0138 | 0.0809 |
| 492 | 507 | KVESRDGTVSGAHHHH | F215C_PGM | 240 | 0.8941 | 0.0998 |
| 494 | 507 | ESRDGTVSGAHHHH | F215C | 0 | 0 | 0 |
| 494 | 507 | ESRDGTVSGAHHHH | F215C | 0.5 | 0.7754 | 0.2336 |
| 494 | 507 | ESRDGTVSGAHHHH | F215C | 1 | 0.8916 | 0.086 |
| 494 | 507 | ESRDGTVSGAHHHH | F215C | 2 | 1.0096 | 0.1128 |
| 494 | 507 | ESRDGTVSGAHHHH | F215C | 5 | 0.6906 | 0.0678 |
| 494 | 507 | ESRDGTVSGAHHHH | F215C | 30 | 0.732 | 0.0745 |
| 494 | 507 | ESRDGTVSGAHHHH | F215C | 60 | 0.8877 | 0.1838 |
| 494 | 507 | ESRDGTVSGAHHHH | F215C | 240 | 0.9317 | 0.063 |
| 494 | 507 | ESRDGTVSGAHHHH | F215C_PGM | 0 | 0 | 0 |
| 494 | 507 | ESRDGTVSGAHHHH | F215C_PGM | 0.5 | 0.7897 | 0.115 |
| 494 | 507 | ESRDGTVSGAHHHH | F215C_PGM | 1 | 0.523 | 0.0262 |

| | | | | | | |
|---|---|---|---|---|---|---|
| 494 | 507 | ESRDGTVSGAHHHH | F215C_PGM | 2 | 0.3085 | 0.0262 |
| 494 | 507 | ESRDGTVSGAHHHH | F215C_PGM | 5 | 0.9005 | 0.117 |
| 494 | 507 | ESRDGTVSGAHHHH | F215C_PGM | 30 | 1.2901 | 0.3454 |
| 494 | 507 | ESRDGTVSGAHHHH | F215C_PGM | 60 | 0.6757 | 0.1678 |
| 494 | 507 | ESRDGTVSGAHHHH | F215C_PGM | 240 | 1.0582 | 0.217 |

| Deuterium uptake data for F215C and F215C_PRG CYP3A4 states | | | | | | |
|---|---|---|---|---|---|---|
| **Start** | **End** | **Sequence** | **State** | **Exposure (min)** | **Uptake (Da)** | **Uptake SD (Da)** |
| 33 | 48 | FKKLGIPGPTPLPFLG | F215C | 0 | 0.000000 | 0.000000 |
| 33 | 48 | FKKLGIPGPTPLPFLG | F215C | 5 | 0.433837 | 0.122801 |
| 33 | 48 | FKKLGIPGPTPLPFLG | F215C | 60 | 0.403173 | 0.100579 |
| 33 | 48 | FKKLGIPGPTPLPFLG | F215C_PRG | 0 | 0.000000 | 0.000000 |
| 33 | 48 | FKKLGIPGPTPLPFLG | F215C_PRG | 5 | 0.471338 | 0.131273 |
| 33 | 48 | FKKLGIPGPTPLPFLG | F215C_PRG | 60 | 0.255826 | 0.133107 |
| 33 | 51 | FKKLGIPGPTPLPFLGNIL | F215C | 0 | 0.000000 | 0.000000 |
| 33 | 51 | FKKLGIPGPTPLPFLGNIL | F215C | 5 | 0.475812 | 0.099766 |
| 33 | 51 | FKKLGIPGPTPLPFLGNIL | F215C | 60 | 0.360873 | 0.168932 |
| 33 | 51 | FKKLGIPGPTPLPFLGNIL | F215C_PRG | 0 | 0.000000 | 0.000000 |
| 33 | 51 | FKKLGIPGPTPLPFLGNIL | F215C_PRG | 5 | 0.622150 | 0.045759 |
| 33 | 51 | FKKLGIPGPTPLPFLGNIL | F215C_PRG | 60 | 0.504159 | 0.041725 |
| 34 | 51 | KKLGIPGPTPLPFLGNIL | F215C | 0 | 0.000000 | 0.000000 |
| 34 | 51 | KKLGIPGPTPLPFLGNIL | F215C | 5 | 0.422684 | 0.048735 |
| 34 | 51 | KKLGIPGPTPLPFLGNIL | F215C | 60 | 0.511415 | 0.063787 |
| 34 | 51 | KKLGIPGPTPLPFLGNIL | F215C_PRG | 0 | 0.000000 | 0.000000 |
| 34 | 51 | KKLGIPGPTPLPFLGNIL | F215C_PRG | 5 | 0.516997 | 0.103176 |
| 34 | 51 | KKLGIPGPTPLPFLGNIL | F215C_PRG | 60 | 0.441847 | 0.036422 |
| 49 | 60 | NILSYHKGFTMF | F215C | 0 | 0.000000 | 0.000000 |
| 49 | 60 | NILSYHKGFTMF | F215C | 5 | 2.017402 | 0.044050 |
| 49 | 60 | NILSYHKGFTMF | F215C | 60 | 2.005704 | 0.163678 |
| 49 | 60 | NILSYHKGFTMF | F215C_PRG | 0 | 0.000000 | 0.000000 |
| 49 | 60 | NILSYHKGFTMF | F215C_PRG | 5 | 2.317295 | 0.069480 |
| 49 | 60 | NILSYHKGFTMF | F215C_PRG | 60 | 2.304271 | 0.059787 |
| 61 | 74 | DMEAHKKYGKVWGF | F215C | 0 | 0.000000 | 0.000000 |
| 61 | 74 | DMEAHKKYGKVWGF | F215C | 5 | 1.003194 | 0.078629 |
| 61 | 74 | DMEAHKKYGKVWGF | F215C | 60 | 1.534021 | 0.114330 |
| 61 | 74 | DMEAHKKYGKVWGF | F215C_PRG | 0 | 0.000000 | 0.000000 |
| 61 | 74 | DMEAHKKYGKVWGF | F215C_PRG | 5 | 1.065498 | 0.090787 |
| 61 | 74 | DMEAHKKYGKVWGF | F215C_PRG | 60 | 1.646033 | 0.093735 |
| 61 | 82 | DMEAHKKYGKVWGFYDGQQPVL | F215C | 0 | 0.000000 | 0.000000 |
| 61 | 82 | DMEAHKKYGKVWGFYDGQQPVL | F215C | 5 | 1.712281 | 0.067264 |
| 61 | 82 | DMEAHKKYGKVWGFYDGQQPVL | F215C | 60 | 2.520236 | 0.116017 |
| 61 | 82 | DMEAHKKYGKVWGFYDGQQPVL | F215C_PRG | 0 | 0.000000 | 0.000000 |
| 61 | 82 | DMEAHKKYGKVWGFYDGQQPVL | F215C_PRG | 5 | 1.863633 | 0.062024 |
| 61 | 82 | DMEAHKKYGKVWGFYDGQQPVL | F215C_PRG | 60 | 2.760389 | 0.116101 |

| | | | | | | |
|---|---|---|---|---|---|---|
| 74 | 82 | FYDGQQPVL | F215C | 0 | 0.000000 | 0.000000 |
| 74 | 82 | FYDGQQPVL | F215C | 5 | 1.021257 | 0.051469 |
| 74 | 82 | FYDGQQPVL | F215C | 60 | 1.412009 | 0.117408 |
| 74 | 82 | FYDGQQPVL | F215C_PRG | 0 | 0.000000 | 0.000000 |
| 74 | 82 | FYDGQQPVL | F215C_PRG | 5 | 1.114111 | 0.026981 |
| 74 | 82 | FYDGQQPVL | F215C_PRG | 60 | 1.449956 | 0.068749 |
| 75 | 82 | YDGQQPVL | F215C | 0 | 0.000000 | 0.000000 |
| 75 | 82 | YDGQQPVL | F215C | 5 | 0.851474 | 0.037782 |
| 75 | 82 | YDGQQPVL | F215C | 60 | 1.099367 | 0.075239 |
| 75 | 82 | YDGQQPVL | F215C_PRG | 0 | 0.000000 | 0.000000 |
| 75 | 82 | YDGQQPVL | F215C_PRG | 5 | 0.862383 | 0.012805 |
| 75 | 82 | YDGQQPVL | F215C_PRG | 60 | 1.154540 | 0.056124 |
| 83 | 89 | AITDPDM | F215C | 0 | 0.000000 | 0.000000 |
| 83 | 89 | AITDPDM | F215C | 5 | 0.485450 | 0.030894 |
| 83 | 89 | AITDPDM | F215C | 60 | 0.602014 | 0.075856 |
| 83 | 89 | AITDPDM | F215C_PRG | 0 | 0.000000 | 0.000000 |
| 83 | 89 | AITDPDM | F215C_PRG | 5 | 0.520982 | 0.007005 |
| 83 | 89 | AITDPDM | F215C_PRG | 60 | 0.710115 | 0.045545 |
| 83 | 94 | AITDPDMIKTVL | F215C | 0 | 0.000000 | 0.000000 |
| 83 | 94 | AITDPDMIKTVL | F215C | 5 | 1.328048 | 0.053532 |
| 83 | 94 | AITDPDMIKTVL | F215C | 60 | 1.995947 | 0.093543 |
| 83 | 94 | AITDPDMIKTVL | F215C_PRG | 0 | 0.000000 | 0.000000 |
| 83 | 94 | AITDPDMIKTVL | F215C_PRG | 5 | 0.973171 | 0.124875 |
| 83 | 94 | AITDPDMIKTVL | F215C_PRG | 60 | 1.484254 | 0.107325 |
| 93 | 102 | VLVKESYSVF | F215C | 0 | 0.000000 | 0.000000 |
| 93 | 102 | VLVKESYSVF | F215C | 5 | 3.014780 | 0.037684 |
| 93 | 102 | VLVKESYSVF | F215C | 60 | 3.044100 | 0.031593 |
| 93 | 102 | VLVKESYSVF | F215C_PRG | 0 | 0.000000 | 0.000000 |
| 93 | 102 | VLVKESYSVF | F215C_PRG | 5 | 2.749955 | 0.031321 |
| 93 | 102 | VLVKESYSVF | F215C_PRG | 60 | 2.837822 | 0.069183 |
| 94 | 106 | LVKESYSVFTNRR | F215C | 0 | 0.000000 | 0.000000 |
| 94 | 106 | LVKESYSVFTNRR | F215C | 5 | 2.218446 | 0.062680 |
| 94 | 106 | LVKESYSVFTNRR | F215C | 60 | 2.542321 | 0.033844 |
| 94 | 106 | LVKESYSVFTNRR | F215C_PRG | 0 | 0.000000 | 0.000000 |
| 94 | 106 | LVKESYSVFTNRR | F215C_PRG | 5 | 1.799088 | 0.045109 |
| 94 | 106 | LVKESYSVFTNRR | F215C_PRG | 60 | 1.980516 | 0.023138 |
| 95 | 102 | VKESYSVF | F215C | 0 | 0.000000 | 0.000000 |
| 95 | 102 | VKESYSVF | F215C | 5 | 1.625188 | 0.043928 |
| 95 | 102 | VKESYSVF | F215C | 60 | 1.764673 | 0.087199 |
| 95 | 102 | VKESYSVF | F215C_PRG | 0 | 0.000000 | 0.000000 |
| 95 | 102 | VKESYSVF | F215C_PRG | 5 | 1.898969 | 0.028250 |
| 95 | 102 | VKESYSVF | F215C_PRG | 60 | 2.103225 | 0.025047 |
| 114 | 122 | MKSAISIAE | F215C | 0 | 0.000000 | 0.000000 |
| 114 | 122 | MKSAISIAE | F215C | 5 | 1.676461 | 0.045615 |
| 114 | 122 | MKSAISIAE | F215C | 60 | 2.620923 | 0.012307 |

| | | | | | | |
|---|---|---|---|---|---|---|
| 114 | 122 | MKSAISIAE | F215C_PRG | 0 | 0.000000 | 0.000000 |
| 114 | 122 | MKSAISIAE | F215C_PRG | 5 | 1.646253 | 0.069841 |
| 114 | 122 | MKSAISIAE | F215C_PRG | 60 | 2.489048 | 0.109403 |
| 120 | 125 | IAEDEE | F215C | 0 | 0.000000 | 0.000000 |
| 120 | 125 | IAEDEE | F215C | 5 | 0.888258 | 0.038403 |
| 120 | 125 | IAEDEE | F215C | 60 | 0.928197 | 0.033075 |
| 120 | 125 | IAEDEE | F215C_PRG | 0 | 0.000000 | 0.000000 |
| 120 | 125 | IAEDEE | F215C_PRG | 5 | 0.782634 | 0.028678 |
| 120 | 125 | IAEDEE | F215C_PRG | 60 | 0.903725 | 0.072679 |
| 123 | 133 | DEEWKRLRSLL | F215C | 0 | 0.000000 | 0.000000 |
| 123 | 133 | DEEWKRLRSLL | F215C | 5 | 1.631899 | 0.095752 |
| 123 | 133 | DEEWKRLRSLL | F215C | 60 | 2.323779 | 0.160533 |
| 123 | 133 | DEEWKRLRSLL | F215C_PRG | 0 | 0.000000 | 0.000000 |
| 123 | 133 | DEEWKRLRSLL | F215C_PRG | 5 | 1.972068 | 0.049233 |
| 123 | 133 | DEEWKRLRSLL | F215C_PRG | 60 | 2.351470 | 0.112164 |
| 123 | 137 | DEEWKRLRSLLSPTF | F215C | 0 | 0.000000 | 0.000000 |
| 123 | 137 | DEEWKRLRSLLSPTF | F215C | 5 | 2.139862 | 0.016592 |
| 123 | 137 | DEEWKRLRSLLSPTF | F215C | 60 | 3.268288 | 0.042248 |
| 123 | 137 | DEEWKRLRSLLSPTF | F215C_PRG | 0 | 0.000000 | 0.000000 |
| 123 | 137 | DEEWKRLRSLLSPTF | F215C_PRG | 5 | 2.254674 | 0.073523 |
| 123 | 137 | DEEWKRLRSLLSPTF | F215C_PRG | 60 | 3.361700 | 0.112259 |
| 126 | 137 | WKRLRSLLSPTF | F215C | 0 | 0.000000 | 0.000000 |
| 126 | 137 | WKRLRSLLSPTF | F215C | 5 | 1.526685 | 0.064055 |
| 126 | 137 | WKRLRSLLSPTF | F215C | 60 | 2.273678 | 0.047244 |
| 126 | 137 | WKRLRSLLSPTF | F215C_PRG | 0 | 0.000000 | 0.000000 |
| 126 | 137 | WKRLRSLLSPTF | F215C_PRG | 5 | 1.615112 | 0.087957 |
| 126 | 137 | WKRLRSLLSPTF | F215C_PRG | 60 | 2.312758 | 0.026500 |
| 134 | 151 | SPTFTSGKLKEMVPIIAQ | F215C | 0 | 0.000000 | 0.000000 |
| 134 | 151 | SPTFTSGKLKEMVPIIAQ | F215C | 5 | 3.361873 | 0.051438 |
| 134 | 151 | SPTFTSGKLKEMVPIIAQ | F215C | 60 | 4.385223 | 0.063278 |
| 134 | 151 | SPTFTSGKLKEMVPIIAQ | F215C_PRG | 0 | 0.000000 | 0.000000 |
| 134 | 151 | SPTFTSGKLKEMVPIIAQ | F215C_PRG | 5 | 3.252570 | 0.057574 |
| 134 | 151 | SPTFTSGKLKEMVPIIAQ | F215C_PRG | 60 | 4.391681 | 0.163598 |
| 135 | 151 | PTFTSGKLKEMVPIIAQ | F215C | 0 | 0.000000 | 0.000000 |
| 135 | 151 | PTFTSGKLKEMVPIIAQ | F215C | 5 | 3.550765 | 0.040179 |
| 135 | 151 | PTFTSGKLKEMVPIIAQ | F215C | 60 | 4.734142 | 0.043618 |
| 135 | 151 | PTFTSGKLKEMVPIIAQ | F215C_PRG | 0 | 0.000000 | 0.000000 |
| 135 | 151 | PTFTSGKLKEMVPIIAQ | F215C_PRG | 5 | 3.478805 | 0.048540 |
| 135 | 151 | PTFTSGKLKEMVPIIAQ | F215C_PRG | 60 | 4.574048 | 0.153064 |
| 138 | 151 | TSGKLKEMVPIIAQ | F215C | 0 | 0.000000 | 0.000000 |
| 138 | 151 | TSGKLKEMVPIIAQ | F215C | 5 | 3.001574 | 0.021701 |
| 138 | 151 | TSGKLKEMVPIIAQ | F215C | 60 | 4.088994 | 0.083572 |
| 138 | 151 | TSGKLKEMVPIIAQ | F215C_PRG | 0 | 0.000000 | 0.000000 |
| 138 | 151 | TSGKLKEMVPIIAQ | F215C_PRG | 5 | 2.963602 | 0.108442 |
| 138 | 151 | TSGKLKEMVPIIAQ | F215C_PRG | 60 | 3.976557 | 0.135461 |

| | | | | | | |
|---|---|---|---|---|---|---|
| 157 | 178 | VRNLRREAETGKPVTLKDVFGA | F215C | 0 | 0.000000 | 0.000000 |
| 157 | 178 | VRNLRREAETGKPVTLKDVFGA | F215C | 5 | 6.540628 | 0.192783 |
| 157 | 178 | VRNLRREAETGKPVTLKDVFGA | F215C | 60 | 7.266910 | 0.095108 |
| 157 | 178 | VRNLRREAETGKPVTLKDVFGA | F215C_PRG | 0 | 0.000000 | 0.000000 |
| 157 | 178 | VRNLRREAETGKPVTLKDVFGA | F215C_PRG | 5 | 6.481874 | 0.126743 |
| 157 | 178 | VRNLRREAETGKPVTLKDVFGA | F215C_PRG | 60 | 7.222749 | 0.316301 |
| 173 | 178 | KDVFGA | F215C | 0 | 0.000000 | 0.000000 |
| 173 | 178 | KDVFGA | F215C | 5 | 1.201806 | 0.039994 |
| 173 | 178 | KDVFGA | F215C | 60 | 1.771405 | 0.043445 |
| 173 | 178 | KDVFGA | F215C_PRG | 0 | 0.000000 | 0.000000 |
| 173 | 178 | KDVFGA | F215C_PRG | 5 | 1.085190 | 0.015027 |
| 173 | 178 | KDVFGA | F215C_PRG | 60 | 1.490471 | 0.041664 |
| 173 | 181 | KDVFGAYSM | F215C | 0 | 0.000000 | 0.000000 |
| 173 | 181 | KDVFGAYSM | F215C | 5 | 1.318286 | 0.105720 |
| 173 | 181 | KDVFGAYSM | F215C | 60 | 1.913944 | 0.216161 |
| 173 | 181 | KDVFGAYSM | F215C_PRG | 0 | 0.000000 | 0.000000 |
| 173 | 181 | KDVFGAYSM | F215C_PRG | 5 | 1.570646 | 0.087041 |
| 173 | 181 | KDVFGAYSM | F215C_PRG | 60 | 2.270595 | 0.090902 |
| 177 | 182 | GAYSMD | F215C | 0 | 0.000000 | 0.000000 |
| 177 | 182 | GAYSMD | F215C | 5 | 0.423805 | 0.044790 |
| 177 | 182 | GAYSMD | F215C | 60 | 0.683070 | 0.027162 |
| 177 | 182 | GAYSMD | F215C_PRG | 0 | 0.000000 | 0.000000 |
| 177 | 182 | GAYSMD | F215C_PRG | 5 | 0.369102 | 0.028883 |
| 177 | 182 | GAYSMD | F215C_PRG | 60 | 0.564120 | 0.043850 |
| 182 | 189 | DVITSTSF | F215C | 0 | 0.000000 | 0.000000 |
| 182 | 189 | DVITSTSF | F215C | 5 | 0.873258 | 0.029543 |
| 182 | 189 | DVITSTSF | F215C | 60 | 1.269042 | 0.097905 |
| 182 | 189 | DVITSTSF | F215C_PRG | 0 | 0.000000 | 0.000000 |
| 182 | 189 | DVITSTSF | F215C_PRG | 5 | 1.047147 | 0.011779 |
| 182 | 189 | DVITSTSF | F215C_PRG | 60 | 1.503471 | 0.006712 |
| 183 | 189 | VITSTSF | F215C | 0 | 0.000000 | 0.000000 |
| 183 | 189 | VITSTSF | F215C | 5 | 0.619015 | 0.025010 |
| 183 | 189 | VITSTSF | F215C | 60 | 0.830090 | 0.100575 |
| 183 | 189 | VITSTSF | F215C_PRG | 0 | 0.000000 | 0.000000 |
| 183 | 189 | VITSTSF | F215C_PRG | 5 | 0.749137 | 0.026952 |
| 183 | 189 | VITSTSF | F215C_PRG | 60 | 1.163511 | 0.007369 |
| 189 | 212 | FGVNIDSLNNPQDPFVENTKKLLR | F215C | 0 | 0.000000 | 0.000000 |
| 189 | 212 | FGVNIDSLNNPQDPFVENTKKLLR | F215C | 5 | 6.456315 | 0.155934 |
| 189 | 212 | FGVNIDSLNNPQDPFVENTKKLLR | F215C | 60 | 7.921372 | 0.089334 |
| 189 | 212 | FGVNIDSLNNPQDPFVENTKKLLR | F215C_PRG | 0 | 0.000000 | 0.000000 |
| 189 | 212 | FGVNIDSLNNPQDPFVENTKKLLR | F215C_PRG | 5 | 6.279265 | 0.176709 |
| 189 | 212 | FGVNIDSLNNPQDPFVENTKKLLR | F215C_PRG | 60 | 7.137933 | 0.149523 |
| 190 | 210 | GVNIDSLNNPQDPFVENTKKL | F215C | 0 | 0.000000 | 0.000000 |
| 190 | 210 | GVNIDSLNNPQDPFVENTKKL | F215C | 5 | 5.175734 | 0.162861 |
| 190 | 210 | GVNIDSLNNPQDPFVENTKKL | F215C | 60 | 5.780868 | 0.087124 |

| | | | | | | |
|---|---|---|---|---|---|---|
| 190 | 210 | GVNIDSLNNPQDPFVENTKKL | F215C_PRG | 0 | 0.000000 | 0.000000 |
| 190 | 210 | GVNIDSLNNPQDPFVENTKKL | F215C_PRG | 5 | 6.451054 | 0.286461 |
| 190 | 210 | GVNIDSLNNPQDPFVENTKKL | F215C_PRG | 60 | 7.190351 | 0.341071 |
| 221 | 226 | LSITVF | F215C | 0 | 0.000000 | 0.000000 |
| 221 | 226 | LSITVF | F215C | 5 | 1.017416 | 0.077418 |
| 221 | 226 | LSITVF | F215C | 60 | 1.560797 | 0.049236 |
| 221 | 226 | LSITVF | F215C_PRG | 0 | 0.000000 | 0.000000 |
| 221 | 226 | LSITVF | F215C_PRG | 5 | 0.871834 | 0.033323 |
| 221 | 226 | LSITVF | F215C_PRG | 60 | 1.395290 | 0.052886 |
| 230 | 235 | IPILEV | F215C | 0 | 0.000000 | 0.000000 |
| 230 | 235 | IPILEV | F215C | 5 | 0.371514 | 0.071435 |
| 230 | 235 | IPILEV | F215C | 60 | 1.154628 | 0.050174 |
| 230 | 235 | IPILEV | F215C_PRG | 0 | 0.000000 | 0.000000 |
| 230 | 235 | IPILEV | F215C_PRG | 5 | 0.377022 | 0.013062 |
| 230 | 235 | IPILEV | F215C_PRG | 60 | 0.861828 | 0.025326 |
| 233 | 247 | LEVLNISVFPREVTN | F215C | 0 | 0.000000 | 0.000000 |
| 233 | 247 | LEVLNISVFPREVTN | F215C | 5 | 2.830988 | 0.028120 |
| 233 | 247 | LEVLNISVFPREVTN | F215C | 60 | 3.703204 | 0.043882 |
| 233 | 247 | LEVLNISVFPREVTN | F215C_PRG | 0 | 0.000000 | 0.000000 |
| 233 | 247 | LEVLNISVFPREVTN | F215C_PRG | 5 | 2.813224 | 0.074288 |
| 233 | 247 | LEVLNISVFPREVTN | F215C_PRG | 60 | 3.651065 | 0.136875 |
| 235 | 241 | VLNISVF | F215C | 0 | 0.000000 | 0.000000 |
| 235 | 241 | VLNISVF | F215C | 5 | 1.970933 | 0.030963 |
| 235 | 241 | VLNISVF | F215C | 60 | 2.848865 | 0.041770 |
| 235 | 241 | VLNISVF | F215C_PRG | 0 | 0.000000 | 0.000000 |
| 235 | 241 | VLNISVF | F215C_PRG | 5 | 2.021958 | 0.050504 |
| 235 | 241 | VLNISVF | F215C_PRG | 60 | 2.736164 | 0.068028 |
| 236 | 241 | LNISVF | F215C | 0 | 0.000000 | 0.000000 |
| 236 | 241 | LNISVF | F215C | 5 | 1.560358 | 0.047760 |
| 236 | 241 | LNISVF | F215C | 60 | 2.207487 | 0.050228 |
| 236 | 241 | LNISVF | F215C_PRG | 0 | 0.000000 | 0.000000 |
| 236 | 241 | LNISVF | F215C_PRG | 5 | 1.603504 | 0.032679 |
| 236 | 241 | LNISVF | F215C_PRG | 60 | 2.106580 | 0.058852 |
| 237 | 248 | NISVFPREVTNF | F215C | 0 | 0.000000 | 0.000000 |
| 237 | 248 | NISVFPREVTNF | F215C | 5 | 3.050890 | 0.090422 |
| 237 | 248 | NISVFPREVTNF | F215C | 60 | 3.498172 | 0.169142 |
| 237 | 248 | NISVFPREVTNF | F215C_PRG | 0 | 0.000000 | 0.000000 |
| 237 | 248 | NISVFPREVTNF | F215C_PRG | 5 | 3.477586 | 0.080233 |
| 237 | 248 | NISVFPREVTNF | F215C_PRG | 60 | 3.771030 | 0.130384 |
| 247 | 259 | NFLRKSVKRMKES | F215C | 0 | 0.000000 | 0.000000 |
| 247 | 259 | NFLRKSVKRMKES | F215C | 5 | 3.156661 | 0.017262 |
| 247 | 259 | NFLRKSVKRMKES | F215C | 60 | 4.014998 | 0.132633 |
| 247 | 259 | NFLRKSVKRMKES | F215C_PRG | 0 | 0.000000 | 0.000000 |
| 247 | 259 | NFLRKSVKRMKES | F215C_PRG | 5 | 3.102405 | 0.036997 |
| 247 | 259 | NFLRKSVKRMKES | F215C_PRG | 60 | 4.105653 | 0.076275 |

| | | | | | | |
|---|---|---|---|---|---|---|
| 249 | 261 | LRKSVKRMKESRL | F215C | 0 | 0.000000 | 0.000000 |
| 249 | 261 | LRKSVKRMKESRL | F215C | 5 | 1.584574 | 0.113040 |
| 249 | 261 | LRKSVKRMKESRL | F215C | 60 | 1.463973 | 0.154752 |
| 249 | 261 | LRKSVKRMKESRL | F215C_PRG | 0 | 0.000000 | 0.000000 |
| 249 | 261 | LRKSVKRMKESRL | F215C_PRG | 5 | 2.253215 | 0.104798 |
| 249 | 261 | LRKSVKRMKESRL | F215C_PRG | 60 | 2.194127 | 0.156353 |
| 262 | 271 | EDTQKHRVDF | F215C | 0 | 0.000000 | 0.000000 |
| 262 | 271 | EDTQKHRVDF | F215C | 5 | 0.947827 | 0.021595 |
| 262 | 271 | EDTQKHRVDF | F215C | 60 | 1.006938 | 0.091569 |
| 262 | 271 | EDTQKHRVDF | F215C_PRG | 0 | 0.000000 | 0.000000 |
| 262 | 271 | EDTQKHRVDF | F215C_PRG | 5 | 0.886863 | 0.029474 |
| 262 | 271 | EDTQKHRVDF | F215C_PRG | 60 | 0.973504 | 0.087130 |
| 277 | 292 | DSQNSKETESHKALSD | F215C | 0 | 0.000000 | 0.000000 |
| 277 | 292 | DSQNSKETESHKALSD | F215C | 5 | 2.629782 | 0.088426 |
| 277 | 292 | DSQNSKETESHKALSD | F215C | 60 | 2.305087 | 0.013189 |
| 277 | 292 | DSQNSKETESHKALSD | F215C_PRG | 0 | 0.000000 | 0.000000 |
| 277 | 292 | DSQNSKETESHKALSD | F215C_PRG | 5 | 3.027152 | 0.108880 |
| 277 | 292 | DSQNSKETESHKALSD | F215C_PRG | 60 | 2.880361 | 0.164885 |
| 277 | 294 | DSQNSKETESHKALSDLE | F215C | 0 | 0.000000 | 0.000000 |
| 277 | 294 | DSQNSKETESHKALSDLE | F215C | 5 | 2.418697 | 0.124561 |
| 277 | 294 | DSQNSKETESHKALSDLE | F215C | 60 | 2.367653 | 0.140933 |
| 277 | 294 | DSQNSKETESHKALSDLE | F215C_PRG | 0 | 0.000000 | 0.000000 |
| 277 | 294 | DSQNSKETESHKALSDLE | F215C_PRG | 5 | 2.827965 | 0.036898 |
| 277 | 294 | DSQNSKETESHKALSDLE | F215C_PRG | 60 | 2.797807 | 0.205426 |
| 296 | 302 | VAQSIIF | F215C | 0 | 0.000000 | 0.000000 |
| 296 | 302 | VAQSIIF | F215C | 5 | 0.039554 | 0.084572 |
| 296 | 302 | VAQSIIF | F215C | 60 | 0.490343 | 0.166591 |
| 296 | 302 | VAQSIIF | F215C_PRG | 0 | 0.000000 | 0.000000 |
| 296 | 302 | VAQSIIF | F215C_PRG | 5 | 0.099300 | 0.062372 |
| 296 | 302 | VAQSIIF | F215C_PRG | 60 | 0.407673 | 0.086905 |
| 301 | 306 | IFIFAG | F215C | 0 | 0.000000 | 0.000000 |
| 301 | 306 | IFIFAG | F215C | 5 | 0.168071 | 0.005883 |
| 301 | 306 | IFIFAG | F215C | 60 | 0.386783 | 0.017392 |
| 301 | 306 | IFIFAG | F215C_PRG | 0 | 0.000000 | 0.000000 |
| 301 | 306 | IFIFAG | F215C_PRG | 5 | 0.194857 | 0.022007 |
| 301 | 306 | IFIFAG | F215C_PRG | 60 | 0.320758 | 0.020699 |
| 307 | 312 | YETTSS | F215C | 0 | 0.000000 | 0.000000 |
| 307 | 312 | YETTSS | F215C | 5 | 2.781619 | 0.031307 |
| 307 | 312 | YETTSS | F215C | 60 | 2.924912 | 0.074875 |
| 307 | 312 | YETTSS | F215C_PRG | 0 | 0.000000 | 0.000000 |
| 307 | 312 | YETTSS | F215C_PRG | 5 | 2.803427 | 0.030698 |
| 307 | 312 | YETTSS | F215C_PRG | 60 | 2.866574 | 0.036035 |
| 307 | 314 | YETTSSVL | F215C | 0 | 0.000000 | 0.000000 |
| 307 | 314 | YETTSSVL | F215C | 5 | 0.549092 | 0.034119 |
| 307 | 314 | YETTSSVL | F215C | 60 | 0.787537 | 0.110334 |

| | | | | | | |
|---|---|---|---|---|---|---|
| 307 | 314 | YETTSSVL | F215C_PRG | 0 | 0.000000 | 0.000000 |
| 307 | 314 | YETTSSVL | F215C_PRG | 5 | 0.867717 | 0.040379 |
| 307 | 314 | YETTSSVL | F215C_PRG | 60 | 1.265972 | 0.025275 |
| 307 | 316 | YETTSSVLSF | F215C | 0 | 0.000000 | 0.000000 |
| 307 | 316 | YETTSSVLSF | F215C | 5 | 0.633763 | 0.021465 |
| 307 | 316 | YETTSSVLSF | F215C | 60 | 1.089605 | 0.063958 |
| 307 | 316 | YETTSSVLSF | F215C_PRG | 0 | 0.000000 | 0.000000 |
| 307 | 316 | YETTSSVLSF | F215C_PRG | 5 | 0.910439 | 0.035823 |
| 307 | 316 | YETTSSVLSF | F215C_PRG | 60 | 1.365145 | 0.032268 |
| 319 | 333 | YELATHPDVQQKLQE | F215C | 0 | 0.000000 | 0.000000 |
| 319 | 333 | YELATHPDVQQKLQE | F215C | 5 | 0.568451 | 0.028625 |
| 319 | 333 | YELATHPDVQQKLQE | F215C | 60 | 1.004232 | 0.086670 |
| 319 | 333 | YELATHPDVQQKLQE | F215C_PRG | 0 | 0.000000 | 0.000000 |
| 319 | 333 | YELATHPDVQQKLQE | F215C_PRG | 5 | 0.897497 | 0.051187 |
| 319 | 333 | YELATHPDVQQKLQE | F215C_PRG | 60 | 1.123165 | 0.048156 |
| 320 | 333 | ELATHPDVQQKLQE | F215C | 0 | 0.000000 | 0.000000 |
| 320 | 333 | ELATHPDVQQKLQE | F215C | 5 | 0.441676 | 0.093651 |
| 320 | 333 | ELATHPDVQQKLQE | F215C | 60 | 0.951984 | 0.074077 |
| 320 | 333 | ELATHPDVQQKLQE | F215C_PRG | 0 | 0.000000 | 0.000000 |
| 320 | 333 | ELATHPDVQQKLQE | F215C_PRG | 5 | 0.574124 | 0.080720 |
| 320 | 333 | ELATHPDVQQKLQE | F215C_PRG | 60 | 1.110045 | 0.098352 |
| 321 | 333 | LATHPDVQQKLQE | F215C | 0 | 0.000000 | 0.000000 |
| 321 | 333 | LATHPDVQQKLQE | F215C | 5 | 0.406908 | 0.096541 |
| 321 | 333 | LATHPDVQQKLQE | F215C | 60 | 0.981392 | 0.070254 |
| 321 | 333 | LATHPDVQQKLQE | F215C_PRG | 0 | 0.000000 | 0.000000 |
| 321 | 333 | LATHPDVQQKLQE | F215C_PRG | 5 | 0.376190 | 0.106716 |
| 321 | 333 | LATHPDVQQKLQE | F215C_PRG | 60 | 0.906891 | 0.031226 |
| 332 | 337 | QEEIDA | F215C | 0 | 0.000000 | 0.000000 |
| 332 | 337 | QEEIDA | F215C | 5 | 0.463965 | 0.017732 |
| 332 | 337 | QEEIDA | F215C | 60 | 1.032951 | 0.037855 |
| 332 | 337 | QEEIDA | F215C_PRG | 0 | 0.000000 | 0.000000 |
| 332 | 337 | QEEIDA | F215C_PRG | 5 | 0.473425 | 0.024297 |
| 332 | 337 | QEEIDA | F215C_PRG | 60 | 1.110039 | 0.045054 |
| 337 | 349 | AVLPNKAPPTYDT | F215C | 0 | 0.000000 | 0.000000 |
| 337 | 349 | AVLPNKAPPTYDT | F215C | 5 | 2.693862 | 0.076806 |
| 337 | 349 | AVLPNKAPPTYDT | F215C | 60 | 3.088066 | 0.184297 |
| 337 | 349 | AVLPNKAPPTYDT | F215C_PRG | 0 | 0.000000 | 0.000000 |
| 337 | 349 | AVLPNKAPPTYDT | F215C_PRG | 5 | 2.862374 | 0.014831 |
| 337 | 349 | AVLPNKAPPTYDT | F215C_PRG | 60 | 3.230834 | 0.180446 |
| 340 | 347 | PNKAPPTY | F215C | 0 | 0.000000 | 0.000000 |
| 340 | 347 | PNKAPPTY | F215C | 5 | 1.395025 | 0.017304 |
| 340 | 347 | PNKAPPTY | F215C | 60 | 1.772481 | 0.071161 |
| 340 | 347 | PNKAPPTY | F215C_PRG | 0 | 0.000000 | 0.000000 |
| 340 | 347 | PNKAPPTY | F215C_PRG | 5 | 1.484500 | 0.059945 |
| 340 | 347 | PNKAPPTY | F215C_PRG | 60 | 1.693179 | 0.119994 |

| | | | | | | |
|---|---|---|---|---|---|---|
| 353 | 358 | MEYLDM | F215C | 0 | 0.000000 | 0.000000 |
| 353 | 358 | MEYLDM | F215C | 5 | 0.385630 | 0.064946 |
| 353 | 358 | MEYLDM | F215C | 60 | 0.599775 | 0.108134 |
| 353 | 358 | MEYLDM | F215C_PRG | 0 | 0.000000 | 0.000000 |
| 353 | 358 | MEYLDM | F215C_PRG | 5 | 0.328755 | 0.028337 |
| 353 | 358 | MEYLDM | F215C_PRG | 60 | 0.438871 | 0.047295 |
| 357 | 363 | DMVVNET | F215C | 0 | 0.000000 | 0.000000 |
| 357 | 363 | DMVVNET | F215C | 5 | 0.036752 | 0.084500 |
| 357 | 363 | DMVVNET | F215C | 60 | 0.082981 | 0.041302 |
| 357 | 363 | DMVVNET | F215C_PRG | 0 | 0.000000 | 0.000000 |
| 357 | 363 | DMVVNET | F215C_PRG | 5 | -0.015312 | 0.028978 |
| 357 | 363 | DMVVNET | F215C_PRG | 60 | -0.029862 | 0.023835 |
| 359 | 366 | VVNETLRL | F215C | 0 | 0.000000 | 0.000000 |
| 359 | 366 | VVNETLRL | F215C | 5 | -0.005319 | 0.072286 |
| 359 | 366 | VVNETLRL | F215C | 60 | 0.094108 | 0.027491 |
| 359 | 366 | VVNETLRL | F215C_PRG | 0 | 0.000000 | 0.000000 |
| 359 | 366 | VVNETLRL | F215C_PRG | 5 | 0.126053 | 0.031636 |
| 359 | 366 | VVNETLRL | F215C_PRG | 60 | 0.252551 | 0.045839 |
| 363 | 370 | TLRLFPIA | F215C | 0 | 0.000000 | 0.000000 |
| 363 | 370 | TLRLFPIA | F215C | 5 | 0.187760 | 0.046126 |
| 363 | 370 | TLRLFPIA | F215C | 60 | 0.465388 | 0.058600 |
| 363 | 370 | TLRLFPIA | F215C_PRG | 0 | 0.000000 | 0.000000 |
| 363 | 370 | TLRLFPIA | F215C_PRG | 5 | 0.200845 | 0.047253 |
| 363 | 370 | TLRLFPIA | F215C_PRG | 60 | 0.377016 | 0.086487 |
| 364 | 371 | LRLFPIAM | F215C | 0 | 0.000000 | 0.000000 |
| 364 | 371 | LRLFPIAM | F215C | 5 | 0.336235 | 0.017197 |
| 364 | 371 | LRLFPIAM | F215C | 60 | 0.467559 | 0.010396 |
| 364 | 371 | LRLFPIAM | F215C_PRG | 0 | 0.000000 | 0.000000 |
| 364 | 371 | LRLFPIAM | F215C_PRG | 5 | 0.694934 | 0.121471 |
| 364 | 371 | LRLFPIAM | F215C_PRG | 60 | 0.861601 | 0.067634 |
| 374 | 385 | ERVCKKDVEING | F215C | 0 | 0.000000 | 0.000000 |
| 374 | 385 | ERVCKKDVEING | F215C | 5 | 1.398765 | 0.022442 |
| 374 | 385 | ERVCKKDVEING | F215C | 60 | 1.481605 | 0.104297 |
| 374 | 385 | ERVCKKDVEING | F215C_PRG | 0 | 0.000000 | 0.000000 |
| 374 | 385 | ERVCKKDVEING | F215C_PRG | 5 | 1.355309 | 0.111353 |
| 374 | 385 | ERVCKKDVEING | F215C_PRG | 60 | 1.433179 | 0.131956 |
| 387 | 393 | FIPKGVV | F215C | 0 | 0.000000 | 0.000000 |
| 387 | 393 | FIPKGVV | F215C | 5 | 0.522279 | 0.026972 |
| 387 | 393 | FIPKGVV | F215C | 60 | 0.635509 | 0.093228 |
| 387 | 393 | FIPKGVV | F215C_PRG | 0 | 0.000000 | 0.000000 |
| 387 | 393 | FIPKGVV | F215C_PRG | 5 | 0.681164 | 0.028546 |
| 387 | 393 | FIPKGVV | F215C_PRG | 60 | 1.191873 | 0.026993 |
| 388 | 393 | IPKGVV | F215C | 0 | 0.000000 | 0.000000 |
| 388 | 393 | IPKGVV | F215C | 5 | 0.437450 | 0.017051 |
| 388 | 393 | IPKGVV | F215C | 60 | 0.662637 | 0.060352 |

| | | | | | | |
|---|---|---|---|---|---|---|
| 388 | 393 | IPKGVV | F215C_PRG | 0 | 0.000000 | 0.000000 |
| 388 | 393 | IPKGVV | F215C_PRG | 5 | 0.349999 | 0.024741 |
| 388 | 393 | IPKGVV | F215C_PRG | 60 | 0.578506 | 0.031708 |
| 388 | 395 | IPKGVVVM | F215C | 0 | 0.000000 | 0.000000 |
| 388 | 395 | IPKGVVVM | F215C | 5 | 0.754749 | 0.074981 |
| 388 | 395 | IPKGVVVM | F215C | 60 | 1.129466 | 0.044802 |
| 388 | 395 | IPKGVVVM | F215C_PRG | 0 | 0.000000 | 0.000000 |
| 388 | 395 | IPKGVVVM | F215C_PRG | 5 | 0.901681 | 0.033178 |
| 388 | 395 | IPKGVVVM | F215C_PRG | 60 | 1.244862 | 0.072218 |
| 394 | 399 | VMIPSY | F215C | 0 | 0.000000 | 0.000000 |
| 394 | 399 | VMIPSY | F215C | 5 | 0.355260 | 0.041496 |
| 394 | 399 | VMIPSY | F215C | 60 | 0.496100 | 0.055969 |
| 394 | 399 | VMIPSY | F215C_PRG | 0 | 0.000000 | 0.000000 |
| 394 | 399 | VMIPSY | F215C_PRG | 5 | 0.138413 | 0.032152 |
| 394 | 399 | VMIPSY | F215C_PRG | 60 | 0.269325 | 0.063071 |
| 394 | 401 | VMIPSYAL | F215C | 0 | 0.000000 | 0.000000 |
| 394 | 401 | VMIPSYAL | F215C | 5 | 0.710683 | 0.097518 |
| 394 | 401 | VMIPSYAL | F215C | 60 | 0.970712 | 0.115180 |
| 394 | 401 | VMIPSYAL | F215C_PRG | 0 | 0.000000 | 0.000000 |
| 394 | 401 | VMIPSYAL | F215C_PRG | 5 | 0.586273 | 0.047934 |
| 394 | 401 | VMIPSYAL | F215C_PRG | 60 | 0.679881 | 0.072788 |
| 396 | 401 | IPSYAL | F215C | 0 | 0.000000 | 0.000000 |
| 396 | 401 | IPSYAL | F215C | 5 | 0.615062 | 0.061245 |
| 396 | 401 | IPSYAL | F215C | 60 | 0.778139 | 0.087799 |
| 396 | 401 | IPSYAL | F215C_PRG | 0 | 0.000000 | 0.000000 |
| 396 | 401 | IPSYAL | F215C_PRG | 5 | 0.615551 | 0.043980 |
| 396 | 401 | IPSYAL | F215C_PRG | 60 | 0.707276 | 0.042260 |
| 400 | 407 | ALHRDPKY | F215C | 0 | 0.000000 | 0.000000 |
| 400 | 407 | ALHRDPKY | F215C | 5 | 0.623924 | 0.015709 |
| 400 | 407 | ALHRDPKY | F215C | 60 | 0.780468 | 0.064645 |
| 400 | 407 | ALHRDPKY | F215C_PRG | 0 | 0.000000 | 0.000000 |
| 400 | 407 | ALHRDPKY | F215C_PRG | 5 | 0.478702 | 0.011112 |
| 400 | 407 | ALHRDPKY | F215C_PRG | 60 | 0.530480 | 0.023746 |
| 402 | 407 | HRDPKY | F215C | 0 | 0.000000 | 0.000000 |
| 402 | 407 | HRDPKY | F215C | 5 | 0.727616 | 0.046015 |
| 402 | 407 | HRDPKY | F215C | 60 | 0.848610 | 0.045762 |
| 402 | 407 | HRDPKY | F215C_PRG | 0 | 0.000000 | 0.000000 |
| 402 | 407 | HRDPKY | F215C_PRG | 5 | 0.543412 | 0.028958 |
| 402 | 407 | HRDPKY | F215C_PRG | 60 | 0.639353 | 0.068386 |
| 408 | 414 | WTEPEKF | F215C | 0 | 0.000000 | 0.000000 |
| 408 | 414 | WTEPEKF | F215C | 5 | 1.285222 | 0.084259 |
| 408 | 414 | WTEPEKF | F215C | 60 | 1.159435 | 0.062964 |
| 408 | 414 | WTEPEKF | F215C_PRG | 0 | 0.000000 | 0.000000 |
| 408 | 414 | WTEPEKF | F215C_PRG | 5 | 1.230232 | 0.022616 |
| 408 | 414 | WTEPEKF | F215C_PRG | 60 | 1.154018 | 0.038421 |

| | | | | | | |
|---|---|---|---|---|---|---|
| 415 | 428 | LPERFSKKNKDNID | F215C | 0 | 0.000000 | 0.000000 |
| 415 | 428 | LPERFSKKNKDNID | F215C | 5 | 1.531150 | 0.070077 |
| 415 | 428 | LPERFSKKNKDNID | F215C | 60 | 1.356475 | 0.020320 |
| 415 | 428 | LPERFSKKNKDNID | F215C_PRG | 0 | 0.000000 | 0.000000 |
| 415 | 428 | LPERFSKKNKDNID | F215C_PRG | 5 | 1.194042 | 0.072342 |
| 415 | 428 | LPERFSKKNKDNID | F215C_PRG | 60 | 1.115721 | 0.035732 |
| 420 | 444 | SKKNKDNIDPYIYTPFGSGPRNCIG | F215C | 0 | 0.000000 | 0.000000 |
| 420 | 444 | SKKNKDNIDPYIYTPFGSGPRNCIG | F215C | 5 | 3.282658 | 0.039254 |
| 420 | 444 | SKKNKDNIDPYIYTPFGSGPRNCIG | F215C | 60 | 3.896878 | 0.052153 |
| 420 | 444 | SKKNKDNIDPYIYTPFGSGPRNCIG | F215C_PRG | 0 | 0.000000 | 0.000000 |
| 420 | 444 | SKKNKDNIDPYIYTPFGSGPRNCIG | F215C_PRG | 5 | 3.040233 | 0.055464 |
| 420 | 444 | SKKNKDNIDPYIYTPFGSGPRNCIG | F215C_PRG | 60 | 3.784681 | 0.165788 |
| 446 | 452 | RFALMNM | F215C | 0 | 0.000000 | 0.000000 |
| 446 | 452 | RFALMNM | F215C | 5 | 0.200274 | 0.051676 |
| 446 | 452 | RFALMNM | F215C | 60 | 0.343515 | 0.042684 |
| 446 | 452 | RFALMNM | F215C_PRG | 0 | 0.000000 | 0.000000 |
| 446 | 452 | RFALMNM | F215C_PRG | 5 | 0.179710 | 0.020803 |
| 446 | 452 | RFALMNM | F215C_PRG | 60 | 0.348818 | 0.019831 |
| 455 | 463 | ALIRVLQNF | F215C | 0 | 0.000000 | 0.000000 |
| 455 | 463 | ALIRVLQNF | F215C | 5 | 0.058376 | 0.036627 |
| 455 | 463 | ALIRVLQNF | F215C | 60 | 0.301026 | 0.078430 |
| 455 | 463 | ALIRVLQNF | F215C_PRG | 0 | 0.000000 | 0.000000 |
| 455 | 463 | ALIRVLQNF | F215C_PRG | 5 | 0.171926 | 0.026435 |
| 455 | 463 | ALIRVLQNF | F215C_PRG | 60 | 0.418468 | 0.026005 |
| 457 | 463 | IRVLQNF | F215C | 0 | 0.000000 | 0.000000 |
| 457 | 463 | IRVLQNF | F215C | 5 | 0.188360 | 0.021824 |
| 457 | 463 | IRVLQNF | F215C | 60 | 0.335792 | 0.041144 |
| 457 | 463 | IRVLQNF | F215C_PRG | 0 | 0.000000 | 0.000000 |
| 457 | 463 | IRVLQNF | F215C_PRG | 5 | 0.160174 | 0.025782 |
| 457 | 463 | IRVLQNF | F215C_PRG | 60 | 0.344007 | 0.028685 |
| 457 | 477 | IRVLQNFSFKPGKETQIPLKL | F215C | 0 | 0.000000 | 0.000000 |
| 457 | 477 | IRVLQNFSFKPGKETQIPLKL | F215C | 5 | 3.811528 | 0.106346 |
| 457 | 477 | IRVLQNFSFKPGKETQIPLKL | F215C | 60 | 3.663356 | 0.190428 |
| 457 | 477 | IRVLQNFSFKPGKETQIPLKL | F215C_PRG | 0 | 0.000000 | 0.000000 |
| 457 | 477 | IRVLQNFSFKPGKETQIPLKL | F215C_PRG | 5 | 3.720469 | 0.024564 |
| 457 | 477 | IRVLQNFSFKPGKETQIPLKL | F215C_PRG | 60 | 3.937125 | 0.169148 |
| 464 | 479 | SFKPGKETQIPLKLSL | F215C | 0 | 0.000000 | 0.000000 |
| 464 | 479 | SFKPGKETQIPLKLSL | F215C | 5 | 4.256176 | 0.042782 |
| 464 | 479 | SFKPGKETQIPLKLSL | F215C | 60 | 4.072812 | 0.163264 |
| 464 | 479 | SFKPGKETQIPLKLSL | F215C_PRG | 0 | 0.000000 | 0.000000 |
| 464 | 479 | SFKPGKETQIPLKLSL | F215C_PRG | 5 | 4.225589 | 0.105149 |
| 464 | 479 | SFKPGKETQIPLKLSL | F215C_PRG | 60 | 4.134392 | 0.171189 |
| 478 | 490 | SLGGLLQPEKPVV | F215C | 0 | 0.000000 | 0.000000 |
| 478 | 490 | SLGGLLQPEKPVV | F215C | 5 | 1.436344 | 0.028368 |
| 478 | 490 | SLGGLLQPEKPVV | F215C | 60 | 1.996869 | 0.098422 |

| | | | | | | |
|---|---|---|---|---|---|---|
| 478 | 490 | SLGGLLQPEKPVV | F215C_PRG | 0 | 0.000000 | 0.000000 |
| 478 | 490 | SLGGLLQPEKPVV | F215C_PRG | 5 | 1.934903 | 0.013951 |
| 478 | 490 | SLGGLLQPEKPVV | F215C_PRG | 60 | 2.521507 | 0.056561 |
| 492 | 507 | KVESRDGTVSGAHHHH | F215C | 0 | 0.000000 | 0.000000 |
| 492 | 507 | KVESRDGTVSGAHHHH | F215C | 5 | 1.106714 | 0.120805 |
| 492 | 507 | KVESRDGTVSGAHHHH | F215C | 60 | 1.073686 | 0.171651 |
| 492 | 507 | KVESRDGTVSGAHHHH | F215C_PRG | 0 | 0.000000 | 0.000000 |
| 492 | 507 | KVESRDGTVSGAHHHH | F215C_PRG | 5 | 1.095793 | 0.061491 |
| 492 | 507 | KVESRDGTVSGAHHHH | F215C_PRG | 60 | 1.052724 | 0.093283 |
| 494 | 507 | ESRDGTVSGAHHHH | F215C | 0 | 0.000000 | 0.000000 |
| 494 | 507 | ESRDGTVSGAHHHH | F215C | 5 | 0.855359 | 0.166710 |
| 494 | 507 | ESRDGTVSGAHHHH | F215C | 60 | 0.885327 | 0.182624 |
| 494 | 507 | ESRDGTVSGAHHHH | F215C_PRG | 0 | 0.000000 | 0.000000 |
| 494 | 507 | ESRDGTVSGAHHHH | F215C_PRG | 5 | 1.428377 | 0.096135 |
| 494 | 507 | ESRDGTVSGAHHHH | F215C_PRG | 60 | 1.452671 | 0.120566 |

| Deuterium uptake data for WT CYP3A4 state | | | | | | |
|---|---|---|---|---|---|---|
| **Start** | **End** | **Sequence** | **State** | **Exposure (min)** | **Uptake (Da)** | **Uptake SD (Da)** |
| 34 | 47 | KKLGIPGPTPLPFL | WT | 0 | 0 | 0 |
| 34 | 47 | KKLGIPGPTPLPFL | WT | 0.5 | 0.288361 | 0.024592 |
| 34 | 47 | KKLGIPGPTPLPFL | WT | 1 | 0.276461 | 0.035186 |
| 34 | 47 | KKLGIPGPTPLPFL | WT | 2 | 0.319475 | 0.037868 |
| 34 | 47 | KKLGIPGPTPLPFL | WT | 5 | 0.238272 | 0.071612 |
| 34 | 47 | KKLGIPGPTPLPFL | WT | 30 | 0.272124 | 0.024427 |
| 34 | 47 | KKLGIPGPTPLPFL | WT | 60 | 0.200194 | 0.029311 |
| 34 | 47 | KKLGIPGPTPLPFL | WT | 240 | 0.350254 | 0.063503 |
| 34 | 48 | KKLGIPGPTPLPFLG | WT | 0 | 0 | 0 |
| 34 | 48 | KKLGIPGPTPLPFLG | WT | 0.5 | 0.312453 | 0.081494 |
| 34 | 48 | KKLGIPGPTPLPFLG | WT | 1 | 0.44931 | 0.042451 |
| 34 | 48 | KKLGIPGPTPLPFLG | WT | 2 | 0.442249 | 0.050705 |
| 34 | 48 | KKLGIPGPTPLPFLG | WT | 5 | 0.475221 | 0.01676 |
| 34 | 48 | KKLGIPGPTPLPFLG | WT | 30 | 0.403124 | 0.077327 |
| 34 | 48 | KKLGIPGPTPLPFLG | WT | 60 | 0.442278 | 0.011946 |
| 34 | 48 | KKLGIPGPTPLPFLG | WT | 240 | 0.387806 | 0.029124 |
| 34 | 51 | KKLGIPGPTPLPFLGNIL | WT | 0 | 0 | 0 |
| 34 | 51 | KKLGIPGPTPLPFLGNIL | WT | 0.5 | 0.532055 | 0.084345 |
| 34 | 51 | KKLGIPGPTPLPFLGNIL | WT | 1 | 0.711536 | 0.143005 |
| 34 | 51 | KKLGIPGPTPLPFLGNIL | WT | 2 | 0.653493 | 0.056847 |
| 34 | 51 | KKLGIPGPTPLPFLGNIL | WT | 5 | 0.730358 | 0.033283 |
| 34 | 51 | KKLGIPGPTPLPFLGNIL | WT | 30 | 0.7745 | 0.116521 |
| 34 | 51 | KKLGIPGPTPLPFLGNIL | WT | 60 | 0.898732 | 0.031169 |
| 34 | 51 | KKLGIPGPTPLPFLGNIL | WT | 240 | 0.909372 | 0.134814 |
| 52 | 59 | SYHKGFCM | WT | 0 | 0 | 0 |
| 52 | 59 | SYHKGFCM | WT | 0.5 | 0.919746 | 0.023313 |

| | | | | | | |
|---|---|---|---|---|---|---|
| 52 | 59 | SYHKGFCM | WT | 1 | 0.983218 | 0.052707 |
| 52 | 59 | SYHKGFCM | WT | 2 | 1.049571 | 0.039628 |
| 52 | 59 | SYHKGFCM | WT | 5 | 0.956794 | 0.053044 |
| 52 | 59 | SYHKGFCM | WT | 30 | 1.044739 | 0.025474 |
| 52 | 59 | SYHKGFCM | WT | 60 | 1.002328 | 0.0219 |
| 52 | 59 | SYHKGFCM | WT | 240 | 1.005325 | 0.034601 |
| 52 | 60 | SYHKGFCMF | WT | 0 | 0 | 0 |
| 52 | 60 | SYHKGFCMF | WT | 0.5 | 0.876167 | 0.074654 |
| 52 | 60 | SYHKGFCMF | WT | 1 | 1.059193 | 0.087479 |
| 52 | 60 | SYHKGFCMF | WT | 2 | 1.184167 | 0.042798 |
| 52 | 60 | SYHKGFCMF | WT | 5 | 1.221389 | 0.072126 |
| 52 | 60 | SYHKGFCMF | WT | 30 | 1.303322 | 0.024208 |
| 52 | 60 | SYHKGFCMF | WT | 60 | 1.259056 | 0.071398 |
| 52 | 60 | SYHKGFCMF | WT | 240 | 1.273114 | 0.060733 |
| 60 | 74 | FDMECHKKYGKVWGF | WT | 0 | 0 | 0 |
| 60 | 74 | FDMECHKKYGKVWGF | WT | 0.5 | 0.572513 | 0.129615 |
| 60 | 74 | FDMECHKKYGKVWGF | WT | 1 | 0.626859 | 0.094879 |
| 60 | 74 | FDMECHKKYGKVWGF | WT | 2 | 0.664525 | 0.096614 |
| 60 | 74 | FDMECHKKYGKVWGF | WT | 5 | 0.72778 | 0.089467 |
| 60 | 74 | FDMECHKKYGKVWGF | WT | 30 | 1.00109 | 0.092726 |
| 60 | 74 | FDMECHKKYGKVWGF | WT | 60 | 1.179976 | 0.173048 |
| 60 | 74 | FDMECHKKYGKVWGF | WT | 240 | 1.383232 | 0.095323 |
| 61 | 74 | DMECHKKYGKVWGF | WT | 0 | 0 | 0 |
| 61 | 74 | DMECHKKYGKVWGF | WT | 0.5 | 0.567141 | 0.05171 |
| 61 | 74 | DMECHKKYGKVWGF | WT | 1 | 0.618468 | 0.034381 |
| 61 | 74 | DMECHKKYGKVWGF | WT | 2 | 0.684872 | 0.074614 |
| 61 | 74 | DMECHKKYGKVWGF | WT | 5 | 0.765599 | 0.038087 |
| 61 | 74 | DMECHKKYGKVWGF | WT | 30 | 1.038159 | 0.025798 |
| 61 | 74 | DMECHKKYGKVWGF | WT | 60 | 1.166808 | 0.097469 |
| 61 | 74 | DMECHKKYGKVWGF | WT | 240 | 1.39977 | 0.076995 |
| 74 | 82 | FYDGQQPVL | WT | 0 | 0 | 0 |
| 74 | 82 | FYDGQQPVL | WT | 0.5 | 0.577375 | 0.021126 |
| 74 | 82 | FYDGQQPVL | WT | 1 | 0.688014 | 0.040329 |
| 74 | 82 | FYDGQQPVL | WT | 2 | 0.78572 | 0.046344 |
| 74 | 82 | FYDGQQPVL | WT | 5 | 0.925278 | 0.039412 |
| 74 | 82 | FYDGQQPVL | WT | 30 | 1.223203 | 0.021467 |
| 74 | 82 | FYDGQQPVL | WT | 60 | 1.315547 | 0.052245 |
| 74 | 82 | FYDGQQPVL | WT | 240 | 1.482378 | 0.02848 |
| 76 | 82 | DGQQPVL | WT | 0 | 0 | 0 |
| 76 | 82 | DGQQPVL | WT | 0.5 | 0.363817 | 0.064968 |
| 76 | 82 | DGQQPVL | WT | 1 | 0.498405 | 0.053894 |
| 76 | 82 | DGQQPVL | WT | 2 | 0.550572 | 0.065643 |
| 76 | 82 | DGQQPVL | WT | 5 | 0.668773 | 0.062943 |
| 76 | 82 | DGQQPVL | WT | 30 | 0.843408 | 0.046317 |
| 76 | 82 | DGQQPVL | WT | 60 | 0.917371 | 0.068118 |

| | | | | | | |
|---|---|---|---|---|---|---|
| 76 | 82 | DGQQPVL | WT | 240 | 0.991597 | 0.091932 |
| 83 | 94 | AITDPDMIKTVL | WT | 0 | 0 | 0 |
| 83 | 94 | AITDPDMIKTVL | WT | 0.5 | 0.457691 | 0.056592 |
| 83 | 94 | AITDPDMIKTVL | WT | 1 | 0.542949 | 0.038076 |
| 83 | 94 | AITDPDMIKTVL | WT | 2 | 0.702472 | 0.053176 |
| 83 | 94 | AITDPDMIKTVL | WT | 5 | 0.897056 | 0.014248 |
| 83 | 94 | AITDPDMIKTVL | WT | 30 | 1.501599 | 0.099249 |
| 83 | 94 | AITDPDMIKTVL | WT | 60 | 1.659104 | 0.092282 |
| 83 | 94 | AITDPDMIKTVL | WT | 240 | 1.933393 | 0.053151 |
| 84 | 89 | ITDPDM | WT | 0 | 0 | 0 |
| 84 | 89 | ITDPDM | WT | 0.5 | 0.320079 | 0.012852 |
| 84 | 89 | ITDPDM | WT | 1 | 0.367042 | 0.037531 |
| 84 | 89 | ITDPDM | WT | 2 | 0.413294 | 0.019549 |
| 84 | 89 | ITDPDM | WT | 5 | 0.682139 | 0.071642 |
| 84 | 89 | ITDPDM | WT | 30 | 0.866915 | 0.030368 |
| 84 | 89 | ITDPDM | WT | 60 | 0.849849 | 0.030049 |
| 84 | 89 | ITDPDM | WT | 240 | 0.834451 | 0.037427 |
| 93 | 99 | VLVKECY | WT | 0 | 0 | 0 |
| 93 | 99 | VLVKECY | WT | 0.5 | 1.210229 | 0.118992 |
| 93 | 99 | VLVKECY | WT | 1 | 1.404369 | 0.057059 |
| 93 | 99 | VLVKECY | WT | 2 | 1.560476 | 0.103176 |
| 93 | 99 | VLVKECY | WT | 5 | 1.570552 | 0.128236 |
| 93 | 99 | VLVKECY | WT | 30 | 1.748168 | 0.068866 |
| 93 | 99 | VLVKECY | WT | 60 | 1.782742 | 0.076442 |
| 93 | 99 | VLVKECY | WT | 240 | 1.983904 | 0.06958 |
| 100 | 113 | SVFTNRRPFGPVGF | WT | 0 | 0 | 0 |
| 100 | 113 | SVFTNRRPFGPVGF | WT | 0.5 | 2.195114 | 0.044316 |
| 100 | 113 | SVFTNRRPFGPVGF | WT | 1 | 2.413714 | 0.126731 |
| 100 | 113 | SVFTNRRPFGPVGF | WT | 2 | 2.677608 | 0.028719 |
| 100 | 113 | SVFTNRRPFGPVGF | WT | 5 | 2.838795 | 0.066887 |
| 100 | 113 | SVFTNRRPFGPVGF | WT | 30 | 3.311278 | 0.039354 |
| 100 | 113 | SVFTNRRPFGPVGF | WT | 60 | 3.330114 | 0.105028 |
| 100 | 113 | SVFTNRRPFGPVGF | WT | 240 | 3.452719 | 0.106605 |
| 102 | 113 | FTNRRPFGPVGF | WT | 0 | 0 | 0 |
| 102 | 113 | FTNRRPFGPVGF | WT | 0.5 | 2.418405 | 0.011388 |
| 102 | 113 | FTNRRPFGPVGF | WT | 1 | 2.694505 | 0.029836 |
| 102 | 113 | FTNRRPFGPVGF | WT | 2 | 2.895016 | 0.070771 |
| 102 | 113 | FTNRRPFGPVGF | WT | 5 | 2.898015 | 0.042283 |
| 102 | 113 | FTNRRPFGPVGF | WT | 30 | 3.195503 | 0.071873 |
| 102 | 113 | FTNRRPFGPVGF | WT | 60 | 3.214159 | 0.051415 |
| 102 | 113 | FTNRRPFGPVGF | WT | 240 | 3.291569 | 0.040305 |
| 103 | 113 | TNRRPFGPVGF | WT | 0 | 0 | 0 |
| 103 | 113 | TNRRPFGPVGF | WT | 0.5 | 2.464388 | 0.018901 |
| 103 | 113 | TNRRPFGPVGF | WT | 1 | 2.641261 | 0.052197 |
| 103 | 113 | TNRRPFGPVGF | WT | 2 | 2.912928 | 0.045039 |

| | | | | | | |
|---|---|---|---|---|---|---|
| 103 | 113 | TNRRPFGPVGF | WT | 5 | 2.838226 | 0.065451 |
| 103 | 113 | TNRRPFGPVGF | WT | 30 | 3.093689 | 0.015301 |
| 103 | 113 | TNRRPFGPVGF | WT | 60 | 3.147605 | 0.068837 |
| 103 | 113 | TNRRPFGPVGF | WT | 240 | 3.213648 | 0.010553 |
| 114 | 122 | MKSAISIAE | WT | 0 | 0 | 0 |
| 114 | 122 | MKSAISIAE | WT | 0.5 | 1.182253 | 0.029977 |
| 114 | 122 | MKSAISIAE | WT | 1 | 1.258464 | 0.076312 |
| 114 | 122 | MKSAISIAE | WT | 2 | 1.381221 | 0.045277 |
| 114 | 122 | MKSAISIAE | WT | 5 | 1.533896 | 0.032093 |
| 114 | 122 | MKSAISIAE | WT | 30 | 2.113082 | 0.053667 |
| 114 | 122 | MKSAISIAE | WT | 60 | 2.372796 | 0.030456 |
| 114 | 122 | MKSAISIAE | WT | 240 | 2.971892 | 0.028425 |
| 114 | 133 | MKSAISIAEDEEWKRLRSLL | WT | 0 | 0 | 0 |
| 114 | 133 | MKSAISIAEDEEWKRLRSLL | WT | 0.5 | 1.570982 | 0.168056 |
| 114 | 133 | MKSAISIAEDEEWKRLRSLL | WT | 1 | 1.790939 | 0.113127 |
| 114 | 133 | MKSAISIAEDEEWKRLRSLL | WT | 2 | 1.991333 | 0.10015 |
| 114 | 133 | MKSAISIAEDEEWKRLRSLL | WT | 5 | 2.553297 | 0.174518 |
| 114 | 133 | MKSAISIAEDEEWKRLRSLL | WT | 30 | 3.878723 | 0.063705 |
| 114 | 133 | MKSAISIAEDEEWKRLRSLL | WT | 60 | 4.462837 | 0.1125 |
| 114 | 133 | MKSAISIAEDEEWKRLRSLL | WT | 240 | 6.501147 | 0.170018 |
| 120 | 125 | IAEDEE | WT | 0 | 0 | 0 |
| 120 | 125 | IAEDEE | WT | 0.5 | 0.918455 | 0.018625 |
| 120 | 125 | IAEDEE | WT | 1 | 0.9815 | 0.031626 |
| 120 | 125 | IAEDEE | WT | 2 | 1.075203 | 0.038781 |
| 120 | 125 | IAEDEE | WT | 5 | 1.138125 | 0.009724 |
| 120 | 125 | IAEDEE | WT | 30 | 1.331157 | 0.038856 |
| 120 | 125 | IAEDEE | WT | 60 | 1.326413 | 0.040076 |
| 120 | 125 | IAEDEE | WT | 240 | 1.382837 | 0.048378 |
| 123 | 133 | DEEWKRLRSLL | WT | 0 | 0 | 0 |
| 123 | 133 | DEEWKRLRSLL | WT | 0.5 | 1.12694 | 0.052186 |
| 123 | 133 | DEEWKRLRSLL | WT | 1 | 1.30977 | 0.11338 |
| 123 | 133 | DEEWKRLRSLL | WT | 2 | 1.566295 | 0.11742 |
| 123 | 133 | DEEWKRLRSLL | WT | 5 | 1.836991 | 0.040053 |
| 123 | 133 | DEEWKRLRSLL | WT | 30 | 2.133149 | 0.069888 |
| 123 | 133 | DEEWKRLRSLL | WT | 60 | 2.315276 | 0.036089 |
| 123 | 133 | DEEWKRLRSLL | WT | 240 | 2.811047 | 0.09361 |
| 123 | 137 | DEEWKRLRSLLSPTF | WT | 0 | 0 | 0 |
| 123 | 137 | DEEWKRLRSLLSPTF | WT | 0.5 | 1.152776 | 0.154284 |
| 123 | 137 | DEEWKRLRSLLSPTF | WT | 1 | 1.396119 | 0.153004 |
| 123 | 137 | DEEWKRLRSLLSPTF | WT | 2 | 1.762218 | 0.133286 |
| 123 | 137 | DEEWKRLRSLLSPTF | WT | 5 | 2.269155 | 0.122614 |
| 123 | 137 | DEEWKRLRSLLSPTF | WT | 30 | 3.201867 | 0.082753 |
| 123 | 137 | DEEWKRLRSLLSPTF | WT | 60 | 3.634515 | 0.068117 |
| 123 | 137 | DEEWKRLRSLLSPTF | WT | 240 | 4.373911 | 0.090555 |
| 126 | 133 | WKRLRSLL | WT | 0 | 0 | 0 |

| | | | | | | |
|---|---|---|---|---|---|---|
| 126 | 133 | WKRLRSLL | WT | 0.5 | 0.158589 | 0.059298 |
| 126 | 133 | WKRLRSLL | WT | 1 | 0.255991 | 0.065627 |
| 126 | 133 | WKRLRSLL | WT | 2 | 0.398748 | 0.056944 |
| 126 | 133 | WKRLRSLL | WT | 5 | 0.566981 | 0.070073 |
| 126 | 133 | WKRLRSLL | WT | 30 | 0.814836 | 0.061679 |
| 126 | 133 | WKRLRSLL | WT | 60 | 0.936622 | 0.088529 |
| 126 | 133 | WKRLRSLL | WT | 240 | 1.192091 | 0.115442 |
| 126 | 151 | WKRLRSLLSPTFTSGKLKEMVPIIAQ | WT | 0 | 0 | 0 |
| 126 | 151 | WKRLRSLLSPTFTSGKLKEMVPIIAQ | WT | 0.5 | 2.663435 | 0.065415 |
| 126 | 151 | WKRLRSLLSPTFTSGKLKEMVPIIAQ | WT | 1 | 3.114553 | 0.112244 |
| 126 | 151 | WKRLRSLLSPTFTSGKLKEMVPIIAQ | WT | 2 | 3.728249 | 0.081285 |
| 126 | 151 | WKRLRSLLSPTFTSGKLKEMVPIIAQ | WT | 5 | 4.548285 | 0.057682 |
| 126 | 151 | WKRLRSLLSPTFTSGKLKEMVPIIAQ | WT | 30 | 6.015619 | 0.066392 |
| 126 | 151 | WKRLRSLLSPTFTSGKLKEMVPIIAQ | WT | 60 | 6.691795 | 0.164346 |
| 126 | 151 | WKRLRSLLSPTFTSGKLKEMVPIIAQ | WT | 240 | 7.811428 | 0.118538 |
| 134 | 142 | SPTFTSGKL | WT | 0 | 0 | 0 |
| 134 | 142 | SPTFTSGKL | WT | 0.5 | 1.144231 | 0.056889 |
| 134 | 142 | SPTFTSGKL | WT | 1 | 1.340539 | 0.055914 |
| 134 | 142 | SPTFTSGKL | WT | 2 | 1.416594 | 0.064064 |
| 134 | 142 | SPTFTSGKL | WT | 5 | 1.437618 | 0.059979 |
| 134 | 142 | SPTFTSGKL | WT | 30 | 1.702483 | 0.088943 |
| 134 | 142 | SPTFTSGKL | WT | 60 | 1.726044 | 0.057596 |
| 134 | 142 | SPTFTSGKL | WT | 240 | 1.732816 | 0.0721 |
| 135 | 151 | PTFTSGKLKEMVPIIAQ | WT | 0 | 0 | 0 |
| 135 | 151 | PTFTSGKLKEMVPIIAQ | WT | 0.5 | 2.130037 | 0.073218 |
| 135 | 151 | PTFTSGKLKEMVPIIAQ | WT | 1 | 2.547309 | 0.065953 |
| 135 | 151 | PTFTSGKLKEMVPIIAQ | WT | 2 | 2.970327 | 0.034534 |
| 135 | 151 | PTFTSGKLKEMVPIIAQ | WT | 5 | 3.44233 | 0.085168 |
| 135 | 151 | PTFTSGKLKEMVPIIAQ | WT | 30 | 4.567106 | 0.016815 |
| 135 | 151 | PTFTSGKLKEMVPIIAQ | WT | 60 | 5.046031 | 0.086723 |
| 135 | 151 | PTFTSGKLKEMVPIIAQ | WT | 240 | 5.981983 | 0.045598 |
| 143 | 151 | KEMVPIIAQ | WT | 0 | 0 | 0 |
| 143 | 151 | KEMVPIIAQ | WT | 0.5 | 0.652144 | 0.012318 |
| 143 | 151 | KEMVPIIAQ | WT | 1 | 0.717022 | 0.023611 |
| 143 | 151 | KEMVPIIAQ | WT | 2 | 0.812896 | 0.029242 |
| 143 | 151 | KEMVPIIAQ | WT | 5 | 0.991709 | 0.010979 |
| 143 | 151 | KEMVPIIAQ | WT | 30 | 1.678354 | 0.032707 |
| 143 | 151 | KEMVPIIAQ | WT | 60 | 2.114544 | 0.024666 |
| 143 | 151 | KEMVPIIAQ | WT | 240 | 2.885109 | 0.026844 |
| 157 | 172 | VRNLRREAETGKPVTL | WT | 0 | 0 | 0 |
| 157 | 172 | VRNLRREAETGKPVTL | WT | 0.5 | 2.623531 | 0.042943 |
| 157 | 172 | VRNLRREAETGKPVTL | WT | 1 | 3.210939 | 0.069913 |
| 157 | 172 | VRNLRREAETGKPVTL | WT | 2 | 3.718018 | 0.069409 |
| 157 | 172 | VRNLRREAETGKPVTL | WT | 5 | 4.068166 | 0.083135 |
| 157 | 172 | VRNLRREAETGKPVTL | WT | 30 | 4.526398 | 0.049763 |

| | | | | | | |
|---|---|---|---|---|---|---|
| 157 | 172 | VRNLRREAETGKPVTL | WT | 60 | 4.580346 | 0.041844 |
| 157 | 172 | VRNLRREAETGKPVTL | WT | 240 | 4.922906 | 0.087173 |
| 173 | 178 | KDVFGA | WT | 0 | 0 | 0 |
| 173 | 178 | KDVFGA | WT | 0.5 | 0.624654 | 0.041707 |
| 173 | 178 | KDVFGA | WT | 1 | 0.826621 | 0.057444 |
| 173 | 178 | KDVFGA | WT | 2 | 0.96806 | 0.043008 |
| 173 | 178 | KDVFGA | WT | 5 | 1.221714 | 0.007116 |
| 173 | 178 | KDVFGA | WT | 30 | 1.723099 | 0.040532 |
| 173 | 178 | KDVFGA | WT | 60 | 1.798262 | 0.010374 |
| 173 | 178 | KDVFGA | WT | 240 | 1.801016 | 0.047179 |
| 173 | 179 | KDVFGAY | WT | 0 | 0 | 0 |
| 173 | 179 | KDVFGAY | WT | 0.5 | 0.616114 | 0.016321 |
| 173 | 179 | KDVFGAY | WT | 1 | 0.807904 | 0.059239 |
| 173 | 179 | KDVFGAY | WT | 2 | 0.970549 | 0.020125 |
| 173 | 179 | KDVFGAY | WT | 5 | 1.246891 | 0.040803 |
| 173 | 179 | KDVFGAY | WT | 30 | 2.050574 | 0.079472 |
| 173 | 179 | KDVFGAY | WT | 60 | 2.146827 | 0.024359 |
| 173 | 179 | KDVFGAY | WT | 240 | 2.347406 | 0.071363 |
| 173 | 181 | KDVFGAYSM | WT | 0 | 0 | 0 |
| 173 | 181 | KDVFGAYSM | WT | 0.5 | 0.962845 | 0.042517 |
| 173 | 181 | KDVFGAYSM | WT | 1 | 1.128317 | 0.059434 |
| 173 | 181 | KDVFGAYSM | WT | 2 | 1.273084 | 0.04553 |
| 173 | 181 | KDVFGAYSM | WT | 5 | 1.487582 | 0.036183 |
| 173 | 181 | KDVFGAYSM | WT | 30 | 2.210751 | 0.061093 |
| 173 | 181 | KDVFGAYSM | WT | 60 | 2.447071 | 0.09156 |
| 173 | 181 | KDVFGAYSM | WT | 240 | 2.851161 | 0.042792 |
| 177 | 182 | GAYSMD | WT | 0 | 0 | 0 |
| 177 | 182 | GAYSMD | WT | 0.5 | 0.54466 | 0.063893 |
| 177 | 182 | GAYSMD | WT | 1 | 0.536209 | 0.018141 |
| 177 | 182 | GAYSMD | WT | 2 | 0.586484 | 0.023781 |
| 177 | 182 | GAYSMD | WT | 5 | 0.626869 | 0.024885 |
| 177 | 182 | GAYSMD | WT | 30 | 0.874035 | 0.038818 |
| 177 | 182 | GAYSMD | WT | 60 | 0.998781 | 0.044861 |
| 177 | 182 | GAYSMD | WT | 240 | 1.464994 | 0.059208 |
| 182 | 189 | DVITSTSF | WT | 0 | 0 | 0 |
| 182 | 189 | DVITSTSF | WT | 0.5 | 0.368351 | 0.032215 |
| 182 | 189 | DVITSTSF | WT | 1 | 0.452032 | 0.048082 |
| 182 | 189 | DVITSTSF | WT | 2 | 0.46323 | 0.057307 |
| 182 | 189 | DVITSTSF | WT | 5 | 0.566369 | 0.033831 |
| 182 | 189 | DVITSTSF | WT | 30 | 0.892722 | 0.033566 |
| 182 | 189 | DVITSTSF | WT | 60 | 1.044047 | 0.119452 |
| 182 | 189 | DVITSTSF | WT | 240 | 1.428061 | 0.080931 |
| 183 | 189 | VITSTSF | WT | 0 | 0 | 0 |
| 183 | 189 | VITSTSF | WT | 0.5 | 0.803026 | 0.062189 |
| 183 | 189 | VITSTSF | WT | 1 | 0.902236 | 0.055522 |

| | | | | | | |
|---|---|---|---|---|---|---|
| 183 | 189 | VITSTSF | WT | 2 | 0.960825 | 0.060375 |
| 183 | 189 | VITSTSF | WT | 5 | 1.039773 | 0.019317 |
| 183 | 189 | VITSTSF | WT | 30 | 1.169448 | 0.030536 |
| 183 | 189 | VITSTSF | WT | 60 | 1.283211 | 0.121465 |
| 183 | 189 | VITSTSF | WT | 240 | 1.627978 | 0.092733 |
| 185 | 209 | TSTSFGVNIDSLNNPQDPFVENTKK | WT | 0 | 0 | 0 |
| 185 | 209 | TSTSFGVNIDSLNNPQDPFVENTKK | WT | 0.5 | 1.747121 | 0.208971 |
| 185 | 209 | TSTSFGVNIDSLNNPQDPFVENTKK | WT | 1 | 1.843971 | 0.179669 |
| 185 | 209 | TSTSFGVNIDSLNNPQDPFVENTKK | WT | 2 | 2.211377 | 0.216144 |
| 185 | 209 | TSTSFGVNIDSLNNPQDPFVENTKK | WT | 5 | 2.444162 | 0.198263 |
| 185 | 209 | TSTSFGVNIDSLNNPQDPFVENTKK | WT | 30 | 2.991939 | 0.204051 |
| 185 | 209 | TSTSFGVNIDSLNNPQDPFVENTKK | WT | 60 | 3.067595 | 0.215633 |
| 185 | 209 | TSTSFGVNIDSLNNPQDPFVENTKK | WT | 240 | 3.397545 | 0.203419 |
| 190 | 210 | GVNIDSLNNPQDPFVENTKKL | WT | 0 | 0 | 0 |
| 190 | 210 | GVNIDSLNNPQDPFVENTKKL | WT | 0.5 | 3.432544 | 0.098034 |
| 190 | 210 | GVNIDSLNNPQDPFVENTKKL | WT | 1 | 4.066843 | 0.122904 |
| 190 | 210 | GVNIDSLNNPQDPFVENTKKL | WT | 2 | 4.716044 | 0.118977 |
| 190 | 210 | GVNIDSLNNPQDPFVENTKKL | WT | 5 | 5.108762 | 0.122605 |
| 190 | 210 | GVNIDSLNNPQDPFVENTKKL | WT | 30 | 6.730416 | 0.062146 |
| 190 | 210 | GVNIDSLNNPQDPFVENTKKL | WT | 60 | 6.907955 | 0.179613 |
| 190 | 210 | GVNIDSLNNPQDPFVENTKKL | WT | 240 | 7.234699 | 0.161963 |
| 190 | 213 | GVNIDSLNNPQDPFVENTKKLLRF | WT | 0 | 0 | 0 |
| 190 | 213 | GVNIDSLNNPQDPFVENTKKLLRF | WT | 0.5 | 4.20585 | 0.153161 |
| 190 | 213 | GVNIDSLNNPQDPFVENTKKLLRF | WT | 1 | 4.867039 | 0.181658 |
| 190 | 213 | GVNIDSLNNPQDPFVENTKKLLRF | WT | 2 | 5.655455 | 0.139156 |
| 190 | 213 | GVNIDSLNNPQDPFVENTKKLLRF | WT | 5 | 6.422188 | 0.155634 |
| 190 | 213 | GVNIDSLNNPQDPFVENTKKLLRF | WT | 30 | 8.428237 | 0.179774 |
| 190 | 213 | GVNIDSLNNPQDPFVENTKKLLRF | WT | 60 | 8.896398 | 0.091004 |
| 190 | 213 | GVNIDSLNNPQDPFVENTKKLLRF | WT | 240 | 9.234595 | 0.180993 |
| 193 | 210 | IDSLNNPQDPFVENTKKL | WT | 0 | 0 | 0 |
| 193 | 210 | IDSLNNPQDPFVENTKKL | WT | 0.5 | 2.32269 | 0.098434 |
| 193 | 210 | IDSLNNPQDPFVENTKKL | WT | 1 | 2.683748 | 0.1261 |
| 193 | 210 | IDSLNNPQDPFVENTKKL | WT | 2 | 3.131675 | 0.031996 |
| 193 | 210 | IDSLNNPQDPFVENTKKL | WT | 5 | 3.547136 | 0.048599 |
| 193 | 210 | IDSLNNPQDPFVENTKKL | WT | 30 | 4.852424 | 0.056552 |
| 193 | 210 | IDSLNNPQDPFVENTKKL | WT | 60 | 5.104006 | 0.051386 |
| 193 | 210 | IDSLNNPQDPFVENTKKL | WT | 240 | 5.260825 | 0.146334 |
| 202 | 213 | PFVENTKKLLRF | WT | 0 | 0 | 0 |
| 202 | 213 | PFVENTKKLLRF | WT | 0.5 | 1.595367 | 0.050849 |
| 202 | 213 | PFVENTKKLLRF | WT | 1 | 1.944863 | 0.075117 |
| 202 | 213 | PFVENTKKLLRF | WT | 2 | 2.316322 | 0.030688 |
| 202 | 213 | PFVENTKKLLRF | WT | 5 | 2.741479 | 0.103747 |
| 202 | 213 | PFVENTKKLLRF | WT | 30 | 3.966653 | 0.08881 |
| 202 | 213 | PFVENTKKLLRF | WT | 60 | 4.453756 | 0.211576 |
| 202 | 213 | PFVENTKKLLRF | WT | 240 | 4.652111 | 0.128805 |

| | | | | | | |
|---|---|---|---|---|---|---|
| 202 | 225 | PFVENTKKLLRFDFLDPFFLSITV | WT | 0 | 0 | 0 |
| 202 | 225 | PFVENTKKLLRFDFLDPFFLSITV | WT | 0.5 | 2.620687 | 0.032956 |
| 202 | 225 | PFVENTKKLLRFDFLDPFFLSITV | WT | 1 | 2.805085 | 0.215793 |
| 202 | 225 | PFVENTKKLLRFDFLDPFFLSITV | WT | 2 | 3.012719 | 0.052604 |
| 202 | 225 | PFVENTKKLLRFDFLDPFFLSITV | WT | 5 | 2.862009 | 0.193069 |
| 202 | 225 | PFVENTKKLLRFDFLDPFFLSITV | WT | 30 | 3.350896 | 0.082672 |
| 202 | 225 | PFVENTKKLLRFDFLDPFFLSITV | WT | 60 | 3.319377 | 0.172794 |
| 202 | 225 | PFVENTKKLLRFDFLDPFFLSITV | WT | 240 | 3.48452 | 0.106365 |
| 214 | 220 | DFLDPFF | WT | 0 | 0 | 0 |
| 214 | 220 | DFLDPFF | WT | 0.5 | 0.318355 | 0.08783 |
| 214 | 220 | DFLDPFF | WT | 1 | 0.401466 | 0.110309 |
| 214 | 220 | DFLDPFF | WT | 2 | 0.42524 | 0.050832 |
| 214 | 220 | DFLDPFF | WT | 5 | 0.452253 | 0.024575 |
| 214 | 220 | DFLDPFF | WT | 30 | 0.57918 | 0.103185 |
| 214 | 220 | DFLDPFF | WT | 60 | 0.513151 | 0.018752 |
| 214 | 220 | DFLDPFF | WT | 240 | 0.750976 | 0.056953 |
| 227 | 233 | PFLIPIL | WT | 0 | 0 | 0 |
| 227 | 233 | PFLIPIL | WT | 0.5 | 0.213102 | 0.028248 |
| 227 | 233 | PFLIPIL | WT | 1 | 0.453692 | 0.072034 |
| 227 | 233 | PFLIPIL | WT | 2 | 0.402406 | 0.158201 |
| 227 | 233 | PFLIPIL | WT | 5 | 0.515622 | 0.091952 |
| 227 | 233 | PFLIPIL | WT | 30 | 0.51066 | 0.135506 |
| 227 | 233 | PFLIPIL | WT | 60 | 0.768793 | 0.024335 |
| 227 | 233 | PFLIPIL | WT | 240 | 1.609218 | 0.024335 |
| 230 | 235 | IPILEV | WT | 0 | 0 | 0 |
| 230 | 235 | IPILEV | WT | 0.5 | 0.119554 | 0.054146 |
| 230 | 235 | IPILEV | WT | 1 | 0.175856 | 0.040194 |
| 230 | 235 | IPILEV | WT | 2 | 0.269941 | 0.048854 |
| 230 | 235 | IPILEV | WT | 5 | 0.536074 | 0.026001 |
| 230 | 235 | IPILEV | WT | 30 | 1.061917 | 0.02862 |
| 230 | 235 | IPILEV | WT | 60 | 1.482054 | 0.047127 |
| 230 | 235 | IPILEV | WT | 240 | 2.098971 | 0.044432 |
| 235 | 241 | VLNICVF | WT | 0 | 0 | 0 |
| 235 | 241 | VLNICVF | WT | 0.5 | 1.123419 | 0.08312 |
| 235 | 241 | VLNICVF | WT | 1 | 1.394991 | 0.081242 |
| 235 | 241 | VLNICVF | WT | 2 | 1.721956 | 0.174841 |
| 235 | 241 | VLNICVF | WT | 5 | 2.148149 | 0.112989 |
| 235 | 241 | VLNICVF | WT | 30 | 2.83841 | 0.131753 |
| 235 | 241 | VLNICVF | WT | 60 | 2.884889 | 0.00476 |
| 235 | 241 | VLNICVF | WT | 240 | 3.097434 | 0.058492 |
| 237 | 248 | NICVFPREVTNF | WT | 0 | 0 | 0 |
| 237 | 248 | NICVFPREVTNF | WT | 0.5 | 2.711526 | 0.150256 |
| 237 | 248 | NICVFPREVTNF | WT | 1 | 3.397545 | 0.151722 |
| 237 | 248 | NICVFPREVTNF | WT | 2 | 3.688646 | 0.100637 |
| 237 | 248 | NICVFPREVTNF | WT | 5 | 3.907446 | 0.114853 |

| | | | | | | |
|---|---|---|---|---|---|---|
| 237 | 248 | NICVFPREVTNF | WT | 30 | 4.554678 | 0.164314 |
| 237 | 248 | NICVFPREVTNF | WT | 60 | 4.430971 | 0.034116 |
| 237 | 248 | NICVFPREVTNF | WT | 240 | 4.48232 | 0.109079 |
| 262 | 271 | EDTQKHRVDF | WT | 0 | 0 | 0 |
| 262 | 271 | EDTQKHRVDF | WT | 0.5 | 0.848334 | 0.021264 |
| 262 | 271 | EDTQKHRVDF | WT | 1 | 0.838679 | 0.025617 |
| 262 | 271 | EDTQKHRVDF | WT | 2 | 0.921236 | 0.056581 |
| 262 | 271 | EDTQKHRVDF | WT | 5 | 0.885109 | 0.048015 |
| 262 | 271 | EDTQKHRVDF | WT | 30 | 1.059906 | 0.059324 |
| 262 | 271 | EDTQKHRVDF | WT | 60 | 1.114012 | 0.06065 |
| 262 | 271 | EDTQKHRVDF | WT | 240 | 1.205896 | 0.052659 |
| 275 | 294 | MIDSQNSKETESHKALSDLE | WT | 0 | 0 | 0 |
| 275 | 294 | MIDSQNSKETESHKALSDLE | WT | 0.5 | 3.364848 | 0.053321 |
| 275 | 294 | MIDSQNSKETESHKALSDLE | WT | 1 | 3.625086 | 0.136037 |
| 275 | 294 | MIDSQNSKETESHKALSDLE | WT | 2 | 4.030052 | 0.087127 |
| 275 | 294 | MIDSQNSKETESHKALSDLE | WT | 5 | 4.205815 | 0.149607 |
| 275 | 294 | MIDSQNSKETESHKALSDLE | WT | 30 | 4.926837 | 0.076232 |
| 275 | 294 | MIDSQNSKETESHKALSDLE | WT | 60 | 4.932695 | 0.123331 |
| 275 | 294 | MIDSQNSKETESHKALSDLE | WT | 240 | 5.049498 | 0.058028 |
| 277 | 293 | DSQNSKETESHKALSDL | WT | 0 | 0 | 0 |
| 277 | 293 | DSQNSKETESHKALSDL | WT | 0.5 | 3.029737 | 0.065971 |
| 277 | 293 | DSQNSKETESHKALSDL | WT | 1 | 3.188209 | 0.157718 |
| 277 | 293 | DSQNSKETESHKALSDL | WT | 2 | 3.50205 | 0.082545 |
| 277 | 293 | DSQNSKETESHKALSDL | WT | 5 | 3.548774 | 0.06943 |
| 277 | 293 | DSQNSKETESHKALSDL | WT | 30 | 3.970327 | 0.042997 |
| 277 | 293 | DSQNSKETESHKALSDL | WT | 60 | 3.879716 | 0.09451 |
| 277 | 293 | DSQNSKETESHKALSDL | WT | 240 | 3.90324 | 0.092535 |
| 301 | 306 | IFIFAG | WT | 0 | 0 | 0 |
| 301 | 306 | IFIFAG | WT | 0.5 | 0.011611 | 0.071005 |
| 301 | 306 | IFIFAG | WT | 1 | -0.018071 | 0.035638 |
| 301 | 306 | IFIFAG | WT | 2 | 0.028 | 0.049569 |
| 301 | 306 | IFIFAG | WT | 5 | 0.02448 | 0.029214 |
| 301 | 306 | IFIFAG | WT | 30 | 0.196481 | 0.045352 |
| 301 | 306 | IFIFAG | WT | 60 | 0.241596 | 0.015536 |
| 301 | 306 | IFIFAG | WT | 240 | 0.587295 | 0.0347 |
| 307 | 314 | YETTSSVL | WT | 0 | 0 | 0 |
| 307 | 314 | YETTSSVL | WT | 0.5 | 0.497592 | 0.03569 |
| 307 | 314 | YETTSSVL | WT | 1 | 0.65505 | 0.028781 |
| 307 | 314 | YETTSSVL | WT | 2 | 0.773646 | 0.022909 |
| 307 | 314 | YETTSSVL | WT | 5 | 0.904149 | 0.037559 |
| 307 | 314 | YETTSSVL | WT | 30 | 1.350802 | 0.159586 |
| 307 | 314 | YETTSSVL | WT | 60 | 1.349216 | 0.080214 |
| 307 | 314 | YETTSSVL | WT | 240 | 1.585206 | 0.102979 |
| 307 | 316 | YETTSSVLSF | WT | 0 | 0 | 0 |
| 307 | 316 | YETTSSVLSF | WT | 0.5 | 0.40548 | 0.039938 |

| | | | | | | |
|---|---|---|---|---|---|---|
| 307 | 316 | YETTSSVLSF | WT | 1 | 0.487248 | 0.02664 |
| 307 | 316 | YETTSSVLSF | WT | 2 | 0.637978 | 0.03203 |
| 307 | 316 | YETTSSVLSF | WT | 5 | 0.805677 | 0.01885 |
| 307 | 316 | YETTSSVLSF | WT | 30 | 1.288376 | 0.04303 |
| 307 | 316 | YETTSSVLSF | WT | 60 | 1.479077 | 0.066883 |
| 307 | 316 | YETTSSVLSF | WT | 240 | 1.92626 | 0.072626 |
| 308 | 316 | ETTSSVLSF | WT | 0 | 0 | 0 |
| 308 | 316 | ETTSSVLSF | WT | 0.5 | 0.226188 | 0.015588 |
| 308 | 316 | ETTSSVLSF | WT | 1 | 0.315655 | 0.023161 |
| 308 | 316 | ETTSSVLSF | WT | 2 | 0.448194 | 0.016959 |
| 308 | 316 | ETTSSVLSF | WT | 5 | 0.592729 | 0.03507 |
| 308 | 316 | ETTSSVLSF | WT | 30 | 0.949799 | 0.044438 |
| 308 | 316 | ETTSSVLSF | WT | 60 | 1.138775 | 0.046693 |
| 308 | 316 | ETTSSVLSF | WT | 240 | 1.560554 | 0.045391 |
| 317 | 333 | IMYELATHPDVQQKLQE | WT | 0 | 0 | 0 |
| 317 | 333 | IMYELATHPDVQQKLQE | WT | 0.5 | 0.10389 | 0.03166 |
| 317 | 333 | IMYELATHPDVQQKLQE | WT | 1 | 0.152361 | 0.061127 |
| 317 | 333 | IMYELATHPDVQQKLQE | WT | 2 | 0.224064 | 0.02763 |
| 317 | 333 | IMYELATHPDVQQKLQE | WT | 5 | 0.370585 | 0.025103 |
| 317 | 333 | IMYELATHPDVQQKLQE | WT | 30 | 0.827423 | 0.018996 |
| 317 | 333 | IMYELATHPDVQQKLQE | WT | 60 | 0.849061 | 0.043469 |
| 317 | 333 | IMYELATHPDVQQKLQE | WT | 240 | 1.21813 | 0.058067 |
| 320 | 333 | ELATHPDVQQKLQE | WT | 0 | 0 | 0 |
| 320 | 333 | ELATHPDVQQKLQE | WT | 0.5 | 0.163185 | 0.102684 |
| 320 | 333 | ELATHPDVQQKLQE | WT | 1 | 0.219159 | 0.090538 |
| 320 | 333 | ELATHPDVQQKLQE | WT | 2 | 0.278502 | 0.08326 |
| 320 | 333 | ELATHPDVQQKLQE | WT | 5 | 0.436909 | 0.106221 |
| 320 | 333 | ELATHPDVQQKLQE | WT | 30 | 0.79915 | 0.069196 |
| 320 | 333 | ELATHPDVQQKLQE | WT | 60 | 0.894425 | 0.083493 |
| 320 | 333 | ELATHPDVQQKLQE | WT | 240 | 1.145383 | 0.091127 |
| 337 | 347 | AVLPNKAPPTY | WT | 0 | 0 | 0 |
| 337 | 347 | AVLPNKAPPTY | WT | 0.5 | 1.216493 | 0.028492 |
| 337 | 347 | AVLPNKAPPTY | WT | 1 | 1.279786 | 0.124199 |
| 337 | 347 | AVLPNKAPPTY | WT | 2 | 1.526244 | 0.075034 |
| 337 | 347 | AVLPNKAPPTY | WT | 5 | 1.793313 | 0.079975 |
| 337 | 347 | AVLPNKAPPTY | WT | 30 | 2.329019 | 0.040273 |
| 337 | 347 | AVLPNKAPPTY | WT | 60 | 2.368831 | 0.019762 |
| 337 | 347 | AVLPNKAPPTY | WT | 240 | 2.38695 | 0.025343 |
| 337 | 349 | AVLPNKAPPTYDT | WT | 0 | 0 | 0 |
| 337 | 349 | AVLPNKAPPTYDT | WT | 0.5 | 2.364323 | 0.059731 |
| 337 | 349 | AVLPNKAPPTYDT | WT | 1 | 2.607805 | 0.077885 |
| 337 | 349 | AVLPNKAPPTYDT | WT | 2 | 2.843949 | 0.084387 |
| 337 | 349 | AVLPNKAPPTYDT | WT | 5 | 3.061756 | 0.041196 |
| 337 | 349 | AVLPNKAPPTYDT | WT | 30 | 3.747301 | 0.099086 |
| 337 | 349 | AVLPNKAPPTYDT | WT | 60 | 3.763214 | 0.034255 |

| | | | | | | |
|---|---|---|---|---|---|---|
| 337 | 349 | AVLPNKAPPTYDT | WT | 240 | 3.805368 | 0.081076 |
| 337 | 351 | AVLPNKAPPTYDTVL | WT | 0 | 0 | 0 |
| 337 | 351 | AVLPNKAPPTYDTVL | WT | 0.5 | 2.200012 | 0.048508 |
| 337 | 351 | AVLPNKAPPTYDTVL | WT | 1 | 2.594051 | 0.074893 |
| 337 | 351 | AVLPNKAPPTYDTVL | WT | 2 | 3.001975 | 0.068491 |
| 337 | 351 | AVLPNKAPPTYDTVL | WT | 5 | 3.476332 | 0.080797 |
| 337 | 351 | AVLPNKAPPTYDTVL | WT | 30 | 4.604474 | 0.060233 |
| 337 | 351 | AVLPNKAPPTYDTVL | WT | 60 | 4.716582 | 0.124898 |
| 337 | 351 | AVLPNKAPPTYDTVL | WT | 240 | 5.076429 | 0.102684 |
| 357 | 362 | DMVVNE | WT | 0 | 0 | 0 |
| 357 | 362 | DMVVNE | WT | 0.5 | 0.232484 | 0.033968 |
| 357 | 362 | DMVVNE | WT | 1 | 0.218137 | 0.023027 |
| 357 | 362 | DMVVNE | WT | 2 | 0.224418 | 0.026253 |
| 357 | 362 | DMVVNE | WT | 5 | 0.213394 | 0.009524 |
| 357 | 362 | DMVVNE | WT | 30 | 0.238513 | 0.017133 |
| 357 | 362 | DMVVNE | WT | 60 | 0.241902 | 0.012472 |
| 357 | 362 | DMVVNE | WT | 240 | 0.240985 | 0.016873 |
| 359 | 364 | VVNETL | WT | 0 | 0 | 0 |
| 359 | 364 | VVNETL | WT | 0.5 | 0.137376 | 0.050386 |
| 359 | 364 | VVNETL | WT | 1 | 0.104066 | 0.0688 |
| 359 | 364 | VVNETL | WT | 2 | 0.083651 | 0.018821 |
| 359 | 364 | VVNETL | WT | 5 | 0.066862 | 0.041534 |
| 359 | 364 | VVNETL | WT | 30 | 0.061391 | 0.032949 |
| 359 | 364 | VVNETL | WT | 60 | 0.146827 | 0.0803 |
| 359 | 364 | VVNETL | WT | 240 | 0.090743 | 0.030535 |
| 364 | 370 | LRLFPIA | WT | 0 | 0 | 0 |
| 364 | 370 | LRLFPIA | WT | 0.5 | 0.077762 | 0.063913 |
| 364 | 370 | LRLFPIA | WT | 1 | 0.053908 | 0.026308 |
| 364 | 370 | LRLFPIA | WT | 2 | 0.101416 | 0.02963 |
| 364 | 370 | LRLFPIA | WT | 5 | 0.129686 | 0.031396 |
| 364 | 370 | LRLFPIA | WT | 30 | 0.287215 | 0.041335 |
| 364 | 370 | LRLFPIA | WT | 60 | 0.390719 | 0.031153 |
| 364 | 370 | LRLFPIA | WT | 240 | 0.518994 | 0.02368 |
| 372 | 385 | RLERVCKKDVEING | WT | 0 | 0 | 0 |
| 372 | 385 | RLERVCKKDVEING | WT | 0.5 | 1.055907 | 0.044983 |
| 372 | 385 | RLERVCKKDVEING | WT | 1 | 1.21087 | 0.066002 |
| 372 | 385 | RLERVCKKDVEING | WT | 2 | 1.427669 | 0.041625 |
| 372 | 385 | RLERVCKKDVEING | WT | 5 | 1.492729 | 0.053503 |
| 372 | 385 | RLERVCKKDVEING | WT | 30 | 1.788426 | 0.041723 |
| 372 | 385 | RLERVCKKDVEING | WT | 60 | 1.840746 | 0.042084 |
| 372 | 385 | RLERVCKKDVEING | WT | 240 | 2.094063 | 0.067299 |
| 374 | 387 | ERVCKKDVEINGMF | WT | 0 | 0 | 0 |
| 374 | 387 | ERVCKKDVEINGMF | WT | 0.5 | 1.406507 | 0.051018 |
| 374 | 387 | ERVCKKDVEINGMF | WT | 1 | 1.642762 | 0.07919 |
| 374 | 387 | ERVCKKDVEINGMF | WT | 2 | 1.817926 | 0.03475 |

| | | | | | | |
|---|---|---|---|---|---|---|
| 374 | 387 | ERVCKKDVEINGMF | WT | 5 | 1.926936 | 0.038789 |
| 374 | 387 | ERVCKKDVEINGMF | WT | 30 | 2.196258 | 0.059836 |
| 374 | 387 | ERVCKKDVEINGMF | WT | 60 | 2.370593 | 0.084678 |
| 374 | 387 | ERVCKKDVEINGMF | WT | 240 | 2.685121 | 0.056199 |
| 387 | 393 | FIPKGVV | WT | 0 | 0 | 0 |
| 387 | 393 | FIPKGVV | WT | 0.5 | 0.324926 | 0.043223 |
| 387 | 393 | FIPKGVV | WT | 1 | 0.322565 | 0.041763 |
| 387 | 393 | FIPKGVV | WT | 2 | 0.348631 | 0.053248 |
| 387 | 393 | FIPKGVV | WT | 5 | 0.385761 | 0.045994 |
| 387 | 393 | FIPKGVV | WT | 30 | 0.486286 | 0.054754 |
| 387 | 393 | FIPKGVV | WT | 60 | 0.540779 | 0.040258 |
| 387 | 393 | FIPKGVV | WT | 240 | 0.622393 | 0.033024 |
| 388 | 393 | IPKGVV | WT | 0 | 0 | 0 |
| 388 | 393 | IPKGVV | WT | 0.5 | 0.220908 | 0.031584 |
| 388 | 393 | IPKGVV | WT | 1 | 0.268613 | 0.035092 |
| 388 | 393 | IPKGVV | WT | 2 | 0.282741 | 0.023521 |
| 388 | 393 | IPKGVV | WT | 5 | 0.35703 | 0.0189 |
| 388 | 393 | IPKGVV | WT | 30 | 0.461565 | 0.038956 |
| 388 | 393 | IPKGVV | WT | 60 | 0.512716 | 0.018328 |
| 388 | 393 | IPKGVV | WT | 240 | 0.59605 | 0.026889 |
| 394 | 407 | VMIPSYALHRDPKY | WT | 0 | 0 | 0 |
| 394 | 407 | VMIPSYALHRDPKY | WT | 0.5 | 0.579737 | 0.050304 |
| 394 | 407 | VMIPSYALHRDPKY | WT | 1 | 0.593187 | 0.020077 |
| 394 | 407 | VMIPSYALHRDPKY | WT | 2 | 0.645994 | 0.049891 |
| 394 | 407 | VMIPSYALHRDPKY | WT | 5 | 0.746208 | 0.057914 |
| 394 | 407 | VMIPSYALHRDPKY | WT | 30 | 0.80989 | 0.073853 |
| 394 | 407 | VMIPSYALHRDPKY | WT | 60 | 0.987136 | 0.007908 |
| 394 | 407 | VMIPSYALHRDPKY | WT | 240 | 1.442154 | 0.036365 |
| 396 | 401 | IPSYAL | WT | 0 | 0 | 0 |
| 396 | 401 | IPSYAL | WT | 0.5 | 0.397488 | 0.053174 |
| 396 | 401 | IPSYAL | WT | 1 | 0.417094 | 0.039857 |
| 396 | 401 | IPSYAL | WT | 2 | 0.416878 | 0.03609 |
| 396 | 401 | IPSYAL | WT | 5 | 0.472234 | 0.019652 |
| 396 | 401 | IPSYAL | WT | 30 | 0.569555 | 0.03043 |
| 396 | 401 | IPSYAL | WT | 60 | 0.545685 | 0.01637 |
| 396 | 401 | IPSYAL | WT | 240 | 0.630862 | 0.033108 |
| 396 | 407 | IPSYALHRDPKY | WT | 0 | 0 | 0 |
| 396 | 407 | IPSYALHRDPKY | WT | 0.5 | 0.48339 | 0.054975 |
| 396 | 407 | IPSYALHRDPKY | WT | 1 | 0.568469 | 0.021669 |
| 396 | 407 | IPSYALHRDPKY | WT | 2 | 0.672727 | 0.032657 |
| 396 | 407 | IPSYALHRDPKY | WT | 5 | 0.745242 | 0.029525 |
| 396 | 407 | IPSYALHRDPKY | WT | 30 | 0.921954 | 0.042002 |
| 396 | 407 | IPSYALHRDPKY | WT | 60 | 0.981628 | 0.022097 |
| 396 | 407 | IPSYALHRDPKY | WT | 240 | 1.314201 | 0.034837 |
| 400 | 414 | ALHRDPKYWTEPEKF | WT | 0 | 0 | 0 |

| | | | | | | |
|---|---|---|---|---|---|---|
| 400 | 414 | ALHRDPKYWTEPEKF | WT | 0.5 | 2.510136 | 0.048291 |
| 400 | 414 | ALHRDPKYWTEPEKF | WT | 1 | 2.615486 | 0.103717 |
| 400 | 414 | ALHRDPKYWTEPEKF | WT | 2 | 2.727659 | 0.065192 |
| 400 | 414 | ALHRDPKYWTEPEKF | WT | 5 | 2.852202 | 0.153014 |
| 400 | 414 | ALHRDPKYWTEPEKF | WT | 30 | 2.880337 | 0.043751 |
| 400 | 414 | ALHRDPKYWTEPEKF | WT | 60 | 2.941318 | 0.102797 |
| 400 | 414 | ALHRDPKYWTEPEKF | WT | 240 | 3.249393 | 0.084337 |
| 408 | 428 | WTEPEKFLPERFSKKNKDNID | WT | 0 | 0 | 0 |
| 408 | 428 | WTEPEKFLPERFSKKNKDNID | WT | 0.5 | 2.409277 | 0.135887 |
| 408 | 428 | WTEPEKFLPERFSKKNKDNID | WT | 1 | 2.524759 | 0.202456 |
| 408 | 428 | WTEPEKFLPERFSKKNKDNID | WT | 2 | 2.891592 | 0.2944 |
| 408 | 428 | WTEPEKFLPERFSKKNKDNID | WT | 5 | 2.476584 | 0.199231 |
| 408 | 428 | WTEPEKFLPERFSKKNKDNID | WT | 30 | 2.711192 | 0.028423 |
| 408 | 428 | WTEPEKFLPERFSKKNKDNID | WT | 60 | 2.671236 | 0.158514 |
| 408 | 428 | WTEPEKFLPERFSKKNKDNID | WT | 240 | 2.706405 | 0.14602 |
| 420 | 444 | SKKNKDNIDPYIYTPFGSGPRNCIG | WT | 0 | 0 | 0 |
| 420 | 444 | SKKNKDNIDPYIYTPFGSGPRNCIG | WT | 0.5 | 2.161745 | 0.081679 |
| 420 | 444 | SKKNKDNIDPYIYTPFGSGPRNCIG | WT | 1 | 2.541107 | 0.099411 |
| 420 | 444 | SKKNKDNIDPYIYTPFGSGPRNCIG | WT | 2 | 2.813967 | 0.104497 |
| 420 | 444 | SKKNKDNIDPYIYTPFGSGPRNCIG | WT | 5 | 3.078602 | 0.136408 |
| 420 | 444 | SKKNKDNIDPYIYTPFGSGPRNCIG | WT | 30 | 3.937071 | 0.017236 |
| 420 | 444 | SKKNKDNIDPYIYTPFGSGPRNCIG | WT | 60 | 4.151412 | 0.166726 |
| 420 | 444 | SKKNKDNIDPYIYTPFGSGPRNCIG | WT | 240 | 4.746206 | 0.156648 |
| 429 | 444 | PYIYTPFGSGPRNCIG | WT | 0 | 0 | 0 |
| 429 | 444 | PYIYTPFGSGPRNCIG | WT | 0.5 | 1.088414 | 0.175226 |
| 429 | 444 | PYIYTPFGSGPRNCIG | WT | 1 | 1.195436 | 0.181211 |
| 429 | 444 | PYIYTPFGSGPRNCIG | WT | 2 | 1.376475 | 0.122619 |
| 429 | 444 | PYIYTPFGSGPRNCIG | WT | 5 | 1.690438 | 0.120153 |
| 429 | 444 | PYIYTPFGSGPRNCIG | WT | 30 | 2.322116 | 0.173494 |
| 429 | 444 | PYIYTPFGSGPRNCIG | WT | 60 | 2.605061 | 0.155105 |
| 429 | 444 | PYIYTPFGSGPRNCIG | WT | 240 | 3.223097 | 0.188388 |
| 445 | 452 | MRFALMNM | WT | 0 | 0 | 0 |
| 445 | 452 | MRFALMNM | WT | 0.5 | 0.18641 | 0.034538 |
| 445 | 452 | MRFALMNM | WT | 1 | 0.286352 | 0.017409 |
| 445 | 452 | MRFALMNM | WT | 2 | 0.394301 | 0.057972 |
| 445 | 452 | MRFALMNM | WT | 5 | 0.418337 | 0.04873 |
| 445 | 452 | MRFALMNM | WT | 30 | 0.527057 | 0.023364 |
| 445 | 452 | MRFALMNM | WT | 60 | 0.537596 | 0.028658 |
| 445 | 452 | MRFALMNM | WT | 240 | 0.726271 | 0.071694 |
| 450 | 456 | MNMKLAL | WT | 0 | 0 | 0 |
| 450 | 456 | MNMKLAL | WT | 0.5 | 0.174744 | 0.06528 |
| 450 | 456 | MNMKLAL | WT | 1 | 0.105409 | 0.10065 |
| 450 | 456 | MNMKLAL | WT | 2 | 0.074876 | 0.03697 |
| 450 | 456 | MNMKLAL | WT | 5 | 0.06959 | 0.051383 |
| 450 | 456 | MNMKLAL | WT | 30 | 0.193575 | 0.0566 |

| | | | | | | |
|---|---|---|---|---|---|---|
| 450 | 456 | MNMKLAL | WT | 60 | 0.087454 | 0.097738 |
| 450 | 456 | MNMKLAL | WT | 240 | 0.226709 | 0.128965 |
| 455 | 463 | ALIRVLQNF | WT | 0 | 0 | 0 |
| 455 | 463 | ALIRVLQNF | WT | 0.5 | 0.11112 | 0.03943 |
| 455 | 463 | ALIRVLQNF | WT | 1 | 0.139042 | 0.023866 |
| 455 | 463 | ALIRVLQNF | WT | 2 | 0.185356 | 0.060574 |
| 455 | 463 | ALIRVLQNF | WT | 5 | 0.202304 | 0.036232 |
| 455 | 463 | ALIRVLQNF | WT | 30 | 0.34756 | 0.02689 |
| 455 | 463 | ALIRVLQNF | WT | 60 | 0.409235 | 0.023219 |
| 455 | 463 | ALIRVLQNF | WT | 240 | 0.61044 | 0.072221 |
| 457 | 465 | IRVLQNFSF | WT | 0 | 0 | 0 |
| 457 | 465 | IRVLQNFSF | WT | 0.5 | 0.174054 | 0.046191 |
| 457 | 465 | IRVLQNFSF | WT | 1 | 0.191938 | 0.028757 |
| 457 | 465 | IRVLQNFSF | WT | 2 | 0.244995 | 0.041517 |
| 457 | 465 | IRVLQNFSF | WT | 5 | 0.336319 | 0.026652 |
| 457 | 465 | IRVLQNFSF | WT | 30 | 0.578721 | 0.019174 |
| 457 | 465 | IRVLQNFSF | WT | 60 | 0.718539 | 0.032332 |
| 457 | 465 | IRVLQNFSF | WT | 240 | 1.115121 | 0.047304 |
| 464 | 477 | SFKPCKETQIPLKL | WT | 0 | 0 | 0 |
| 464 | 477 | SFKPCKETQIPLKL | WT | 0.5 | 2.998562 | 0.078043 |
| 464 | 477 | SFKPCKETQIPLKL | WT | 1 | 3.186661 | 0.102191 |
| 464 | 477 | SFKPCKETQIPLKL | WT | 2 | 3.399694 | 0.03668 |
| 464 | 477 | SFKPCKETQIPLKL | WT | 5 | 3.465326 | 0.098996 |
| 464 | 477 | SFKPCKETQIPLKL | WT | 30 | 3.83654 | 0.096322 |
| 464 | 477 | SFKPCKETQIPLKL | WT | 60 | 3.952768 | 0.123676 |
| 464 | 477 | SFKPCKETQIPLKL | WT | 240 | 3.784958 | 0.075977 |
| 464 | 479 | SFKPCKETQIPLKLSL | WT | 0 | 0 | 0 |
| 464 | 479 | SFKPCKETQIPLKLSL | WT | 0.5 | 3.99117 | 0.100423 |
| 464 | 479 | SFKPCKETQIPLKLSL | WT | 1 | 4.3165 | 0.127968 |
| 464 | 479 | SFKPCKETQIPLKLSL | WT | 2 | 4.567933 | 0.089892 |
| 464 | 479 | SFKPCKETQIPLKLSL | WT | 5 | 4.543018 | 0.193126 |
| 464 | 479 | SFKPCKETQIPLKLSL | WT | 30 | 4.924918 | 0.031426 |
| 464 | 479 | SFKPCKETQIPLKLSL | WT | 60 | 4.882702 | 0.190728 |
| 464 | 479 | SFKPCKETQIPLKLSL | WT | 240 | 4.895189 | 0.150771 |
| 464 | 491 | SFKPCKETQIPLKLSLGGLLQPEKP | WT | 0 | 0 | 0 |
| 464 | 491 | SFKPCKETQIPLKLSLGGLLQPEKP | WT | 0.5 | 6.180708 | 0.087459 |
| 464 | 491 | SFKPCKETQIPLKLSLGGLLQPEKP | WT | 1 | 6.83061 | 0.122002 |
| 464 | 491 | SFKPCKETQIPLKLSLGGLLQPEKP | WT | 2 | 7.504459 | 0.055678 |
| 464 | 491 | SFKPCKETQIPLKLSLGGLLQPEKP | WT | 5 | 7.89599 | 0.193945 |
| 464 | 491 | SFKPCKETQIPLKLSLGGLLQPEKP | WT | 30 | 8.893352 | 0.071292 |
| 464 | 491 | SFKPCKETQIPLKLSLGGLLQPEKP | WT | 60 | 9.081659 | 0.182847 |
| 464 | 491 | SFKPCKETQIPLKLSLGGLLQPEKP | WT | 240 | 9.255642 | 0.158928 |
| 471 | 477 | TQIPLKL | WT | 0 | 0 | 0 |
| 471 | 477 | TQIPLKL | WT | 0.5 | 2.675564 | 0.046618 |
| 471 | 477 | TQIPLKL | WT | 1 | 2.798798 | 0.066025 |

| | | | | | | |
|---|---|---|---|---|---|---|
| 471 | 477 | TQIPLKL | WT | 2 | 2.894955 | 0.093012 |
| 471 | 477 | TQIPLKL | WT | 5 | 2.857614 | 0.055565 |
| 471 | 477 | TQIPLKL | WT | 30 | 2.871971 | 0.165942 |
| 471 | 477 | TQIPLKL | WT | 60 | 2.915935 | 0.047649 |
| 471 | 477 | TQIPLKL | WT | 240 | 2.919089 | 0.058136 |
| 478 | 491 | SLGGLLQPEKPVVL | WT | 0 | 0 | 0 |
| 478 | 491 | SLGGLLQPEKPVVL | WT | 0.5 | 3.076216 | 0.050013 |
| 478 | 491 | SLGGLLQPEKPVVL | WT | 1 | 3.556918 | 0.034746 |
| 478 | 491 | SLGGLLQPEKPVVL | WT | 2 | 4.060428 | 0.060372 |
| 478 | 491 | SLGGLLQPEKPVVL | WT | 5 | 4.328249 | 0.112176 |
| 478 | 491 | SLGGLLQPEKPVVL | WT | 30 | 5.153987 | 0.072639 |
| 478 | 491 | SLGGLLQPEKPVVL | WT | 60 | 5.240485 | 0.103898 |
| 478 | 491 | SLGGLLQPEKPVVL | WT | 240 | 5.454744 | 0.078604 |
| 480 | 491 | GGLLQPEKPVVL | WT | 0 | 0 | 0 |
| 480 | 491 | GGLLQPEKPVVL | WT | 0.5 | 2.254753 | 0.034737 |
| 480 | 491 | GGLLQPEKPVVL | WT | 1 | 2.621439 | 0.092948 |
| 480 | 491 | GGLLQPEKPVVL | WT | 2 | 2.947375 | 0.068902 |
| 480 | 491 | GGLLQPEKPVVL | WT | 5 | 3.291881 | 0.10747 |
| 480 | 491 | GGLLQPEKPVVL | WT | 30 | 3.996846 | 0.049707 |
| 480 | 491 | GGLLQPEKPVVL | WT | 60 | 4.158267 | 0.10324 |
| 480 | 491 | GGLLQPEKPVVL | WT | 240 | 4.301888 | 0.055647 |
| 485 | 491 | PEKPVVL | WT | 0 | 0 | 0 |
| 485 | 491 | PEKPVVL | WT | 0.5 | 1.436924 | 0.042954 |
| 485 | 491 | PEKPVVL | WT | 1 | 1.653079 | 0.026989 |
| 485 | 491 | PEKPVVL | WT | 2 | 1.874679 | 0.019166 |
| 485 | 491 | PEKPVVL | WT | 5 | 2.006098 | 0.061767 |
| 485 | 491 | PEKPVVL | WT | 30 | 2.4269 | 0.020045 |
| 485 | 491 | PEKPVVL | WT | 60 | 2.500689 | 0.050276 |
| 485 | 491 | PEKPVVL | WT | 240 | 2.583039 | 0.042018 |
| 492 | 507 | KVESRDGTVSGAHHHH | WT | 0 | 0 | 0 |
| 492 | 507 | KVESRDGTVSGAHHHH | WT | 0.5 | 1.245708 | 0.053611 |
| 492 | 507 | KVESRDGTVSGAHHHH | WT | 1 | 1.317352 | 0.127103 |
| 492 | 507 | KVESRDGTVSGAHHHH | WT | 2 | 1.314784 | 0.077977 |
| 492 | 507 | KVESRDGTVSGAHHHH | WT | 5 | 1.108957 | 0.167699 |
| 492 | 507 | KVESRDGTVSGAHHHH | WT | 30 | 1.227316 | 0.057315 |
| 492 | 507 | KVESRDGTVSGAHHHH | WT | 60 | 1.148085 | 0.068809 |
| 492 | 507 | KVESRDGTVSGAHHHH | WT | 240 | 1.272796 | 0.100602 |
